# A personalized multi-platform assessment of somatic mosaicism in the human frontal cortex

**DOI:** 10.1101/2024.12.18.629274

**Authors:** Weichen Zhou, Camille Mumm, Yanming Gan, Jessica A. Switzenberg, Jinhao Wang, Paulo De Oliveira, Kunal Kathuria, Steven J. Losh, Brandt Bessell, Torrin L. McDonald, Michael J. McConnell, Alan P. Boyle, Ryan E. Mills

## Abstract

Somatic mutations in individual cells create genomic mosaicism, influencing genetic disorders and cancers. While clonal mutations in cancers are well-studied, rarer somatic variants in normal tissues remain poorly characterized. This study systematically evaluates detection methods using a personalized donor-specific assembly (DSA) from a neurotypical individual’s dorsolateral prefrontal cortex assessed with Oxford Nanopore, NovaSeq, linked-read sequencing, Cas9-targeted long-read sequencing (TEnCATS), and single-neuron MALBAC amplification. The haplotype-resolved DSA improved cross-platform analysis, dramatically increasing phasing rates. Germline SNVs, structural variations (SVs), and transposable elements (TEs) were recalled with 99.4%–99.7% accuracy in bulk tissue, and phased haplotype analysis reduced false positives by 15.4%–75.1% for putative somatic candidates. Long-read single-neuron sequencing detected nine somatic SV candidates, demonstrating enhanced sensitivity for rare variants, while TEnCATS identified eight low-frequency somatic TE candidates. These findings highlight advanced methodologies for precise somatic variant detection, critical for understanding mosaicism’s role in health and disease.

## Main

Human genomes harbor significant variation both between and within individuals. Numerous studies have explored inherited variation across human populations and linked various germline polymorphisms to traits and disease susceptibility^1–7^. Genomic sequences can also vary within an individual due to post-zygotic mutations, leading to somatic mosaicism^8^. As early as 1929, it was recognized that cancers frequently possess abnormal karyotypes and somatic mutations^9,10^. Since then, several studies have identified driver genes and a wide range of somatic mutations across multiple cancer types^11–15^, including single nucleotide variants (SNVs), copy number alterations or variants (CNVs), and structural variations (SVs), that shape tumor evolution and heterogeneity^11,16,17^.

In addition to cancer genomics, somatic mosaicism has been observed throughout the human body, occurring at variable frequencies ranging from individual cells to entire tissues and across different developmental stages^18–24^. In the 1970s, somatic gene rearrangement was found in healthy human tissues to create functional diversity of immunoglobulin and T-cell receptor genes^25^. The development of cytogenetic techniques, including karyotyping and G-banding, allowed researchers to observe large-scale chromosomal abnormalities in human cells^26,27^, such as mosaic Turner syndrome^28^ and Down syndrome^29^. This is particularly true in the human brain, where neural progenitor and cortical neurons have been shown to harbor extensive tissue -specific somatic mutations, including SNVs^30–32^, transposable elements (TEs)^33–35^, and large SVs^21,36^. Since neurons are among the longest-lived cells in the body, the accumulation of somatic mutations within neural progenitors or postmitotic neurons could influence neuronal development and diversity, potentially contributing to the etiology of numerous neuropsychiatric disorders^37–41^.

The expansion of high-throughput techniques, e.g. next-generation sequencing technology, has significantly enhanced the ability to detect smaller genetic changes with much higher resolution^42^. In oncogenomics, tumor and matched normal cell pairs have been used along with targeted exome sequencing (WES) or whole genome sequencing (WGS) to identify somatic variation in cancerous tissues^42,43^. However, the natural accumulation of low-frequency somatic mutations across different cell populations in healthy tissue makes it difficult to distinguish true mutations from background noise without streamlined analysis techniques^44,45^. With the help of single-cell and higher-coverage bulk DNA sequencing approaches, we have begun to explore the extent of somatic mutations within individual tissues, though there still remain many challenges^19,46^. For example, while short-read whole genome sequencing can be highly accurate in detecting somatic point mutations from bulk tissues, such as SNVs^30,45^, it faces limitations due to the repetitive nature of large portions of the human genome which are less accessible to short reads^47–49^. These obstacles are compounded when investigating larger mutations, e.g. CNVs. Although large somatic deletions and duplications can be identified from whole-genome-amplified (WGA) single cells^21,36,50^, they are often restricted to uniquely mapped regions of the genome and the associated breakpoints are typically imprecise^48,49^. Furthermore, inaccuracies can arise due to improperly aligned reads and, when compounded with amplification bias across different alleles, can lead to additional complications^21,36,45^. The same holds true when examining somatic TEs, where despite the use of whole-genome and targeted approaches, their repetitive nature exacerbates the challenges for accurate detection and characterization^3,35,51^.

Long-read sequencing has emerged as a powerful tool to overcome the limitations associated with short-read sequencing, particularly in detecting larger genetic variants within complex genomic regions, such as SVs and TEs^3,49,52–54^. In the era of genome assembly^55,56^, long-read technologies have demonstrated the capability to generate megabase-scale phase blocks and *de novo* diploid contigs^57–60^ that provide essential haplotype information for phasing and significantly enhance genetic variation calling^61–63^. Notably, in the discovery of low-frequency somatic mosaicism, phasing information can substantially improve variant detection by reducing false positive signals^21^. The inclusion of a donor-specific assembly (DSA) has the potential to offer reference-free insights, further aiding in the discovery of somatic variants^45^.

Here we present a systematic investigation of somatic mosaicism discovery in a human dorsolateral prefrontal cortex (DLPFC) using multiple sequencing platforms and computational tools (**Fig. 1**). We employed Oxford Nanopore Technologies (ONT) long-read sequencing, TE nanopore Cas9-Targeted Sequencing (TEnCATS), 10x Genomics linked-read sequencing, and Illumina NovaSeq WGS for characterizing somatic mosaicism in bulk tissue and single neurons. Additionally, we developed experimental protocols and computational packages for ONT long-read sequencing and somatic variant calling in single cells. By constructing a diploid genome assembly and leveraging phase block information, we assessed the capabilities of different technologies to detect germline variants and somatic mosaicism in bulk tissue and single cells. Our prototype analysis spans from personalized assembly to single-cell somatic variant calling, providing a comprehensive *ab initio ad finem* approach and resource in non-diseased human tissue.

**Figure 1.**
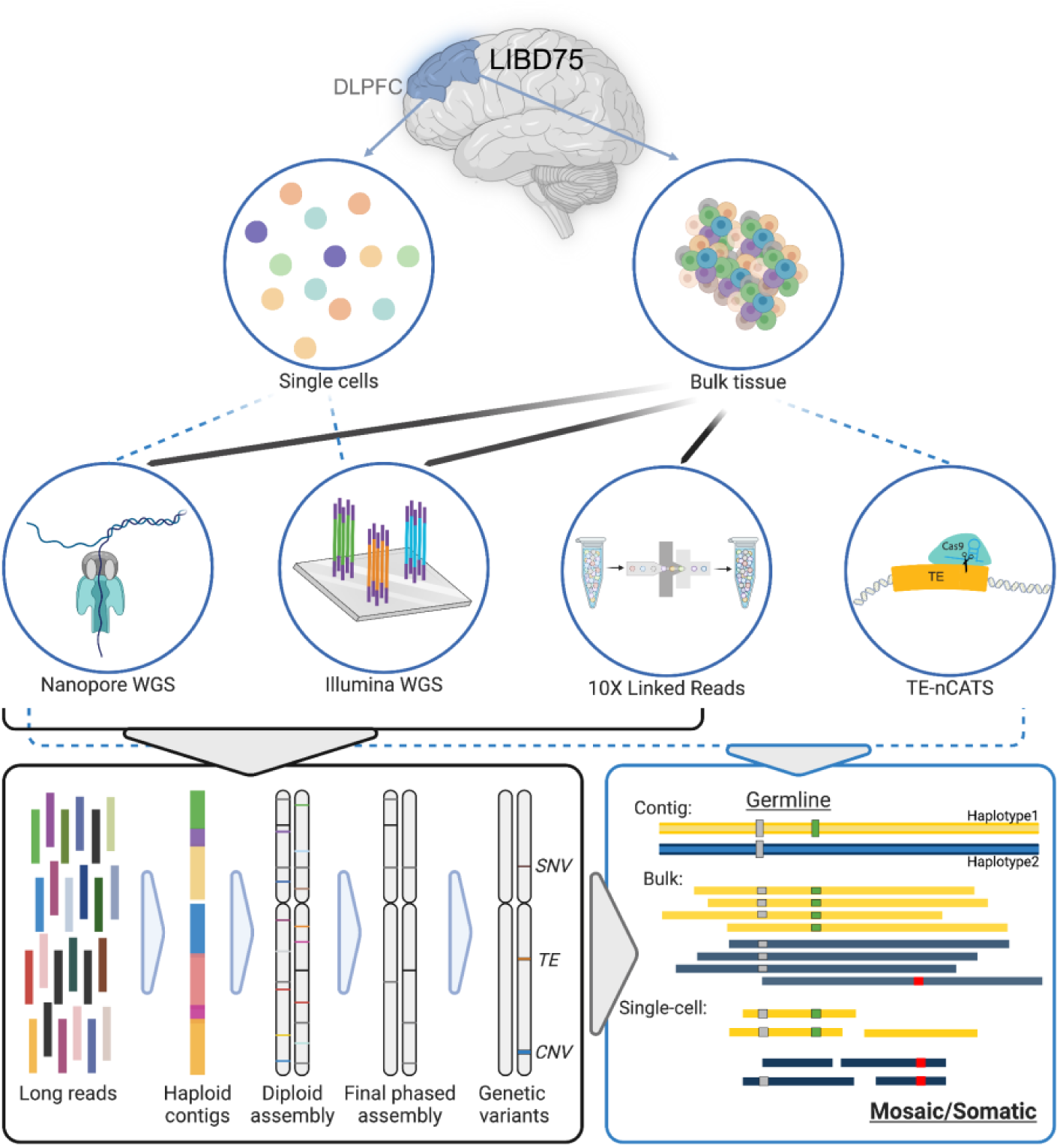
Diagram of multi-platform DNA sequencing data generation for the LIBD75 frontal cortex. Black arrows correspond to relevant methods and data used for genome assembly, while blue arrows and dotted lines indicate methods and data used for variant calling. DLPFC, dorsolateral prefrontal cortex.

## Results

### Multi-platform sequencing of a donor dorsolateral prefrontal cortex

We obtained tissue from the DLPFC of a post-mortem neurotypical 31-year-old individual of African ancestry from the Lieber Institute for Brain Development (LIBD, ID: LIBD75) to assess our ability to detect genetic variation, particularly somatic mutations, by various assays and tools using a donor-specific genome assembly (**Fig. 1**). This individual was also examined as part of the Brain Somatic Mosaicism Network (BSMN)^37^, which provided us with additional data, including Illumina NovaSeq and 10x Genomics linked-read WGS data^64^. We isolated 160 mg of DLPFC tissue to perform bulk long-read WGS. We generated 193 Gb of sequence comprising 93.5 million reads, with an average N50 of 3.7 kb (**Table 1, Supplementary Table 1**). We further applied our TE nanopore Cas9-Targeted Sequencing (TEnCATS)^53^ approach to the bulk DLPFC tissue to specifically capture active *Alu* elements and Long Interspersed Element-1 (LINE-1s or L1s). The on-target rates of our TEnCATS approach ranged from 9.48% to 48.7%, consistent with our previous findings^53^.

**Table 1.** Multi-platform DNA sequences of LIBD75 frontal cortex.

| Sample | Assay | Platform | Yield | Reads | N50 | Coverage | On-target Rate |
| --- | --- | --- | --- | --- | --- | --- | --- |
| Bulk Tissue | WGS | PromethION x3 | 195Gb | 93.5M | 3.7kb (avg) | 61x | - |
|  | TEnCATS, L1Hs | MinION x2 | 3.7Gb | 2.5M | 3.3kb | 77.4x* | 0.0948 |
|  | TEnCATS, *Alu*Ya5 & *Alu*Yb8 | PromethION x3, MinION | 35.5Gb | 20.8M | 3.3kb | 224x* | 0.56 |
|  | WGS | NovaSeq** | 94.3Gb | 624M | 151bp | 29.7x | - |
|  | WGS | Chromium** | 96.3Gb | 753M | 128bp | 28.4x | - |
|  | Assay | Platform | Yield per cell | Reads per cell | N50 | Genome Covered per cell | Cell # |
| Neurons | MALBAC, WGS | MinION | 1.4Gb*** | 1.4M | 1.2kb | 0.22 | 115 |
|  | MALBAC, WGS | PromethION | 6.0Gb**** | 5.7M | 1.3kb | 0.53 | 5 |
|  | MALBAC, WGS | PromethION | 89Gb | 93M | 1.2kb | 0.80 | 1 |
|  | MALBAC, WGS | NovaSeq | 1.3Gb | 8.3M | 151bp | 0.21 | 94 |

\*, On-target coverage: coverage over/on the targeted regions.
\*\*, Data obtained from Garrison, M. A. et al. 2023.
\*\*\*, One, five or ten cells per flow cell (Supplementary Table1).
\*\*\*\*, Five cells per flow cell.
Three sets of WGS data were generated from LIBD75 DLPFC using ONT, 10x Chromium and Illumina NovaSeq platforms. TEnCATs was also performed using this tissue to characterize L1Hs, *Alu*Ya5, and *Alu*Yb8 elements. Additionally, single neurons were sequenced using MALBAC WGA and sequenced in batches of one, five, and ten cell batches on MinION and PromethION flow cells. 94 MALBAC samples (89 in common with samples sequenced using ONT platforms) were sequenced using Illumina NovaSeq.

We next employed ONT sequencing on Multiple Annealing and Loop-Based Amplification Cycles (MALBAC)^65^ single-cell libraries to investigate potential somatic mutations in individual neurons. This MALBAC protocol^66^ was optimized to produce longer (>1kb) amplicons for long-read sequencing. MALBAC has also been shown to generate quasi-linear amplification, reducing biases and providing more uniform genomic coverage compared to other WGA methods^65,67–69^. To evaluate the efficiency of ONT sequencing on MALBAC-amplified genomes, we examined different batch sizes of pooled single-neuronal cell amplifications (**Table 1**). Specifically, we first sequenced 115 single cell libraries in batches of one, five, and ten cells on MinION flow cells, achieving an average 22% overall physical genome coverage, defined as the percent of the genome covered by at least one read (**Supplementary Table 1**). Next, we randomly chose five of these cells (9100, 9102, 9103, 9104, 9107) to sequence on a PromethION flow cell to examine the impact of higher sequencing depth on overall physical genome coverage and observed a commensurate increase (53%). Finally, we selected one cell (9203) that initial analysis suggested harbored several large CNVs to deeply sequence using a PromethION flow cell, achieving an 80% physical genome coverage (**Table 1, Supplementary Figure 1, Supplementary Table 1**). The read lengths, measured as N50, consistently ranged from 1.2 kb to 1.5 kb across the 121 samples, marking an almost 10-fold increase compared to standard short-read sequencing technologies. Additionally, we used the NovaSeq platform to sequence 94 MALBAC-amplified neurons with short-reads, 89 of which were a subset of those with ONT single-cell sequencing. The short-read-based single-cell sequencing achieved an average physical genome coverage rate of approximately 21% per cell, which was similar to the physical coverage obtained by ONT MinION single-cell sequencing. (**Table 1, Supplementary Figure 1, Supplementary Table 1**).

### Construction of a haplotype-resolved assembly for the LIBD75 DLPFC tissue

Using our sequencing data produced across multiple platforms, we generated a haplotype-resolved DSA to provide phasing information and facilitate germline and somatic variation calling as we have previously shown that using haplotype information can greatly improve accuracy in somatic variant discovery^21,30^. First, we established a scalable and efficient pipeline for a personalized, haplotype-resolved assembly (**Fig. 2a**). The raw diploid assembly was generated using Shasta and HapDup using ONT reads^60^. Illumina short-reads were then incorporated to polish the draft diploid assembly to resolve inaccuracies due to the more error-prone ONT long-read sequences. We produced a haplotype-resolved genome with a size of 5.77 Gb for the donor tissue sample (**Supplementary Table 2**). The phased contigs had an N50 of 0.75 Mb before further refinement, with other quality metrics comparable to prior studies^60,70^, suggesting a high-quality assembly (**Supplementary Figure 2, Supplementary Table 2**). We achieved a 93.18% recall rate of the phased SNVs called from the phased linked-read data, which were not used in the initial assembly construction at this stage. We then identified a high-confidence collection of phased genetic variants from the final assembly, including 4,310,781 SNVs, 625,451 INDELs (1-49 bp), and 26,248 SVs ranging from 50 bp to 95,192 bp using established assembly-based variant calling methods^3^ (**Fig. 2b, c, Supplementary Table 3,** see **Methods**). Out of the 26,712 SVs we detected, we observed 10,613 deletions, 16,069 insertions, and 30 inversions. These findings are consistent with the levels reported in our previous research on samples of similar ancestry^3^. We also annotated 1,818 TEs, including 172 L1Hs, 1565 *Alu*Ys, and 81 SVAs, projecting expected peaks in the overall length distribution at 320 bp and 6 kb for full-length *Alu* and L1, respectively (**Fig. 2b**).

**Figure 2.**
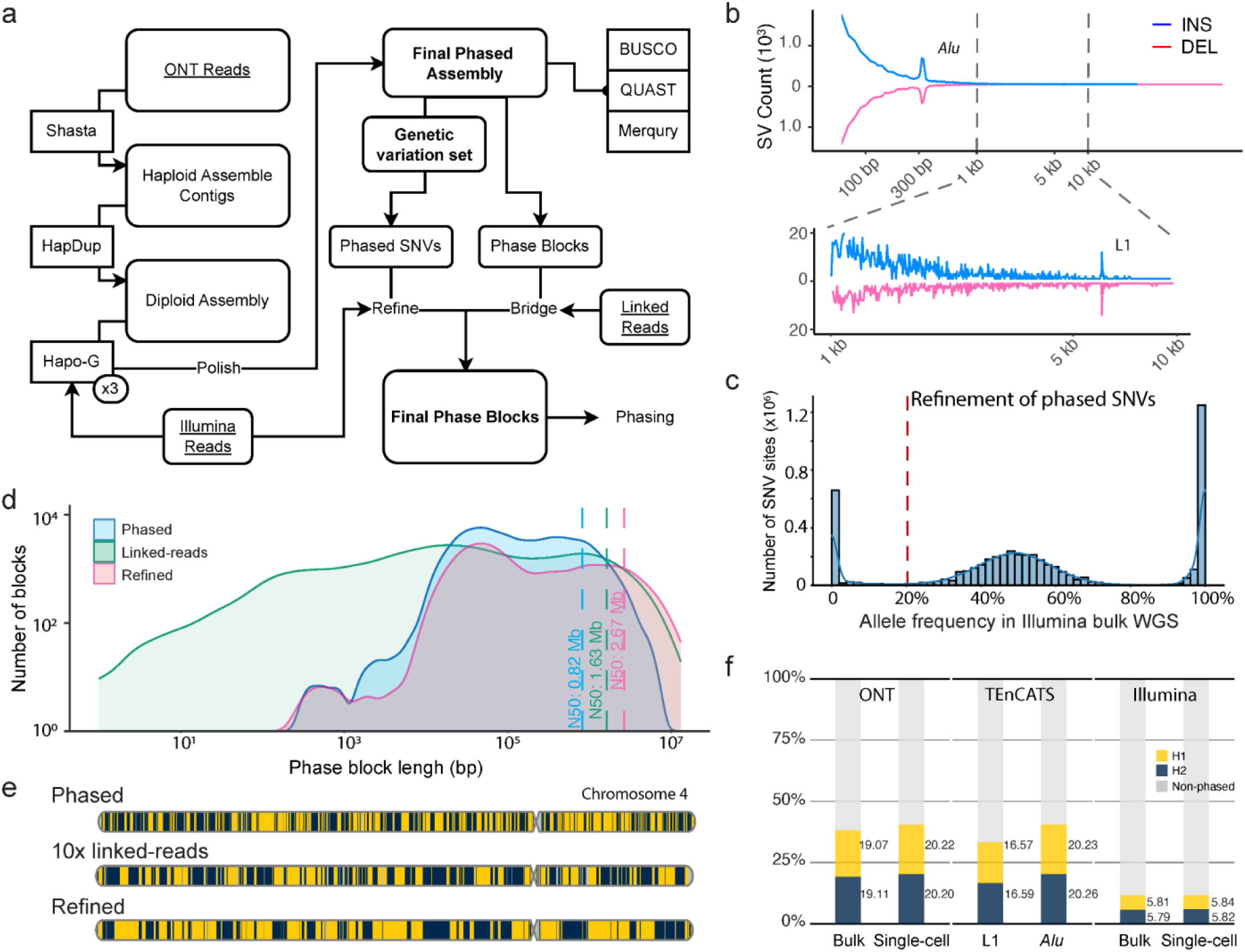
Construction of a haplotype-resolved donor-specific assembly to facilitate genetic variation calling in the LIBD75 DLPFC tissue. **a,** Pipeline to generate a haplotype-resolved assembly for LIBD75 DLPFC tissue using bulk sequences (underlined). ONT reads were used to build the raw diploid assemblies. Illumina reads were used to refine the raw assemblies and phased SNVs due to its high accuracy in point mutations. Linked reads were used to bridge the phase block for the two haplotypes. The three deliverables are highlighted in bold font within the diagram. **b,** Number and length distributions of assembly contig-based genetic variations. **c,** Refinement of phased contig-based SNVs was based on the allele frequency distribution in the Illumina bulk WGS. A 20% allele frequency cutoff is denoted by the red dotted line. **d,** Length distribution of phase blocks from the phased assembly (blue), linked reads called by LongRanger2.0 (green), and the final refined assembly (red) by bridging those from phased assembly and linked reads. The N50 length is denoted by dotted lines based on the phase blocks after filtering out reads without heterozygous phased SNVs in each category. **e,** An example (chromosome 4) shows the improvement of the final refined phased blocks versus those from phased assembly and linked reads. Adjacent blocks are colored in maize and blue. **f,** Phasing rates across the various platform sequences based on assembly information. Maize represents reads in haplotype 1 (H1), blue represents reads in haplotype 2 (H2), and grey represents non-phased reads.

To maximize the length of our phase blocks that contain resolved and distinguished maternal and paternal haplotype information, we next used these initial haplotype-resolved germline variants and developed a pipeline designed to bridge the phase blocks (N50 = 0.82 Mb) from the haplotype-resolved assembly with those (N50 = 1.63 Mb) derived from linked-reads from the same tissue (see **Methods**). Through this approach, we produced a refined set of phase blocks with an N50 of 2.67 Mb, significantly extending the length of the original blocks (**Fig. 2a, d, e**). We then utilized this refined phase block set in downstream analyses to phase the ONT long-reads and Illumina short-reads, thereby providing essential phasing information to resolve genetic variation. Post-phasing, we observed an average 3.27-fold increase in the phasing rate for long-reads (38.06%) compared to short-reads (11.64%) (**Fig. 2f**). These rates were consistent across various experiments, including bulk tissue, single cells, and TEnCATS.

### Assessment of germline genetic variants in bulk tissue across sequencing platforms

We next interrogated our set of germline variants across different technologies to establish an upper bound of calling efficacy to inform our somatic discovery (**Supplementary Table 3**). First, we identified a high-confidence subset of our assembly-based germline variants by examining their derived allele frequency from both the short- and long-read bulk WGS sequence data and filtered out variation that fell below empirically derived variant allele frequencies (VAF) (**Supplementary Figure 3**, see **Methods**). This high-confidence set was then used to assess the calling efficacy across sequencing platforms with different germline variant callers.

The long-read sequencing technologies overall achieved better recall rates than short-reads in bulk tissue for germline SNVs (99.4% vs. 98.1%), SVs (99.6% vs. 29.6%), and TEs (99.7% vs. 92.7%) (**Fig. 3a, Supplementary Table 4**). The VAF of these germline calls conformed to expected homozygous and heterozygous distributions within each data type^71^, though some calls were missing in ONT or Illumina bulk tissue sequencing (**Fig. 3a**). We observed an average 2.5-fold increase in recall rates for long-reads compared to short-reads in SV detection across various SV and TE types (**Fig. 3b, Supplementary Table 4**), consistent with previous reports^49,72^. Given the sparsity of single-cell sequencing data, we were unable to directly interrogate the majority of germline variants in individual cells. Instead, we constructed a pseudo-bulk sample by combining individual single cells to assess the upper bound of germline genetic variant calling using single-cell long-read and short-read sequencing data (see **Methods**). Similar to the bulk tissue sequencing, we observed higher recall rates (95.4% vs 92.9% for SNV, 42.8% vs 13.7% for SV, and 73.0% vs 58.2% for TE) from single-cell ONT sequencing compared to Illumina sequencing (**Fig. 3c**, **Supplementary Figure 4**). Additionally, increased yield through pooling cells resulted in better recall rates (**Fig. 3c**, **Supplementary Figure 4**), suggesting that the MALBAC-amplified DNA libraries have high complexity that was not completely saturated at the current sequencing coverage. Overall, this analysis shows that high-fidelity germline genetic variants can be called across multiple sequencing platforms and provides a ground truth to compare putative somatic variation using these same technologies.

**Figure 3.**
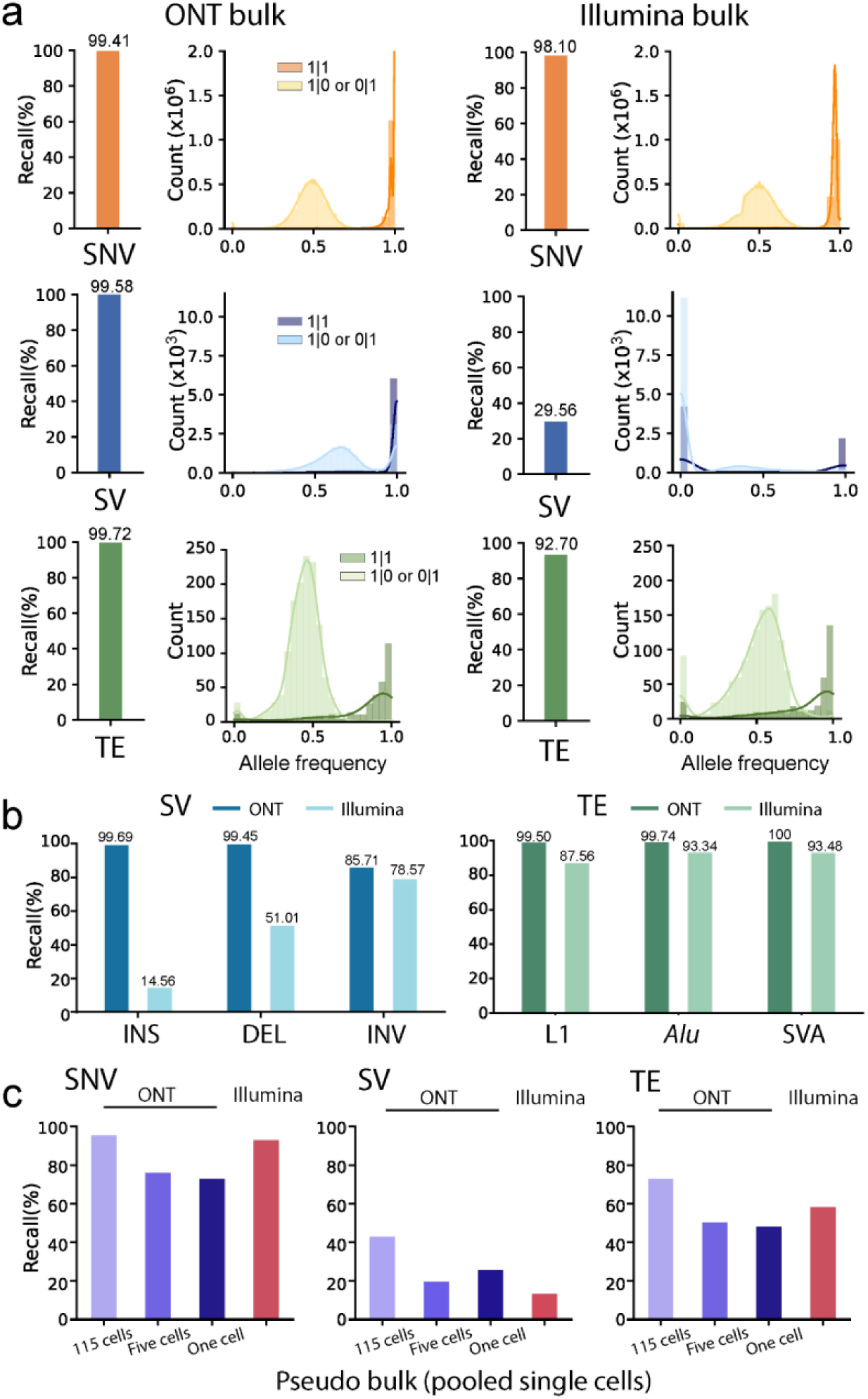
Assessment of germline genetic variants in bulk tissue across sequencing platforms. **a,** The recall rates (bar plots) and allele frequency distributions (histograms) in ONT bulk WGS sequencing (left) and Illumina bulk WGS sequencing (right). Orange represents SNVs, blue represents SVs, and green represents TEs. b, Recall rates of SV (left) and TE (right) subtypes in the analysis of ONT and Illumina bulk tissue. c, Recall rates of SNV (left), SV (middle), and TE (right) in pseudobulk samples derived from pooled single-cell runs. 115 cells (light purple) are single-cell libraries in batches of one, five, and ten cells on MinION flow cells. Five cells (purple) are in batch on one PromethION flow cell. And one cell (dark purple) is sequenced by one PromethION.

### Donor-specific assembly refines somatic mosaicism in bulk tissue

In contrast to cancer studies that typically used matched normal tissue as a control and are focused on higher frequency somatic events, there are a limited number of available tools for the discovery of somatic mosaicism in bulk tissue alone. Somatic Mosaicism across Human Tissues Network (SMaHT)^73^ is starting benchmarking efforts to fill these gaps and advance detection of low-VAF somatic mutations (SNVs, indels, MEIs, SVs)^74–78^.

To identify somatic mosaicism within our dataset, we utilized MosaicForecast^79^, Sniffles2 mosaic model^52^, and an enhanced mosaic model of PALMER^51^ for somatic mosaicism discovery of SNVs, SVs, and TEs, respectively (**Supplementary Table 3**). Theoretically, a somatic mutation should be observed only on the haplotype from which it originated, and thus there are three primary categories of potential false positives when identifying somatic variations in bulk tissue (**Fig. 2a**): those introduced by the unequal representation of haplotypes during sequencing (*hapErrors*), those by mapping errors introduced by recurrent sequencing errors (*seqErrors*), and those from mapping errors likely due to genomic repetitive context (*mapErrors*). Such errors cannot be distinguished from *bona fide* somatic variation by assessing overall allele frequency alone. Based on previous studies, we posited that leveraging the haplotype information from our DSA would enable us to filter out alleles present at low frequencies on both haplotypes that are unlikely to be true somatic events^21,30,80^. To do this, we first calculated the allele frequency (AF) for each candidate variant within each haplotype and compared their relative abundances. For germline SNVs, homozygous variants were enriched near the AF=1 position for both haplotypes (x>= 0.8 and y >=0.8, 95.2% 1,262,689 out of 1,326,778), and heterozygous calls clustered near AF=1 for one haplotype and AF=0 for the other (x >=0.8 and y <= 0.2, 95.8%, 2,787,756 out of 2,910,400) (**Fig. 4b**). For germline SVs and TEs, we observed a lower enrichment towards expected allele frequencies, with 70.1% (3,129/4,461) and 35.4% (127/359) for homozygous variants, and 72.1% (8,439/11,704) and 36.2% (550/1,519) for heterozygous variants, respectively (**Supplementary Figure 5**). This suggests that the complexity and repetitive nature of the genetic variant may affect the accuracy of AF estimation.

**Figure 4.**
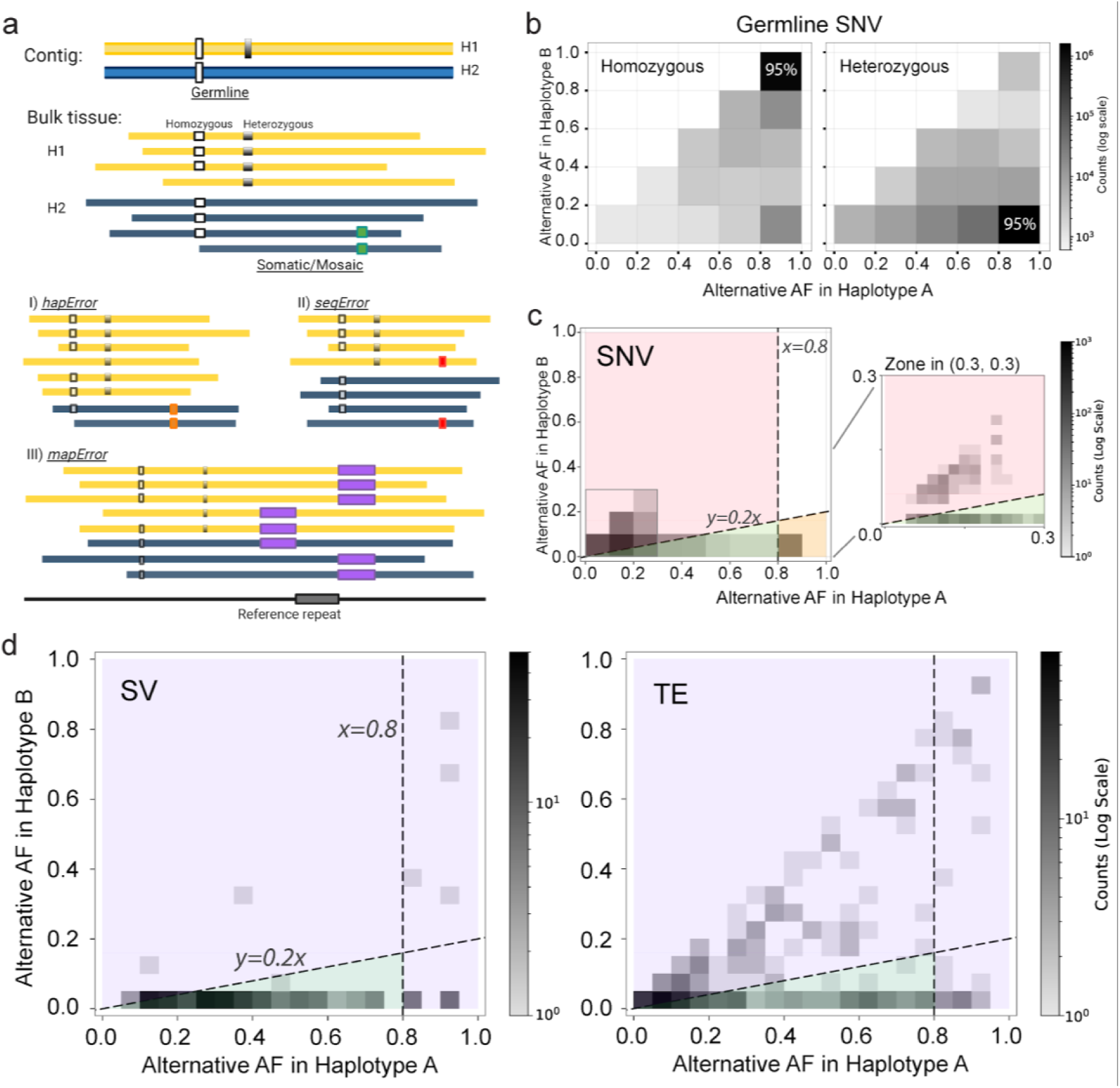
Haplotype-based analysis enables the removal of false positive somatic calls in bulk tissue. **a,** Schematic illustrating i) the germline (white and black) and somatic (green) genetic variants in the diploid contigs and reads, and ii) the use of phasing information to eliminate false positive somatic calls due to unequal representation of haplotypes (*hapErrors*, orange), sequencing noise (*seqErrors*, red), or misalignment errors (*mapErrors*, purple). **b,** Heatmaps of germline homozygous SNVs (left) and heterozygous SNVs (right) in ONT WGS bulk tissue sequences. The X-axis represents the allele frequency (AF) in Haplotype A, which contains the highest alternative allele frequency, and the Y-axis represents the AF in Haplotype B, the second haplotype. c, 2D kernel density plot (left, bin=0.1AF) for putative somatic SNVs showing the exclusion of false positives by *hapErrors* (bottom right, orange block) and *seqErrors* (upper left, red block) using two dotted lines (*x=0.8* and *y=0.2x*). The plot (right, bin=0.02AF) offers a magnified view of the range between (0,0) and (0.3, 0.3). **d,** 2D kernel density plots (bin=0.03AF) for putative somatic variants: SVs (left) and TEs (right). *mapErrors* is indicated by purple blocks.

We next identified 6,516 candidate somatic variants in bulk tissue using the tools above, including 5,353 SNVs, 769 SV, and 394 TEs. Among these, 1,407 SNVs, 312 SVs and 363 TEs were located in regions with balanced reads across both haplotypes (see **Methods**). When applying the same analysis, potential true somatic mutations are expected to be observed on only one haplotype with an AF<0.8 and near AF=0 for the other haplotype (i.e. AF_A_<0.8 and AF_B_=0). For SNVs, we observed that calls falling at the AF_A_≥0.8 positions could be potential false positive *hapErrors* introduced by the unequal representation of haplotypes, and excluded such variants (**Fig. 4c, Supplementary Table 5, 6**). We also set an empirical boundary (*y* = 0.2*x*) to exclude false positives (*seqErrors or mapErrors*) introduced by sequencing or mapping errors (**Fig. 4c, Supplementary Figure 5, 6**). Using haplotype-based analysis from the DSA haplotype information, we achieved a removal rate of 75.1% for SNVs for false positive somatic calls in bulk tissue. Specifically, the SNV removal rate was 68.2% (959 out of 1,407) for false positive *hapErrors* and 7.0% (98 out of 1,407) for *seqErrors*. For SVs and TEs we observed that calls falling at the AF_A_≥0.8 positions or the empirical boundary (*y* = 0.2*x*) could be potential false positive *mapErrors* introduced by the mapping errors due to genomic context (**Fig. 2a**). With the haplotype information, we were able to remove 15.4% (48 out of 312) of SVs and 38.9% (141 out of 363) of TEs for false positive somatic calls in bulk tissue as *mapErrors* (**Fig. 4d, e**). As expected, the larger rate observed in *mapErrors* for SVs and TEs, compared to other variants, suggests that the detection of somatic SVs and TEs are likely impacted more by genomic content due to their complex and repetitive nature^51,81^. More specifically, we found that the somatic calling of SVs is significantly affected by tandem repeats, while the detection of TEs is impacted by repetitive TEs (**Supplementary Figure 7, Supplementary Table 7, 8**). Finally, we randomly selected a subset of our false positive candidates (115 out of 1,246) and manually confirmed that they were likely due to unequal representation of haplotypes, sequencing errors, and mapping errors (**Supplementary Table 5-8**). Ultimately, we identified 836 candidate somatic calls: 350 SNVs, 264 SVs, and 222 TEs (see **Data Availability**).

### TEnCATS detects non-reference germline TEs efficiently in bulk tissue and observes somatic TEs only at low frequency

As an alternate method to investigate TEs within the DLPFC tissue, we employed Transposable Element nanopore Cas9-Targeted Sequencing (TEnCATS). This technique leverages CRISPR-Cas9 with guide RNAs targeted to specific sequences that selectively identify transposable elements^53^. TEnCATS is capable of achieving high coverage over the target elements, such as L1Hs and *Alu*, facilitating the characterization of low-frequency somatic events.

We applied TEnCATS to reference and non-reference L1Hs and *Alu* elements (AluYa5 and AluYb8) in the genome, achieving average read coverages of 77.4x and 224x at reference targeted sites, respectively (**Table 1**). Using NanoPal, our previously published pipeline for detecting germline TEs in long-read Cas9-enrichment data^53^, we identified 180 non-reference germline L1Hs and 1,165 *Alu*Ya and Yb elements (**Supplementary Table 9,** see **Method**). We first examined the germline L1Hs and *Alu*s targeted by TEnCATS and observed similarly high recall rates compared to the ones in ONT WGS in bulk tissue (98.8% versus 99.5% for L1 and 92.5-95.2% versus 99.7% for *Alu*) (**Fig. 3b, 5a, Supplementary Table 9**). Notably, despite a significantly lower total sequencing yield compared to ONT WGS (3.7 Gb in L1Hs and 35.5 Gb in *Alu*s versus 195 Gb), NanoPal combined with TEnCATS showed a similar number of supporting reads for non-reference L1Hs compared to PALMER using ONT WGS data (mean 36.9 versus 32.0) and exhibited a substantial enrichment in supporting reads for non-reference *Alu*s (mean 225.0 versus 32.4) (**Fig. 5b**).

**Figure 5.**
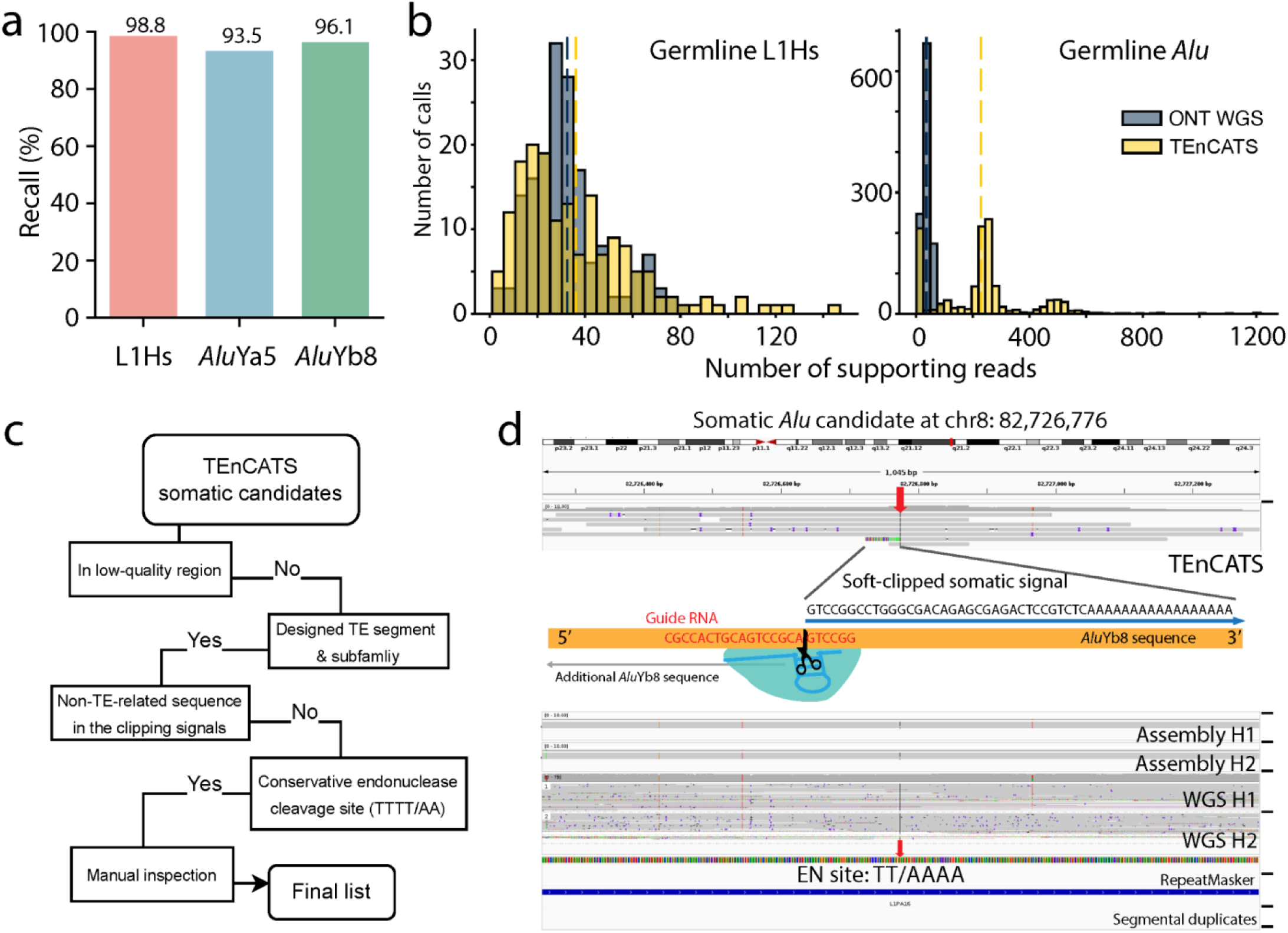
TEnCATS methodology detects non-reference transposable elements (TEs) within donor DLPFC tissue. **a,** Recall rates for targeted active TE subfamilies by TEnCATS based on the assembly-based TE callset. **b,** Number of supporting reads of non-reference TEs reported by NanoPal from TEnCATS versus PALMER from ONT WGS. Dotted line represents the mean value of the group. **c**, A semi-automatic pipeline with manual inspection was used to stringently refine somatic TE candidates into a final list. Low-quality regions include any segmental duplicates, low-confidence mask regions used in this project, and reference *Alu* repeats or LINE-1 repeat regions in RepeatMask for *Alu* candidates or L1Hs candidates, respectively. Non-TE-related sequences refer to any genomic content that is not TE sequence, polyA tracts, target site duplications, or potential transduction sequences between two polyA tract signals ^51^. We ran BLAT and IGV for the final manual inspection process. **d,** IGV screenshot of a somatic *Alu* element candidate captured by TEnCATS at chr8:82,726,776 (H1, haplotype1; H2, haplotype2). The red arrow indicates the insertion site. The middle illustration demostrates the soft-clipped sequence in the supporting read representing the somatic signal at the 3’ end of *Alu*Yb8. This sequence aligns with the cut site of guide RNA (marked in red) and the AluYb8 consensus sequence (orange bar).

We then investigated potential somatic TE candidates identified by TEnCATS. We gathered all reads with putative somatic signals, excluding those associated with identified non-reference germline TEs, and applied a stringent pipeline to refine these signals (**Fig. 5c**). Through this process, we identified 14 putative somatic TE candidates, each supported by a single read. Combining these with 34 putative non-reference L1Hs elements and 1,462 *Alu*Y elements, each supported by more than two reads and at least one read in each direction (see **Data Availability**), we conducted a meticulous manual inspection. Ultimately, we concluded with eight somatic TE candidates (**Supplementary Table 9**), all supported by only one read but with all expected features of a *bona fide* TE insertion. Although we cannot rule out potential ligation artifacts in these instances, these ultra-low frequency candidates suggest possible TE somatic activity only in the late stages of tissue development in this individual. An example of a detected somatic *Alu* candidate inserted in chromosome 8p21.13 is presented in **Figure 5d**.

### Haplotype-aware detection for somatic CNVs in single cells

We next analyzed sequencing data from MALBAC whole-genome amplified single neurons. As ONT long-read single-cell data has not been widely implemented yet, there are currently no tools available for identifying large CNVs and TE insertions from such data. Recently, our group utilized phase information from Illumina short-read single-cell data and developed a novel tool to investigate more than 2,000 human neurons^21^. Building on this, we developed a pipeline called GARLIC (Genome-wide Allelic copy numbeR variation Locator In Cells) to detect large somatic deletions in single neurons using long-read data. GARLIC uses a circular binary segmentation (CBS) algorithm to process a statistic called physical phased coverage (PPC), which leverages phase information from a donor-specific assembly (**Supplementary Figure 8**, see **Methods**). Additionally, we applied a modified version of Ginkgo^21,50^ to generate a companion callset from our short-read single-cell sequencing data.

For deletions larger than 1 Mb, the long-read data identified 5.9 times more candidate deletions than the short-read data. Specifically, we detected 279 candidate deletions (median size 1.48 Mb) from long-reads, compared to 47 (median size 2.63 Mb) from short-reads (**Fig. 6a, Supplementary Table 10**). Interestingly, 6.1% (seven out of 115) of neurons from the long-read data and 5.3% (five out of 94) from the short-read data showed aggregated deletions exceeding 10 Mb, aligning with previous observations^21^ (**Fig. 6b**). Nine somatic deletion candidates were reported by both technologies, with five occurring in the same neurons (one in neuron 9095 and four in neuron 9203) (**Fig. 6c**). We then leveraged both sequencing coverage and PPC to manually inspect these calls. Interestingly, we identified two adjacent somatic deletion candidates on chromosome 7 in neuron 9203, detected by both technologies, each approximately 5 Mb in length (**Fig. 6d**). Visualizations of read coverage and PPC metrics in these regions showed consistent lower and higher signals, respectively, supporting the predicted deletions while no significant signals were observed in other cells at these positions. By leveraging phase information in long-read single-cell sequences, we identified at least five high-confidence somatic deletion candidates supported by both long-read and short-read technologies, achieving a refined callset of somatic deletion candidates in terms of number and length in single neurons using long-read data compared to short-read data.

**Figure 6.**
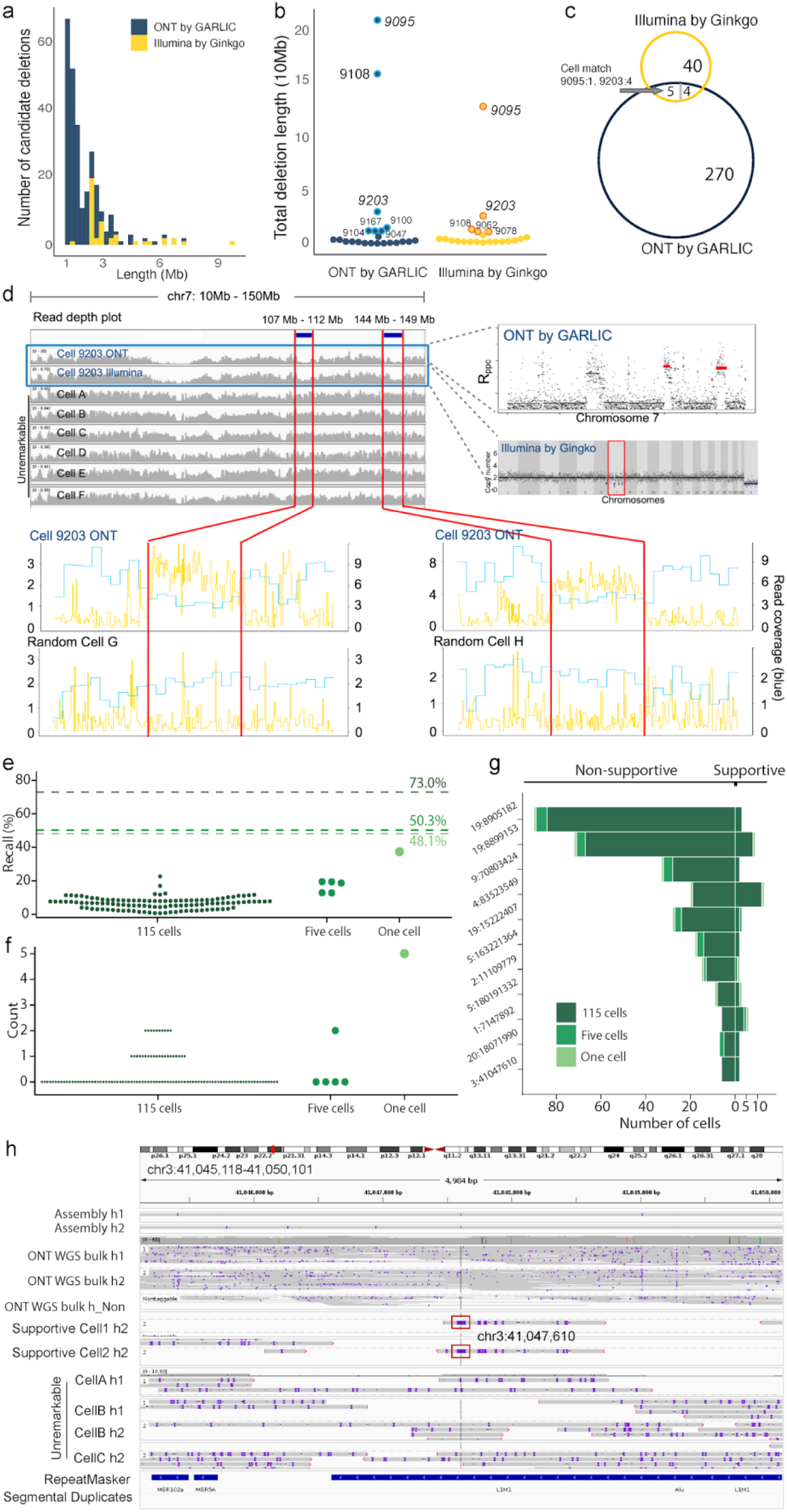
Haplotype-aware detection of somatic CNVs and TEs in single neurons. **a.** Length distribution of candidate somatic deletions (>1 Mb) identified using ONT long-read sequencing by GARLIC and Illumina short-read sequencing by Ginkgo. **b**, Swarm plot of aggregated potential somatic deletion detected in each cell by two methods. Cells with aggregated deletions more than 10Mb were highlighted by bold circles. **c**, Overlap calls by two methods. **a**, **b**, and **c** share the same color legend: ONT by GARLIC (blue) and Illumina by Ginkgo (maize). **d**. An example of two candidate somatic deletions on chr7 in a single neuron (9203) detected by two methods. The main panel shows the read depth plots for the 9203 single neuron from ONT single-cell sequencing (above, blue frame) and Illumina single-cell sequencing (below for six random single neurons). The left panel shows the signal distribution from Rppc by GARLIC for chr7 in neuron 9203 (above) and the copy number states by Ginkgo (below). The bottom panel illustrates the signals from ONT single-cell sequences for two mutations, Rppc (yellow) and read coverage (blue). The plots for neuron 9203 are depicted above, while plots for three random cells are depicted below. Signals representing the three candidate somatic deletions across all panels are highlighted between red bars. **e**, Swarm plot of recall rates for high-confidence assembly-based germline TEs in individual single cells. Each point represents one cell, and dashed lines represent pooled single cells. 115 cells (dark green) are single-cell libraries in batches of one, five, and ten cells on MinION flow cells. Five cells (green) are in batch on one PromethION flow cell. And one cell (light purple) is sequenced by one PromethION. The same color legend for dashed lines applies to **f** and **g**. **f**, Number of candidate somatic calls per individual single cell. Each point represents one cell. **g**, Number of cells in which each candidate somatic call is detected. Left bar, non-supportive cells with non-supportive go-through reads; right bar, supportive cells with supportive go-through reads, or left or right clipped reads. **h**, A candidate somatic Alu insertion at chromosome 3 was observed in two out of 121 cell samples and was not detected by ONT WGS in bulk tissue.

### Haplotype-aware detection for somatic TE detection in single cells

To identify TEs from single-cell long-read sequencing data, we developed a pipeline called PalmeSom, building on our earlier PALMER method^51^ (and Wang et al. in prep.^82^). PalmeSom can annotate reads from different haplotypes that show potential non-reference TE signals. In annotating TE signals, PalmeSom evaluates the coordinates of the aligned consensus sequences and the structural variation signals within the reads to enhance accuracy (**Supplementary Figure 8**, see **Methods**). When applied to 121 single-neuron samples, PalmeSom identified 5,423 non-redundant TE clusters including 4,947 *Alu* elements, 46 SVA elements, and 430 L1 elements. Benchmarking against high-confidence germline callsets derived from the DSA, the average recall rate per MinION flow cell was 6.55%, compared to 16.60% per PromethION flow cell and 37.37% for single cell 9203 sequenced on a single PromethION, with pooled recall rates for these groups being 72.97%, 50.28%, and 48.11%, respectively (**Fig. 3c, 6e**).

PalmeSom further refines high-confidence TE calls to identify somatic TEs by incorporating additional features, such as the number of non-supportive reads, signal read counts, number of cells with signals, and haplotype information. This analysis revealed 61 candidate somatic *Alu* elements and one candidate somatic L1 element (**Fig. 6f**). After thoroughly examining all 62 candidates, we identified eleven *Alu* candidates as likely somatic insertions. These insertions were supported by single cells containing supportive or non-supportive reads, with more cells exhibiting non-supportive reads than supportive reads (**Fig. 6g**). Five of these candidates were assessed as high-confidence (**Supplementary Table 11**). We show an example high-confidence somatic *Alu* insertion candidate at chromosome 3p22.1, observed in two out of 121 single cell samples (**Fig. 6h**). This insertion was not detected by ONT WGS in bulk tissue or TEnCATS, likely due to being a very low frequency mosaicism. Overall, our analysis demonstrates the ability to identify somatic TEs from whole-genome amplified single-cell data, though the process is constrained by the total number of reads available per cell and the number of cells sequenced.

## Discussion

In this study we aimed to refine the detection of somatic mosaicism in the human dorsolateral prefrontal cortex (DLPFC) by constructing a high-quality, personalized, haplotype-resolved assembly using multiple sequencing platforms. This approach combined Oxford Nanopore Technologies (ONT) long-read sequencing, Illumina NovaSeq short-read sequencing, 10x linked-read sequencing, and Cas9-targeted sequencing (TEnCATS), enabling us to investigate somatic mutations in bulk tissue and individual neurons. Over the past four years there have been significant efforts to survey germline genetic variation within large cohorts, usually by large genomic consortia^2,3,81,83^. While most were limited to cell lines, they demonstrated the ability to accurately characterize variants using multiple sequencing platforms. Other consortia, such as the Somatic Mosaicism across Human Tissues Network (SMaHT)^73^ and the Brain Somatic Mosaicism Network (BSMN)^40^, have made strides in systematically documenting tissue DNA variation using donor-specific genomes coupled with state-of-the-art sequencing technologies and analysis tools. By leveraging these advances, including single-cell sequencing and novel long-read single-cell variant detection callers, we were able to identify new insights into somatic variation in non-cancerous human tissue, demonstrating the potential of donor-specific assemblies (DSA) to enhance variant detection.

A key finding from our work is the substantial improvement in somatic variant calling that we achieved by incorporating haplotype-resolved genome assembly. Using this DSA, we were able to significantly reduce false positive somatic calls in bulk tissue. The use of haplotype phasing enabled us to filter out variants that were likely due to sequencing errors or unequal representation of haplotypes, achieving up to a 75.1% reduction in false positives for SNVs, 15.4% for SVs, and 38.9% for TEs. This underscores the importance of using donor-specific phasing to enhance the accuracy of somatic variant detection, especially in tissues with non-clonal genetic variation, such as the brain. In addition, long-read sequencing played a crucial role in overcoming the limitations of short-read sequencing, particularly in detecting SVs and TEs. The repetitive nature of these regions makes them difficult to resolve with short-read technologies; however, long-read sequencing allowed us to generate megabase-scale phase blocks and *de novo* diploid contigs that facilitated more accurate variant calling. This was particularly evident in our ability to identify large candidate somatic deletions and insertions in both bulk tissue and single neurons, and would have been challenging with short reads alone.

Our work also introduced efficient technology and innovative tools for large somatic variant detection at both the bulk tissue and single-cell levels. GARLIC, a novel pipeline we developed for detecting large somatic deletions in single neurons, showed enhanced performance compared to existing methods^50^. By leveraging haplotype information, GARLIC was able to detect the same number of somatic deletions reported by traditional short-read single-cell sequencing approaches and numerous smaller somatic deletions with greater sensitivity. For TE detection, we showed that TEnCATS is not only able to identify non-reference germline TEs from bulk tissue, as demonstrated in this and previous studies^53^, but also provide a superior orthogonal technology to WGS for somatic TE detection. To note, eight somatic *Alu* candidates identified by TEnCATS were supported by a single read after we rigorously inspected and ruled out all signals with more than one supporting read. Our TEnCATS capture experiments achieve over 250x and 80x coverage on the target homozygous germline and reference *Alu* and L1 on average, respectively. This may indicate a lack of somatic TEs at an allele frequency of more than 0.4% (1/250) for *Alu*s and 1.25% (1/80) for L1s in the DLPFC of this LIBD75 individual. Additionally, PalmeSom, our pipeline for detecting somatic TEs in single-cell long-read sequencing data, successfully identified at least five high-confidence somatic TE candidates from the small number of cells sequenced. Since all detected elements were *Alu*s and we did not observe the somatic L1 mutation reported in previous publications (ranging from 0.2 to 0.6 per neuron)^84,85^, our ability to detect larger TE families may be limited by the current single-cell ONT read length. However, this still highlights the potential of long-read sequencing for single-cell variant detection.

Whole genome amplification (WGA) has previously been applied to explore genetic variation at the single cell level^86–89^. This approach benefits from the integration of long-read sequencing technologies, such as those applied in SMOOTH-seq^90^ and droplet MDA^91^. The longer reads provided by these platforms improve mappability and phasing capabilities, enabling more accurate detection of somatic variation. However, these advancements are limited by lower per-base accuracy, chimeric artifacts, and jackpotting^91,92^. In this study, we observed similar trends when sequencing MALBAC libraries using ONT, where the average N50 of 1.5 kb provided longer fragments, yet still resulted in segmented genome coverage. Despite these limitations, our findings underscore that further refinement of WGA techniques combined with long-read sequencing holds significant potential for improving the detection of genetic variation at both the tissue and single-cell levels, particularly in complex genomic regions such as SVs and TEs.

While our results are promising, there are limitations that should be addressed in future research. Targeted ONT sequencing from frozen post-mortem brain tissue produced shorter read fragments compared to studies using cell lines^93^. This led to a decreased proportion of long, mappable reads, impacting both bulk tissue WGS and TEnCATS sequencing analysis. Additionally, the relatively low sequencing coverage in some of the single -cell experiments may have restricted our ability to detect rarer somatic variants with lower allele frequencies. Although TEnCATS can detect somatic TE candidates at very low frequencies, the potential for experimental ligation artifacts from a single read cannot be entirely dismissed. Conducting a multi-tissue study could likely address this issue. Furthermore, while we observed a number of somatic variants, their significance in the context of neuronal function remains to be fully explored. It is evident that somatic mutations contribute to neuronal diversity, but further studies are needed to understand their role in disease.

Looking ahead, our study lays the groundwork for several future directions. Expanding the analysis to a larger and more diverse cohort of individuals would help assess the broader applicability of our findings. Increasing sequencing depth and coverage, particularly for single-cell sequencing, would allow us to detect even rarer somatic mutations. Furthermore, integrating other genomic and transcriptomic data could provide a more comprehensive view of the functional impact of somatic mutations in the brain.

## Methods

### Single-cell isolation and MALBAC amplification from frontal cortex tissue

Flow-sorted single-cell dorsolateral prefrontal cortical neuron MALBAC libraries from a 31.3-year-old male neurotypical individual of African ancestry^64^ (LIBD75) were prepared using the method described in Burbulis et al.^66^. Briefly, 2,000-40,000 neuronal cells in 2mL of 1X Phosphate Buffered Saline (PBS) were applied to rafts, and single cells were isolated using the CellRaft system (Cell Microsystems). 120 isolated single neurons were transferred to PCR tubes and lysed in 2.5µL of lysis buffer with 25µL of PCR-grade mineral oil laid on top. Following lysis, 2.5µL of 2X amplification buffer was added to the tubes, heated to 95°C for 3 minutes then snap-cooled on ice before the addition of 0.6µL of enzyme mix. 6 cycles of amplification were completed using the following protocol: 10°C for 45 seconds (sec), 15°C for 45 sec, 20°C for 45 sec, 30°C for 45 sec, 40°C for 45 sec, 50°C for 45 sec, 65°C for 10 minutes, 95°C for 20 sec and 58°C for 1 minute. *Pfu* DNA polymerase PCR Master Mix was added to the amplified samples to bring the volume to 50µL total followed by 1µL of *Pfu* DNA polymerase. Samples were further amplified for 14 cycles using the following protocol: 94°C for 40 sec, 94°C for 20 sec, 59°C for 20 sec, 68°C for 7 minutes, with a final extension of 68°C for 7 minutes. Ethanol precipitation or the QIAquick PCR Purification Kit (28104, Qiagen) was used to purify the whole genome amplified (WGA) samples followed by storage at -20°C.

### Genomic DNA isolation from bulk tissue

High molecular weight (HMW) gDNA was isolated from 160mg of bulk LIBD75 frontal cortex tissue using the Monarch HMW DNA Extraction Kit for Tissue (T3060S, NEB) following the manufacturer’s instructions with the following changes to the lysis step. 40µL of 10 mg/mL Proteinase K (3115879001, Roche) was added to 580µL of Tissue Lysis Buffer. The tissue was placed at 56°C for 15 minutes on a ThermoMixer (Eppendorf) at 2000 rpm, then incubated at 56°C for 30 minutes without agitation.

### Library preparation and sequencing

#### ONT library preparation for a single MALBAC neuronal cell

A sequencing library consisting of a single-cell MALBAC-amplified product was prepared using the ONT Ligation Sequencing Kit (SQK-LSK109). 293ng of the WGA sample was end-prepped using the NEBNext UltraII End-Repair/dA-tailing module (E7546S, NEB) in a 60µl reaction. The end-prepped sample was then added to a 95µL ligation reaction (5µL T4 DNA ligase (M0202M, NEB), 25µL Ligation Buffer (LNB), 5µL Adapter Mix (AMX)) and rotated for 10 minutes at room temperature (RT). Next, 0.4X CleanNGS (CNGS005, Bulldog Bio) beads were added and incubated for an additional 10 minutes at RT with rotation. The library was placed on a magnet and the supernatant was removed. This was then cleaned with 2X Small Fragment Buffer (SFB) wash with resuspension following the addition of SFB. Finally, the supernatant was removed and the bead pellet was allowed to air dry for 30 seconds. The library was eluted in 16µL of Elution Buffer (EB) and 1µL was quantified using a Qubit (Thermo Scientific). The final library was prepared with 15µL adapted sample, 37.5µL Sequencing Buffer (SQB), 25.5µL Loading Beads (LB), and sequenced on a MinION R9.4.1 flow cell.

#### ONT library preparation for the first 20 neuronal cells

Sequencing libraries consisting of single-cell MALBAC-amplified products were prepared using the ONT Native Barcoding Expansion 1-12 kit (EXP-NBD104, ONT) as described here. (2.5-50ng) of each amplified product were end-prepped using NEBNext UltraII End-Repair/dA-tailing module in 10µL reactions. 10µL of end-prepped product was ligated with barcodes in a 25µL reaction with 1.5µL Native Barcode (NBD01-12), 6.25µL LNB, and 1.25µL T4 DNA ligase. 1µL 0.5M EDTA was added to stop the ligation. These ligation reactions were then pooled and incubated with 1X CleanNGS beads for 10 minutes at RT with rotation. The pooled, barcoded samples were then placed on a magnet and washed 2X with 70% ethanol. Following the washes, the sample was eluted in 65µL DNase/RNase-free water. Next, the eluate was added to a 100µL ligation reaction (5µL T4 DNA ligase, 25µL LNB, 5µL Adapter Mix II (AMII)). This reaction was rotated for 10 minutes at RT. 0.4X CleanNGS beads were added to the reaction and incubated for 10 minutes at RT with rotation. The library was placed on a magnet and the supernatant was removed followed by 2X SFB washes with resuspension after SFB addition. The supernatant was removed and the beads were air dried for 30 seconds. The adapted library was eluted in 16µL of EB and 1µL was quantified on a Qubit. The sequencing library was prepared with 15µL adapted sample, 37.5µL SQB, 25.5µL LB, and sequenced on a MinION R9.4.1 flow cell.

#### ONT library preparation for 95 additional neuronal cells

Sequencing libraries consisting of 95 additional single cell MALBAC-amplified products were prepared using the ONT Native Barcoding kit 24 V14 (SQK-NBD114.24, ONT) as described here with a slight modification. 400ng of each amplified product was end-prepped using the NEBNext UltraII End-Repair/dA-tailing module in 20µL reactions. 1µL of each of the samples was used to check the DNA concentration on the Qubit. The equimolar end-prepped products were ligated with in separate PCR tubes with barcodes in a 20µL reaction with 2.5µL Native Barcode (NB01-24), 5µL of LNB, and 2µL T4 DNA ligase for 20 minutes at RT with rotation. 2µL of 0.5M EDTA was added to stop the barcode ligation and pooled into a single 1.5mL microcentrifuge tube. The barcoded reactions were incubated with 1X CleanNGS beads for 10 minutes at RT with rotation. The samples were then placed on a magnet and washed twice with 700µL of 80% ethanol. Following the washes, the pooled samples were eluted in 36µL of water, and 1µL was used to quantify the DNA concentration on the Qubit. The Native Adapter (NA) was ligated to the samples in a 100µL ligation reaction (5µL T4 DNA ligase, 25µL LNB, 5µL NA) and rotated for 20 minutes at RT. Next, 0.4X CleanNGS beads were added and incubated for an additional 10 minutes at RT with rotation. The library was placed on a magnet and the supernatant was removed followed by two 125µL washes with SFB. After the final wash, the bead pellet was allowed to air dry for 30 seconds and the library was eluted in 16µL EB, and 1µL was used to quantify the DNA on the Qubit. The sequencing library was prepared with 300ng of barcoded-adapted sample, 37.5µL Sequencing Buffer (SB), 25.5µL Library Beads (LIB), and sequenced on a MinION R10.4.1 flow cell following the ONT MinION loading method.

#### Deep sequencing of 5 MALBAC WGA samples and 9203 single neuronal cell

The ONT barcoded library containing 5 single MALBAC WGA cells (9100, 9102, 9103, 9104, 9107) was loaded onto a PromethION R10.4.1 flow cell following the manufacturer’s instructions with a total of 250ng adapted library. The 9203 WGA sample was end-prepped using our protocol detailed above starting with 400ng of amplified DNA. The ligation sequencing preparation and PromethION loading followed the ONT protocols for SQK-LSK114 and R10.4.1 chemistry with 176ng of the adapted library.

#### Illumina NovaSeq sequencing of matched single-cell MALBAC WGA samples

94 MALBAC-amplified samples were prepared in a 96-well plate so that there was 3ng of a single neuron WGA sample per well. Samples were submitted to the University of Michigan Advanced Genomics Core for NovaSeq S4 300 cycle library preparation and sequencing.

#### Transposable Element nanopore Cas9-Targeted Sequencing (TEnCATS) library preparation and sequencing

TEnCATS library preparation was performed following McDonald et. al.^53^ with changes described here for SQK-LSK114 and R10.4.1 flow cells (see **Code availability**). 30µL of gDNA was dephosphorylated in a 40μL reaction with 6μL Quick CIP (M0525S, NEB) and 4μL 10X rCutSmart buffer (B7204S, NEB). This reaction was inverted and gently tapped to mix, and then incubated at 37°C for 30 minutes followed by a 2 minute heat inactivation at 80°C. The Cas9 ribonucleoprotein (RNP) was formed by combining 850ng of *in vitro* transcribed guide RNA, 1µL of a 1:5 dilution of Alt-R *S.p*. HiFi Cas9 Nuclease V3 (1081060, IDT), and 1X rCutSmart buffer (B7204S, NEB) in a total of 30µL. This reaction was incubated at RT for 20 minutes. Next, both the prepped gDNA and RNP were placed on ice and the RNP was added to the dephosphorylated gDNA. 1μL 10mM dATP and 1.5μL Taq DNA Polymerase (M0273S, NEB) were added to the gDNA:RNP reaction, then inverted and gently tapped to mix. This reaction was incubated at 37°C for 30 minutes for Cas9 cutting and brought to 75°C for a-tailing for 10 minutes. For adapter ligation, the cut reaction was transferred to a 1.5mL microcentrifuge tube. We then added 5μL T4 DNA ligase (M0202M, NEB) and 5μL Ligation Adapter (LA; SQK-LSK114, ONT). This reaction was inverted to mix and incubated at RT for 20 minutes with rotation. Following ligation, we added 1 volume of 1X Tris EDTA (TE) and inverted to mix. Next 0.3X Ampure beads (SQK-LSK114, ONT) are added and incubated for 5 minutes with rotation followed by 5 minutes at RT without rotation. The beads were then washed twice with 150μL Long Fragment Buffer (LFB; SQK-LSK114, ONT) followed by incubation with 20-50μL Elution Buffer (EB; SQK-LSK114, ONT) at 37°C for 30 minutes. Finally, we loaded the R10.4.1 MinION flow cell following the ONT protocol using 12μL of the library and sequenced for 72 hrs on a MinION. For sequencing of the *Alu* libraries on PromethION flow cells, 12μL of the same library sequenced on a MinION was used and brought to a total volume of 32μL with EB then sequenced for 72 hours on a PromethION2 Solo.

#### 10x Genomics linked-read and Illumina NovaSeq library preparation and sequencing on bulk tissue

10x Genomics linked-read and Illumina NovaSeq sequencing on bulk tissue is described in Garrison et al.^64^ and briefly outlined here: Genomic DNA was isolated using the MagAttract High Molecular Weight DNA Kit. 1 -5ug gDNA aliquots were used to generate both 10x Genomics linked-read sequencing libraries and Illumina short-read sequencing libraries. 10x libraries were sequenced on the 10x Chromium platform to 53x, and the Illumina library was sequenced on the NovaSeq 6000 platform to 30x.

### Basecalling and alignment

Data was basecalled and aligned to hg38 human reference genome (GCA_000001405.15_GRCh38_no_alt, https://ftp.ncbi.nlm.nih.gov/genomes/all/GCA/000/001/405/GCA_000001405.15_GRCh38/seqs_for_alignment_ pipelines.ucsc_ids/) using dorado 7.2 (dna_r10.4.1_e8.2_400bps_sup@v5.0.0) with CG methylation calling and a minimum qscore of 9.

TEnCATs data was basecalled using dorado 7.2 (dna_r10.4.1_e8.2_400bps_sup@v5.0.0) with CG methylation calling and a minimum qscore of 9. Reads were aligned to hg38 (same as above) alongside basecalling with dorado using the “map-ont” preset.

Single-cell sequencing was performed for up to 168 hours. Data was basecalled using Guppy v6.2.11 or 6.4.6 (ONT) using the high accuracy mode (dna_r10.4.1_e8.2_400bps_hac) with a minimum qscore of 9. Chimeric MALBAC reads were split using duplex_tools (ONT) where native adapters were replaced with the MALBAC primer. Next, reads were aligned to hg38 using minimap2^94^ v2.26-r1175.

### Donor-specific genome assembly

The initial haploid genome assembly was generated using Shasta v0.11.1^58^ from ONT WGS data. Both Flye^95^ v2.9.2 and Shasta were utilized for draft assembly generation. Shasta achieved nearly a 10 -fold increase in speed while yielding results comparable to Flye, consistent with previous findings^58^ highlighting Shasta’s superior speed and resource efficiency. Subsequently, we chose the draft haploid assembly from Shasta and proceeded to generate the diploid assembly. First, the original ONT reads were realigned to the haploid draft using minimap2^94^ v2.28. HapDup^60^ v0.12 was then employed to convert the haploid draft to a diploid format by constructing haplotypes from the realignment. Two versions of the assembly were generated: a dual assembly, possessing the same continuity as the original diploid assembly with potential phase switches and optimized for variant calling, and a more fragmented phased assembly that contains haplotype-resolved contigs without switch errors. Hapo-G^96^ v.1.38 was used three times sequentially and polished the both genome assembly versions using Illumina short-reads. Contigs from two haplotypes are annotated and merged into one file to be processed. The quality assessment of the final polished diploid assemblies was performed thoroughly by four pipelines: BUSCO^97^ v5.7.1, QUAST^98^ v5.2.0, Merqury^70^ v.1.3, and a customized pipeline to assess the recall rate of heterozygous phased SNVs from linked-read data (see **Code availability**). BUSCO and Merqury were used to evaluate assembly completeness and accuracy, while QUAST analyzed key assembly metrics (**Supplementary Table 2**), e.g. NG50 and number of contigs. For BUSCO, we used the *primates_odb10* dataset to assess the completeness of the human genome assemblies. The linked-read SNV recall rate was evaluated by examining each heterozygous phased SNP identified in the 10x linked-read data against the corresponding locus in the assembly. For each SNP, we assessed whether the nucleotides from the two haplotypes of the assembly at the given locus accurately matched the alternate alleles reported in the 10x data. The recall rate was calculated as the percentage of heterozygous phased SNPs for which both alleles were correctly identified in the assembly. We chose to use the phased assembly (instead of the dual assembly) with haplotype-resolved contigs in the downstream analysis.

### Refinement of phased SNVs and phase blocks

Heterozygous phased SNVs were called from the phased assembly using Phased Assembly Variant Caller (PAV)^3^ v2.3.4. To obtain a conservative set of high-confidence heterozygous SNVs for phase block extension, we examined the VAF of each SNV in Illumina WGS data. Any SNV with a VAF below 0.2 was excluded. The refined SNV callset was then used to extend the phase blocks. We obtained linked phase blocks from both the phased assembly and linked-read data, enabling the correction of potential phase-switch errors within each extended block. The phase block information from the phased assembly was provided by Hapdup with the start and end positions for each haplotype-resolved contig. The information from linked-reads was extracted based on the positions of the first and last SNVs within the same phase block reported by LongRanger ^99^ v2.2. Blocks lacking heterozygous phased SNVs from the phased assembly were excluded from further analysis. The phase blocks from two data sources were merged and phase switches were identified by comparing the genotypes of matching SNVs at the same locus between two data sources. Specifically, a phase switch was pinpointed when discordance was observed between two bridged phase blocks from the phased assembly and the corresponding connecting linked-read phase block. Subsequently, the haplotype information of all the SNVs in the second bridged phase block was flipped until another switch was detected. Merged phase blocks with no matching SNVs were subdivided into smaller segments confidently free of phase switches. Finally, a set of heterozygous SNVs with phasetag information was reconstructed from the phase blocks and utilized for the downstream phasing process.

### Phasing

We used HaploTaglr^62^ to phase the long-read sequencing data. HaploTaglr assigns haplotags to long sequencing reads based on a multinomial model and existing phased variant lists, incorporating a basic error model to control the empirical false discovery rate (FDR) in its output. We built up a haplotype assignment pipeline for short-reads based on the reconstructed phased heterozygous SNVs from the phase blocks. For each read overlapping a phased heterozygous SNV locus, an initial haplotype was assigned based on the allele at the locus on the read. We first assigned an initial haplotype to each read based on the SNVs it contains across the genome, then applied the following rules to determine the final haplotype for each read pair: 1) Consistent Haplotype Agreement: If all overlapping phased heterozygous SNVs for a read pair agreed on one haplotype, that haplotype was assigned to the pair. 2) Discrepant Haplotype Assignment: If the two reads in a pair had different haplotypes, the pair was classified as unphased. 3) Unmapped Reads: If one read in a pair was assigned a haplotype and the other read was unmapped, the entire read pair was considered unphased.

### Genetic variant calling

#### 10x Genomics Linked-reads

We used LongRanger^99^ v2.2 to analyze the linked-read sequencing data from 10x Genomics for the LIBD75 bulk tissue. Phase block information was derived from the phased linked-read SNV callset directly and proceeded into the phase block extension process. The heterozygous SNVs from the phased SNV callset were also used for assembly assessment analysis.

#### Phased assembly

We utilized PAV^3^ v2.3.4 to identify SNVs, indels, and SVs of the phased assembly in comparison to the reference genome. We retained all variants labeled as "SNV" from the PAV callset for downstream analysis as SNVs. Variants tagged as"DEL" and "INS" were categorized as either indels or SVs based on a length cutoff of 50 base pairs. Variants tagged as "INV" were categorized as inversions. Additionally, SNVs and indels overlapped with any defined SVs were excluded. Indels within tandem repeat regions and homopolymer regionswere excluded. We implemented two pipelines to identify TEs from the phased assembly. First, we annotated the insertion (INS) sequences reported from PAV using RepeatMasker^100^ v4.1.2. Further refinement was conducted based on subfamily information such as L1Hs, *Alu*Y, and SVA, with an additional criterion of a minimum 6 bp polyA tail length to confirm active TEs. Second, we used PALMER^3,51^ v2.0.1 to identify TEs (LINE-1, *Alu*, SVA) directly from the phased assembly with the ‘--mode asm’ option. We collected the calls from the annotated PAV callset that intersected with the PALMER assembly callset, resulting in a high-confidence, assembly-based TE callset. For the downstream TEnCATS recall rate analysis, we only used the subset of L1Hs, *Alu*Ya5, and *Alu*Yb8 from the assembly-based TE callset.

#### Bulk tissue

We utilized DeepVariant^101^ (v1.6.0) model, Clair3^102^ (v1.0.6) model r1041_e82_400bps_sup_v410, and ClairS-TO (v0.0.2, https://github.com/HKU-BAL/ClairS-TO) model ont_r10_dorado_sup_4khz to identify SNVs from the ONT WGS sequences in the bulk tissue. SNVs from Illumina WGS sequences were identified using GATK Mutect2^103^ (v4.3.0), DeepVariant (v1.6.0) WGS model, and ClairS-TO (v0.0.2) ilmn model. GATK Mutect2, Clair3, and DeepVariant VCFs were processed to retain only those with ’FILTER = PASS’, and ClairS-TO callsets were filtered for using either ’NonSomatic’ or ’PASS’.

We utilized DELLY2^104^ (v1.2.6) (lr mode) and Sniffles2^52^ (v2.4) (default mode) to generate SV callsets for ONT bulk tissue data. For Illumina bulk data, we employed DELLY2 (v1.2.6) in its default ‘--call’ mode, and Manta^105^ (v1.6.0) (default mode) to generate the SV callsets. We pooled the category "DUP" from DELLY2 with "INS" as “INS/DUP” to facilitate the comparison with the output of other tools. For Manta, based on the diploidSV.vcf, we then applied the provided function convertInversion.py to reformat INV out of reported breakends to get the final callset. We used PALMER^3,51,53^ (v2.0.1) and xTea^72^ (v0.1.0 xTea_long_release) to identify TEs from the ONT WGS sequences in the bulk tissue. In the PALMER callset, TE calls were required to have at least one high-confidence supporting read. Additionally, SVA calls were refined to ensure ‘start_inVariant’ ≥ 420 and ‘end_inVariant’ ≥ 1355. For Illumina WGS data, xTea (v0.1.9) and MELT^106^ (v2.2.2) were used to detect TEs, utilizing their built-in consensus library. Calls with a "PASS" tag were selected for downstream analysis.

We used MosaicForecast^79^ to identify mosaic SNVs from Illumina WGS data. Following MosaicForecast’s guidelines, we processed the Mutect2 output as the input file and executed the steps of "extracting read-level features" and "genotype prediction" using the 50xRFmodel_addRMSK_Refine.rds model. Variants labeled as "mosaic" in the final call set were selected for downstream analysis. Sniffles2 somatic mode with default parameters was used to identify somatic SVs from ONT bulk tissue data. We relaxed the cutoff for the number of supporting reads in PALMER to detect both somatic and germline TE insertions. For somatic TEs, we derived PALMER mosaic calls by masking TE calls from the assembly-based callset if they shared the same TE subfamily and insertion orientation, classifying the remaining calls as potential somatic TE calls.

#### TEnCATS

For both L1 and *Alu* datasets, on-target rate, and TE calling were characterized using NanoPal (see **Code availability**), adapted from our prior study^53^. Briefly, reads were aligned to the reference genome and non-reference TE insertions were detected with PALMER. Next, TEnCATS reads were then classified into on-target using BLASTn and reads supporting the MEIs are clustered by location. Variant calls with fewer than two read support were filtered from the final results.

We compared the TEnCATS putative TEs with TE callsets derived from assembly, bulk ONT WGS, and single-cell ONT sequencing. The recall rate for germline TEs was determined by intersecting with the assembly-based germline callset (see above). We applied a semi-automatic pipeline to stringently refine somatic TEs after gathering somatic signals by excluding germline calls. The candidate list was processed through the pipeline using the following steps: a, Candidates should not be located in low-quality regions, including any segmental duplicates identified by UCSC, low-confidence mask regions used in this project (see below), and reference *Alu* or LINE-1 repeat regions from RepeatMask for *Alu* or L1Hs candidates, respectively; b. Candidates should contain only the signals associated with the designated TE sequence segment and subfamily^53^ and should not include any genomic content that is not TE-related sequence, such as polyA tracts, target site duplications, or potential transduction sequences between two polyA tract signals^51^; c. Candidates should be inserted in a region containing a conserved endonuclease site motif (TTTT/AA). We used BLAT^107^ and IGV^108^ for the final manual inspection process and concluded a final list. We did not observe any overlap with the single-cell putative somatic TEs.

#### Single cells

We developed GARLIC and PalmeSom (see below) to identify somatic CNVs and TEs in the single -cell ONT sequences, respectively. In single-cell Illumina sequences, we utilized an adapted version of Ginkgo^21,50^ for identifying somatic CNVs. We used xTea v0.1.9 and MELT v2.2.2 for TEs with default parameters.

#### GARLIC

We developed a pipeline called GARLIC (Genome-wide Allelic copy numbeR variation Locator In Cells, https://github.com/WeichenZhou/GARLIC) for identifying large somatic deletions in single neurons using long-read sequencing data, based on the tool from our previous study^21^. GARLIC leverages haplotype information by introducing a statistic called physical phase coverage (PPC). By using PPC, GARLIC minimizes the effects of PCR bias introduced by single-cell DNA amplification. The concept of PPC involves calculating the proportion of the physically covered genome by any reads, rather than read coverage, and producing a separate PPC for the two individual haplotypes. Subsequently, GARLIC calculates the log_2_ ratio of PPC between the two haplotypes and derives an absolute value of this log_2_ ratio, termed R_ppc_. GARLIC then segments the genome into small bins, dynamically selecting bin sizes based on regions covering an arbitrary 100 SNPs in a single phase block. It filters out low-confidence regions, identified as bad bins, using the mask file in this study (see **Data availability**).

GARLIC calculates the R_ppc_ value for each bin across the genome, and implements a Circular Binary Segmentation (CBS)^109^ algorithm to segment the R_ppc_ signals and determine copy number variations in each bin as follows:

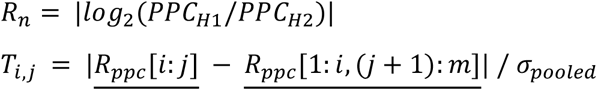

where "i" and "j" represent potential changepoint locations within the data, "R" is the data vector for the ratios of physical phase coverage across bins in the genome, and σ is the pooled standard deviation; the algorithm searches for the maximum value of T(i,j) across all possible combinations of i and j, signifying the most significant changepoint location.

GARLIC filters out germline CNVs using existing data sets^3,81^, refining a set of candidate somatic CNVs. Lastly, GARLIC generates curve plots for both R_ppc_ and sequence coverage to facilitate further manual inspection.

#### PalmeSom

We developed an enhanced version of PALMER, PalmeSom (https://github.com/mills-lab/PALMESOM), to identify TEs from ONT single-cell sequences. PalmeSom incorporates three fundamental steps : a) Initial calling and information merging: it implements modules in PALMER to identify *Alu*, L1, and SVA signals from each single cell. To further facilitate the analysis of putative somatic TE signals, PalmeSom merges TE signals as putative insertion positions from each single cell into a large data frame. Centered around the putative insertion positions, a bin of ±25 bp was used to examine all reads across single cells within bins to determine the presence of TEs in each cell. Phase information, mapping quality, CIGAR, and coordinates in consensus TE sequence are also recorded. b) Read categorizing: based on the TE signal and read information, PalmeSom categorized reads with signal into three types: read-through, left-side soft-clipped supportive, and right-side soft-clipped supportive reads. In addition, read-through reads with no TE signal are documented as non-signal supportive reads. TE signals in the supportive reads were considered based on the coordinate range in the TE consensus sequences reported by our previous study^51^. A dedicated module to exclude reads with the false positive signal introduced by misalignment and genomic rearrangement, e.g. large deletion, is also implemented in this step. For each insertion position, PalmeSom tallies the number of right-side, left-side soft-clipped supportive reads, read-through supportive reads, and read-through non-supportive reads in all haplotypes (h1, h2 and non-phased) for each single cell. c) Summarization: PalmeSom reports the putative somatic TE calls in different tiers.

In this project, we analyzed 121 cell samples and focused on putative TE calls with at least 5 supportive reads. To define a high-confidence TE callset, we applied criteria requiring a) read-through supportive reads from the same haplotype greater than zero or supportive read pairs from the same haplotype (with both left-side and right-side directions) greater than one, and b) the count of cell with supportive signals greater than one. In addition, we annotated the callset with additional information, including RepeatMasker and the Segmental Duplication track from UCSC genome browser^110^. Based on the high-confidence TE calls, we further filtered somatic calls by requiring AF of the call in ONT bulk tissue less than 20%, sum of supporting reads equal or less than ten, not reported by other population-level studies^3,81^, and at least one non-signal supportive read from at least one cell in both haplotypes.

### Refinement of genetic variants

We merged the low-confidence region information from HGSVC^3^ and ENCODE^111^ as a universal mask file in this study to mask regions where genome assembly results in erroneous signal or variant calls were found to be difficult to reproduce. All variants within this mask file and sex chromosomes were removed. In addition, we applied de-redundancy filters^3^ and filters^112,113^ of homopolymer and low-confidence regions for indels. In addition, we applied a low-confidence region mask file from Genome In A Bottle^114^ as well as a customized mask file derived from the LIBD75 phased assembly indicating the gap regions among assembled contigs when working on the comparison analysis of the germline recall rate (**Supplementary Table 4**). Relevant files are listed under **Data Availability**.

### Integration of the genetic variant callsets

For non-somatic callsets in bulk tissue, we used assembly-based calls as the primary set and overlapped other callsets onto it to generate a unified set. For SNVs, we considered calls as identical when both their position and alternative allele matched exactly. For insertions (INS), we merged calls by applying an open window at the insertion site, based on the data source: ±10 bp for assembly contigs, ±30 bp for long-reads, and ±50 bp for short-reads. Additionally, the length difference between the insertion from another tool and the primary call needed to be less than 50% of the longer insertion. For deletions (DEL) and inversions (INV), we required at least a 50% reciprocal overlap between two calls for merging. For transposable elements (TEs), we used the same open window extension as for insertions and ensured that the TE family and insertion orientation matched.

For somatic bulk tissue callsets, we used the callsets from somatic callers as the primary reference and overlapped assembly-based variants with them. We applied the same strategy as described above to intersect candidate somatic calls from MosaicForecast for SNVs, Sniffles2 mosaic model for SVs, and PALMER mosaic calls for TEs.

### Generating high-confidence assembly-based callsets

To produce high-confidence contig-based callsets, we assessed the variant allele frequencies (VAFs) of the assembly-based calls in both ONT and Illumina bulk tissue data. We obtained these VAFs by utilizing non-somatic variant detection tools and assigning their values to the corresponding calls. For analyzing VAFs in ONT bulk tissue sequences for assembly-based variant calls, we approached SNVs by randomly selecting a representative VAF from the values reported by Clair3, ClairS-TO, and DeepVariant. For SVs, representative VAFs were randomly chosen from the frequencies reported by DELLY2 and Sniffles2. For transposable elements (TEs), we manually calculated VAFs by randomly using the support provided by PALMER or xTea_long. Specifically, the VAF was derived by dividing the number of potential supporting reads, reported by PALMER or annotated by BLASTn^115^ at the loci by xTea_long, by the mean sequencing coverage of a ±30 bp window around the variant, as determined by samtools coverage. In Illumina bulk tissue sequences, the VAFs for SNVs were randomly selected from the values reported by Clair3, ClairS-TO, DeepVariant, and GATK Mutect. For SVs, the VAFs were randomly selected from either DELLY2 or Manta. For TEs, we randomly chose VAFs from the MELT and xTea callsets. We then plotted the VAF distribution in ONT and Illumina data for each variant type (see **Supplementary Figure 3**). From this analysis, we identified that the transition points between the first and second peaks in the distribution occur at VAF = 0.2 for both SNVs and SVs, and VAF = 0.15 for TEs. Accordingly, we set these as our cutoff VAFs. Variants with VAFs exceeding the cutoff in either ONT or Illumina are categorized as high-confidence germline variants, while those with VAFs below the cutoffs in both platforms are classified as potential somatic variants.

### Recall rate calculation in bulk tissue and pooled single cells

#### Construction pseudo-bulk samples from pooled single cells

We used samtools merge to create pooled BAM files from single-cell data. Specifically, we pooled the single cells based on their sequencing platforms: all 115 cells sequenced by MinION flow cells were merged, and the 5 cells sequenced by one PromethION flow cell were merged separately. Additionally, the BAM file for cell 9203, sequenced by one PromethION flow cell, was analyzed individually. These two pooled BAM files, along with the individual BAM file for cell 9203, were treated as three pseudo-bulk samples. These pseudo-bulk samples were used to generate germline integrated non-somatic SV and TE callsets using the same tools employed for ONT sequences.

#### Calculating germline recall rate

For bulk tissue, we used high-confidence assembly-based germline callsets as the reference set to calculate the recall rates of SNVs, SVs, and TEs detected by at least one non-somatic variant detection tool from ONT and Illumina bulk tissue data. For the pseudo-bulk, we performed the same analysis for SVs and TEs. However, for SNVs, we used samtools mpileup to check for the presence of the alternative allele signal at the coordinates of the germline SNVs to determine the recall rate. Furthermore, we recorded the recall rates under various masking conditions (**Supplementary Table 4**). In the single-cell sequences, due to the sparsity of the data, we were unable to calculate the recall rate for SVs. For TEs, we used the high-confidence assembly-based germline TE callset as the reference set and calculated the recall rate for each individual cell detected by PalmeSom.

### VAF Calculation in individual haplotypes in bulk tissue data

#### Construction of split phased BAM files

For the phased ONT and Illumina bulk tissue data, we extracted reads tagged with HP:Z:1 as haplotype 1 and HP:Z:2 as haplotype 2, creating two split BAM files. The original header was then added to each of the split BAM files to retain metadata consistency.

#### Calculation of VAF in different haplotypes

We first calculated the VAFs in the callsets from non-somatic callers. For SNVs, we used samtools mpileup to determine the alternative allele read count and total depth at specified coordinates in the haplotype 1 and haplotype 2 BAM files for both ONT and Illumina bulk tissue data. VAFs were then calculated by dividing the alternative allele read count by the total depth. For SVs, we applied Sniffles2 and DELLY2 to the split ONT BAM files separately. To determine the VAF for each haplotype, we prioritized the VAF reported by Sniffles2 and used the VAF provided by DELLY2 only when Sniffles2 did not detect the variant. For TEs, we ran xTea_long and PALMER on the split ONT BAM files and calculated the VAFs using the same method as calculating AF in the WGS data, after merging the callsets from xTea_long and PALMER for each haplotype.

For the VAFs in the callsets from somatic callers, we applied the same strategy above for the callsets generated by MosaicForecast for SNVs, and Sniffles2 (somatic mode) for SVs. For TEs, we ran PALMER on the split ONT BAM files, extracted the PALMER mosaic callsets, and calculated the VAF for each haplotype. Supporting reads were counted from intermediate files^51^ reported by PALMER, and the final VAF was computed by dividing the supporting read count by the read depth obtained using samtools coverage. For all genetic variants, we filtered out calls in regions with a read depth of less than five in either haplotype to exclude the drifting effect introduced by small numbers and false negatives caused by the sparsity of reads. Haplotype A was defined as the haplotype with the highest alternative allele frequency, while Haplotype B was defined as the other haplotype. Additionally, we intersected the callsets with the assembly-based callsets using the same merging strategy mentioned in the previous section.

To refine candidate somatic variants in bulk tissue data, we established an empirical cutoff line at *y* = 0.2*x*. As an example, for a heterozygous variant supported by ten reads in Haplotype A, it is permissible to have up to three reads in Haplotype B due to errors, such as phasing errors. If the number of reads in Haplotype B exceeds three, we consider it unlikely to be a phasing error. Generally, if the ratio of Haplotype A and Haplotype B exceeds the cutoff line in heterozygous calls, it could be indicative of other types of errors, leading to false positives (*hapErrors* or *mapErrors*). To reject calls where the frequency in Haplotype A exceeds 80%, we also established a cutoff at *x* = 0.8 to exclude calls, as these could be false positive *hapErrors*.

### Genomic annotation

To examine the potential enrichment in repetitive genomic regions or bias from reference repeat classes, we annotated the callsets with genomic information, including RepeatMasker^100^ and the Segmental Duplication track from UCSC genome browser^110^.

For the analysis for putative somatic callsets from ONT bulk WGS, we also added tandem repeats finder (TRF) annotations^116^ and performed an overlap analysis using pyranges^117^. We extended the coordinates of all somatic calls by a slack value of 300 bp on each side for TRF annotation. We used fractional counting in events where somatic calls overlapped multiple RM annotations for GRCh38. We calculated the expected counts under the null hypothesis that the counts would be distributed based on the slack-corrected proportion of each TRF or RM class falling outside the low-confidence regions. We employed Fisher’s Exact Test to evaluate enrichment of somatic variant overlap with various reference repeat classes.

### Manual inspection for candidate somatic calls

We utilized IGV^108^ to inspect the read distribution, genomic content, and annotations for all candidate somatic calls in bulk and single cells (see **Supplementary Table 5-9**). For large deletions, we also leveraged the sequence coverage and PPC plots reported by GARLIC for further evaluation. All manual inspections leveraged the phasing information from DSA.

## Data availability

This study was conducted as part of the NIH Common Fund consortium initiative, Somatic Mosaicism across Human Tissues (SMaHT). The long-read WGS, TEnCATS, and MALBAC sequencing data can be found in SRA under BioProject: <u>PRJNA1291649.</u> Illumina NovaSeq and 10x Genomics linked-read WGS data can be found in the BSMN data resouces^64^.

The assemblies, phase blocks, mask file, and callsets from different technologies can be found at Zenodo https://doi.org/10.5281/zenodo.15733390118.

We used GRCh38 (GenBank accession: GCA_000001405.15) as the primary reference in this project. The reference can be downloaded at https://ftp.ncbi.nlm.nih.gov/genomes/all/GCA/000/001/405/GCA_000001405.15_GRCh38/seqs_for_alignment_pipelines.ucsc_ids/. The CHM13-T2T reference (GenBank accession: GCA_009914755.4) used in the assembly assessment can be found at https://www.ncbi.nlm.nih.gov/datasets/genome/GCF_009914755.1/.

The filters we applied for indels can be found here: de-redundancy filters (https://ftp-trace.ncbi.nlm.nih.gov/ReferenceSamples/giab/release/genome-stratifications/v3.5/GRCh38@all), homopolymer (../LowComplexity/GRCh38_AllTandemRepeatsandHomopolymers_slop5.bed.gz), and low-confidence regions in chrX (/XY/GRCh38_chrX_XTR.bed.gz and ../XY/GRCh38_chrX_ampliconic.bed.gz).

## Code availability

The scripts in this project can be found at https://github.com/mills-lab/LIBD75.

PALMER: https://github.com/WeichenZhou/PALMER

GARLIC: https://github.com/mills-lab/GARLIC

PalmeSom: https://github.com/mills-lab/PALMESOM

TEnCATS:

For the molecular protocol, https://dx.doi.org/10.17504/protocols.io.kqdg3q66ev25/v1

For NanoPal, https://github.com/Boyle-Lab/NanoPal-Snakemake

## Acknowledgments

We thank the Brain Somatic Mosaicism Network (BSMN) for providing the NovaSeq and 10x linked-read bulk tissue sequencing data for LIBD75. Library prep and Illumina NovaSeq sequencing of MALBAC libraries was carried out in the Advanced Genomics Core at the University of Michigan. W.Z. was partially supported by and provided salary support from the NIH/NIA-funded Michigan Alzheimer’s Disease Research Center (P30AG072931) and the University of Michigan Alzheimer’s Disease Center Berger Endowment. C.M. and B.B. were supported in part by the National Institute of Health training grant T32 [HG000040]. This research is supported by the NIH Common Fund, through the Office of Strategic Coordination/Office of the NIH Director under award UG3NS132084 to M.J.M., A.P.B., and R.E.M, and UH3NS132084 to A.P.B. and R.E.M. This research was supported by the National Institutes of Health (NIH) under awards R21HG011493 to A.P.B. and R.E.M. and R03AG087485 to W.Z.

## Author information

### Author Contributions

R.E.M., A.P.B., M.J.M., and W.Z. conceived the project. M.J.M. isolated single neurons from bulk tissue and prepared MALBAC libraries. P.D.O. helped prepare MALBAC libraries and aliquoted sections from the LIBD75 donor brain tissue. C.M., T.L.M., and J.A.S. performed the gDNA extractions and Cas9 targeted enrichment and nanopore sequencing. C.M. and J.A.S. performed nanopore sequencing of the MALBAC libraries and bulk tissue WGS. J.A.S. prepared and submitted the MALBAC libraries for Illumina NovaSeq sequencing. J.W. and W.Z. constructed the personalized assembly. Y.G. and W.Z. performed the genetic variant calling. W.Z. developed the GARLIC pipeline. W.Z. and Y.G. developed the PalmeSom software. W.Z., Y.G., J.W., C.M., K.K., S.J.L, and B.B. performed computational analysis. W.Z., Y.G., J.W., C.M., and B.B. did the visualization. All authors guided the data analysis strategy. W.Z., C.M., Y.G., J.W., J.A.S., R.E.M., and A.P.B. wrote the manuscript. All authors reviewed and edited the manuscript. All authors approved the final manuscript.

## Supplementary information

**Supplementary Figure 1.** Percent genome covered for long-read (ONT MinION and PromethION) and short-read (Illumina NovaSeq) WGS from MALBAC amplified single-cell DNA.

**Supplementary Figure 2.** Assessment of assemblies (Dual and Phased) compared to reference genomes (HG38 and CHM13-T2T). The left panel shows the number of single-copy complete BUSCOs (expected marker genes present as single copies), while the right panels display duplicated (marker genes present more than once), fragmented (partially recovered marker genes), and missing BUSCOs (marker genes not detected).

**Supplementary Figure 3.** Variant allele frequency of assembly-based germline variants in ONT bulk and Illumina bulk WGS sequencing, dash lines indicate the cutoffs. **a**, SNV. **b**, SV. **c**, TE.

**Supplementary Figure 4.** Germline variant analysis in pseudobulk from single cell sequences. **a**, Germline recall rates across different libraries and platforms. Variant recall rates by yield using the various mask files across ONT WGS, Illumina WGS, and pseudobulk MALBAC data. **b**, Allele frequency of assembly-based germline variant distributions in pooled MinION, PromethION and NovaSeq sequencing. Main mask file: the low-confidence region information from HGSVC^3^ and ENCODE^111^; GIAB mask file: a low-confidence region mask file from Genome In A Bottle^114^; In-house mask file: a list of regions derived from the LIBD75 phased assembly indicating the gaps among assembled contigs.

**Supplementary Figure 5.** Two-dimensional kernel density plots of allele frequency of assembly-based germline homozygous variants (left) and heterozygous variants (right) in ONT WGS bulk tissue sequences. **a**, SNVs in Illumina. **b**, SNVs in ONT. **c**, SVs in ONT. **d**, TEs in ONT.

**Supplementary Figure 6.** Two-dimensional kernel density plots of allele frequency of putative somatic variants which overlapped with assembly-based potential somatic variants (left) and bulk somatic caller-specific variants (right). **a**, SNVs, **b**, SVs, **c**, TEs.

**Supplementary Figure 7.** Enrichment analysis of all refined putative somatic candidate classes, including deletions, insertions, Alu, L1 and SVA, and false positive somatic calls through tandem repeats finder (TRF) and RepeatMasker (RM) annotation. **a**, Enrichment of putative somatic SV candidates by TRF annotations. **b**, Enrichment of putative somatic TE candidates by RM annotations. **c**, Enrichment of putative somatic SV candidates by RM annotations. **d**, Enrichment of putative somatic TE candidates by TRF annotations. *** denotes a p-value of less than 0.0001, and ** corresponds to a p-value of less than 0.001 by Fisher’s Exact Test.

**Supplementary Figure 8.** Schematic illustration of haplotype-aware detection in single neurons. **a**, Somatic CNVs by GARLIC. **b**, Somatic TEs by PalmeSom.

**Supplementary Table 1.** Data generation. **Sheet1**, MALBAC read statistics by single cell. Read stats, alignment stats, and sequencing details for each MALBAC library. **Sheet2**, Bulk tissue read statistics by acquisition.

**Supplementary Table 2.** Assembly statistics. **Sheet1**, General assembly statistics for both dual and phased assemblies, including total size, number of contigs, N50, NG50, genome fraction covered by the assembly, linked-read SNV recall rate, QV and k-mer completeness from Merqury. A dual assembly refers to the creation of two distinct assemblies representing both haplotypes of the diploid human genome. These assemblies maintain the same contiguity as the draft assembly, with each contig assigned to one haplotype, with potential phase switches within individual contigs. In contrast, a phased assembly, also known as a haplotype-resolved assembly, ensures accurate phasing of each haplotype from the dual assembly. This process eliminates phase switches within contigs, but results in a more fragmented assembly. **Sheet2**, Quality metrics acquired from QUAST for both dual and phased assemblies. **Sheet3**, Quality metrics acquired from Merqury for both dual and phased assemblies.

**Supplementary Table 3.** Application of existing and novel variant discovery tools across multiple sequencing platforms and assays.

**Supplementary Table 4.** Recall rates of germline genetic variants under different mask regions. Recall rates of high-confidence assembly-based germline variants for unified callsets from **Sheet1** ONT WGS and Illumina WGS, **Sheet2** TEnCATs, and **Sheet3** pooled single-cell, under different mask regions. The table summarizes the total number of high-confidence assembly-based germline variants and the corresponding recall rates for each sequencing technology. Applied filters include masks from Encode and HGSVC, in-house, GIAB, and their combinations.

**Supplementary Table 5.** Curation of 30 false positive SNV somatic mosaicisms by *hapErrors*.

**Supplementary Table 6.** Curation of 15 false positive SNV somatic mosaicisms by *seqErrors*.

**Supplementary Table 7.** Curation of 30 false positive SV somatic mosaicisms by *mapErrors*.

**Supplementary Table 8.** Curation of 40 false positive TE somatic mosaicisms by *mapErrors*.

**Supplementary Table 9.** Non-reference germline and putative somatic TEs captured by TEnCATs in bulk tissue with annotations. **Sheet1**, Germline TEs captured by TEnCATs. **Sheet2**, Candidate somatic TEs captured by TEnCATs with manual inspection.

**Supplementary Table 10.** Somatic DEL candidates reported by Gingko from single-cell Illumina sequencing and GARLIC from single-cell ONT sequencing. **Sheet1**, Somatic DEL candidates reported by Gingko. **Sheet2**, Somatic DEL candidates reported by GARLIC. **Sheet3**, Intersection set of Ginkgo and GARLIC.

**Supplementary Table 11.** Somatic TE candidates reported by PalmeSom from single-cell ONT sequencing with annotations.

