## Supplementary material for "A personalized multi-platform assessment of somatic mosaicism in the human frontal cortex": Supplementary Table 5.pdf

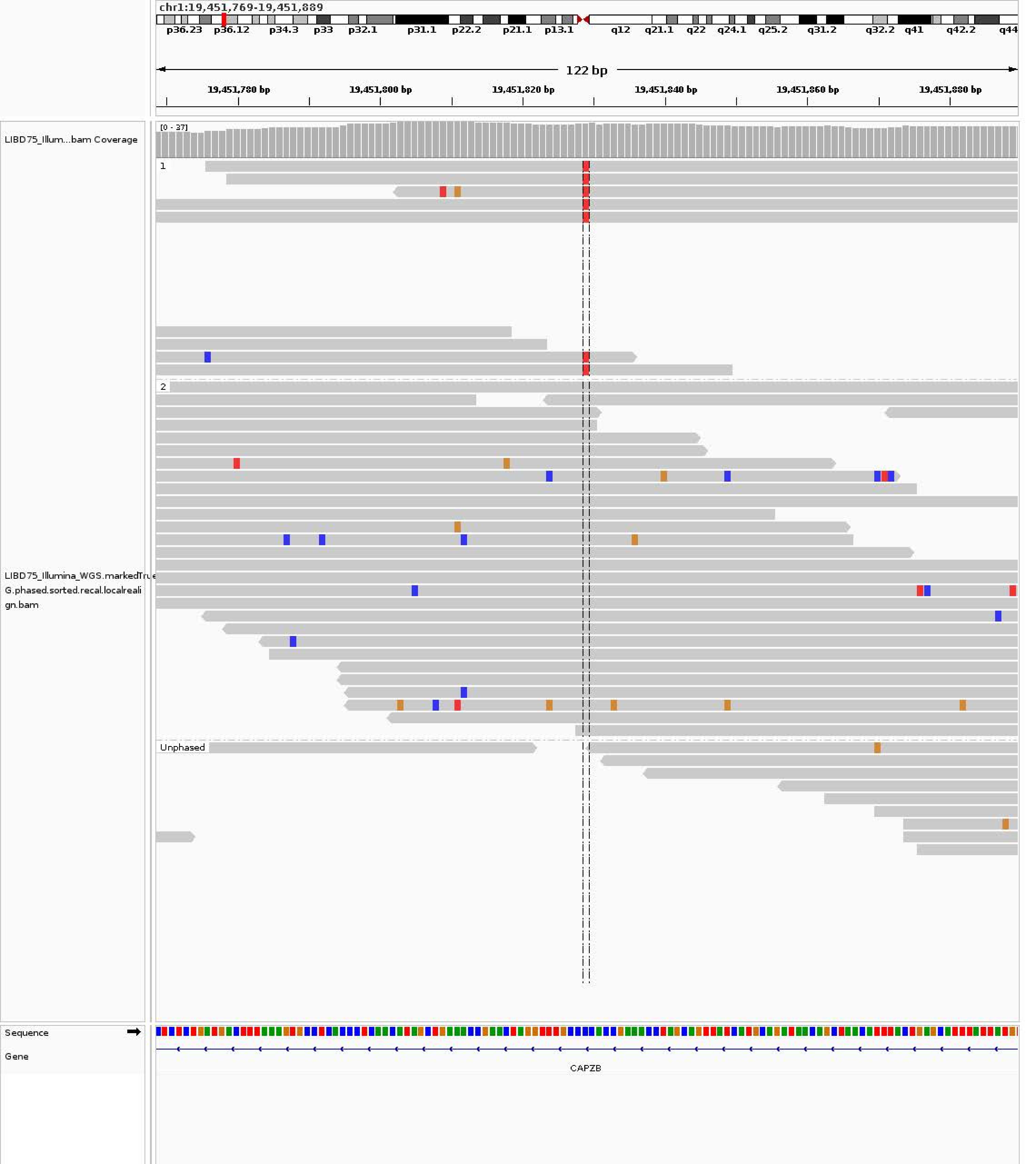
chr1:19,451,769-19,451,889
p36.23 p36.12 p34.3 p33 p32.1 p31.1 p22.2 p21.1 p13.1 q12 q21.1 q22 q24.1 q25.2 q31.2 q32.2 q41 q42.2 q44
122 bp
19,451,780 bp
19,451,800 bp
19,451,820 bp
19,451,840 bp
19,451,860 bp
19,451,880 bp
LIBD75_Illum...bam Coverage
[0 - 37]
1
2
Unphased
LIBD75_Illumina_WGS.markedTrueG.phased.sorted.recal.localrealign.bam
Sequence
Gene
CAPZB

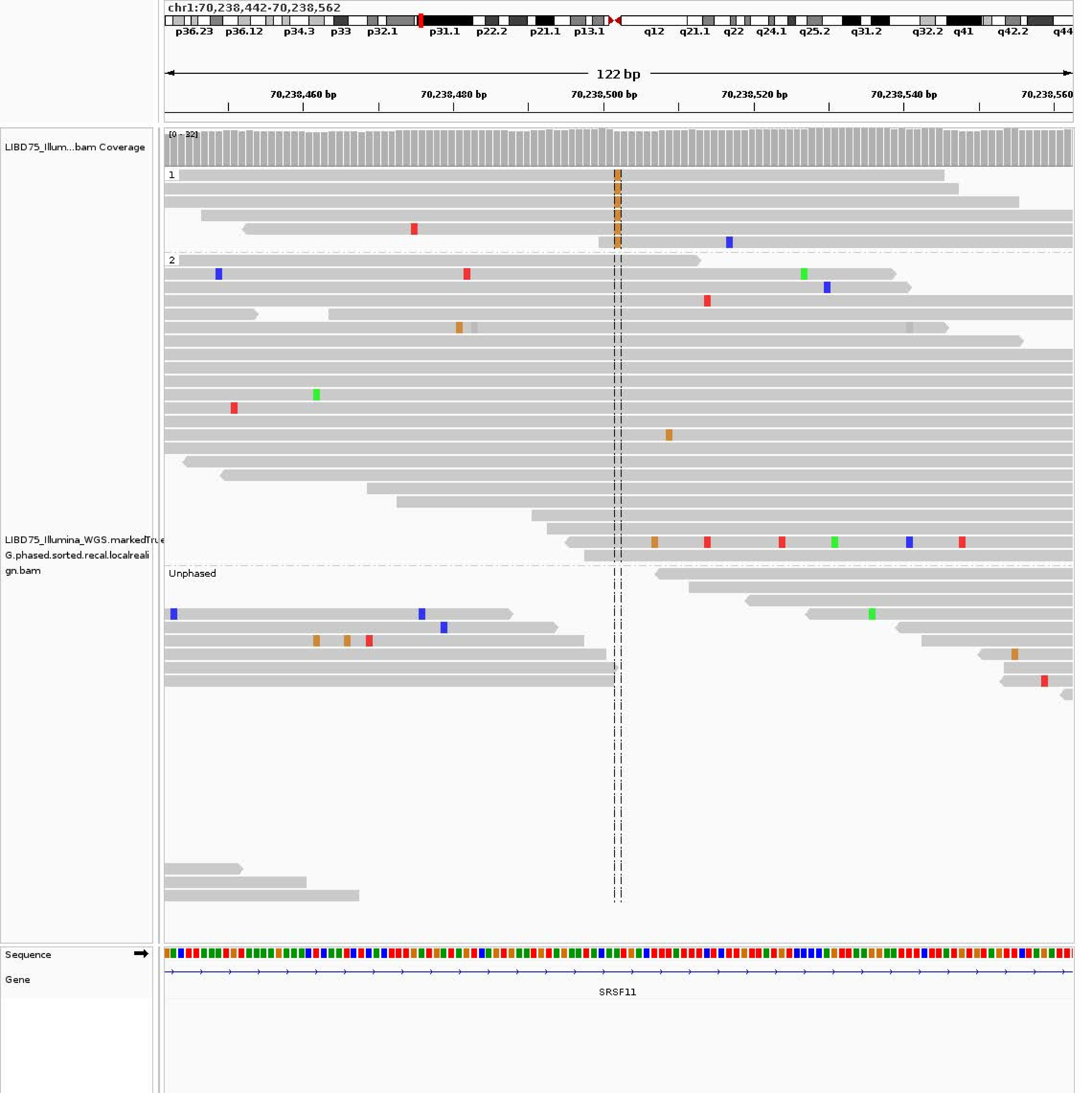

chr1:70,238,442-70,238,562
p36.23 p36.12 p34.3 p33 p32.1 p31.1 p22.2 p21.1 p13.1 q12 q21.1 q22 q24.1 q25.2 q31.2 q32.2 q41 q42.2 q44
122 bp
70,238,460 bp
70,238,480 bp
70,238,500 bp
70,238,520 bp
70,238,540 bp
70,238,560
LIBD75_Illum...bam Coverage
[0 - 32]
1
2
Unphased
LIBD75_Illumina_WGS.markedTrue
G.phased.sorted.recal.localreali
gn.bam
Sequence
Gene
SRSF11

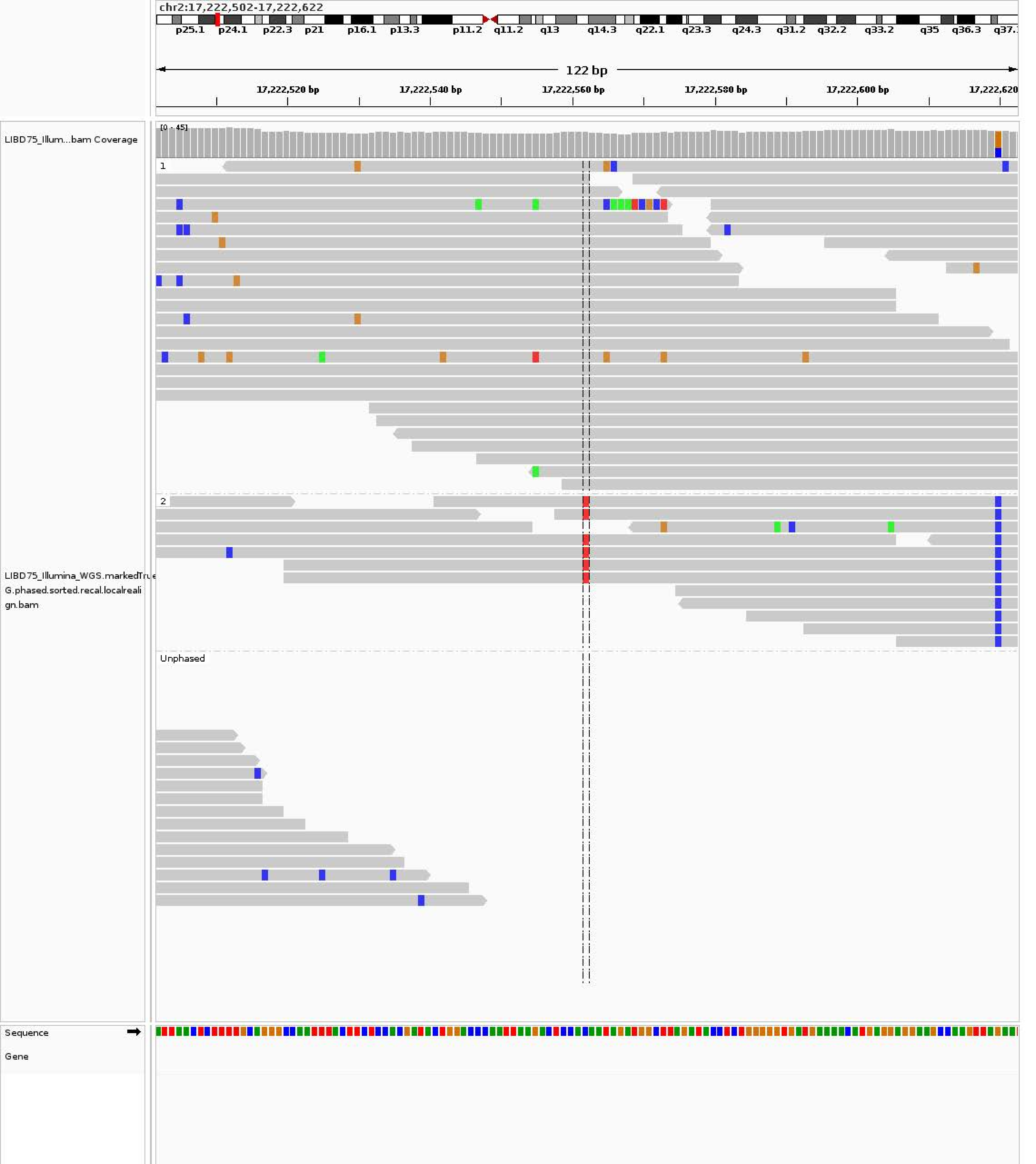

chr2:17,222,502-17,222,622
p25.1 p24.1 p22.3 p21 p16.1 p13.3 p11.2 q11.2 q13 q14.3 q22.1 q23.3 q24.3 q31.2 q32.2 q33.2 q35 q36.3
122 bp
17,222,520 bp
17,222,540 bp
17,222,560 bp
17,222,580 bp
17,222,600 bp
17,222,620
LIBD75_Illum...bam Coverage
[0 - 45]
1
2
Unphased
LIBD75_Illumina_WGS.markedTrue
G.phased.sorted.recal.localreali
gn.bam
Sequence
Gene

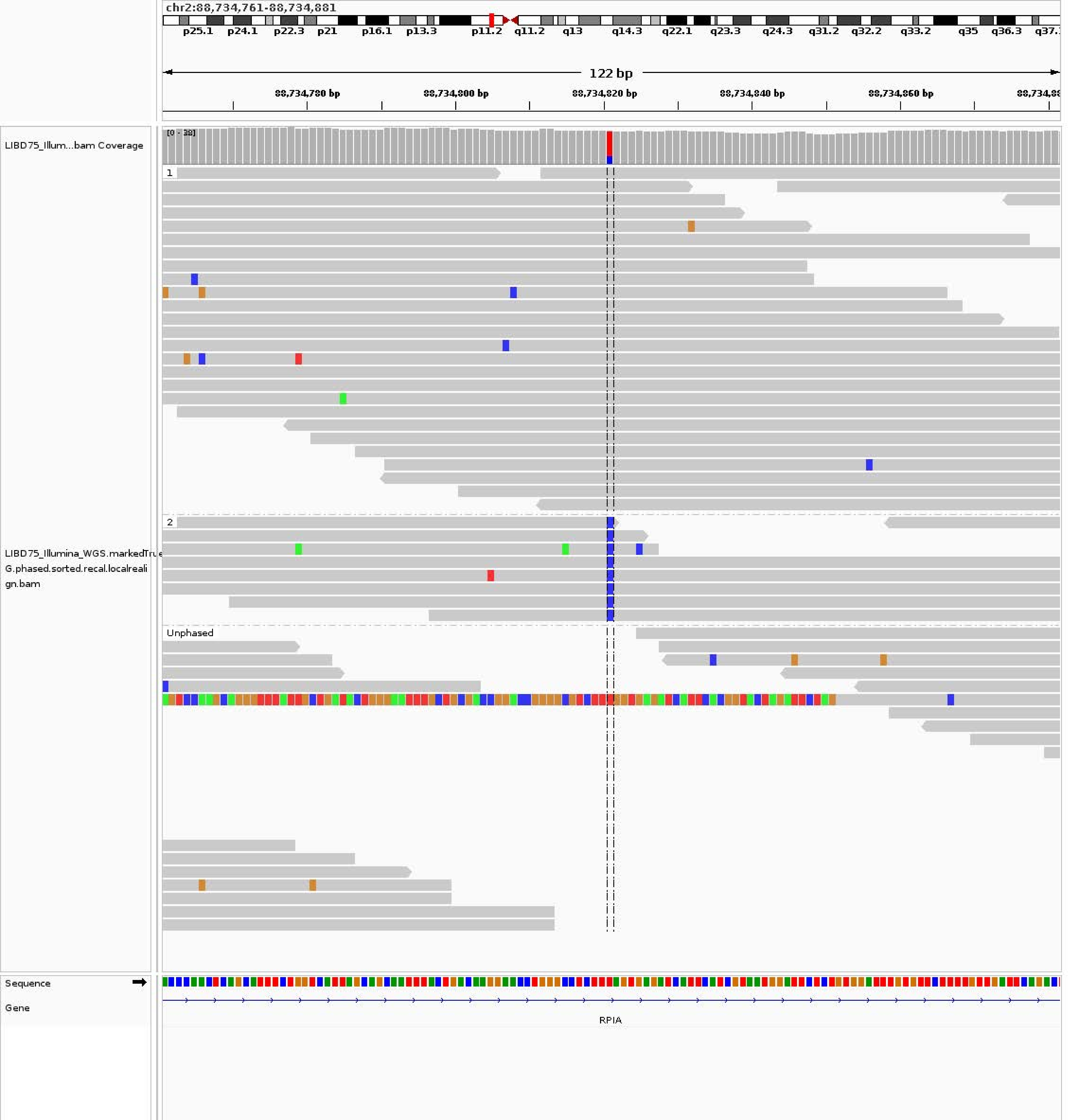
chr2:88,734,761-88,734,881
p25.1 p24.1 p22.3 p21 p16.1 p13.3 p11.2 q11.2 q13 q14.3 q22.1 q23.3 q24.3 q31.2 q32.2 q33.2 q35 q36.3
122 bp
88,734,780 bp
88,734,800 bp
88,734,820 bp
88,734,840 bp
88,734,860 bp
LIBD75_Illum...bam Coverage
[0 - 38]
1
2
Unphased
LIBD75_Illumina_WGS.markedTru
G.phased.sorted.recal.localreali
gn.bam
Sequence
Gene
RPIA

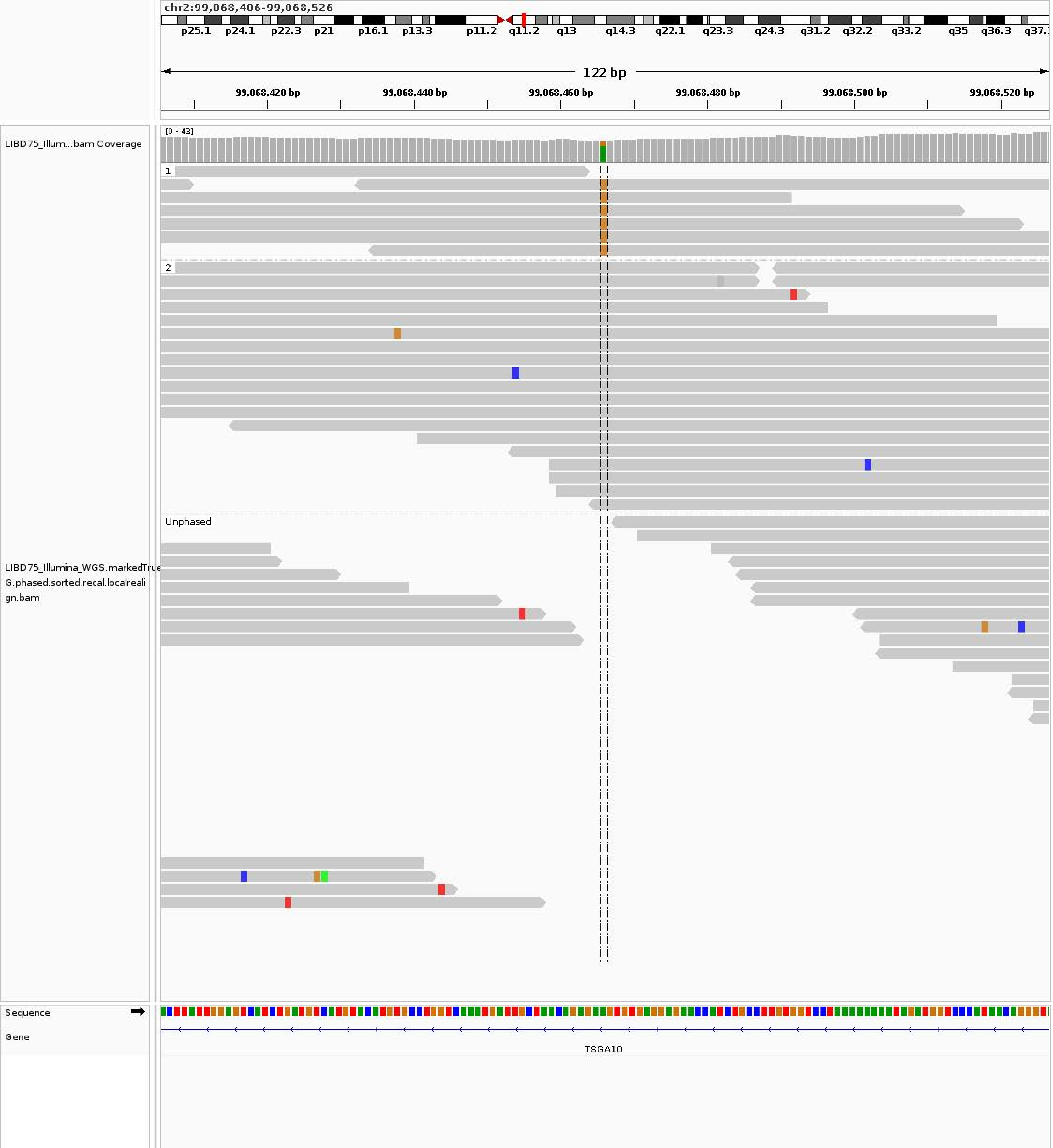

chr2:99,068,406-99,068,526
p25.1 p24.1 p22.3 p21 p16.1 p13.3 p11.2 q11.2 q13 q14.3 q22.1 q23.3 q24.3 q31.2 q32.2 q33.2 q35 q36.3
122 bp
99,068,420 bp
99,068,440 bp
99,068,460 bp
99,068,480 bp
99,068,500 bp
99,068,520 bp
LIBD75_Illum...bam Coverage
[0 - 43]
1
2
Unphased
LIBD75_Illumina_WGS.markedTrue
G.phased.sorted.recal.localreali
gn.bam
Sequence
Gene
TSGA10

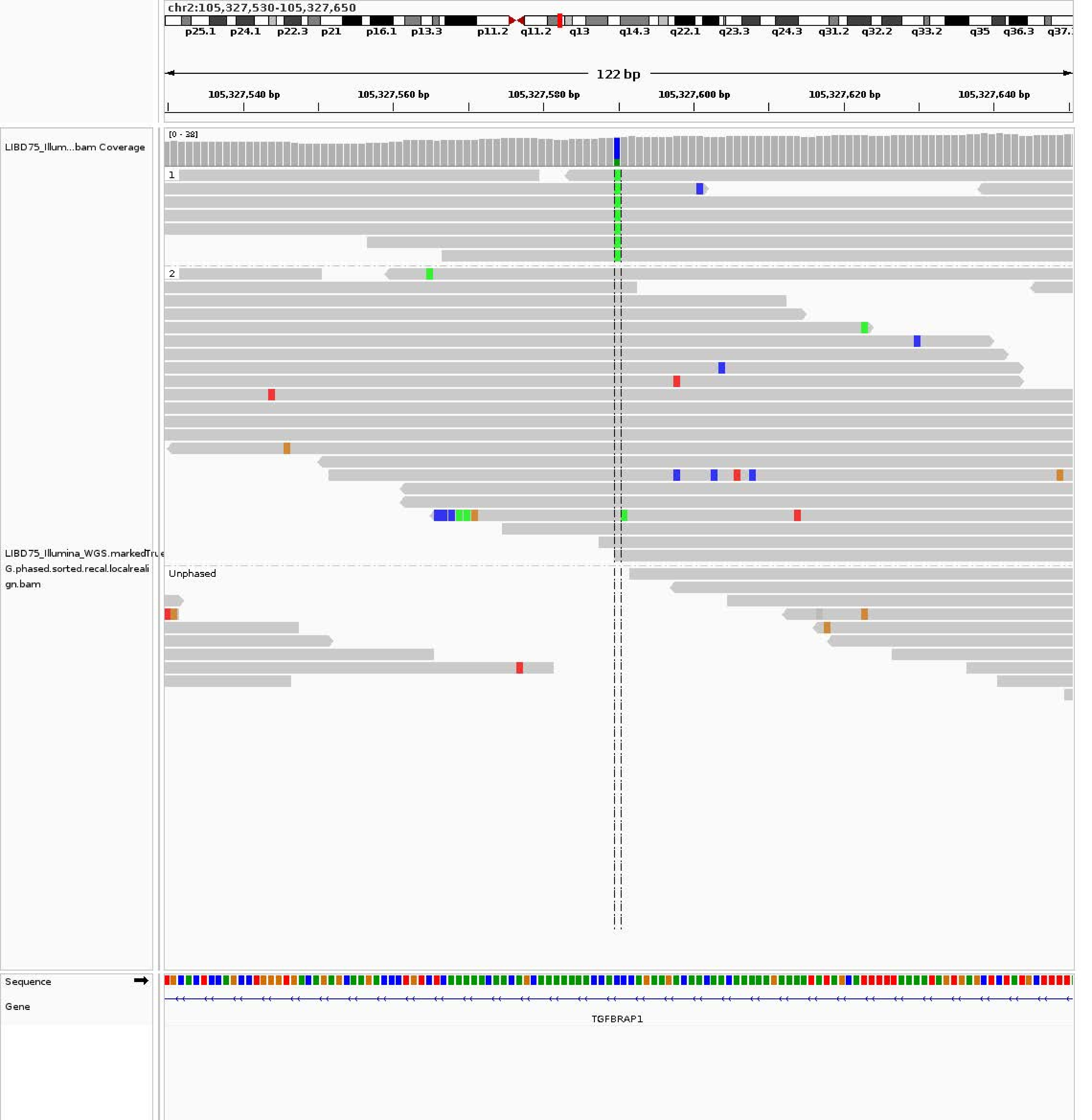
chr2:105,327,530-105,327,650
p25.1
p24.1
p22.3
p21
p16.1
p13.3
p11.2
q11.2
q13
q14.3
q22.1
q23.3
q24.3
q31.2
q32.2
q33.2
q35
q36.3
122 bp
105,327,540 bp
105,327,560 bp
105,327,580 bp
105,327,600 bp
105,327,620 bp
105,327,640 bp
LIBD75_Illum...bam Coverage
[0 - 38]
1
2
Unphased
LIBD75_Illumina_WGS.markedTrue
G.phased.sorted.recal.localreali
gn.bam
Sequence
Gene
TGFBRAP1

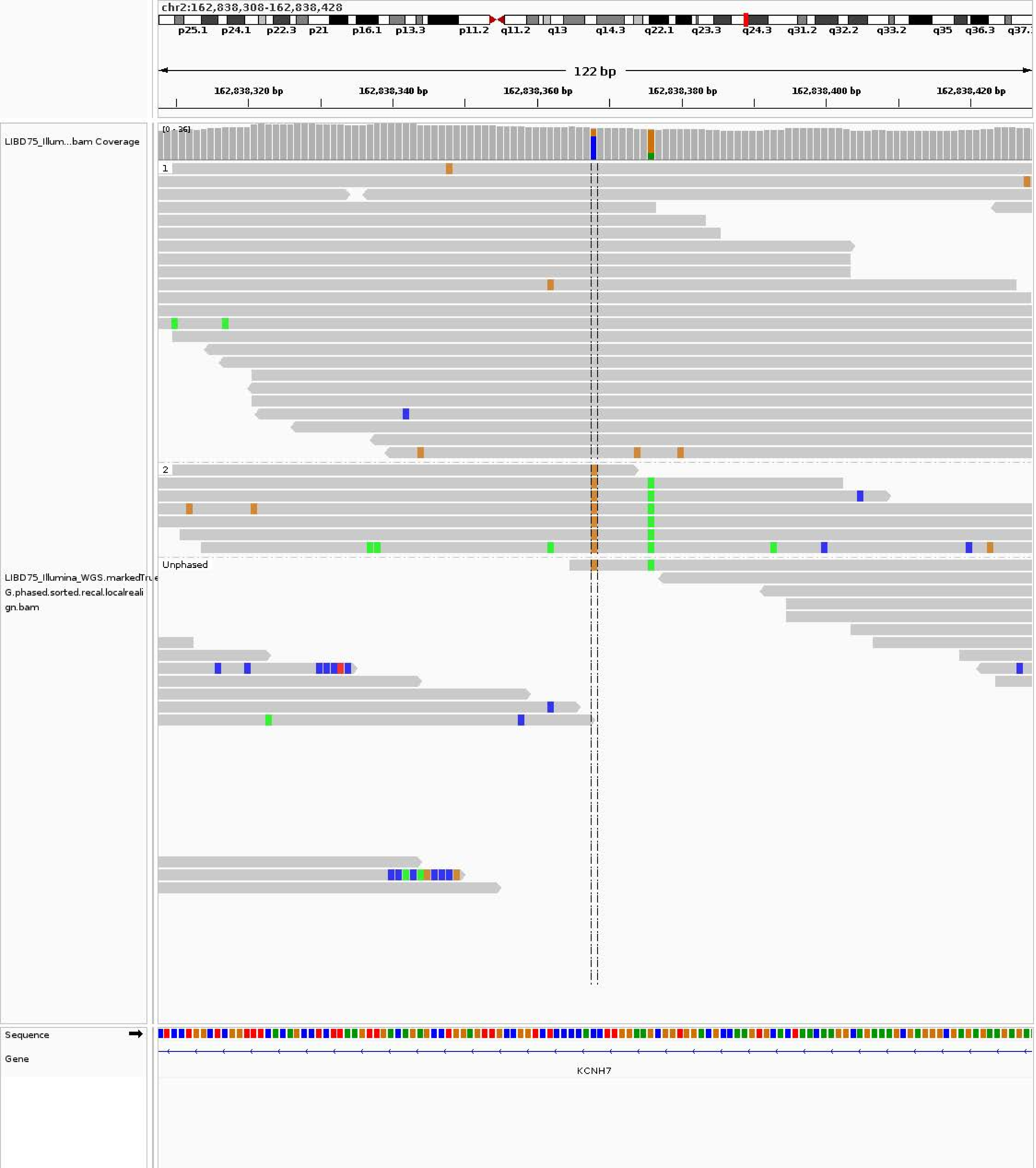
chr2:162,838,308-162,838,428
p25.1 p24.1 p22.3 p21 p16.1 p13.3 p11.2 q11.2 q13 q14.3 q22.1 q23.3 q24.3 q31.2 q32.2 q33.2 q35 q36.3
122 bp
162,838,320 bp
162,838,340 bp
162,838,360 bp
162,838,380 bp
162,838,400 bp
162,838,420 bp
LIBD75_Illum...bam Coverage
[0 - 36]
1
2
Unphased
LIBD75_Illumina_WGS.markedTrue
G.phased.sorted.recal.localreali
gn.bam
Sequence
Gene
KCNH7

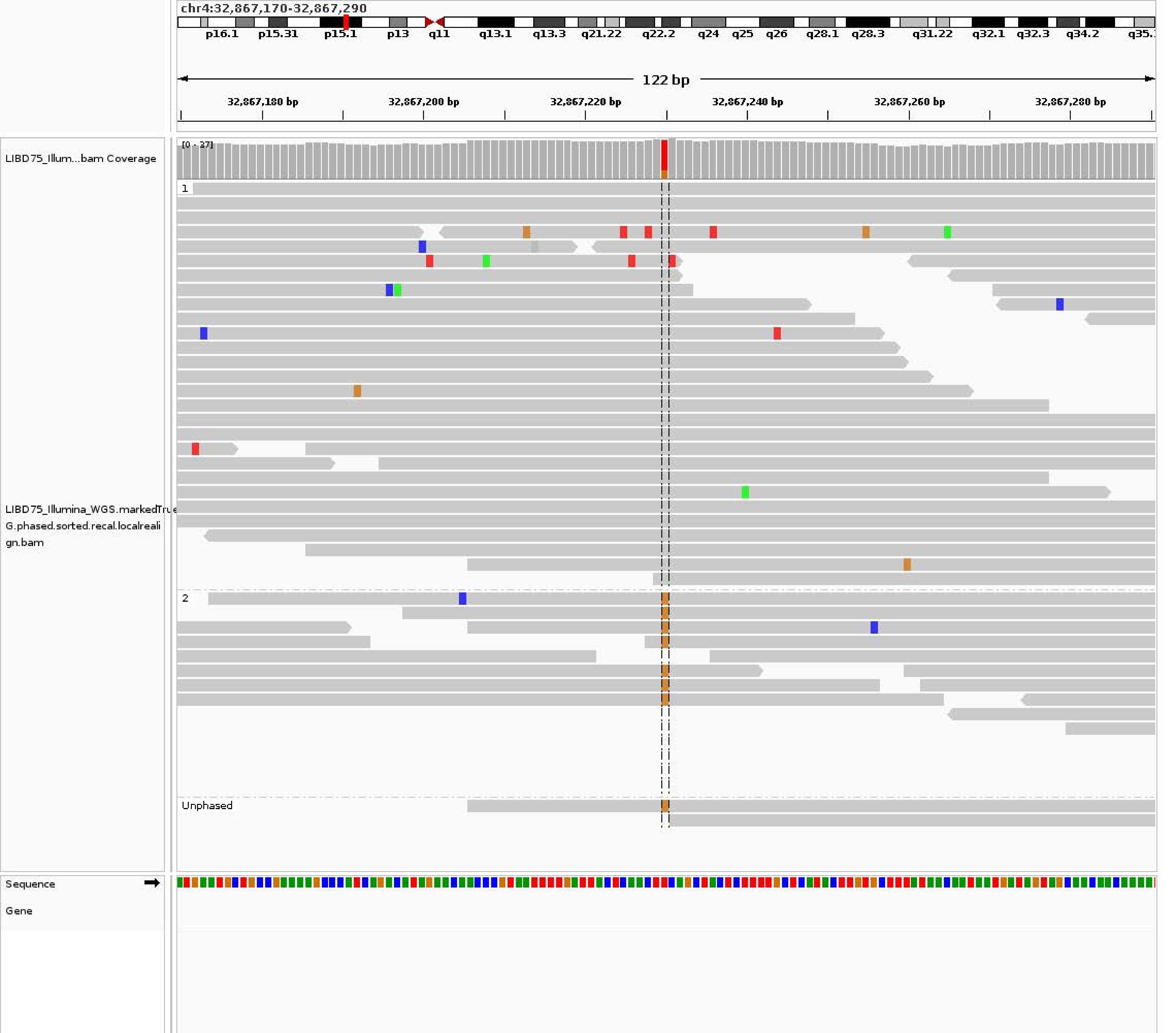
chr4:32,867,170-32,867,290
p16.1
p15.31
p15.1
p13
q11
q13.1
q13.3
q21.22
q22.2
q24
q25
q26
q28.1
q28.3
q31.22
q32.1
q32.3
q34.2
122 bp
32,867,180 bp
32,867,200 bp
32,867,220 bp
32,867,240 bp
32,867,260 bp
32,867,280 bp
LIBD75_Illum...bam Coverage
[0 - 37]
1
LIBD75_Illumina_WGS.markedTrue
G.phased.sorted.recal.localreali
gn.bam
2
Unphased
Sequence
Gene

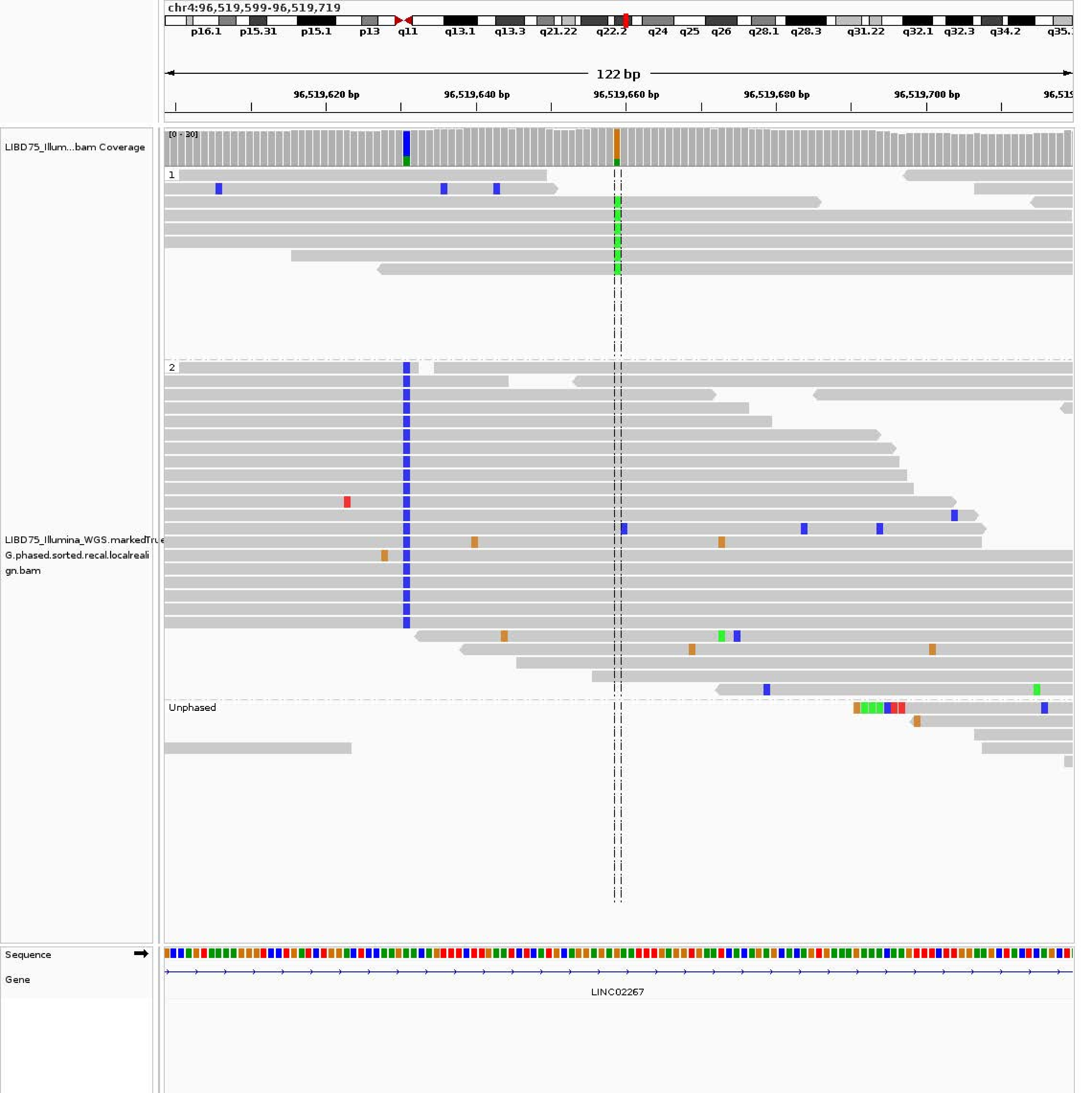

chr4:96,519,599-96,519,719
p16.1
p15.31
p15.1
p13
q11
q13.1
q13.3
q21.22
q22.2
q24
q25
q26
q28.1
q28.3
q31.22
q32.1
q32.3
q34.2
122 bp
96,519,620 bp
96,519,640 bp
96,519,660 bp
96,519,680 bp
96,519,700 bp
LIBD75_Illum...bam Coverage
[0 - 30]
1
2
Unphased
LIBD75_Illumina_WGS.markedTrue
G.phased.sorted.recal.localreali
gn.bam
Sequence
Gene
LINC02267

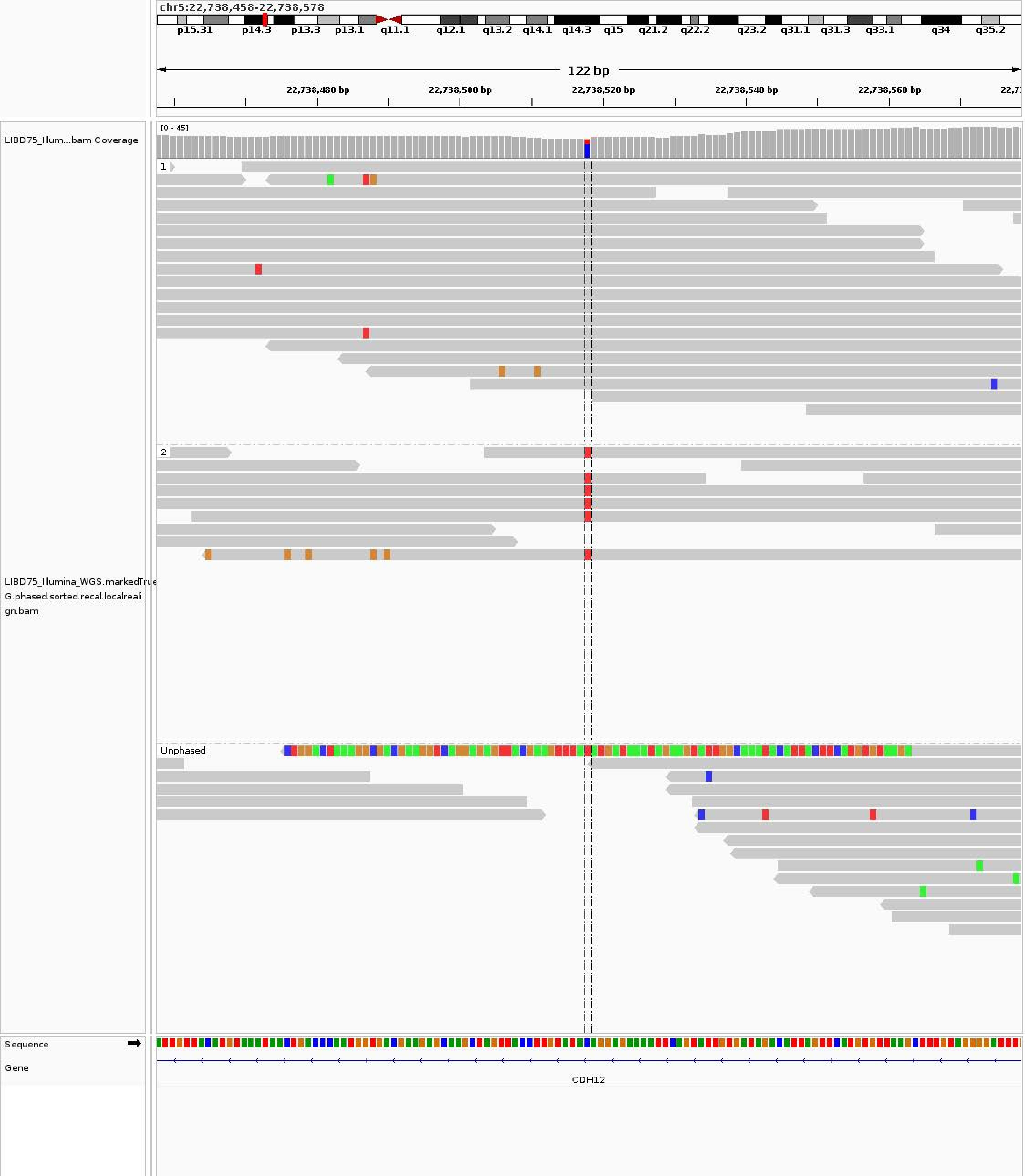

chr5:22,738,458-22,738,578
p15.31 p14.3 p13.3 p13.1 q11.1 q12.1 q13.2 q14.1 q14.3 q15 q21.2 q22.2 q23.2 q31.1 q31.3 q33.1 q34 q35.2
122 bp
22,738,480 bp
22,738,500 bp
22,738,520 bp
22,738,540 bp
22,738,560 bp
LIBD75_Illum...bam Coverage
[0 - 45]
1
2
Unphased
LIBD75_Illumina_WGS.markedTrue
G.phased.sorted.recal.localreali
gn.bam
Sequence
Gene
CDH12

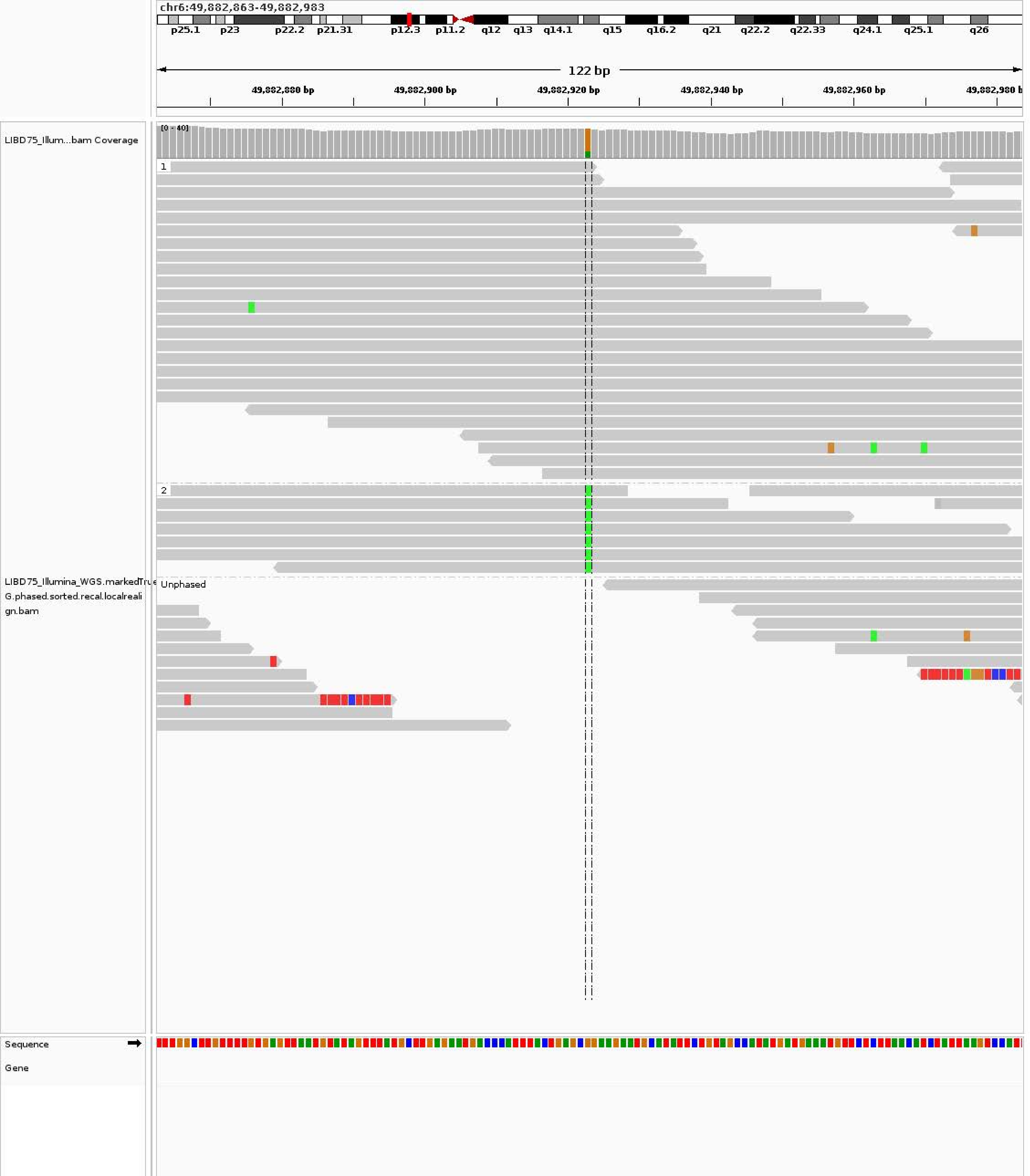
chr6:49,882,863-49,882,983
p25.1
p23
p22.2
p21.31
p12.3
p11.2
q12
q13
q14.1
q15
q16.2
q21
q22.2
q22.33
q24.1
q25.1
q26
122 bp
49,882,880 bp
49,882,900 bp
49,882,920 bp
49,882,940 bp
49,882,960 bp
LIBD75_Illum...bam Coverage
[0 - 40]
1
2
Unphased
LIBD75_Illumina_WGS.markedTrue G.phased.sorted.recal.localrealign.bam
Sequence
Gene

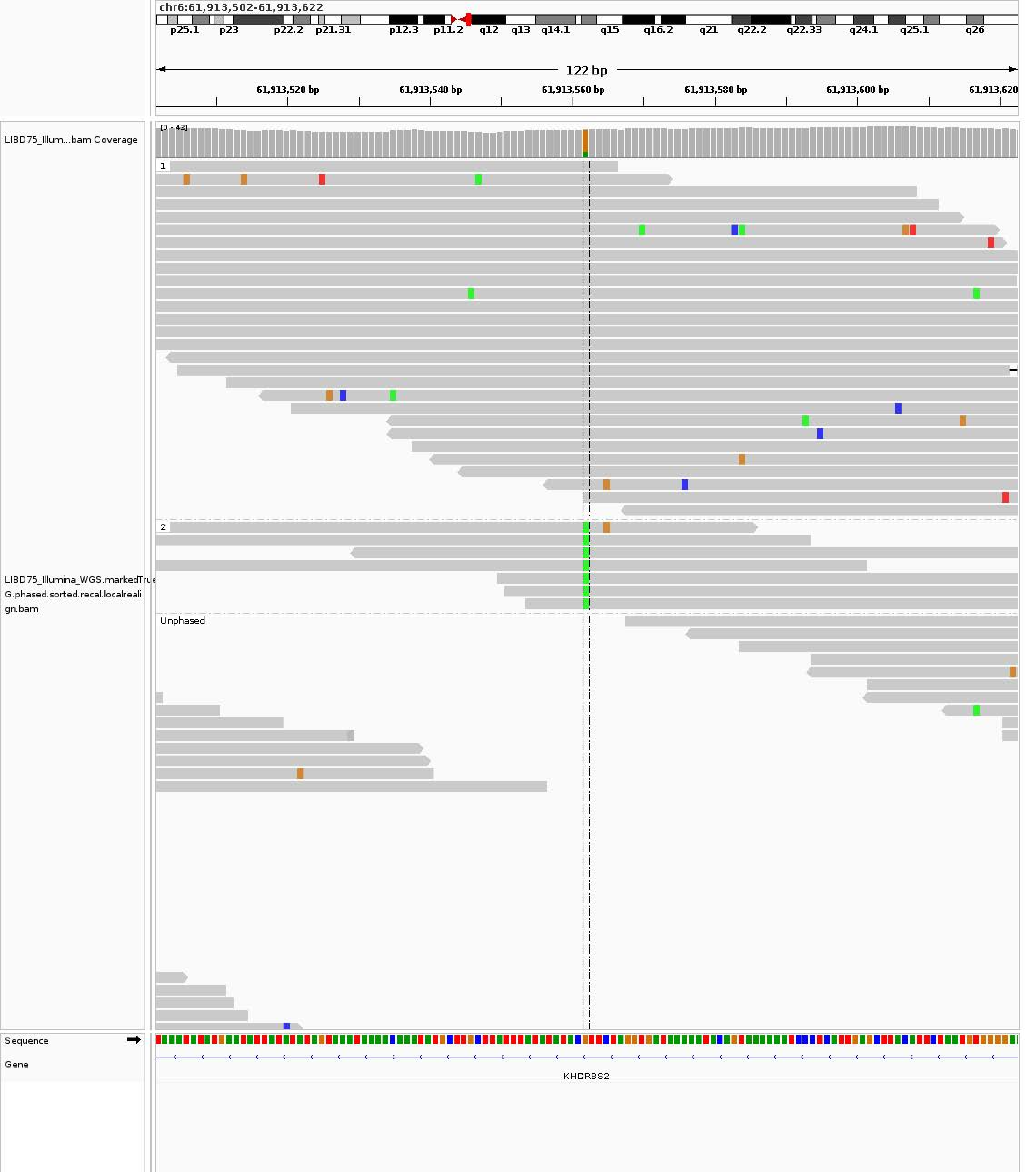
chr6:61,913,502-61,913,622
p25.1 p23 p22.2 p21.31 p12.3 p11.2 q12 q13 q14.1 q15 q16.2 q21 q22.2 q22.33 q24.1 q25.1 q26
122 bp
61,913,520 bp
61,913,540 bp
61,913,560 bp
61,913,580 bp
61,913,600 bp
61,913,620
LIBD75_Illum...bam Coverage
[0 - 43]
1
2
Unphased
LIBD75_Illumina_WGS.markedTrue G.phased.sorted.recal.localrealign.bam
Sequence
Gene
KHDRBS2

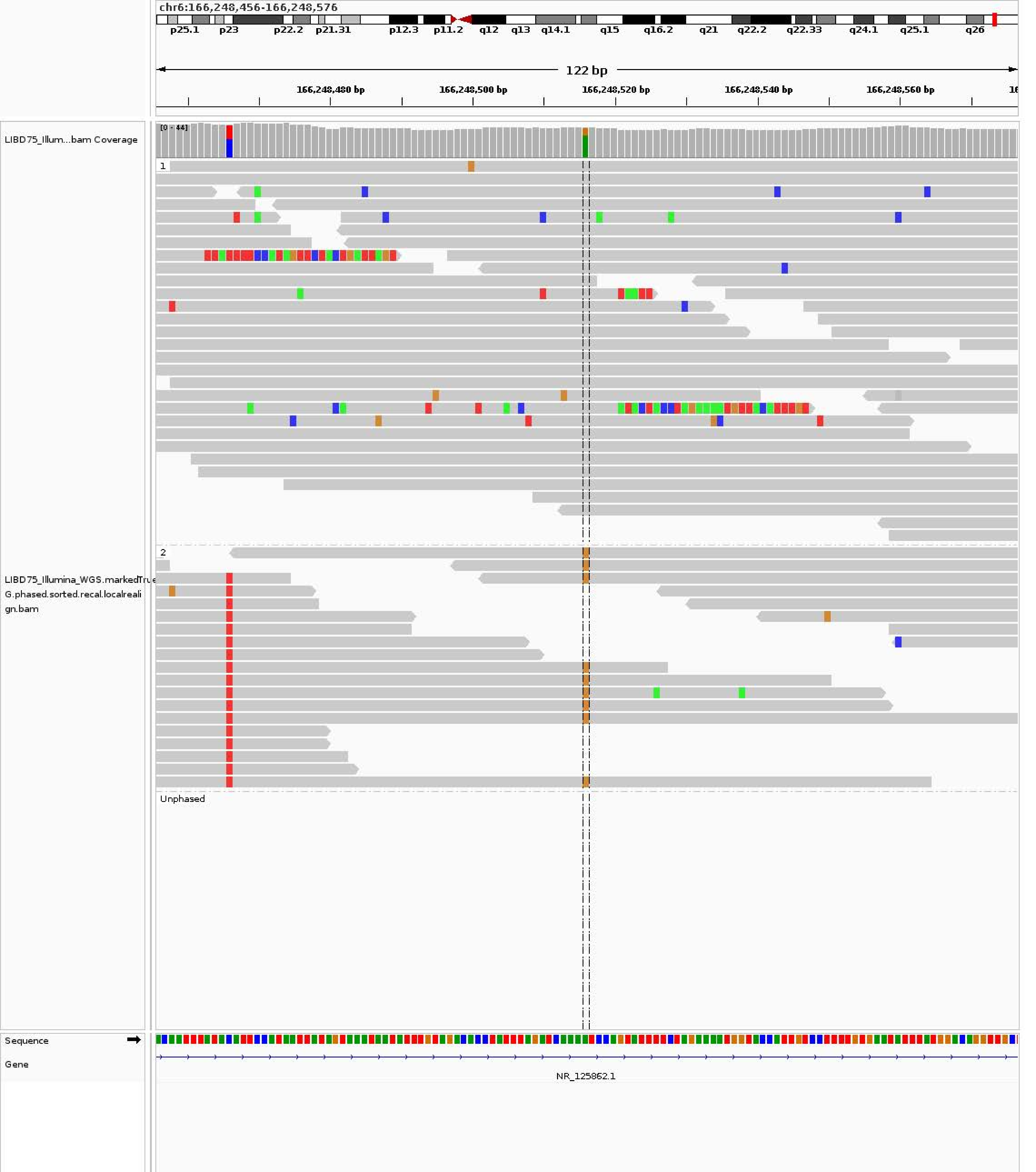
chr6:166,248,456-166,248,576
p25.1 p23 p22.2 p21.31 p12.3 p11.2 q12 q13 q14.1 q15 q16.2 q21 q22.2 q22.33 q24.1 q25.1 q26
122 bp
166,248,480 bp
166,248,500 bp
166,248,520 bp
166,248,540 bp
166,248,560 bp
LIBD75_Illum...bam Coverage
[0 - 44]
1
2
Unphased
LIBD75_Illumina_WGS.markedTrue
G.phased.sorted.recal.localreali
gn.bam
Sequence
Gene
NR_125862.1

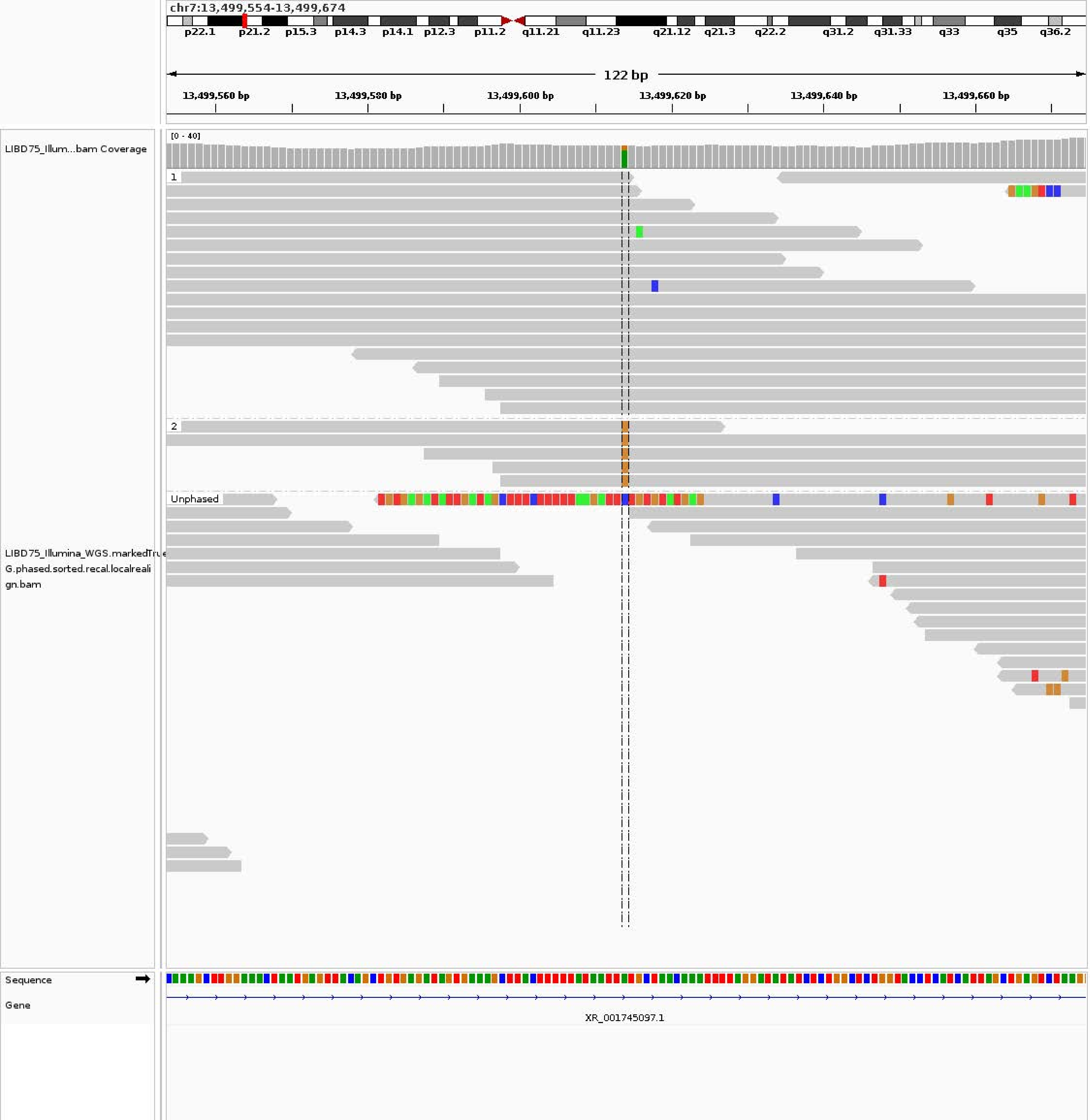
chr7:13,499,554-13,499,674
p22.1 p21.2 p15.3 p14.3 p14.1 p12.3 p11.2 q11.21 q11.23 q21.12 q21.3 q22.2 q31.2 q31.33 q33 q35 q36.2
122 bp
13,499,560 bp
13,499,580 bp
13,499,600 bp
13,499,620 bp
13,499,640 bp
13,499,660 bp
LIBD75_Illum...bam Coverage
[0 - 40]
1
2
Unphased
LIBD75_Illumina_WGS.markedTrue
G.phased.sorted.recal.localreali
gn.bam
Sequence
Gene
XR_001745097.1

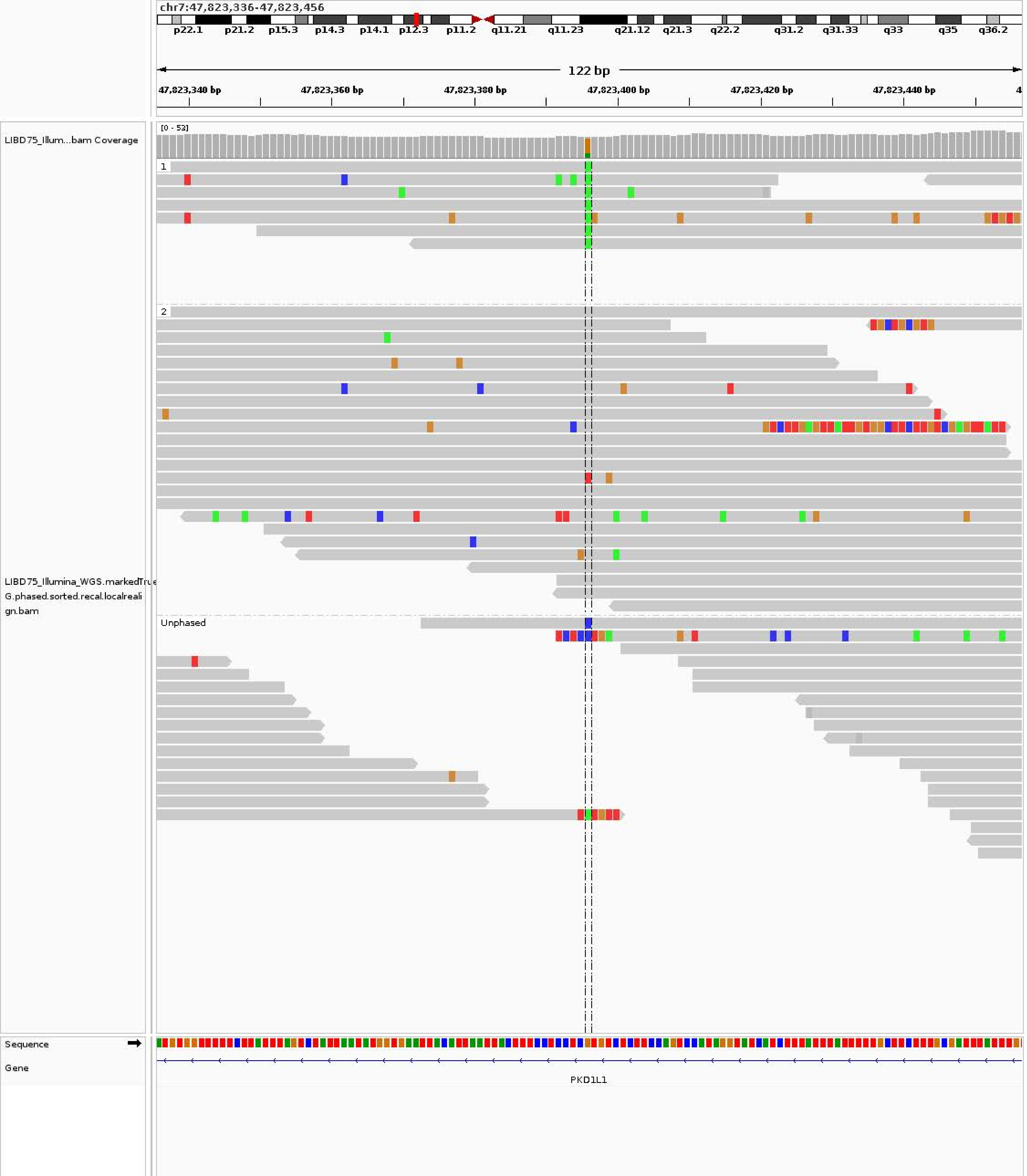
chr7:47,823,336-47,823,456
p22.1 p21.2 p15.3 p14.3 p14.1 p12.3 p11.2 q11.21 q11.23 q21.12 q21.3 q22.2 q31.2 q31.33 q33 q35 q36.2
122 bp
47,823,340 bp 47,823,360 bp 47,823,380 bp 47,823,400 bp 47,823,420 bp 47,823,440 bp
LIBD75_Illum...bam Coverage
[0 - 53]
1
2
Unphased
LIBD75_Illumina_WGS.markedTrueG.phased.sorted.recal.localrealign.bam
Sequence
Gene
PKD1L1

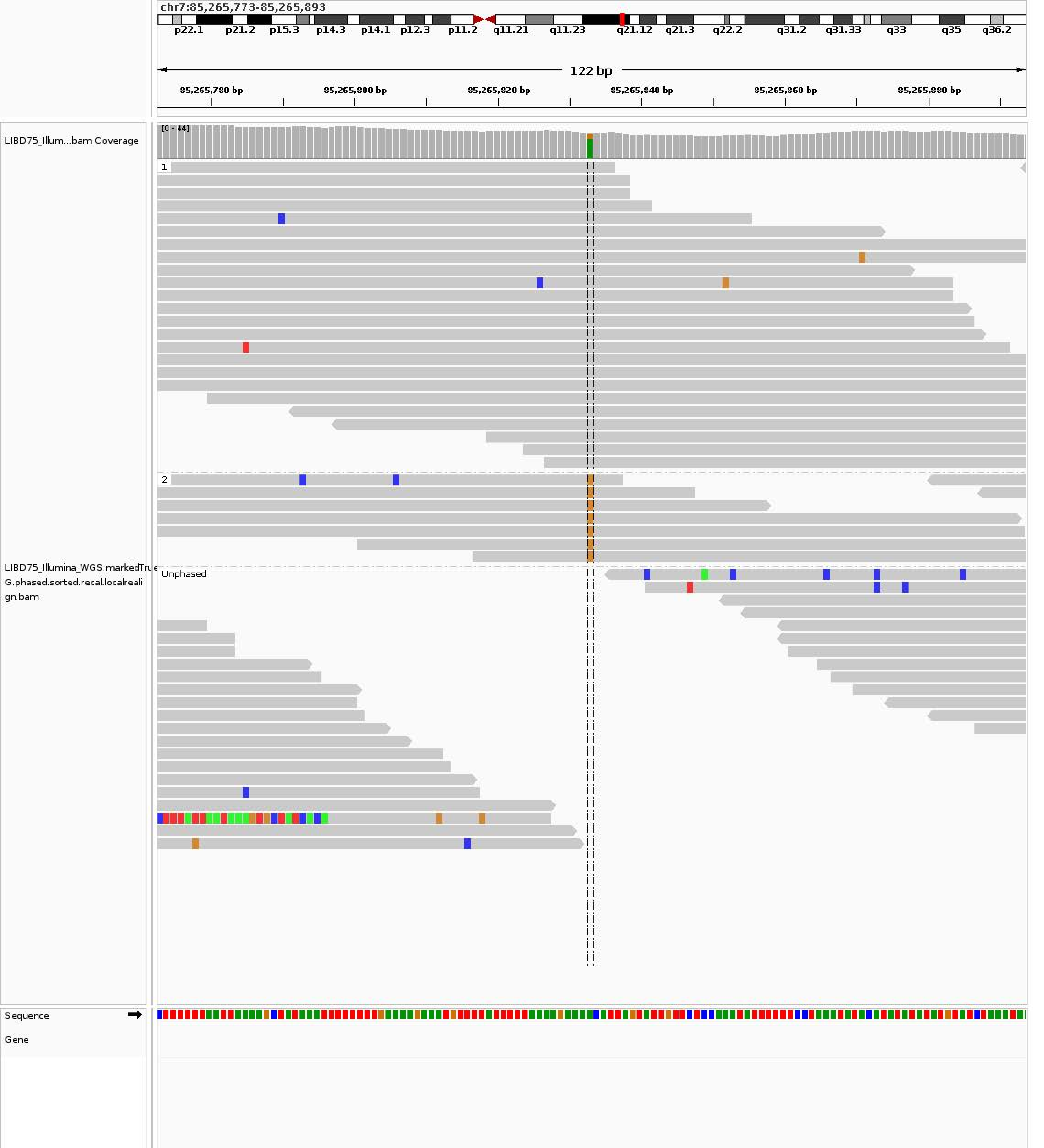
chr7:85,265,773-85,265,893
p22.1
p21.2
p15.3
p14.3
p14.1
p12.3
p11.2
q11.21
q11.23
q21.12
q21.3
q22.2
q31.2
q31.33
q33
q35
q36.2
122 bp
85,265,780 bp
85,265,800 bp
85,265,820 bp
85,265,840 bp
85,265,860 bp
85,265,880 bp
LIBD75_Illum...bam Coverage
[0 - 44]
1
2
Unphased
LIBD75_Illumina_WGS.markedTrue
G.phased.sorted.recal.localreali
gn.bam
Sequence
Gene

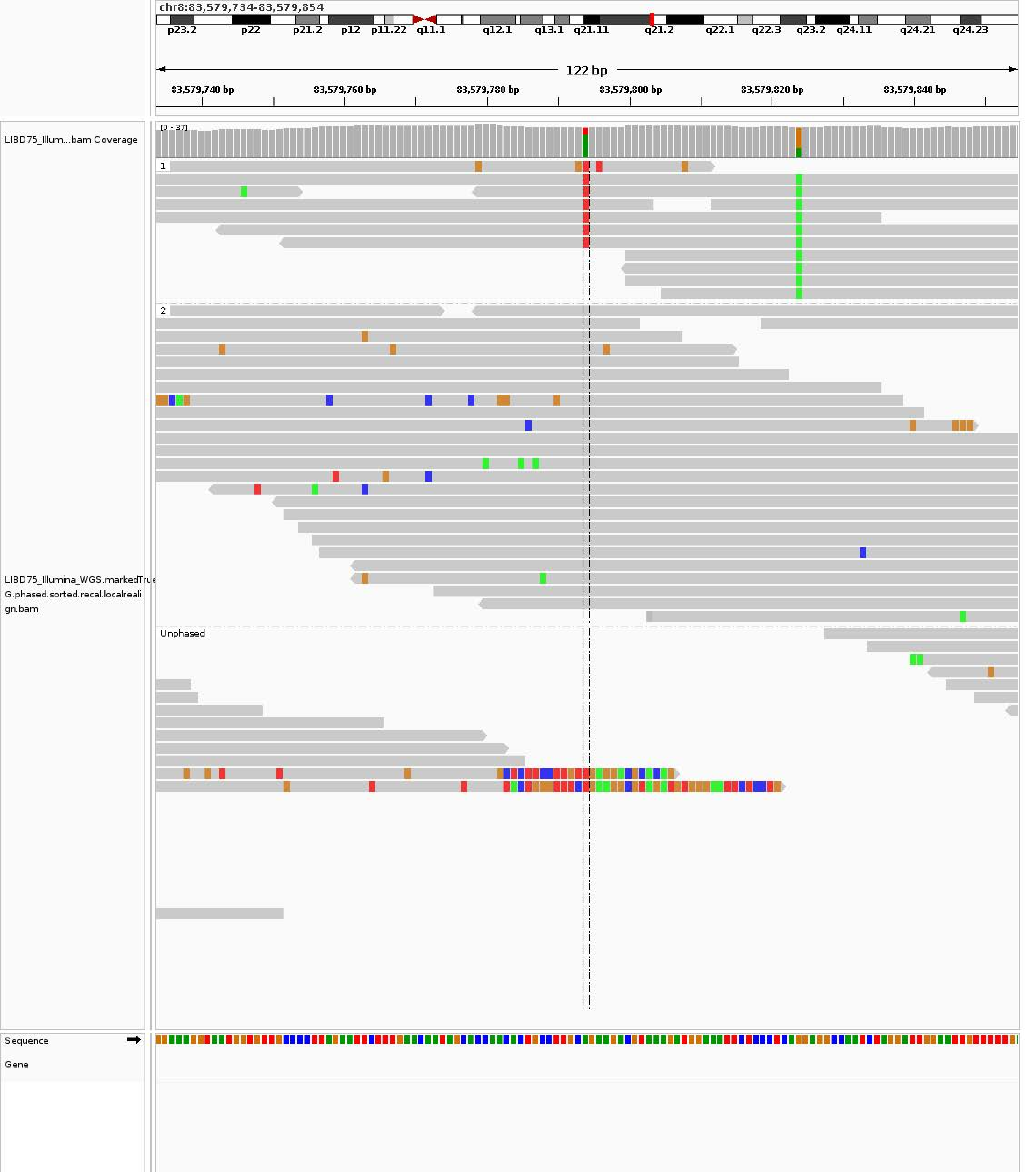
chr8:83,579,734-83,579,854
p23.2 p22 p21.2 p12 p11.22 q11.1 q12.1 q13.1 q21.11 q21.2 q22.1 q22.3 q23.2 q24.11 q24.21 q24.23
122 bp
83,579,740 bp
83,579,760 bp
83,579,780 bp
83,579,800 bp
83,579,820 bp
83,579,840 bp
LIBD75_Illum...bam Coverage
[0 - 37]
1
2
Unphased
LIBD75_Illumina_WGS.markedTrueG.phased.sorted.recal.localrealign.bam
Sequence
Gene

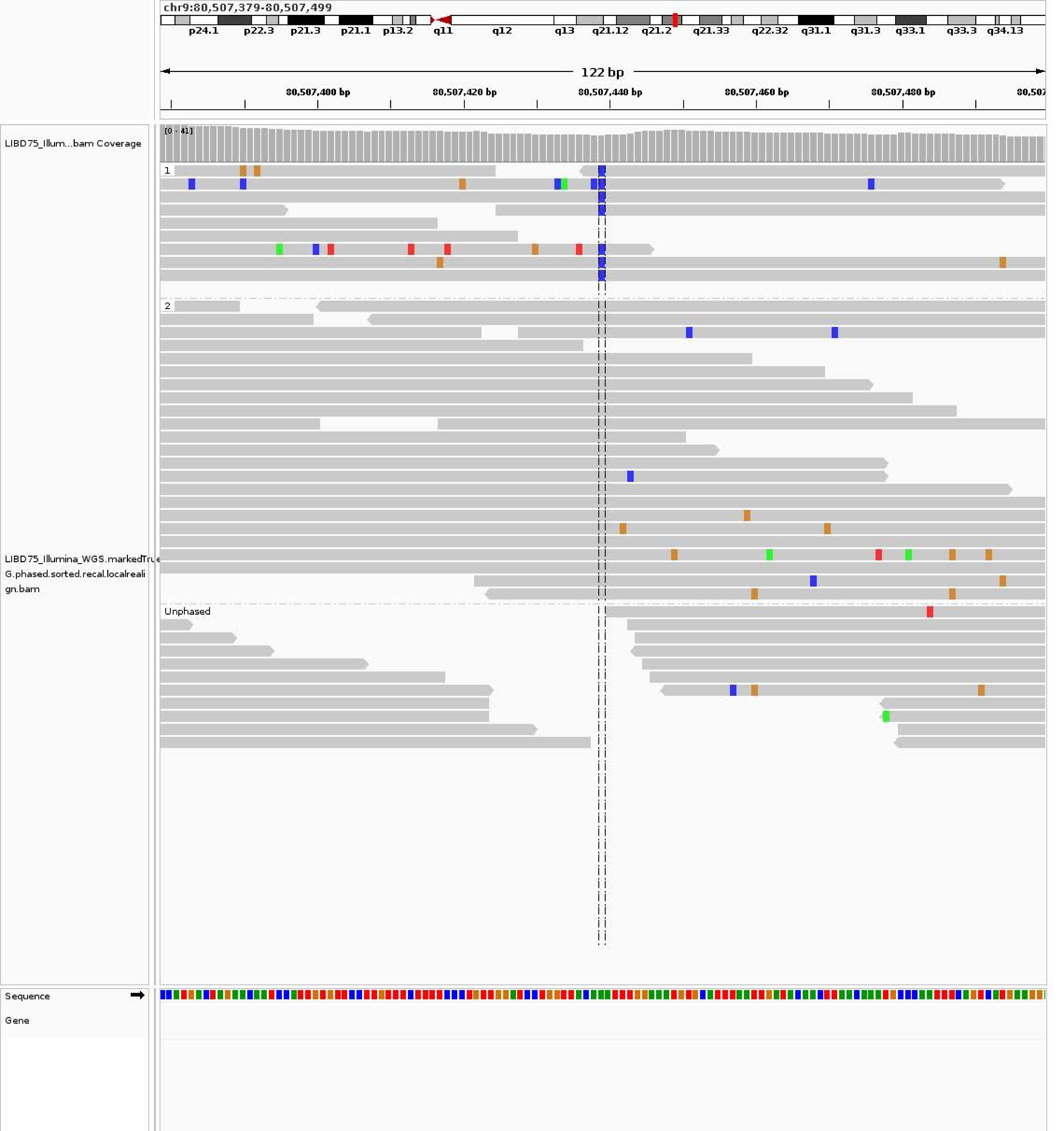

chr9:80,507,379-80,507,499
p24.1
p22.3
p21.3
p21.1
p13.2
q11
q12
q13
q21.12
q21.2
q21.33
q22.32
q31.1
q31.3
q33.1
q33.3
q34.13
122 bp
80,507,400 bp
80,507,420 bp
80,507,440 bp
80,507,460 bp
80,507,480 bp
LIBD75_Illum...bam Coverage
[0 - 41]
1
2
Unphased
LIBD75_Illumina_WGS.markedTrue
G.phased.sorted.recal.localreali
gn.bam
Sequence
Gene

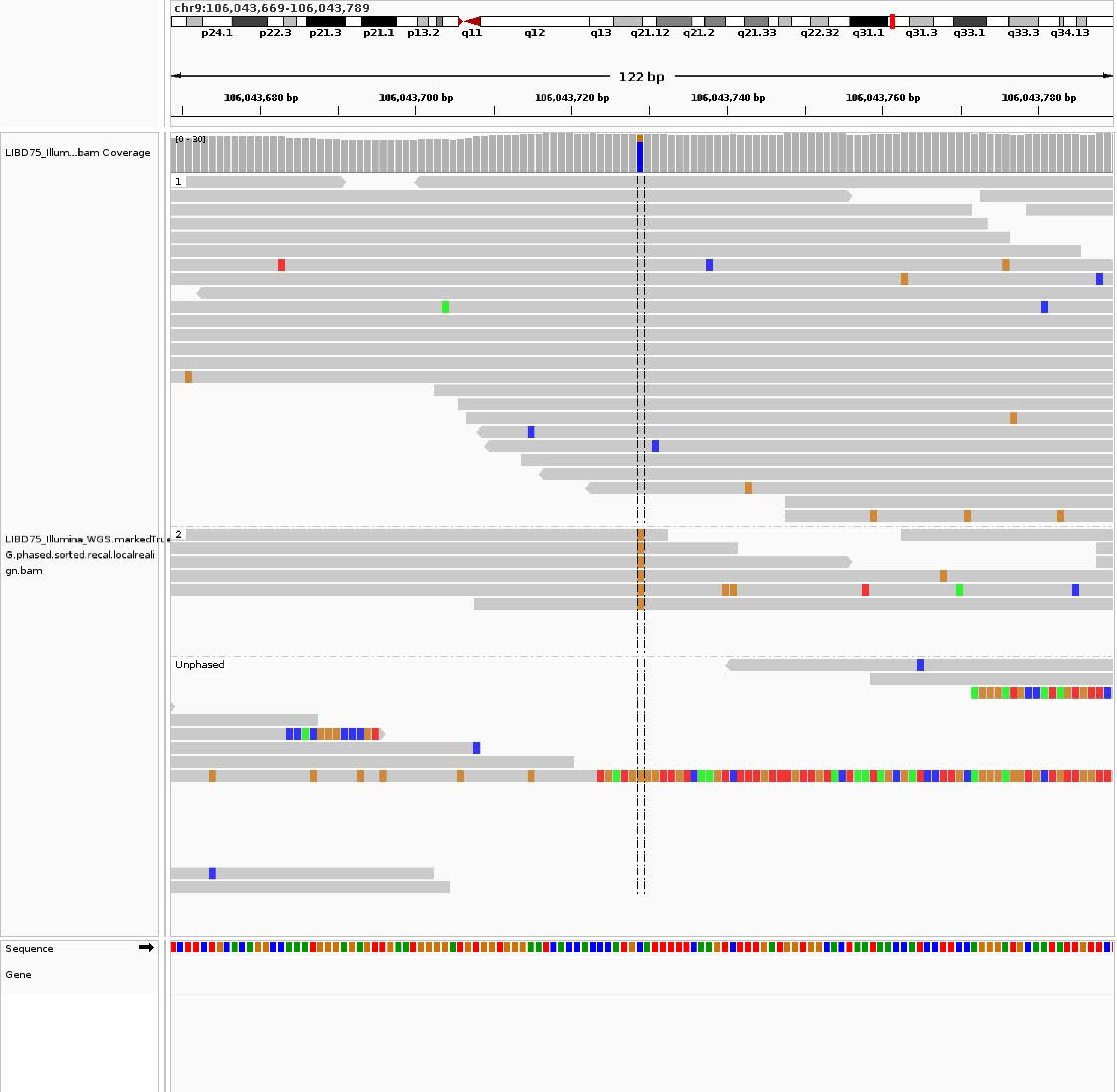
chr9:106,043,669-106,043,789
p24.1
p22.3
p21.3
p21.1
p13.2
q11
q12
q13
q21.12
q21.2
q21.33
q22.32
q31.1
q31.3
q33.1
q33.3
q34.13
122 bp
106,043,680 bp
106,043,700 bp
106,043,720 bp
106,043,740 bp
106,043,760 bp
106,043,780 bp
LIBD75_Illum...bam Coverage
[0 - 30]
1
LIBD75_Illumina_WGS.markedTrue
G.phased.sorted.recal.localreali
gn.bam
2
Unphased
Sequence
Gene

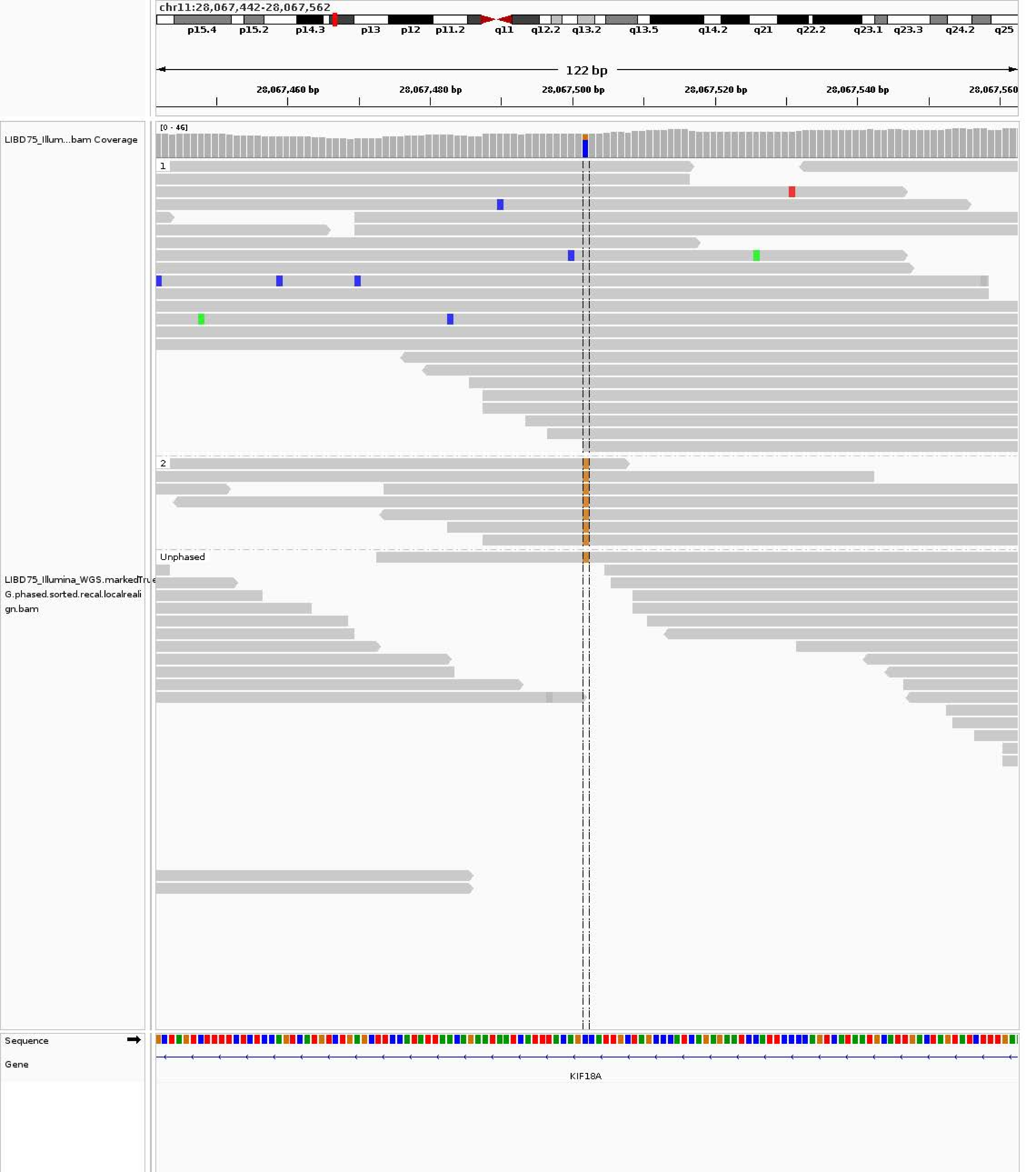

chr11:28,067,442-28,067,562
p15.4
p15.2
p14.3
p13
p12
p11.2
q11
q12.2
q13.2
q13.5
q14.2
q21
q22.2
q23.1
q23.3
q24.2
q25
122 bp
28,067,460 bp
28,067,480 bp
28,067,500 bp
28,067,520 bp
28,067,540 bp
28,067,560
LIBD75_Illum...bam Coverage
[0 - 46]
1
2
Unphased
LIBD75_Illumina_WGS.markedTrueG.phased.sorted.recal.localrealign.bam
Sequence
Gene
KIF18A

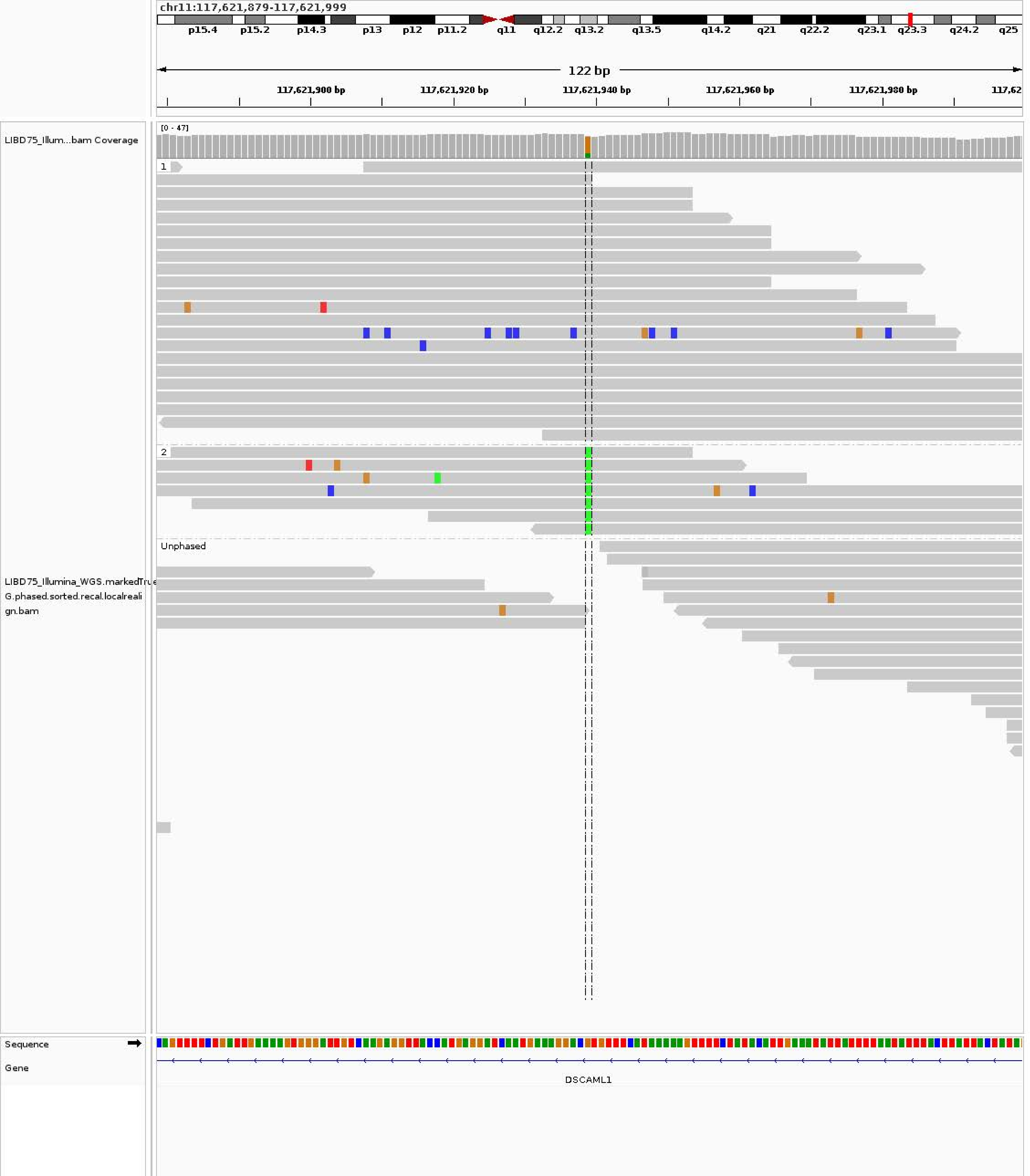
chr11:117,621,879-117,621,999
p15.4
p15.2
p14.3
p13
p12
p11.2
q11
q12.2
q13.2
q13.5
q14.2
q21
q22.2
q23.1
q23.3
q24.2
q25
122 bp
117,621,900 bp
117,621,920 bp
117,621,940 bp
117,621,960 bp
117,621,980 bp
LIBD75_Illum...bam Coverage
[0 - 47]
1
2
Unphased
LIBD75_Illumina_WGS.markedTrue
G.phased.sorted.recal.localreali
gn.bam
Sequence
Gene
DSCAML1

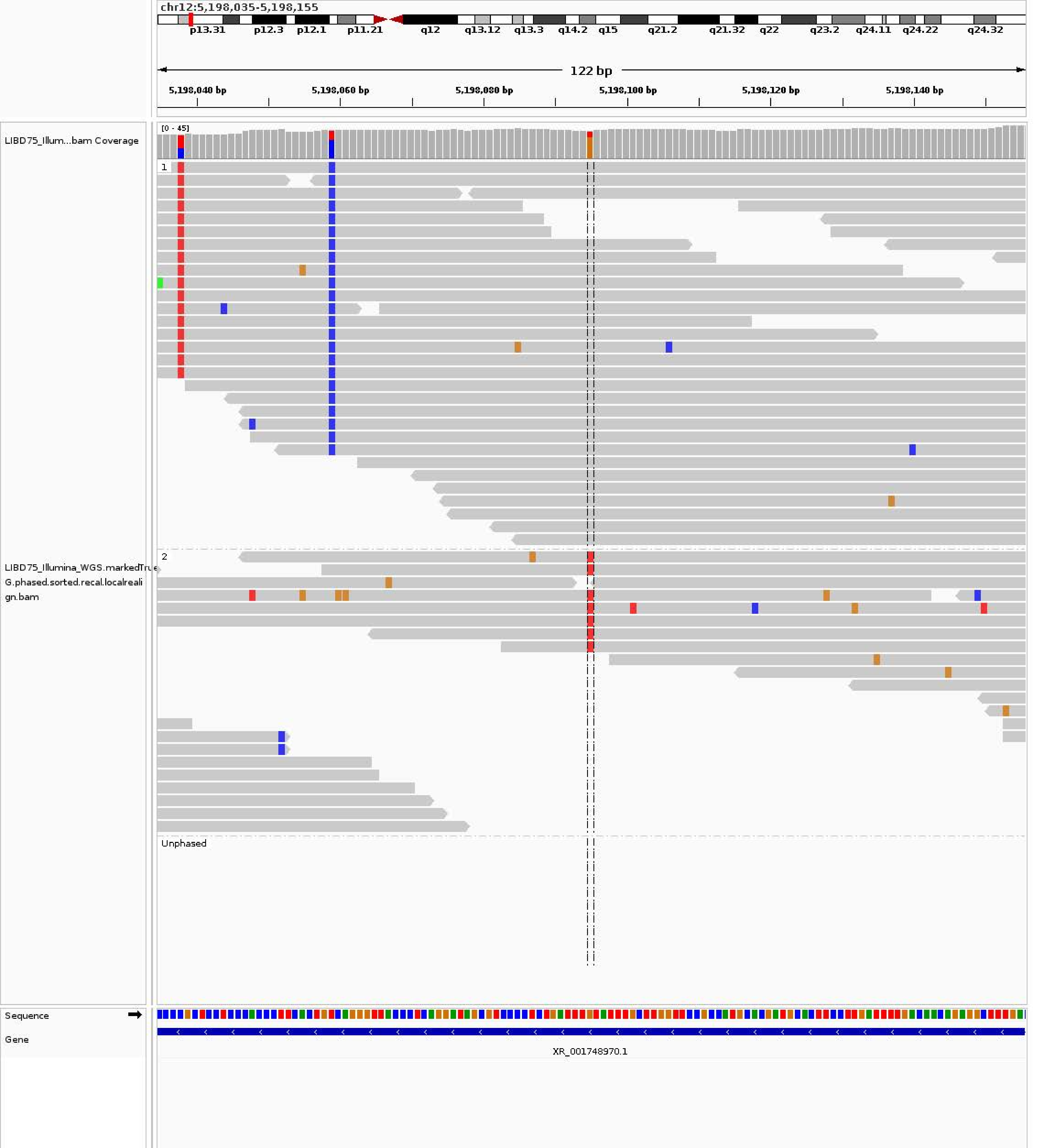
chr12:5,198,035-5,198,155
p13.31 p12.3 p12.1 p11.21 q12 q13.12 q13.3 q14.2 q15 q21.2 q21.32 q22 q23.2 q24.11 q24.22 q24.32
122 bp
5,198,040 bp 5,198,060 bp 5,198,080 bp 5,198,100 bp 5,198,120 bp 5,198,140 bp
LIBD75_Illum...bam Coverage
[0 - 45]
1
2
LIBD75_Illumina_WGS.markedTrue
G.phased.sorted.recal.localreali
gn.bam
Unphased
Sequence
Gene
XR_001748970.1

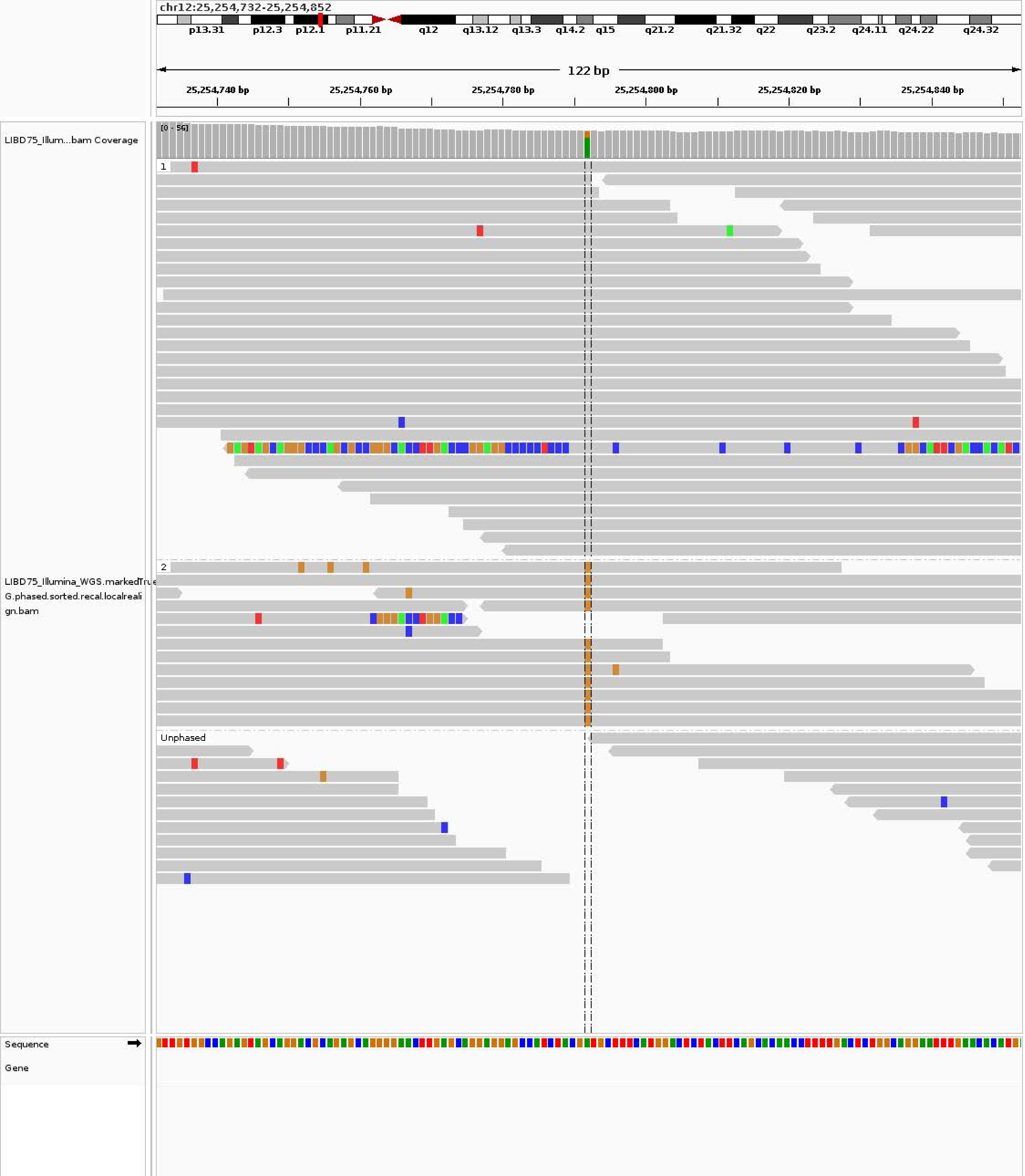
chr12:25,254,732-25,254,852
p13.31
p12.3
p12.1
p11.21
q12
q13.12
q13.3
q14.2
q15
q21.2
q21.32
q22
q23.2
q24.11
q24.22
q24.32
122 bp
25,254,740 bp
25,254,760 bp
25,254,780 bp
25,254,800 bp
25,254,820 bp
25,254,840 bp
LIBD75_Illum...bam Coverage
[0 - 56]
1
2
Unphased
LIBD75_Illumina_WGS.markedTrue G.phased.sorted.recal.localreali gn.bam
Sequence
Gene

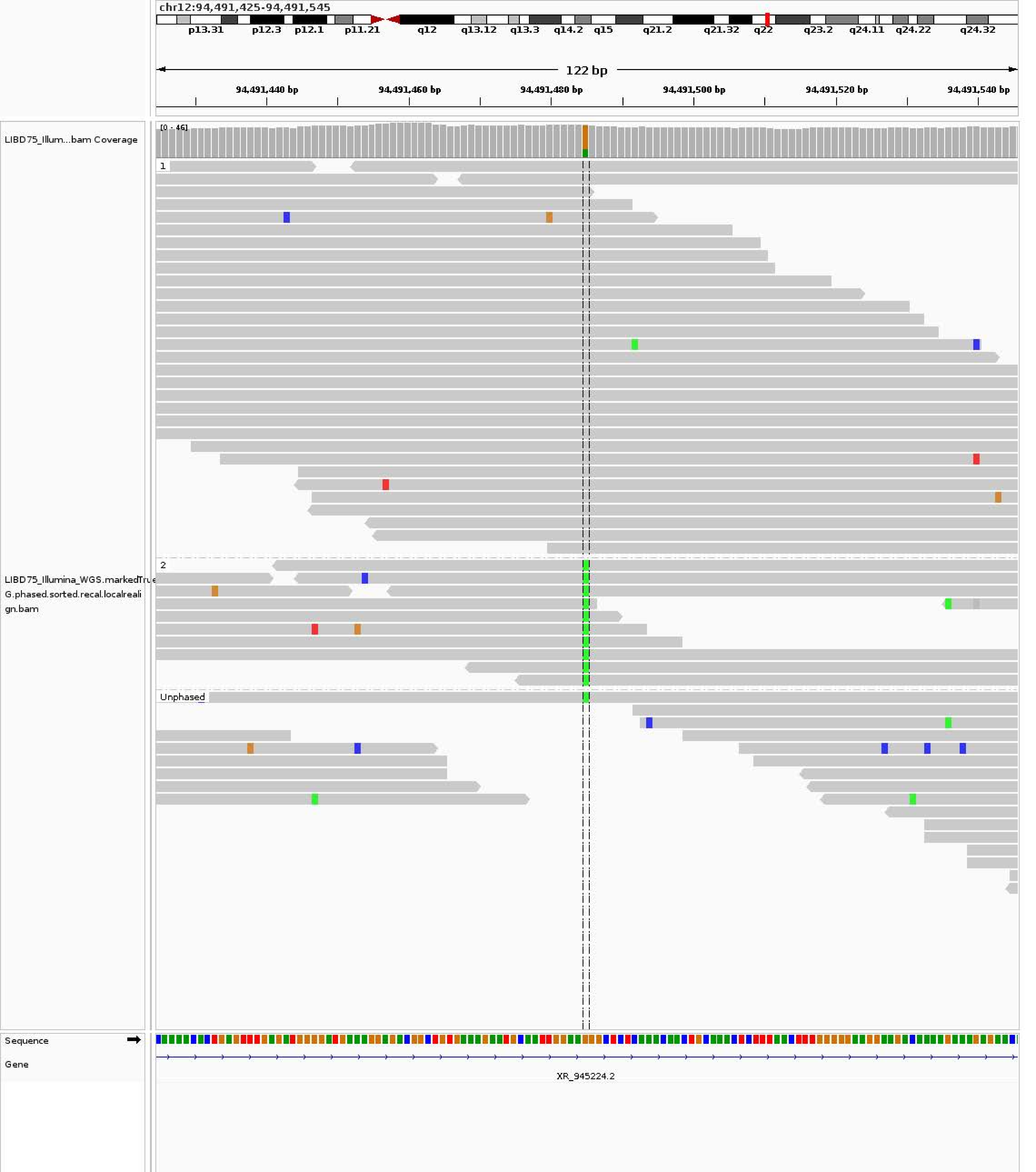
chr12:94,491,425-94,491,545
p13.31
p12.3
p12.1
p11.21
q12
q13.12
q13.3
q14.2
q15
q21.2
q21.32
q22
q23.2
q24.11
q24.22
q24.32
122 bp
94,491,440 bp
94,491,460 bp
94,491,480 bp
94,491,500 bp
94,491,520 bp
94,491,540 bp
LIBD75_Illum...bam Coverage
[0 - 46]
1
2
Unphased
LIBD75_Illumina_WGS.markedTrueG.phased.sorted.recal.localrealign.bam
Sequence
Gene
XR_945224.2

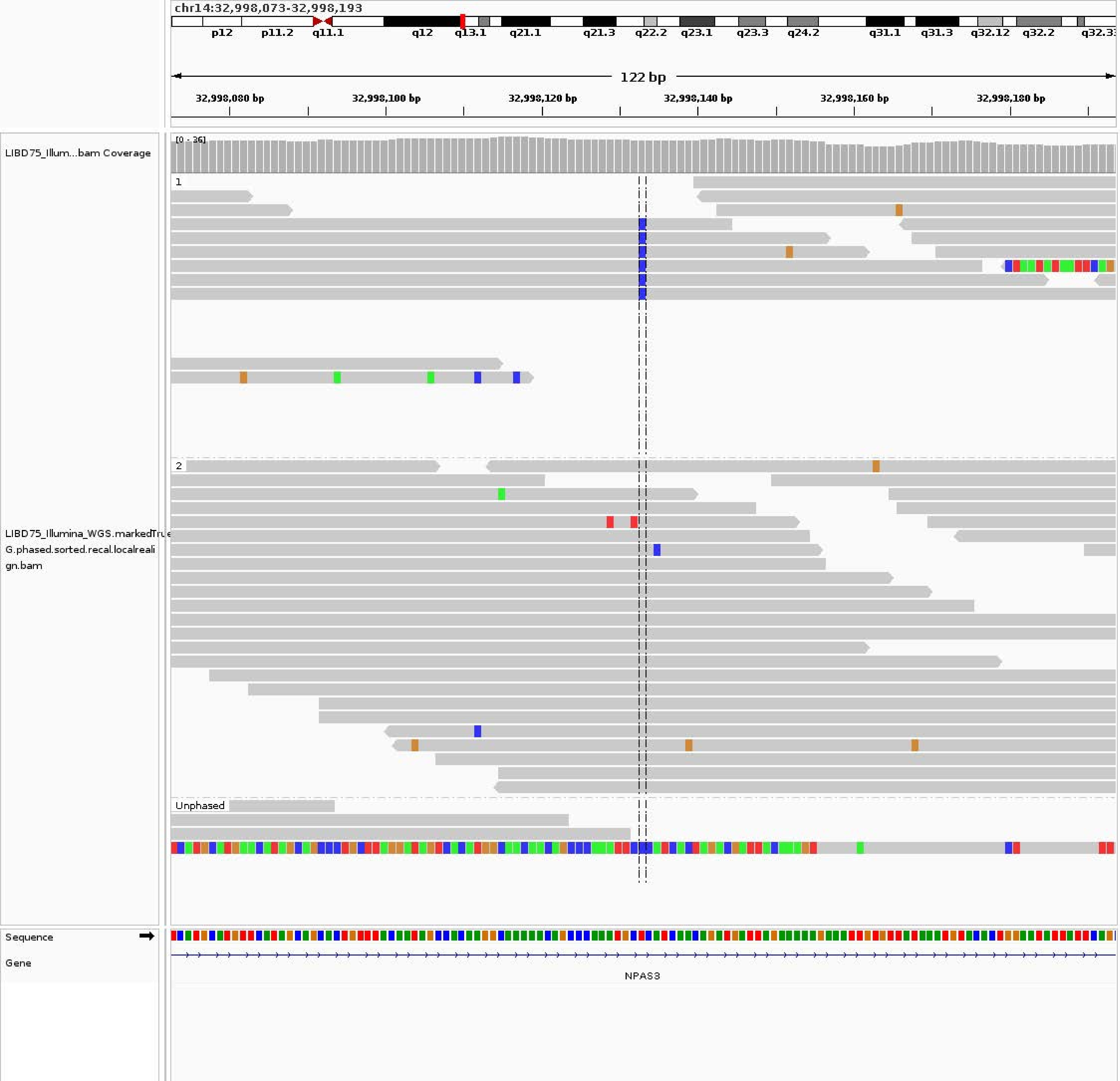

chr14:32,998,073-32,998,193
p12
p11.2
q11.1
q12
q13.1
q21.1
q21.3
q22.2
q23.1
q23.3
q24.2
q31.1
q31.3
q32.12
q32.2
122 bp
32,998,080 bp
32,998,100 bp
32,998,120 bp
32,998,140 bp
32,998,160 bp
32,998,180 bp
LIBD75_Illum...bam Coverage
[0 - 36]
1
2
Unphased
LIBD75_Illumina_WGS.markedTrue
G.phased.sorted.recal.localreali
gn.bam
Sequence
Gene
NPAS3

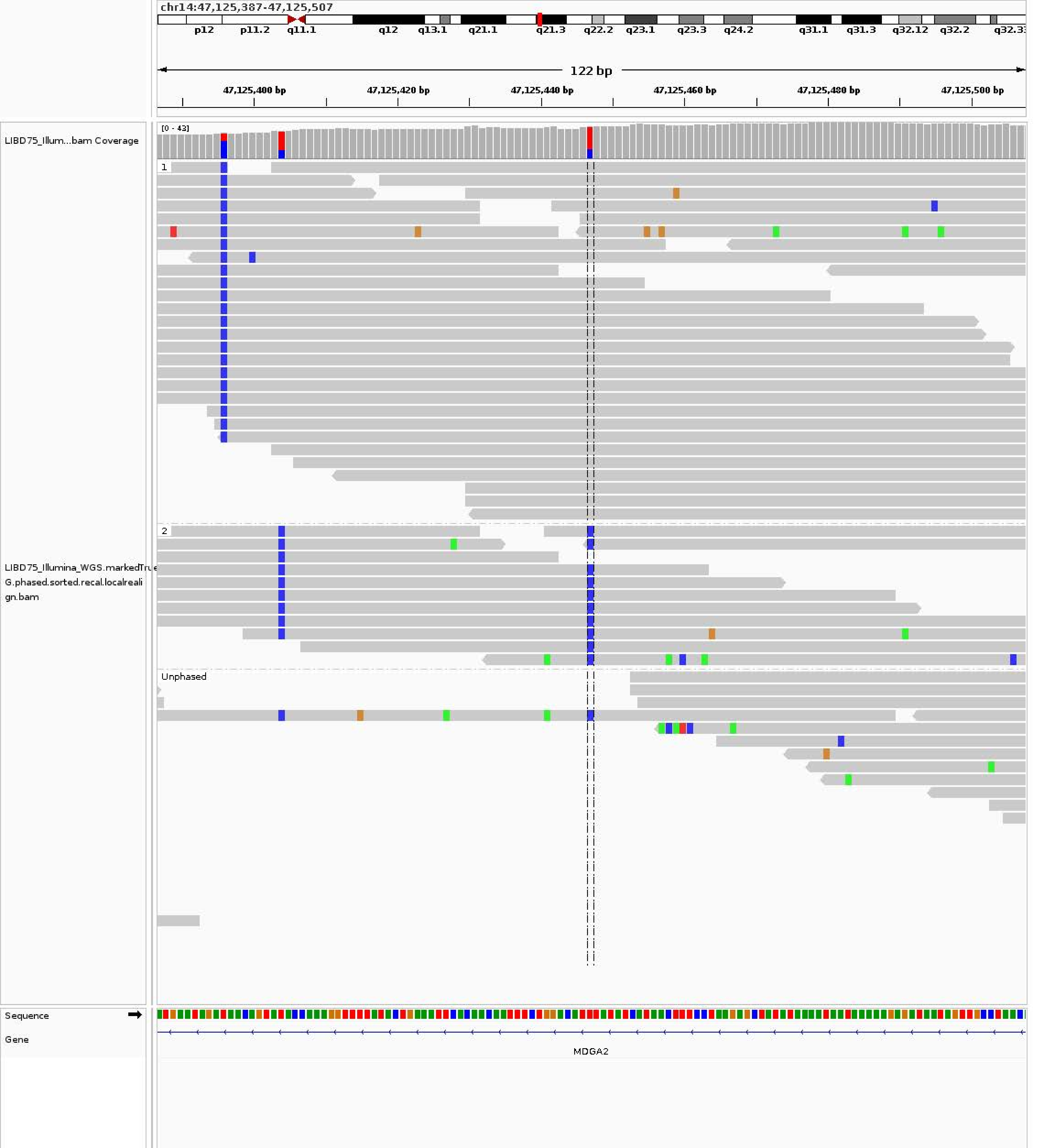
chr14:47,125,387-47,125,507
p12
p11.2
q11.1
q12
q13.1
q21.1
q21.3
q22.2
q23.1
q23.3
q24.2
q31.1
q31.3
q32.12
q32.2
122 bp
47,125,400 bp
47,125,420 bp
47,125,440 bp
47,125,460 bp
47,125,480 bp
47,125,500 bp
LIBD75_Illum...bam Coverage
[0 - 43]
1
2
Unphased
LIBD75_Illumina_WGS.markedTrue
G.phased.sorted.recal.localreali
gn.bam
Sequence
Gene
MDGA2

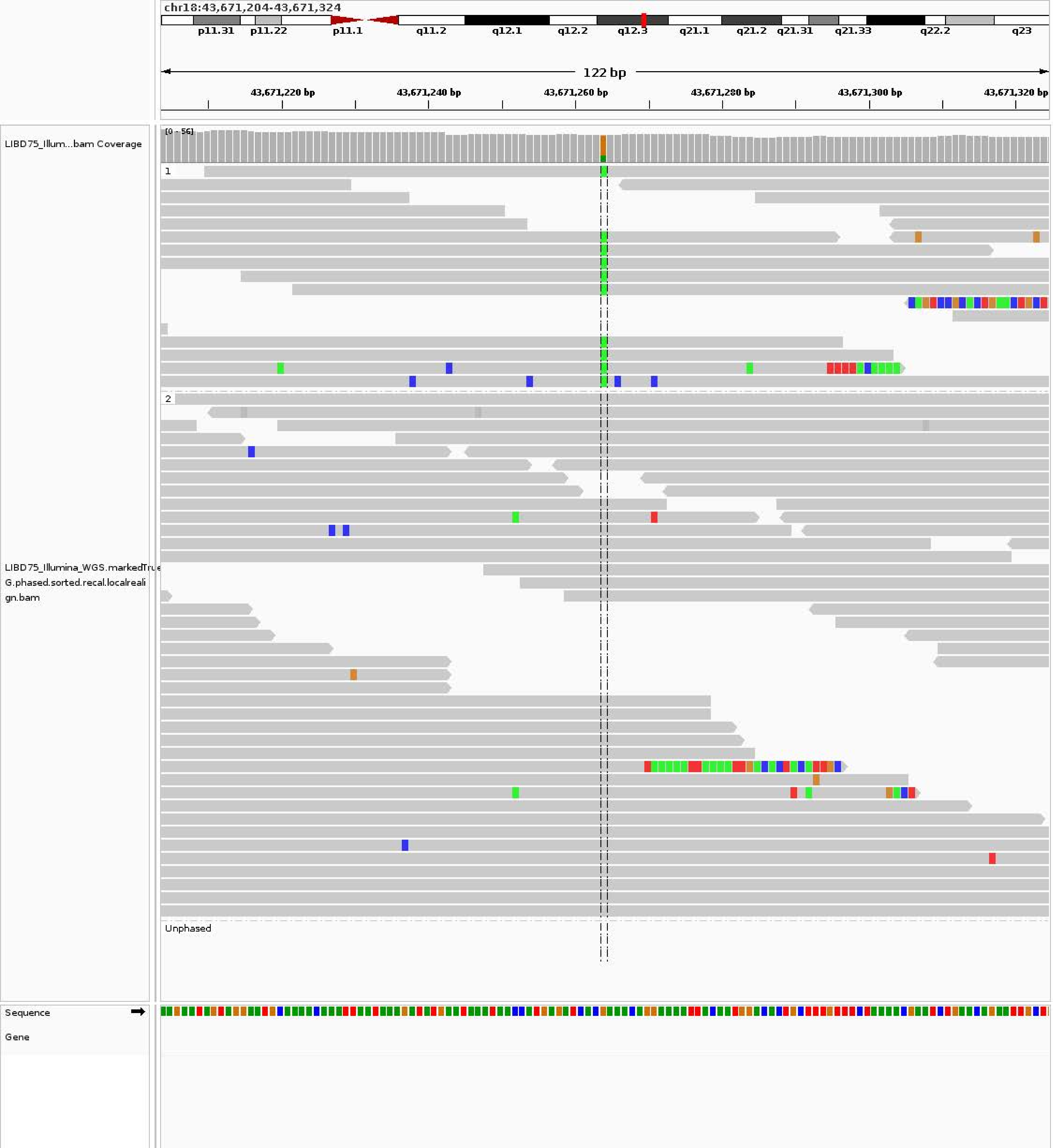

chr18:43,671,204-43,671,324
p11.31
p11.22
p11.1
q11.2
q12.1
q12.2
q12.3
q21.1
q21.2
q21.31
q21.33
q22.2
q23
122 bp
43,671,220 bp
43,671,240 bp
43,671,260 bp
43,671,280 bp
43,671,300 bp
43,671,320 bp
LIBD75_Illum...bam Coverage
[0 - 56]
1
2
Unphased
LIBD75_Illumina_WGS.markedTrue G.phased.sorted.recal.localrealign.bam
Sequence
Gene

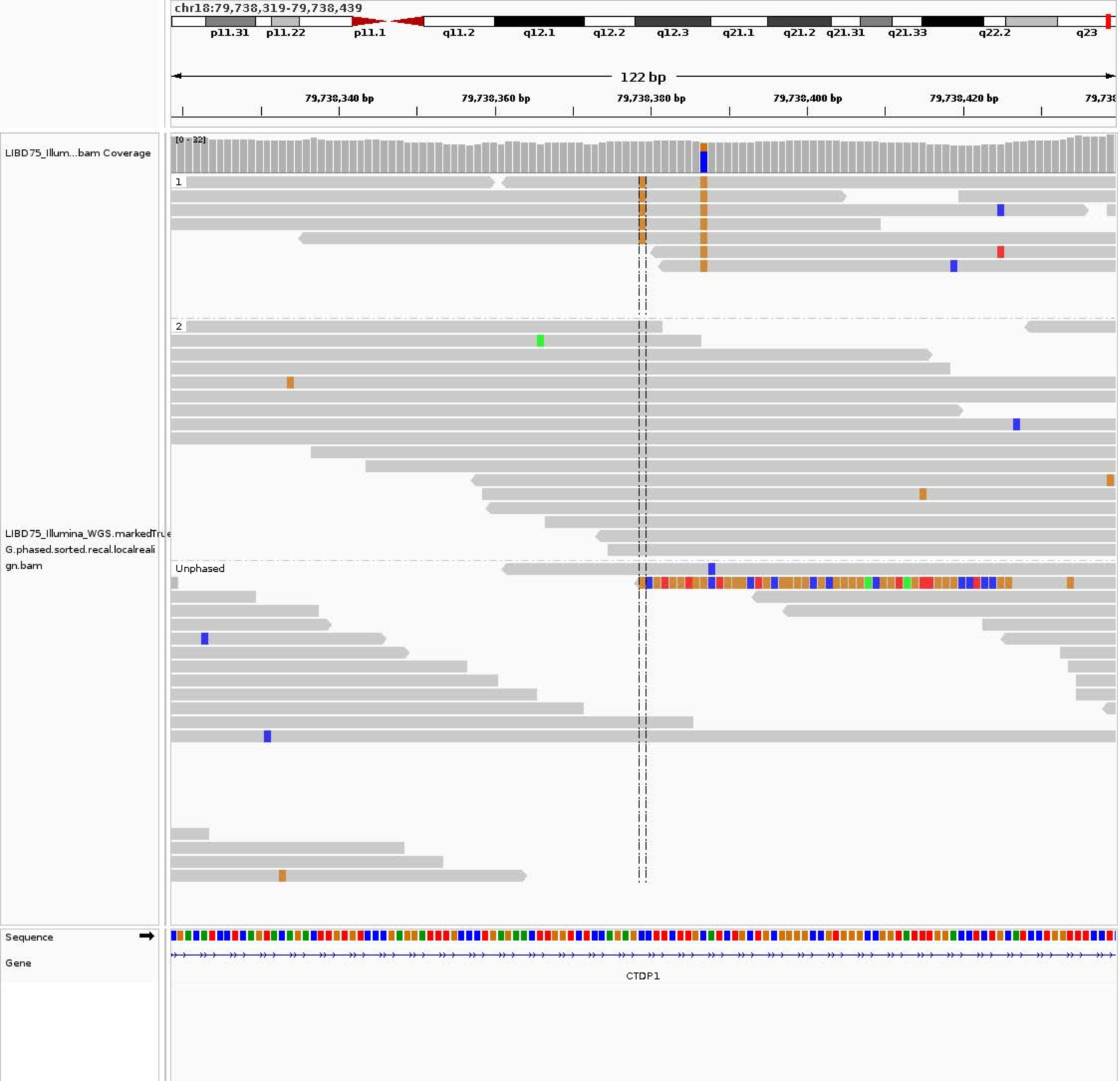

chr18:79,738,319-79,738,439
p11.31
p11.22
p11.1
q11.2
q12.1
q12.2
q12.3
q21.1
q21.2
q21.31
q21.33
q22.2
q23
122 bp
79,738,340 bp
79,738,360 bp
79,738,380 bp
79,738,400 bp
79,738,420 bp
LIBD75_Illum...bam Coverage
[0 - 32]
1
2
Unphased
LIBD75_Illumina_WGS.markedTrue
G.phased.sorted.recal.localreali
gn.bam
Sequence
Gene
CTDP1

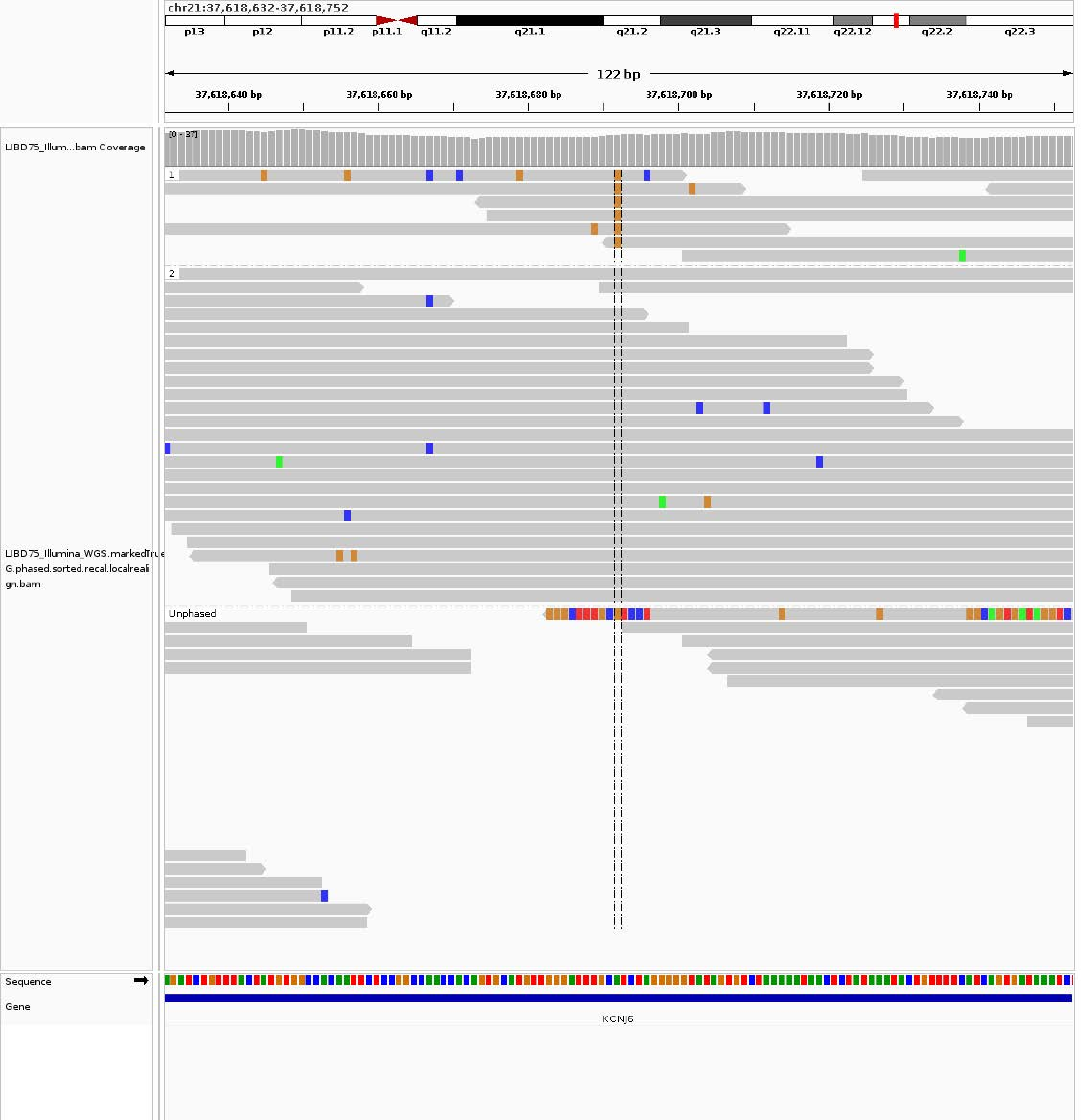
chr21:37,618,632-37,618,752
p13
p12
p11.2
p11.1
q11.2
q21.1
q21.2
q21.3
q22.11
q22.12
q22.2
q22.3
122 bp
37,618,640 bp
37,618,660 bp
37,618,680 bp
37,618,700 bp
37,618,720 bp
37,618,740 bp
LIBD75_Illum...bam Coverage
[0 - 37]
1
2
Unphased
LIBD75_Illumina_WGS.markedTrue G.phased.sorted.recal.localrealign.bam
Sequence
Gene
KCNJ6

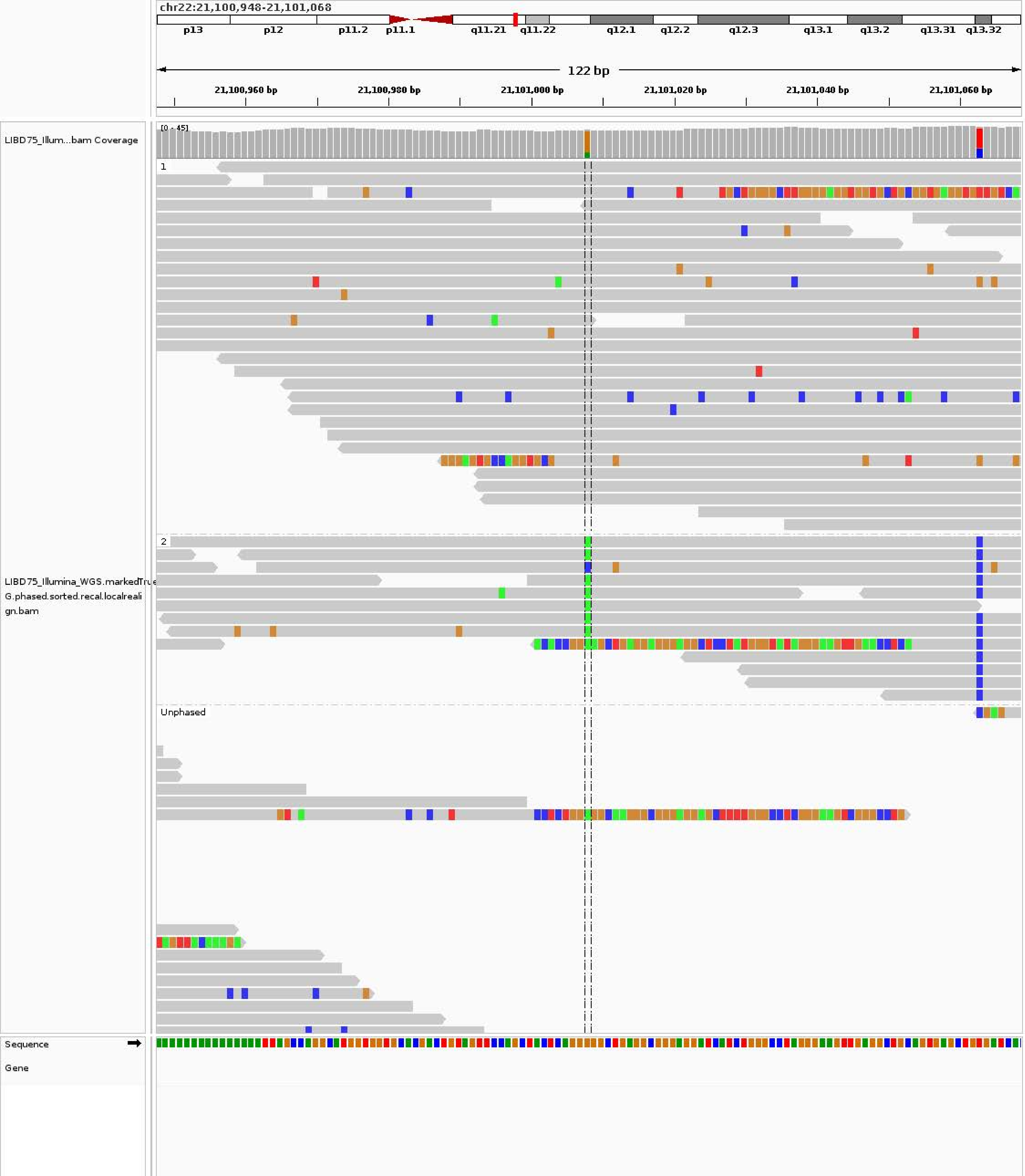
chr22:21,100,948-21,101,068
p13
p12
p11.2
p11.1
q11.21
q11.22
q12.1
q12.2
q12.3
q13.1
q13.2
q13.31
q13.32
122 bp
21,100,960 bp
21,100,980 bp
21,101,000 bp
21,101,020 bp
21,101,040 bp
21,101,060 bp
LIBD75_Illum...bam Coverage
[0 - 45]
LIBD75_Illumina_WGS.markedTrueG.phased.sorted.recal.localrealign.bam
1
2
Unphased
Sequence
Gene
