## Supplementary material for "A personalized multi-platform assessment of somatic mosaicism in the human frontal cortex": Supplementary Table 6.pdf

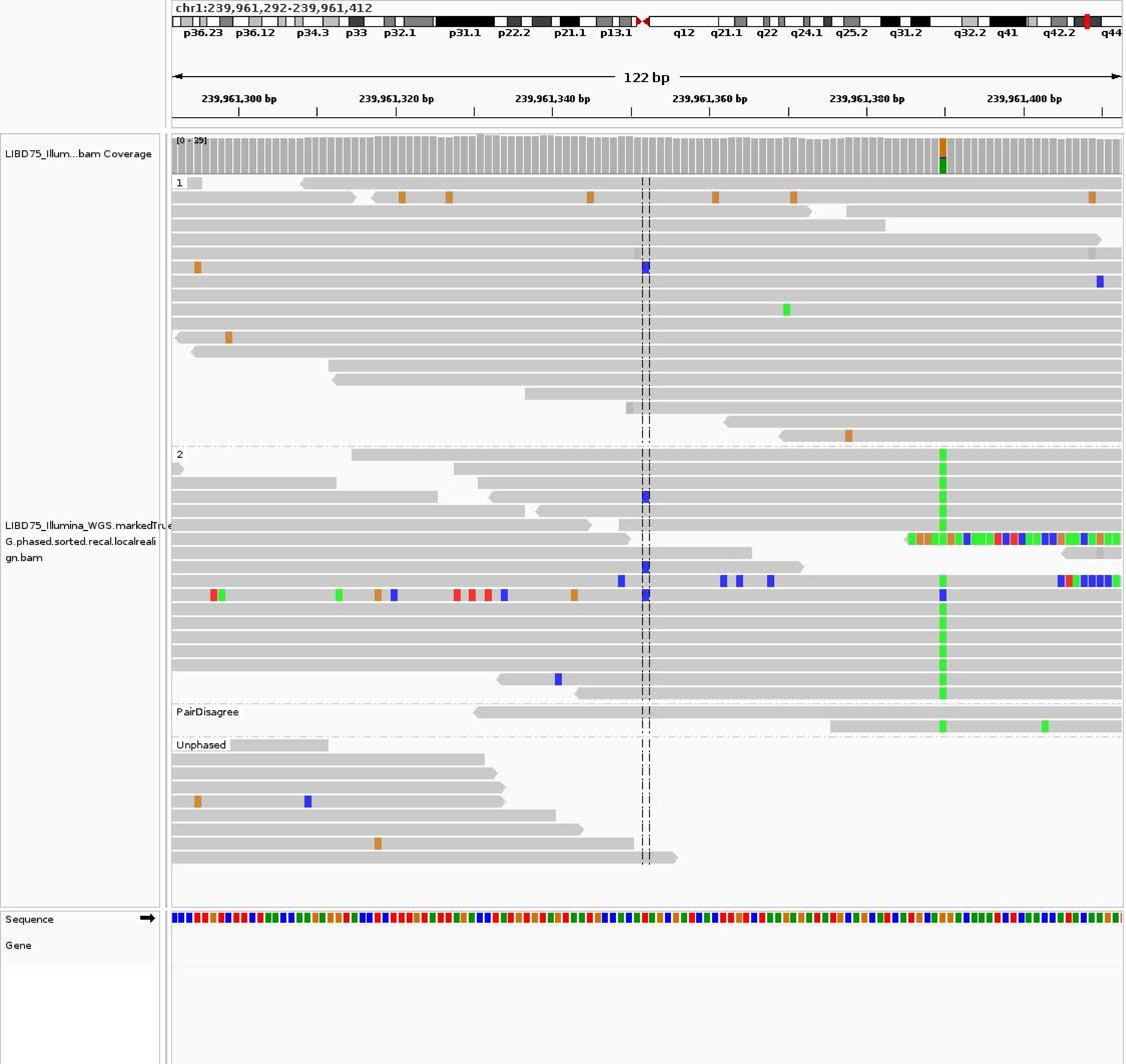
chr1:239,961,292-239,961,412
p36.23
p36.12
p34.3
p33
p32.1
p31.1
p22.2
p21.1
p13.1
q12
q21.1
q22
q24.1
q25.2
q31.2
q32.2
q41
q42.2
q44
122 bp
239,961,300 bp
239,961,320 bp
239,961,340 bp
239,961,360 bp
239,961,380 bp
239,961,400 bp
LIBD75_Illum...bam Coverage
[0 - 39]
1
2
PairDisagree
Unphased
LIBD75_Illumina_WGS.markedTrue
G.phased.sorted.recal.localreali
gn.bam
Sequence
Gene

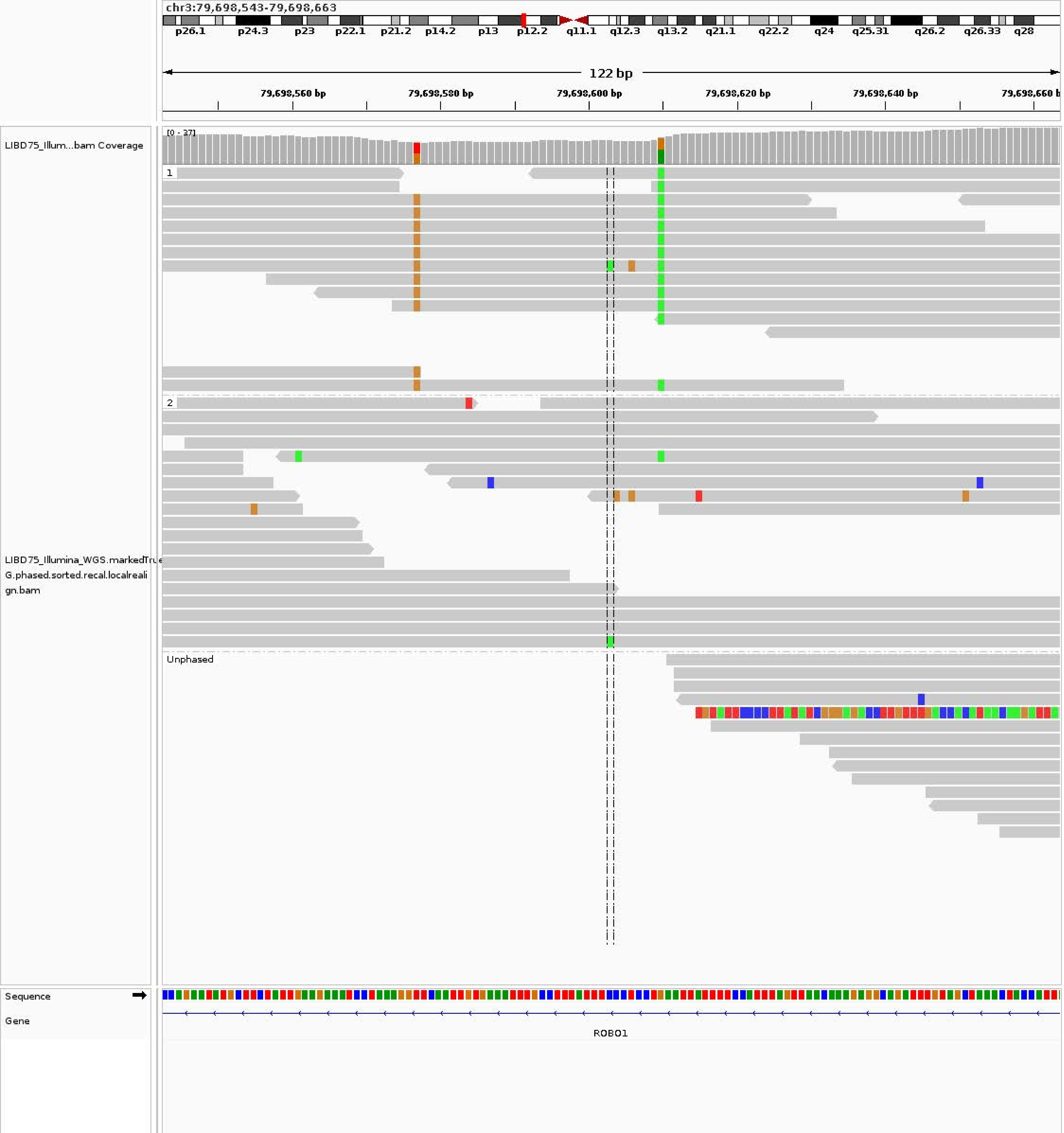
chr3:79,698,543-79,698,663
p26.1 p24.3 p23 p22.1 p21.2 p14.2 p13 p12.2 q11.1 q12.3 q13.2 q21.1 q22.2 q24 q25.31 q26.2 q26.33 q28
122 bp
79,698,560 bp
79,698,580 bp
79,698,600 bp
79,698,620 bp
79,698,640 bp
79,698,660 b
LIBD75_Illum...bam Coverage
[0 - 37]
1
2
Unphased
LIBD75_Illumina_WGS.markedTru
G.phased.sorted.recal.localreali
gn.bam
Sequence
Gene
ROBO1

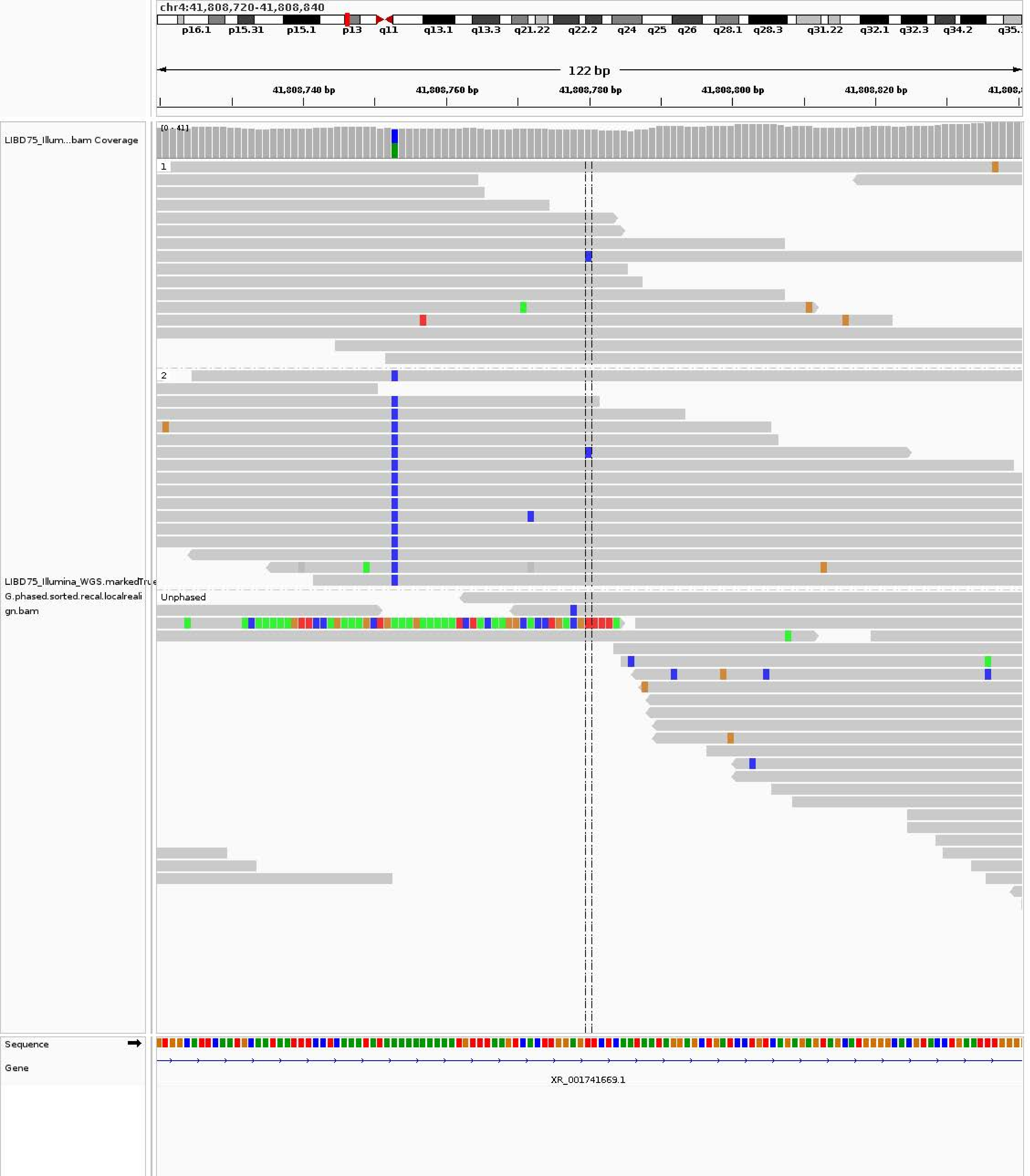
chr4:41,808,720-41,808,840
p16.1 p15.31 p15.1 p13 q11 q13.1 q13.3 q21.22 q22.2 q24 q25 q26 q28.1 q28.3 q31.22 q32.1 q32.3 q34.2
122 bp
41,808,740 bp
41,808,760 bp
41,808,780 bp
41,808,800 bp
41,808,820 bp
LIBD75_Illum...bam Coverage
[0 - 41]
1
2
Unphased
LIBD75_Illumina_WGS.markedTrue G.phased.sorted.recal.localrealign.bam
Sequence
Gene
XR_001741669.1

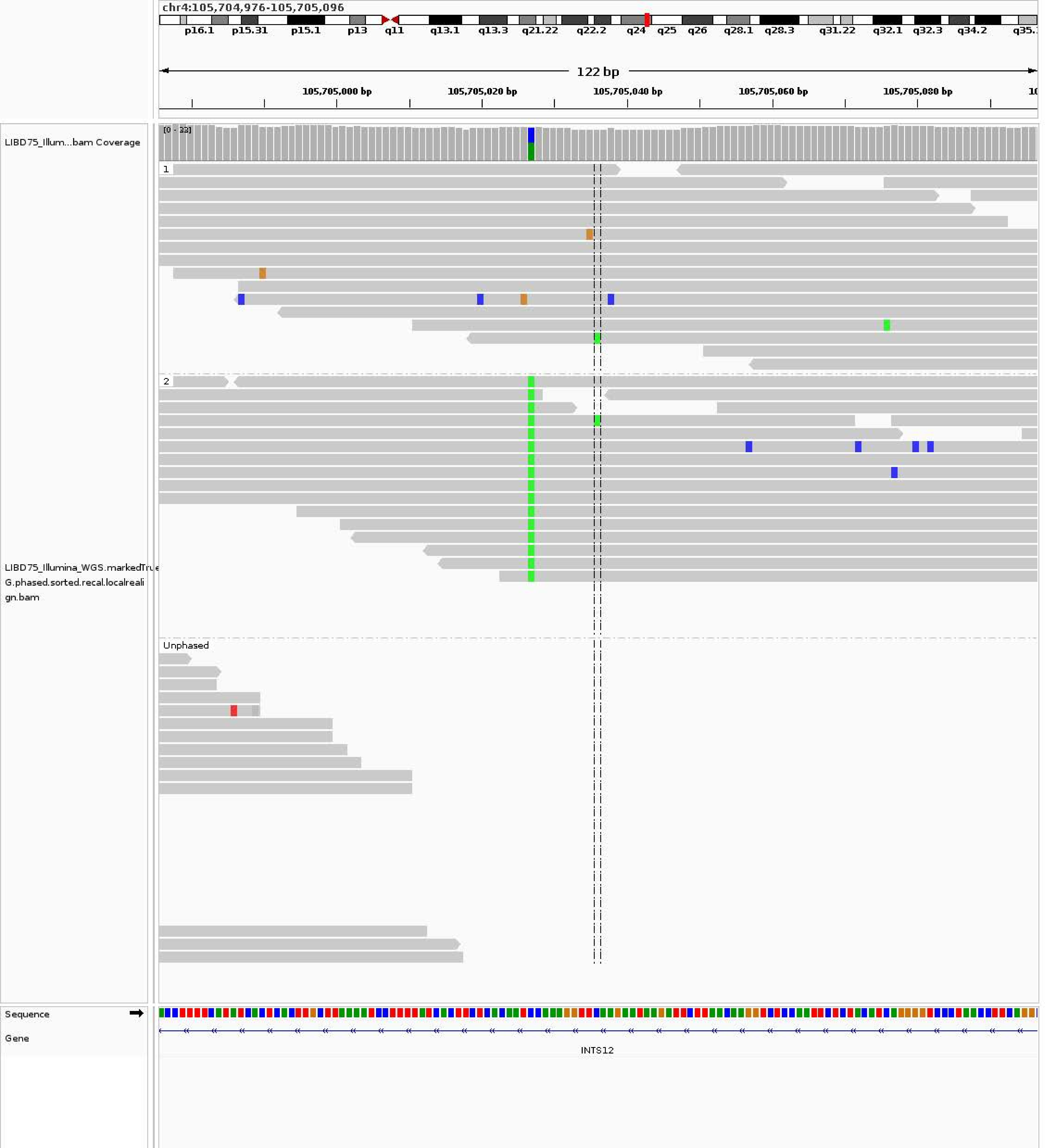
chr4:105,704,976-105,705,096
p16.1 p15.31 p15.1 p13 q11 q13.1 q13.3 q21.22 q22.2 q24 q25 q26 q28.1 q28.3 q31.22 q32.1 q32.3 q34.2
122 bp
105,705,000 bp
105,705,020 bp
105,705,040 bp
105,705,060 bp
105,705,080 bp
LIBD75_Illum...bam Coverage
[0 - 33]
1
2
Unphased
LIBD75_Illumina_WGS.markedTru
G.phased.sorted.recal.localreali
gn.bam
Sequence
Gene
INTS12

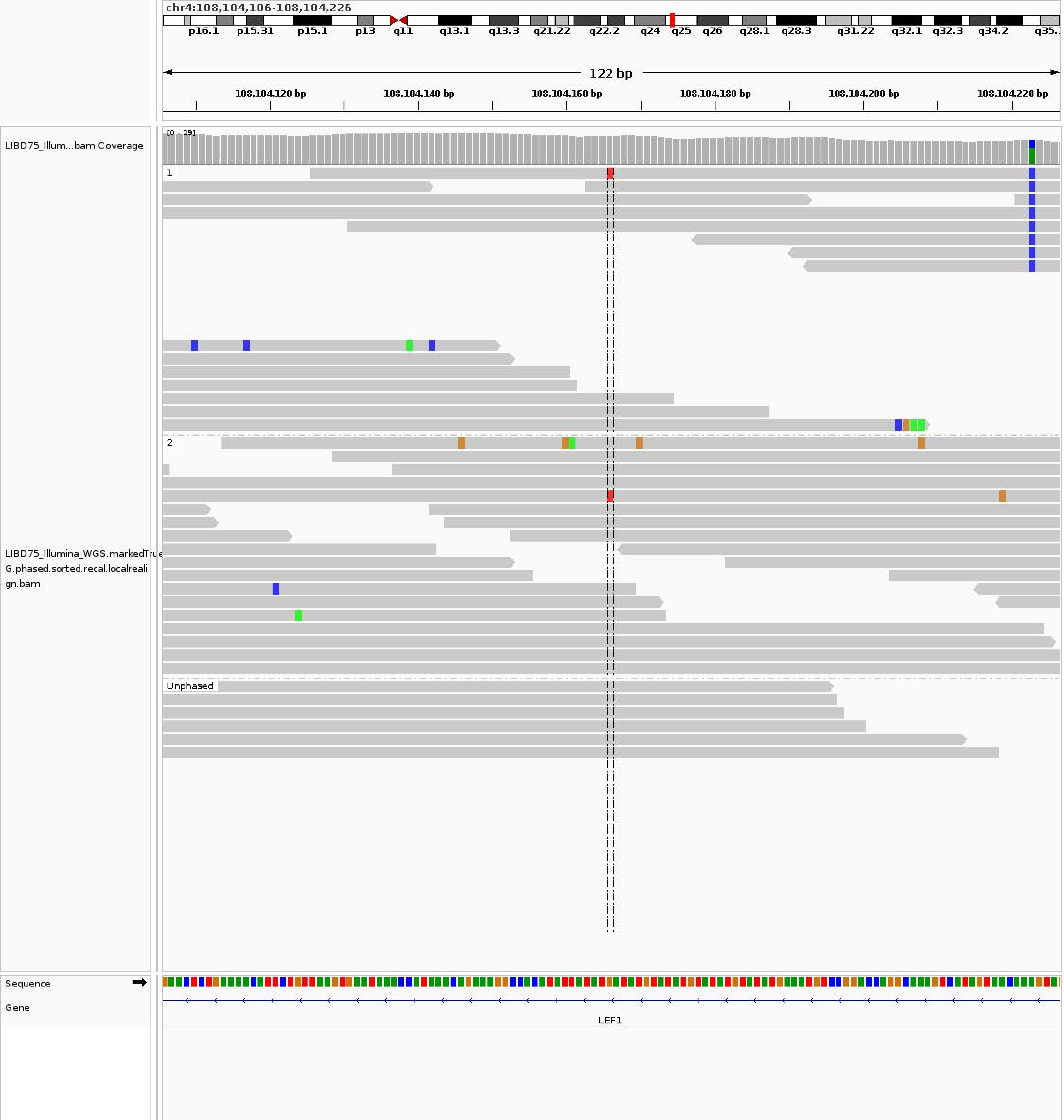
chr4:108,104,106-108,104,226
p16.1 p15.31 p15.1 p13 q11 q13.1 q13.3 q21.22 q22.2 q24 q25 q26 q28.1 q28.3 q31.22 q32.1 q32.3 q34.2
122 bp
108,104,120 bp
108,104,140 bp
108,104,160 bp
108,104,180 bp
108,104,200 bp
108,104,220 bp
LIBD75_Illum...bam Coverage
[0 - 39]
1
2
Unphased
LIBD75_Illumina_WGS.markedTru
G.phased.sorted.recal.localreali
gn.bam
Sequence
Gene
LEF1

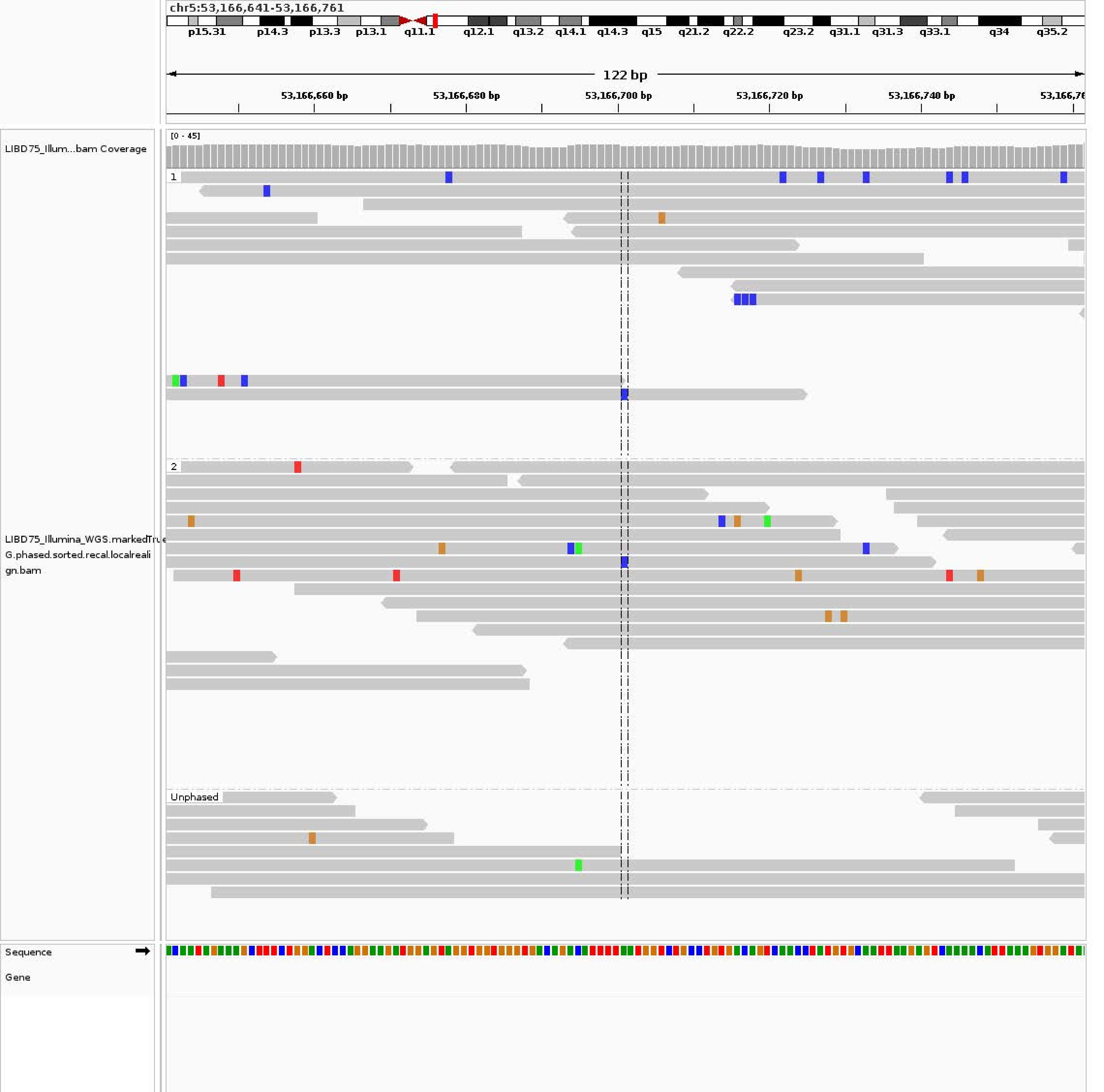
chr5:53,166,641-53,166,761
p15.31
p14.3
p13.3
p13.1
q11.1
q12.1
q13.2
q14.1
q14.3
q15
q21.2
q22.2
q23.2
q31.1
q31.3
q33.1
q34
q35.2
122 bp
53,166,660 bp
53,166,680 bp
53,166,700 bp
53,166,720 bp
53,166,740 bp
LIBD75_Illum...bam Coverage
[0 - 45]
1
2
Unphased
LIBD75_Illumina_WGS.markedTrue
G.phased.sorted.recal.localreali
gn.bam
Sequence
Gene

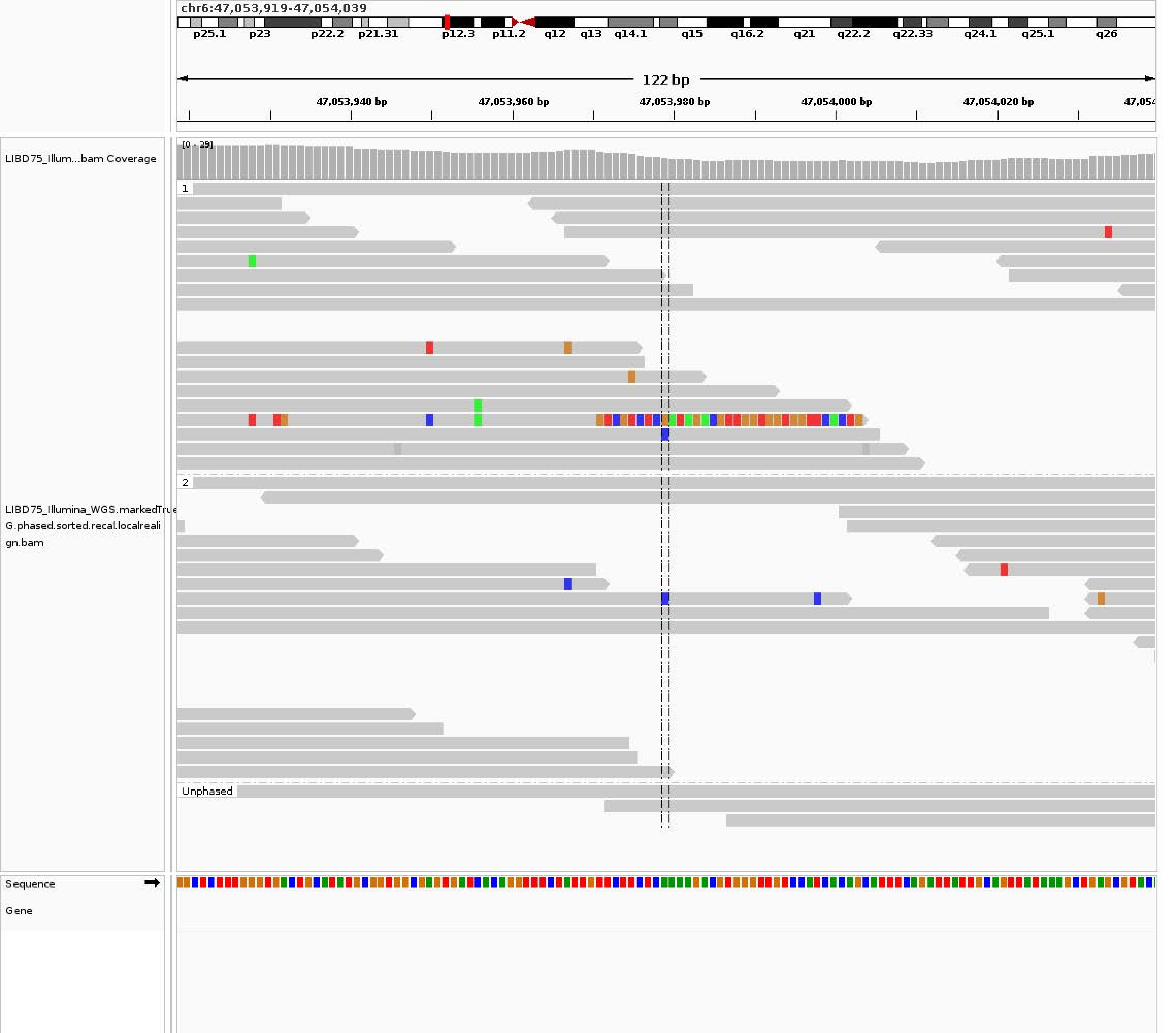
chr6:47,053,919-47,054,039
p25.1 p23 p22.2 p21.31 p12.3 p11.2 q12 q13 q14.1 q15 q16.2 q21 q22.2 q22.33 q24.1 q25.1 q26
122 bp
47,053,940 bp
47,053,960 bp
47,053,980 bp
47,054,000 bp
47,054,020 bp
LIBD75_Illum...bam Coverage
[0 - 39]
1
2
Unphased
LIBD75_Illumina_WGS.markedTrue
G.phased.sorted.recal.localreali
gn.bam
Sequence
Gene

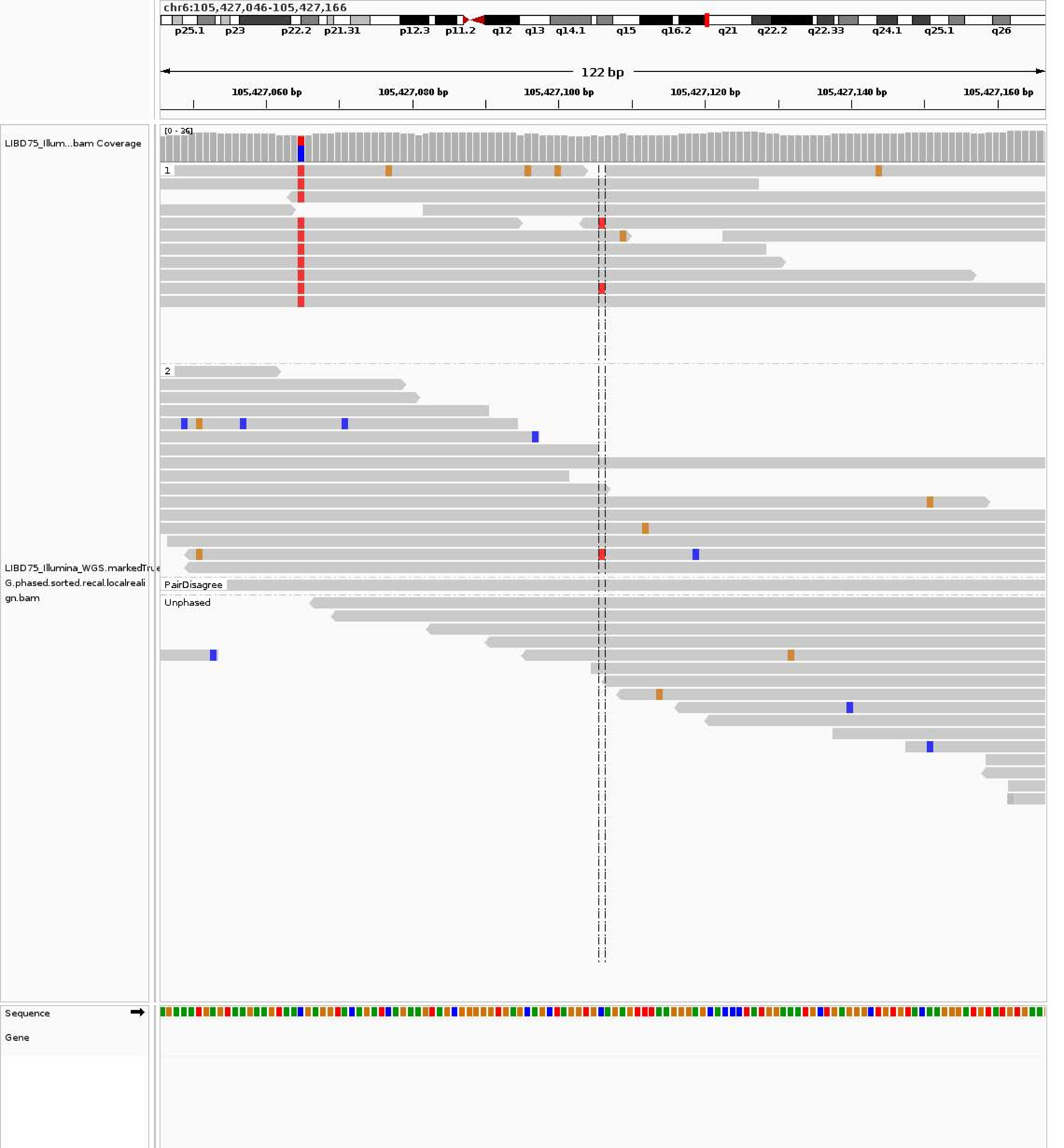
chr6:105,427,046-105,427,166
p25.1 p23 p22.2 p21.31 p12.3 p11.2 q12 q13 q14.1 q15 q16.2 q21 q22.2 q22.33 q24.1 q25.1 q26
122 bp
105,427,060 bp 105,427,080 bp 105,427,100 bp 105,427,120 bp 105,427,140 bp 105,427,160 bp
LIBD75_Illum...bam Coverage
[0 - 36]
1
2
LIBD75_Illumina_WGS.markedTrue
G.phased.sorted.recal.localreali
gn.bam
PairDisagree
Unphased
Sequence
Gene

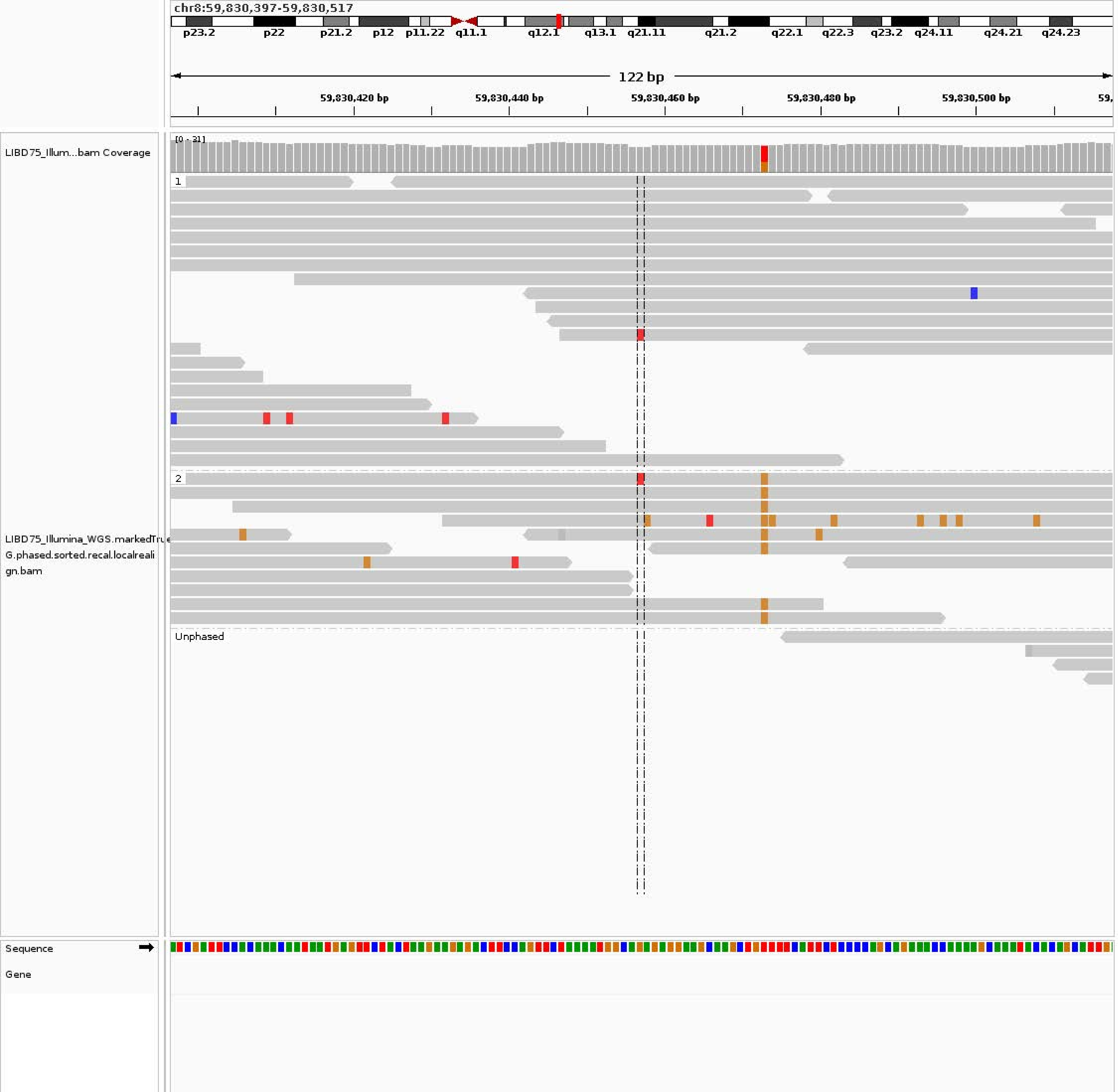
chr8:59,830,397-59,830,517
p23.2
p22
p21.2
p12
p11.22
q11.1
q12.1
q13.1
q21.11
q21.2
q22.1
q22.3
q23.2
q24.11
q24.21
q24.23
122 bp
59,830,420 bp
59,830,440 bp
59,830,460 bp
59,830,480 bp
59,830,500 bp
LIBD75_Illum...bam Coverage
[0 - 31]
1
2
Unphased
LIBD75_Illumina_WGS.markedTrue
G.phased.sorted.recal.localreali
gn.bam
Sequence
Gene

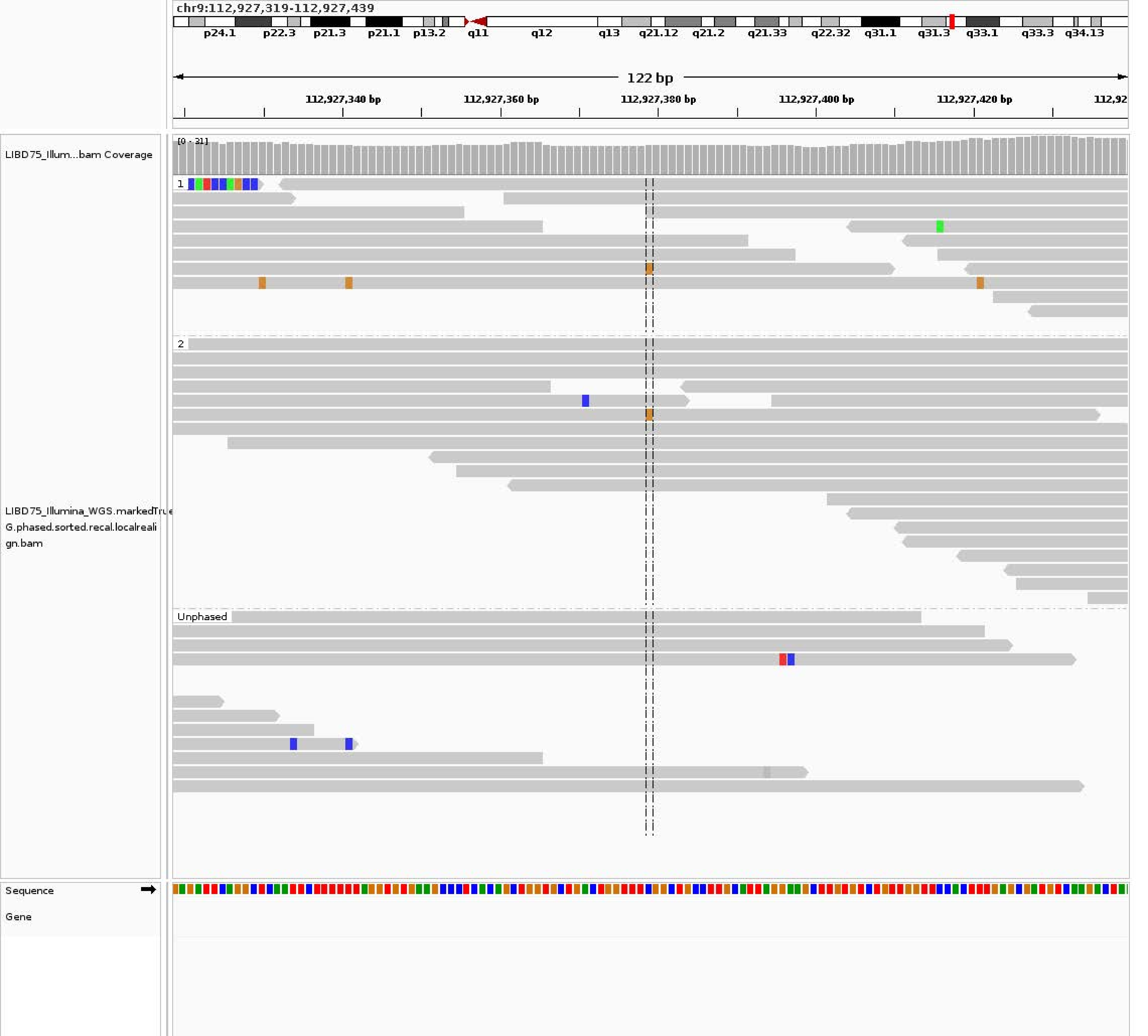
chr9:112,927,319-112,927,439
p24.1
p22.3
p21.3
p21.1
p13.2
q11
q12
q13
q21.12
q21.2
q21.33
q22.32
q31.1
q31.3
q33.1
q33.3
q34.13
122 bp
112,927,340 bp
112,927,360 bp
112,927,380 bp
112,927,400 bp
112,927,420 bp
112,92
LIBD75_Illum...bam Coverage
[0 - 31]
1
2
Unphased
LIBD75_Illumina_WGS.markedTru
G.phased.sorted.recal.localreali
gn.bam
Sequence
Gene

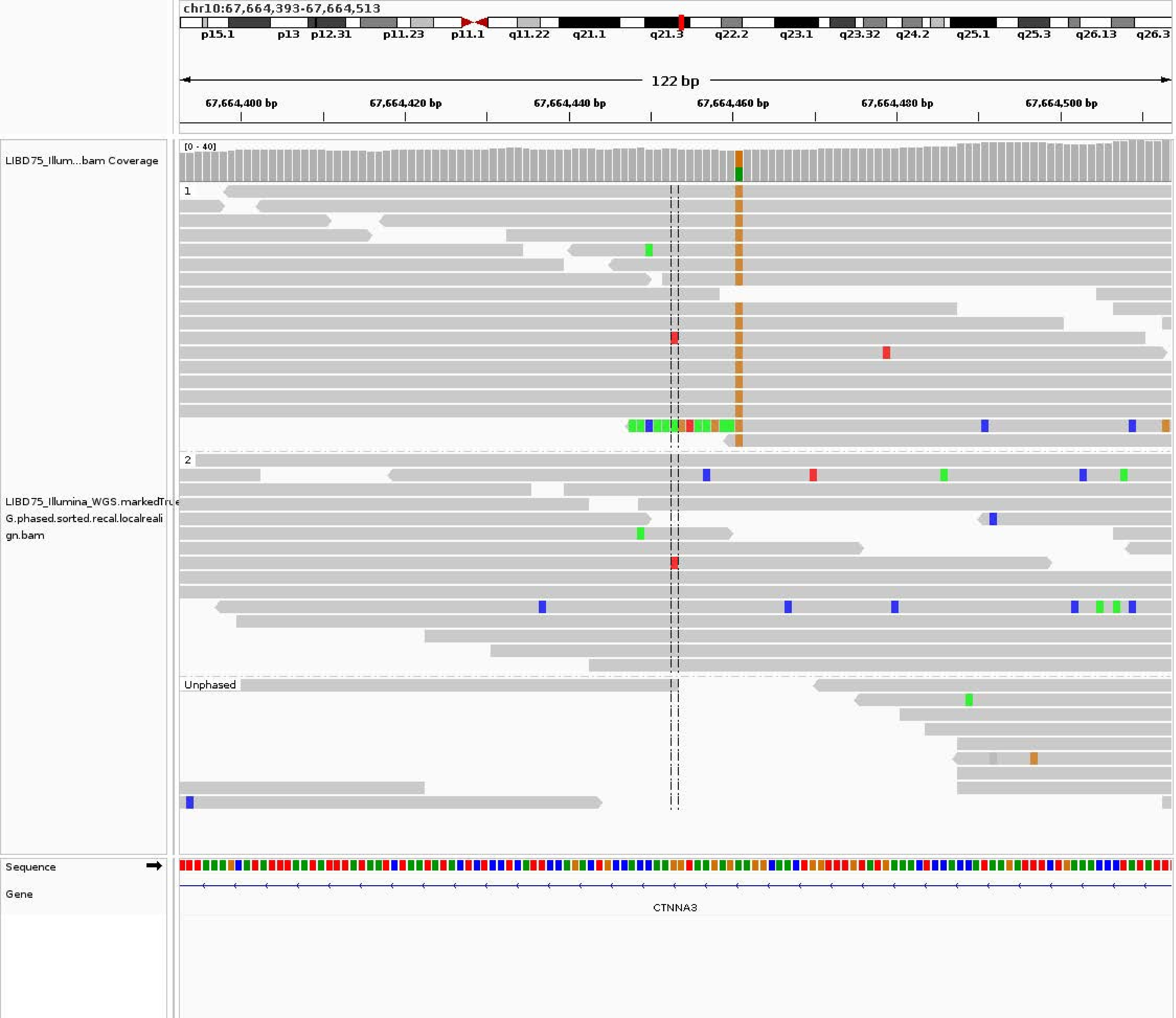
chr10:67,664,393-67,664,513
p15.1
p13
p12.31
p11.23
p11.1
q11.22
q21.1
q21.3
q22.2
q23.1
q23.32
q24.2
q25.1
q25.3
q26.13
q26.3
122 bp
67,664,400 bp
67,664,420 bp
67,664,440 bp
67,664,460 bp
67,664,480 bp
67,664,500 bp
LIBD75_Illum...bam Coverage
[0 - 40]
1
2
Unphased
LIBD75_Illumina_WGS.markedTrue
G.phased.sorted.recal.localreali
gn.bam
Sequence
Gene
CTNNA3

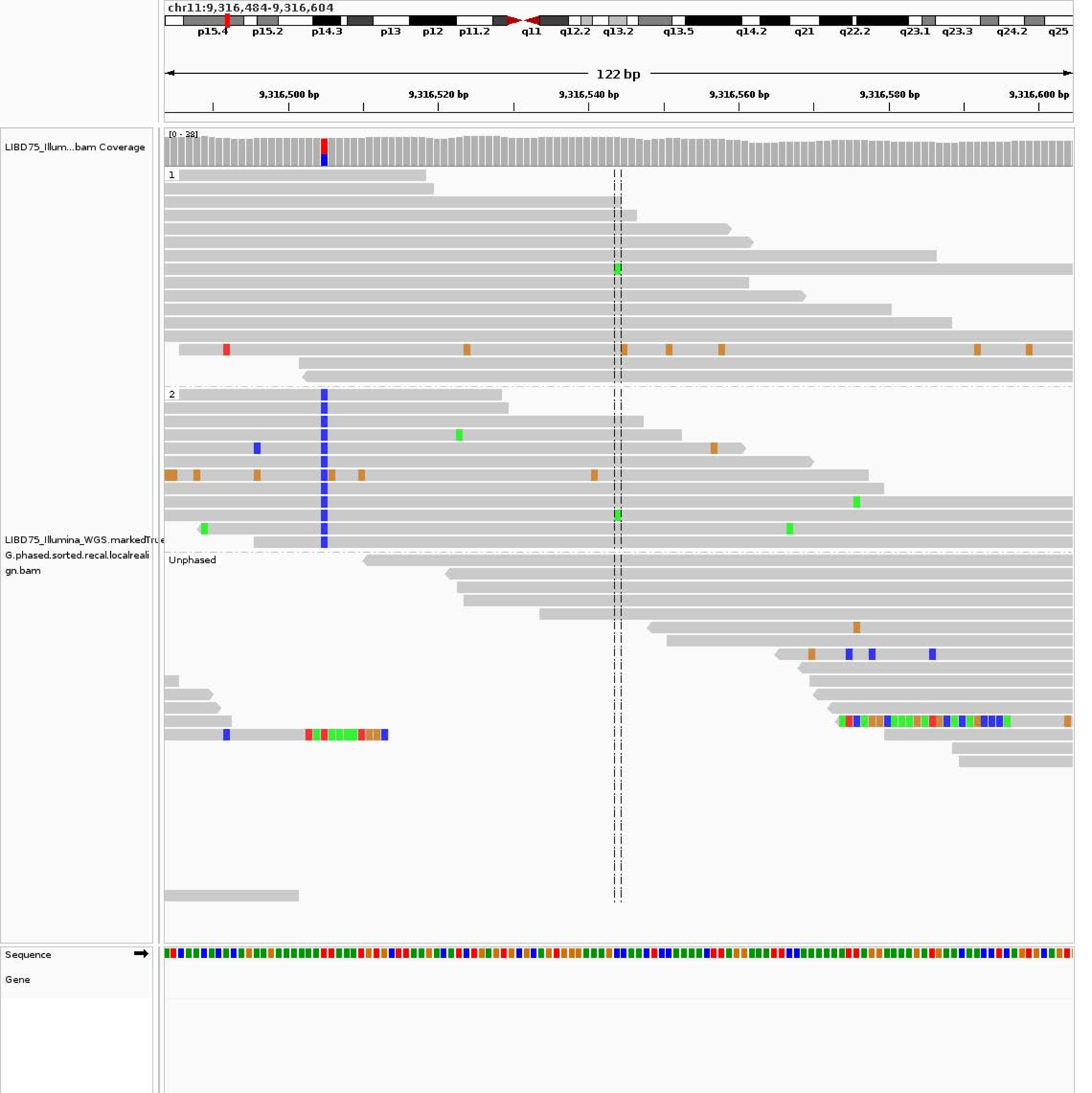
chr11:9,316,484-9,316,604
p15.4
p15.2
p14.3
p13
p12
p11.2
q11
q12.2
q13.2
q13.5
q14.2
q21
q22.2
q23.1
q23.3
q24.2
q25
122 bp
9,316,500 bp
9,316,520 bp
9,316,540 bp
9,316,560 bp
9,316,580 bp
9,316,600 bp
LIBD75_Illum...bam Coverage
[0 - 38]
1
2
Unphased
LIBD75_Illumina_WGS.markedTrue
G.phased.sorted.recal.localreali
gn.bam
Sequence
Gene

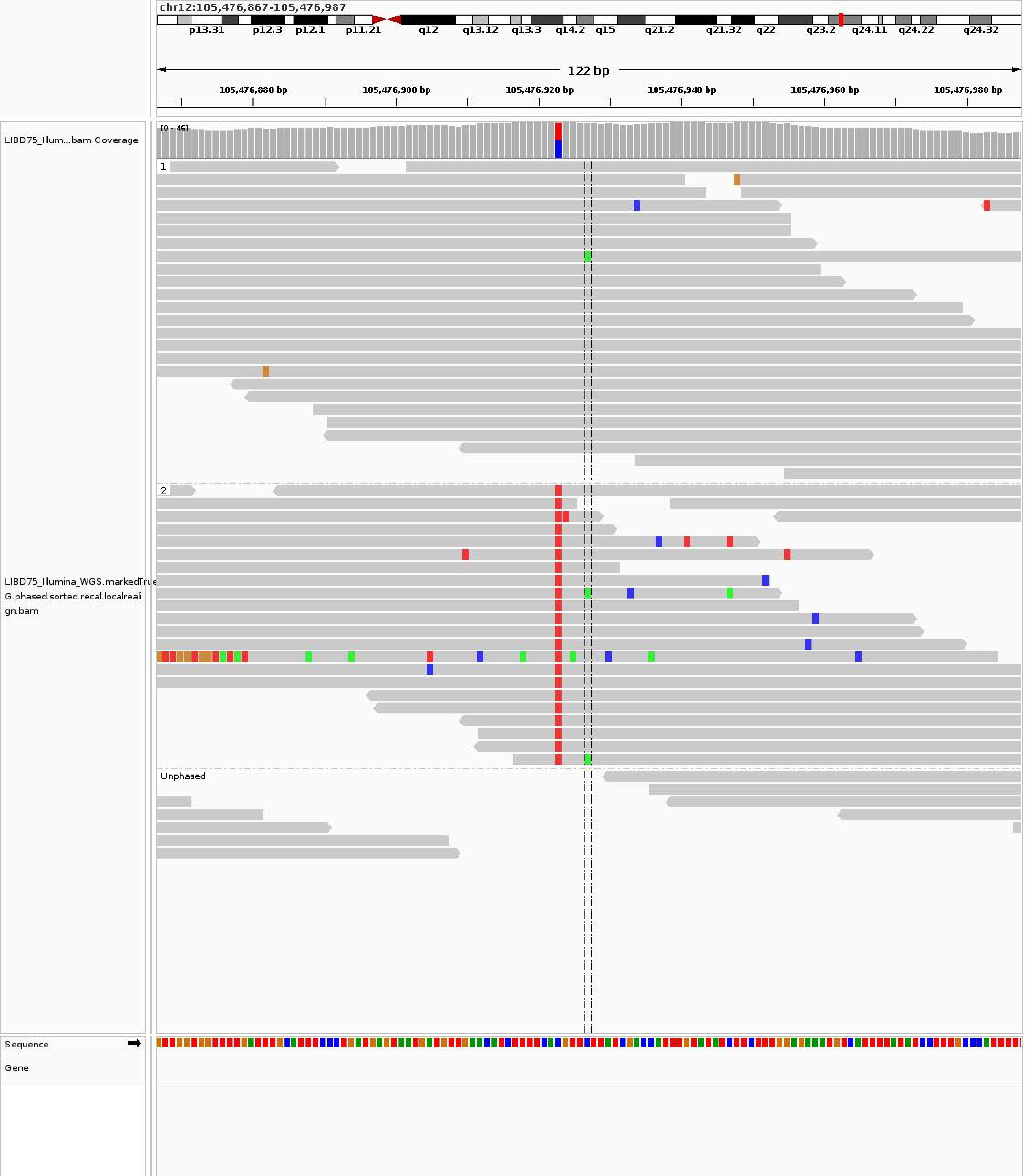

chr12:105,476,867-105,476,987
p13.31
p12.3
p12.1
p11.21
q12
q13.12
q13.3
q14.2
q15
q21.2
q21.32
q22
q23.2
q24.11
q24.22
q24.32
122 bp
105,476,880 bp
105,476,900 bp
105,476,920 bp
105,476,940 bp
105,476,960 bp
105,476,980 bp
LIBD75_Illum...bam Coverage
[0 - 46]
1
2
Unphased
LIBD75_Illumina_WGS.markedTrue
G.phased.sorted.recal.localreali
gn.bam
Sequence
Gene

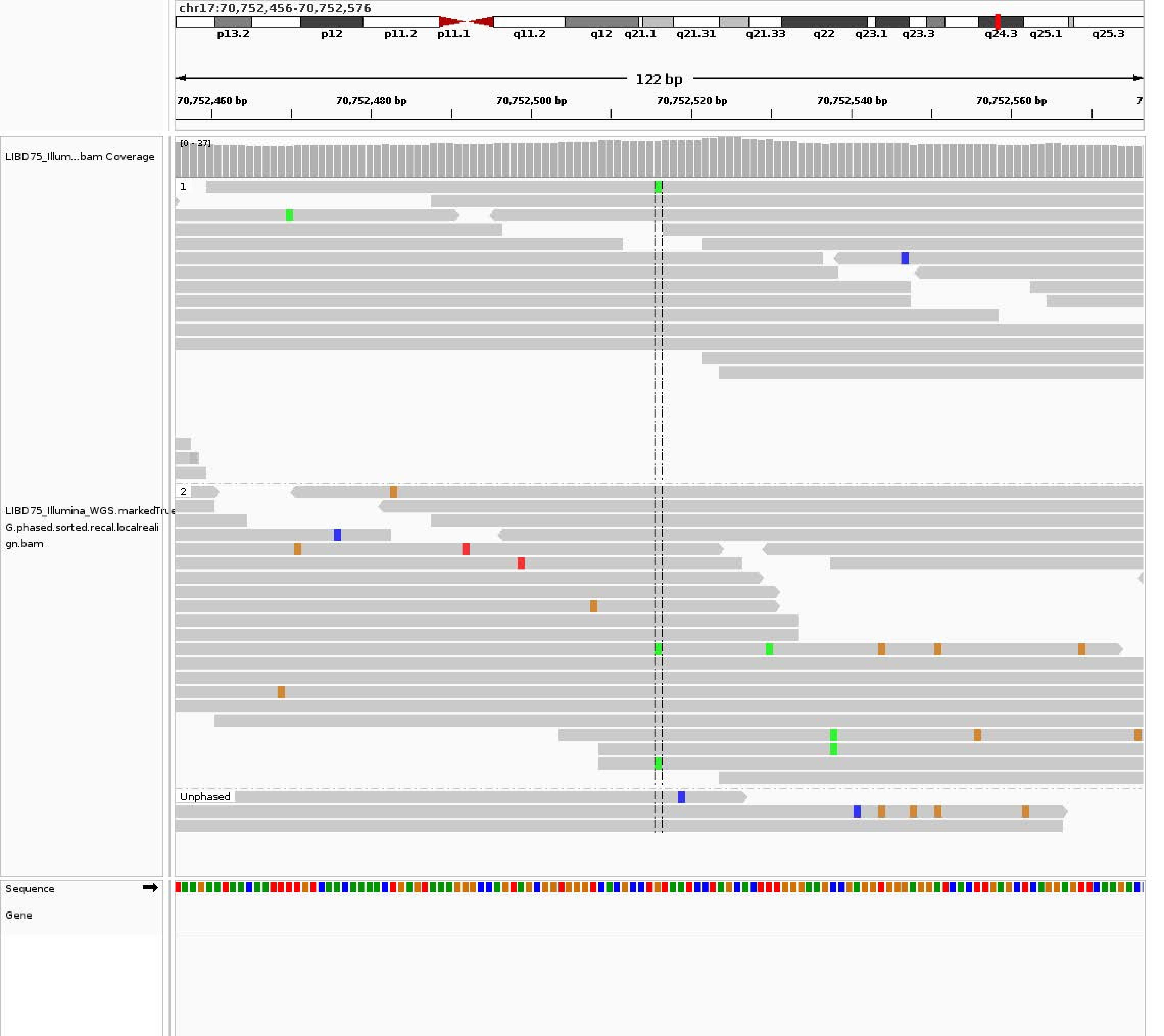
chr17:70,752,456-70,752,576
p13.2
p12
p11.2
p11.1
q11.2
q12
q21.1
q21.31
q21.33
q22
q23.1
q23.3
q24.3
q25.1
q25.3
122 bp
70,752,460 bp
70,752,480 bp
70,752,500 bp
70,752,520 bp
70,752,540 bp
70,752,560 bp
LIBD75_Illum...bam Coverage
[0 - 37]
1
2
Unphased
LIBD75_Illumina_WGS.markedTrue
G.phased.sorted.recal.localreali
gn.bam
Sequence
Gene

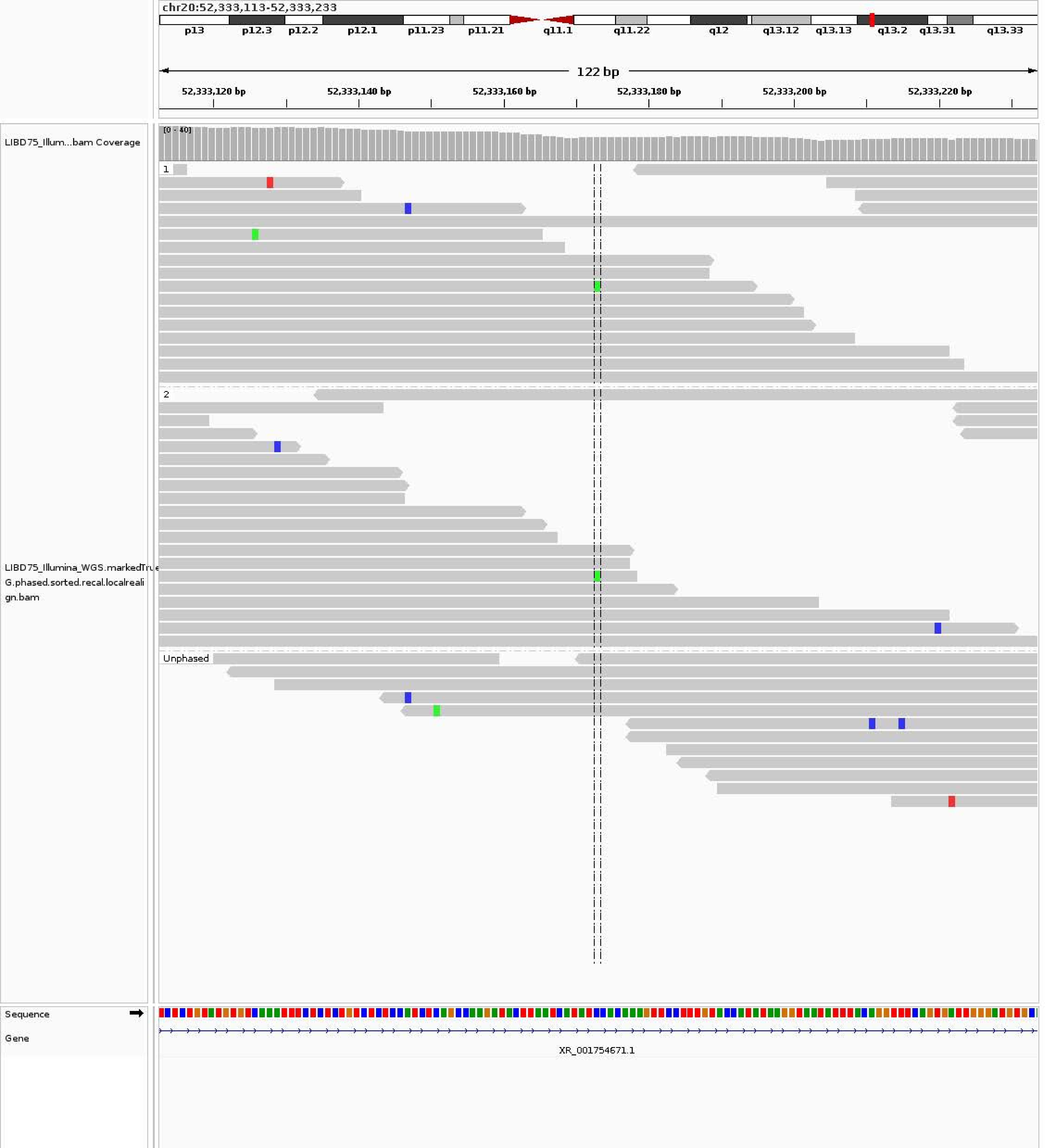
chr20:52,333,113-52,333,233
p13
p12.3
p12.2
p12.1
p11.23
p11.21
q11.1
q11.22
q12
q13.12
q13.13
q13.2
q13.31
q13.33
122 bp
52,333,120 bp
52,333,140 bp
52,333,160 bp
52,333,180 bp
52,333,200 bp
52,333,220 bp
LIBD75_Illum...bam Coverage
[0 - 40]
1
2
Unphased
LIBD75_Illumina_WGS.markedTrue
G.phased.sorted.recal.localreali
gn.bam
Sequence
Gene
XR_001754671.1
