## Supplementary material for "A personalized multi-platform assessment of somatic mosaicism in the human frontal cortex": Supplementary Table 7.pdf

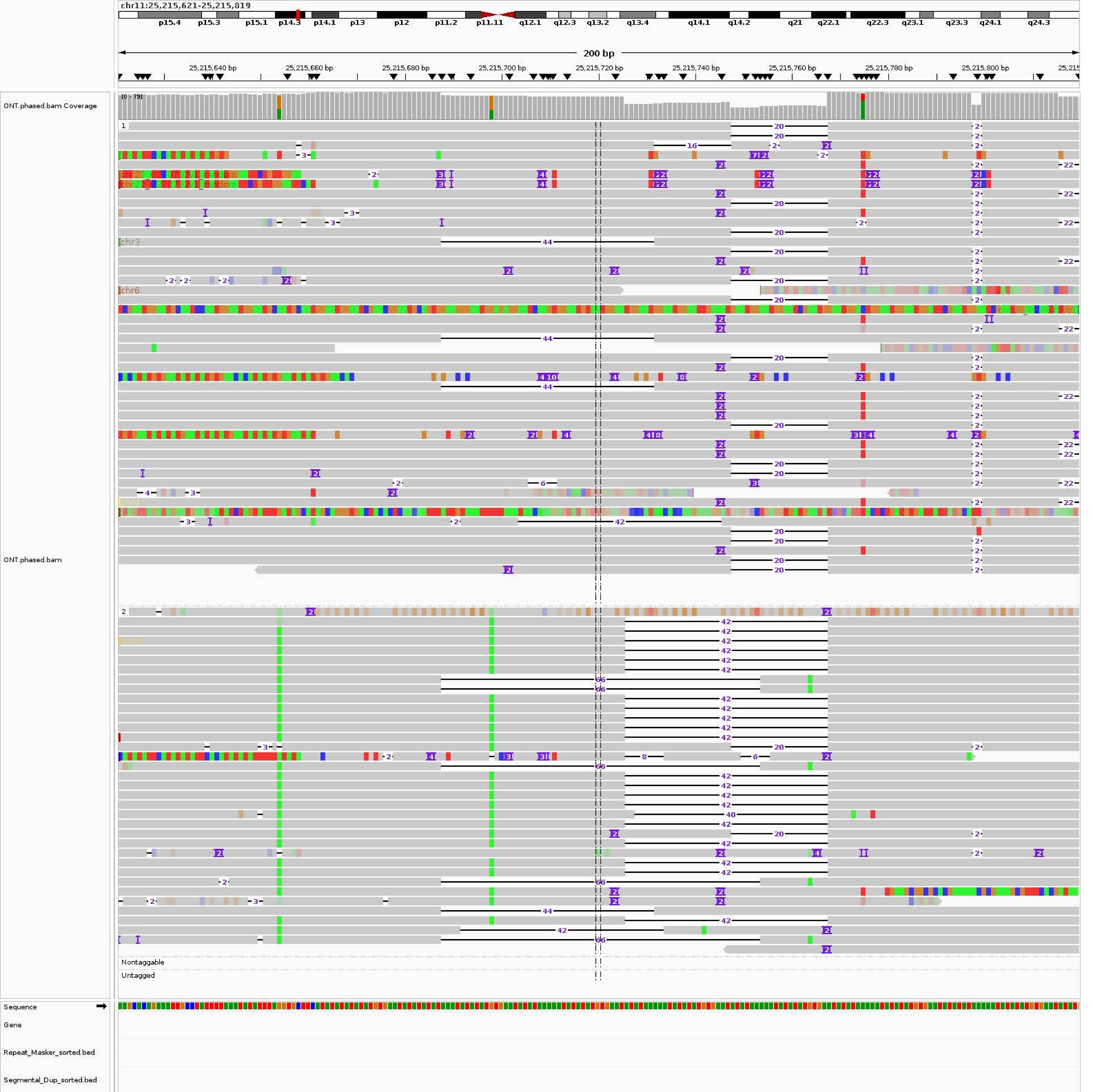
chr11:25,215,621-25,215,819
p15.4 p15.3 p15.1 p14.3 p14.1 p13 p12 p11.2 p11.11 q12.1 q12.3 q13.2 q13.4 q14.1 q14.2 q21 q22.1 q22.3 q23.1 q23.3 q24.1 q24.3
200 bp
25,215,640 bp
25,215,660 bp
25,215,680 bp
25,215,700 bp
25,215,720 bp
25,215,740 bp
25,215,760 bp
25,215,780 bp
25,215,800 bp
ONT.phased.bam Coverage
[0 - 79]
ONT.phased.bam
chr3
chr6
Nontaggable
Untagged
Sequence
Gene
Repeat_Masker_sorted.bed
Segmental_Dup_sorted.bed

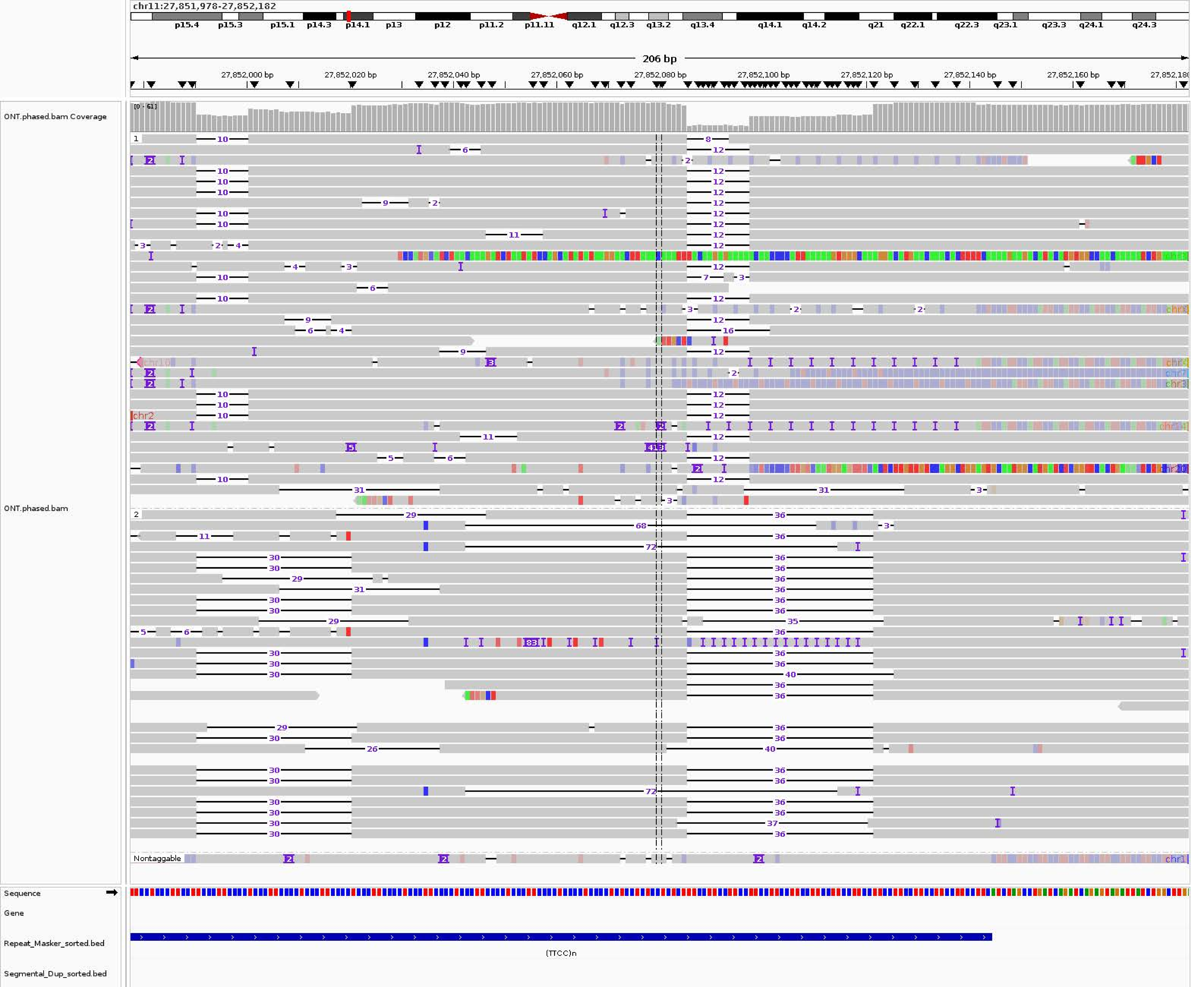
chr11:27,851,978-27,852,182
p15.4
p15.3
p15.1
p14.3
p14.1
p13
p12
p11.2
p11.11
q12.1
q12.3
q13.2
q13.4
q14.1
q14.2
q21
q22.1
q22.3
q23.1
q23.3
q24.1
q24.3
206 bp
27,852,000 bp
27,852,020 bp
27,852,040 bp
27,852,060 bp
27,852,080 bp
27,852,100 bp
27,852,120 bp
27,852,140 bp
27,852,160 bp
ONT.phased.bam Coverage
[0 - 61]
ONT.phased.bam
Nontaggable
Sequence
Gene
Repeat_Masker_sorted.bed
(TTCC)n
Segmental_Dup_sorted.bed

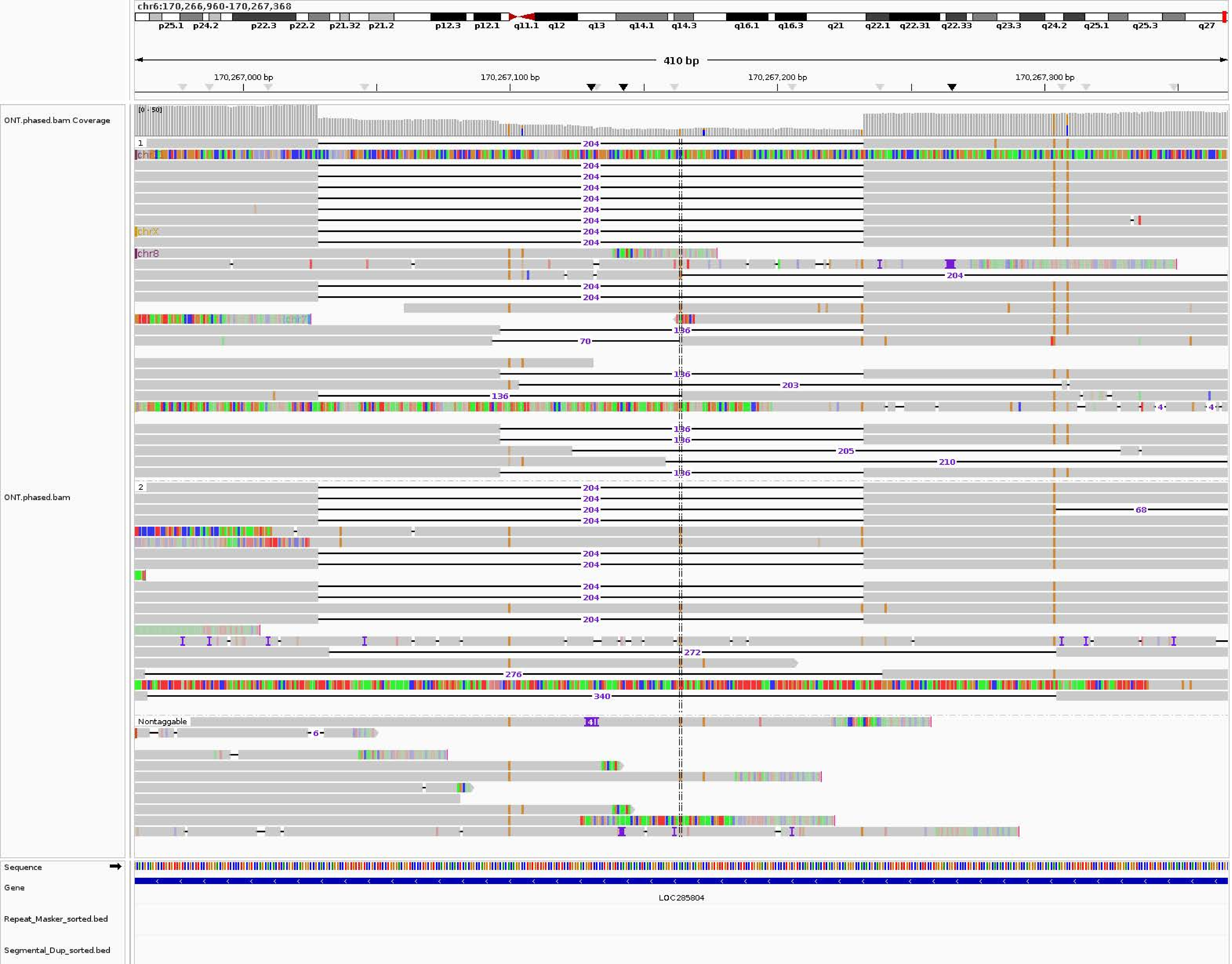

chr6:170,266,960-170,267,368
p25.1
p24.2
p22.3
p22.2
p21.32
p21.2
p12.3
p12.1
q11.1
q12
q13
q14.1
q14.3
q16.1
q16.3
q21
q22.1
q22.31
q22.33
q23.3
q24.2
q25.1
q25.3
q27
410 bp
170,267,000 bp
170,267,100 bp
170,267,200 bp
170,267,300 bp
ONT.phased.bam Coverage
[0 - 50]
ONT.phased.bam
1
2
Nontaggable
chr8
chrX
chr7
Sequence
Gene
LOC285804
Repeat_Masker_sorted.bed
Segmental_Dup_sorted.bed

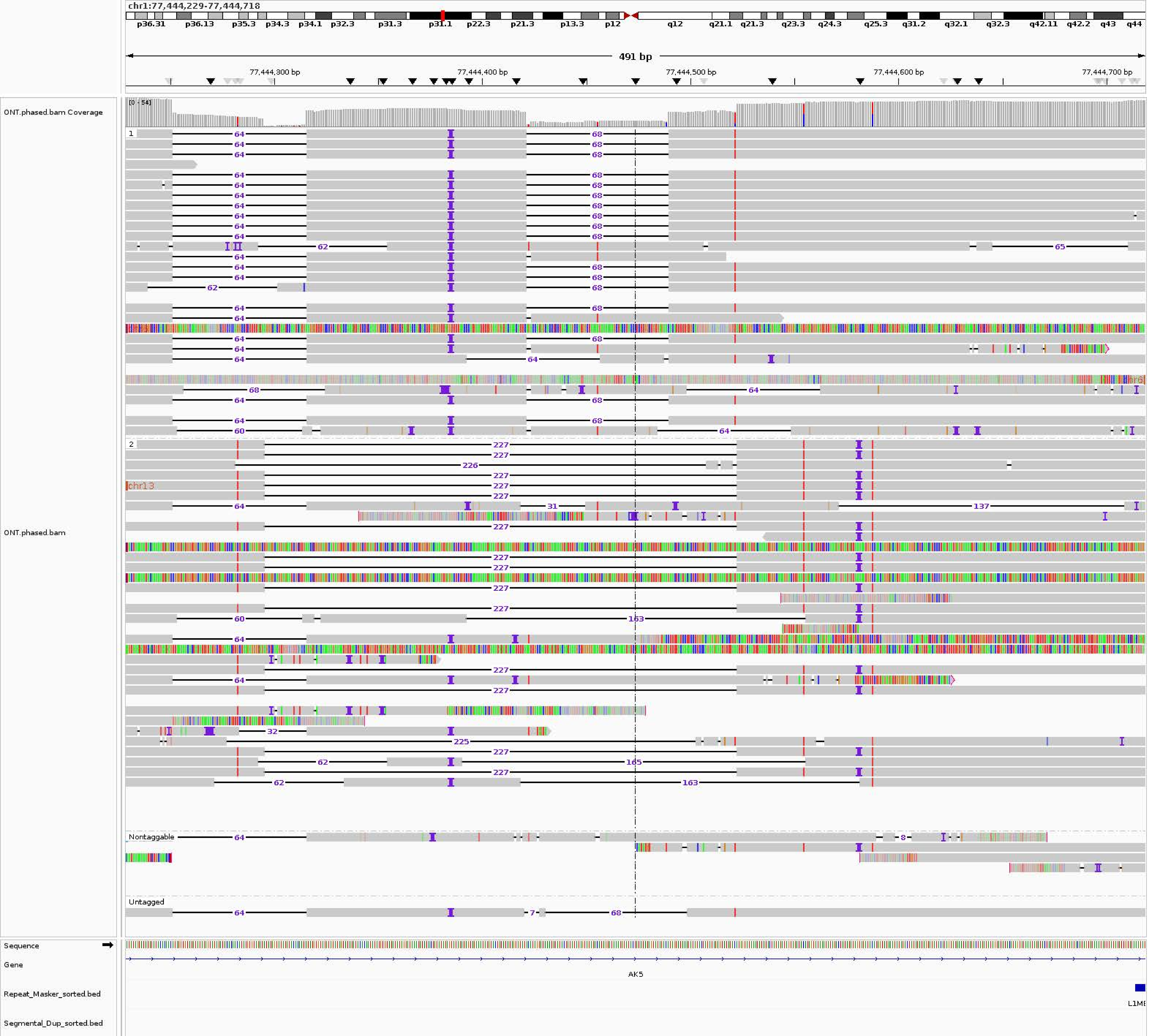
chr1:77,444,229-77,444,718
p36.31 p36.13 p35.3 p34.3 p34.1 p32.3 p31.3 p31.1 p22.3 p21.3 p13.3 p12 q12 q21.1 q21.3 q23.3 q24.3 q25.3 q31.2 q32.1 q32.3 q42.11 q42.2 q43 q44
491 bp
77,444,300 bp
77,444,400 bp
77,444,500 bp
77,444,600 bp
77,444,700 bp
ONT.phased.bam Coverage
[0 - 54]
ONT.phased.bam
1
2
chr13
Nontaggable
Untagged
Sequence
Gene
AK5
Repeat_Masker_sorted.bed
Segmental_Dup_sorted.bed

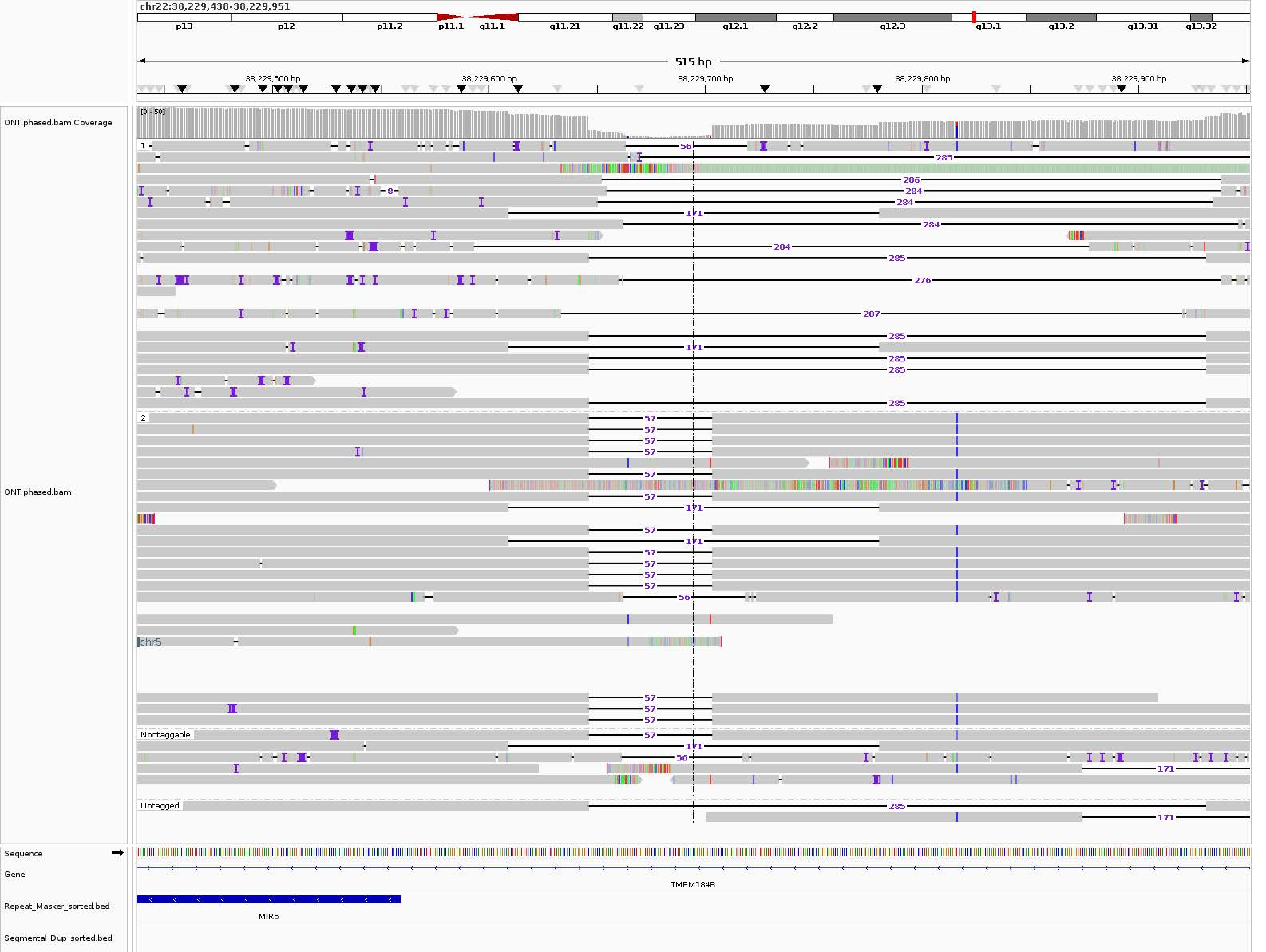

chr22:38,229,438-38,229,951
p13
p12
p11.2
p11.1
q11.1
q11.21
q11.22
q11.23
q12.1
q12.2
q12.3
q13.1
q13.2
q13.31
q13.32
515 bp
38,229,500 bp
38,229,600 bp
38,229,700 bp
38,229,800 bp
38,229,900 bp
ONT.phased.bam Coverage
[0 - 50]
ONT.phased.bam
1
2
chr5
Nontaggable
Untagged
Sequence
Gene
TMEM184B
Repeat_Masker_sorted.bed
MIRb
Segmental_Dup_sorted.bed

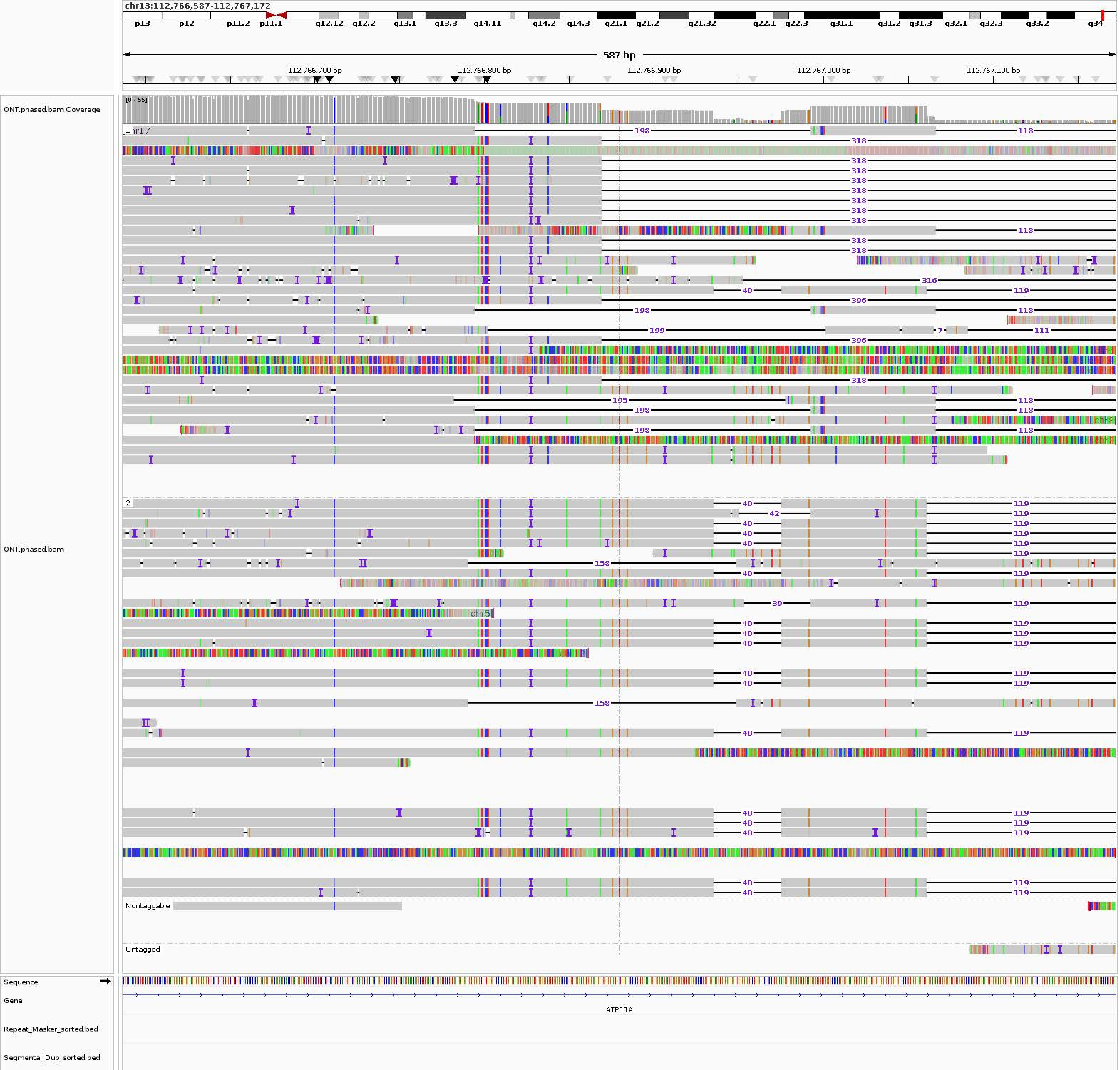
chr13:112,766,587-112,767,172
p13
p12
p11.2
p11.1
q12.12
q12.2
q13.1
q13.3
q14.11
q14.2
q14.3
q21.1
q21.2
q21.32
q22.1
q22.3
q31.1
q31.2
q31.3
q32.1
q32.3
q33.2
q34
587 bp
112,766,700 bp
112,766,800 bp
112,766,900 bp
112,767,000 bp
112,767,100 bp
ONT.phased.bam Coverage
[0 - 55]
ONT.phased.bam
1
2
Nontaggable
Untagged
Sequence
Gene
ATP11A
Repeat_Masker_sorted.bed
Segmental_Dup_sorted.bed

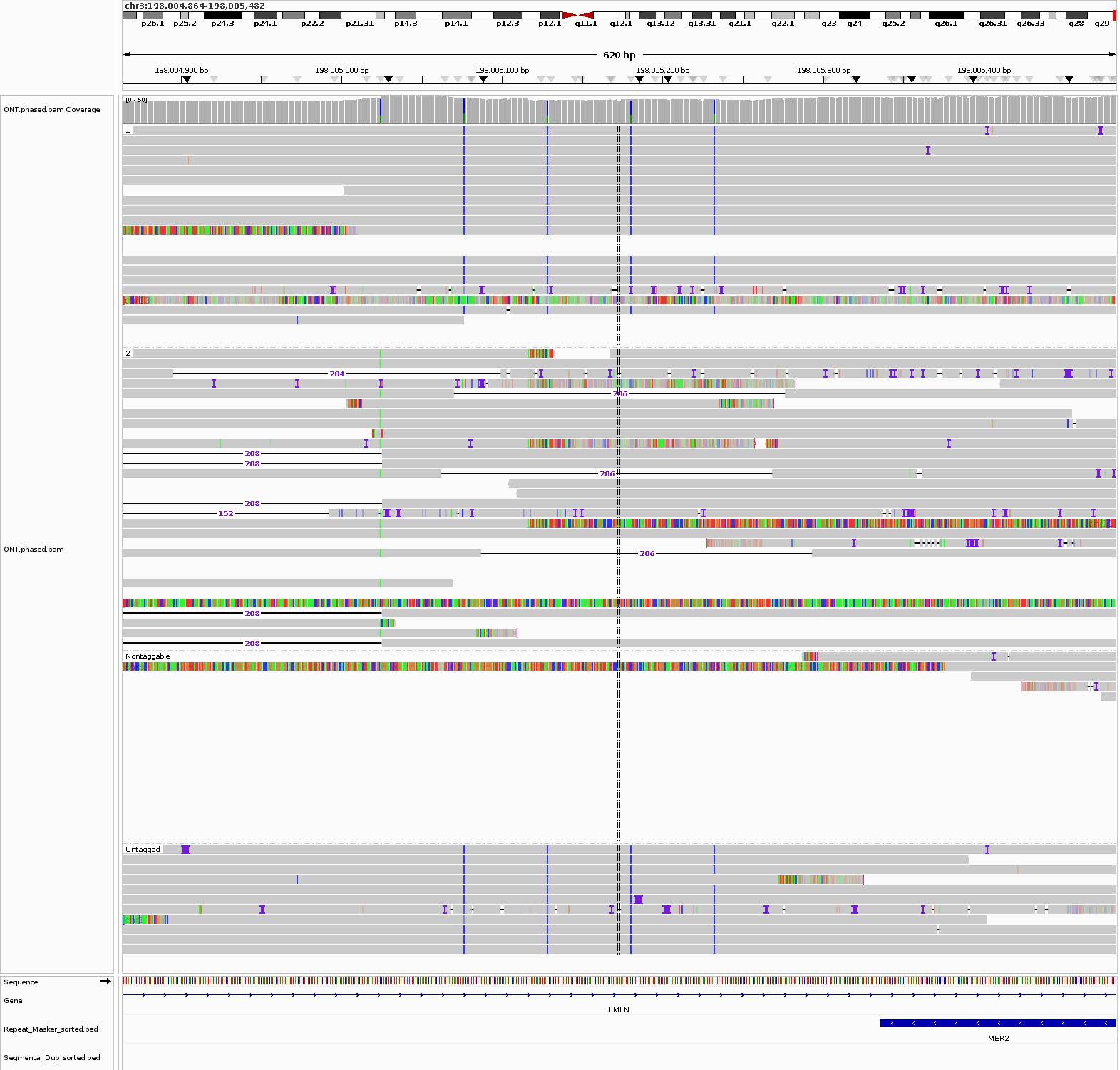
chr3:198,004,864-198,005,482
p26.1 p25.2 p24.3 p24.1 p22.2 p21.31 p14.3 p14.1 p12.3 p12.1 q11.1 q12.1 q13.12 q13.31 q21.1 q22.1 q23 q24 q25.2 q26.1 q26.31 q26.33 q28 q29
620 bp
198,004,900 bp
198,005,000 bp
198,005,100 bp
198,005,200 bp
198,005,300 bp
198,005,400 bp
ONT.phased.bam Coverage
[0 - 50]
1
2
204
206
208
152
Nontaggable
Untagged
ONT.phased.bam
Sequence
Gene
LMLN
Repeat_Masker_sorted.bed
MER2
Segmental_Dup_sorted.bed

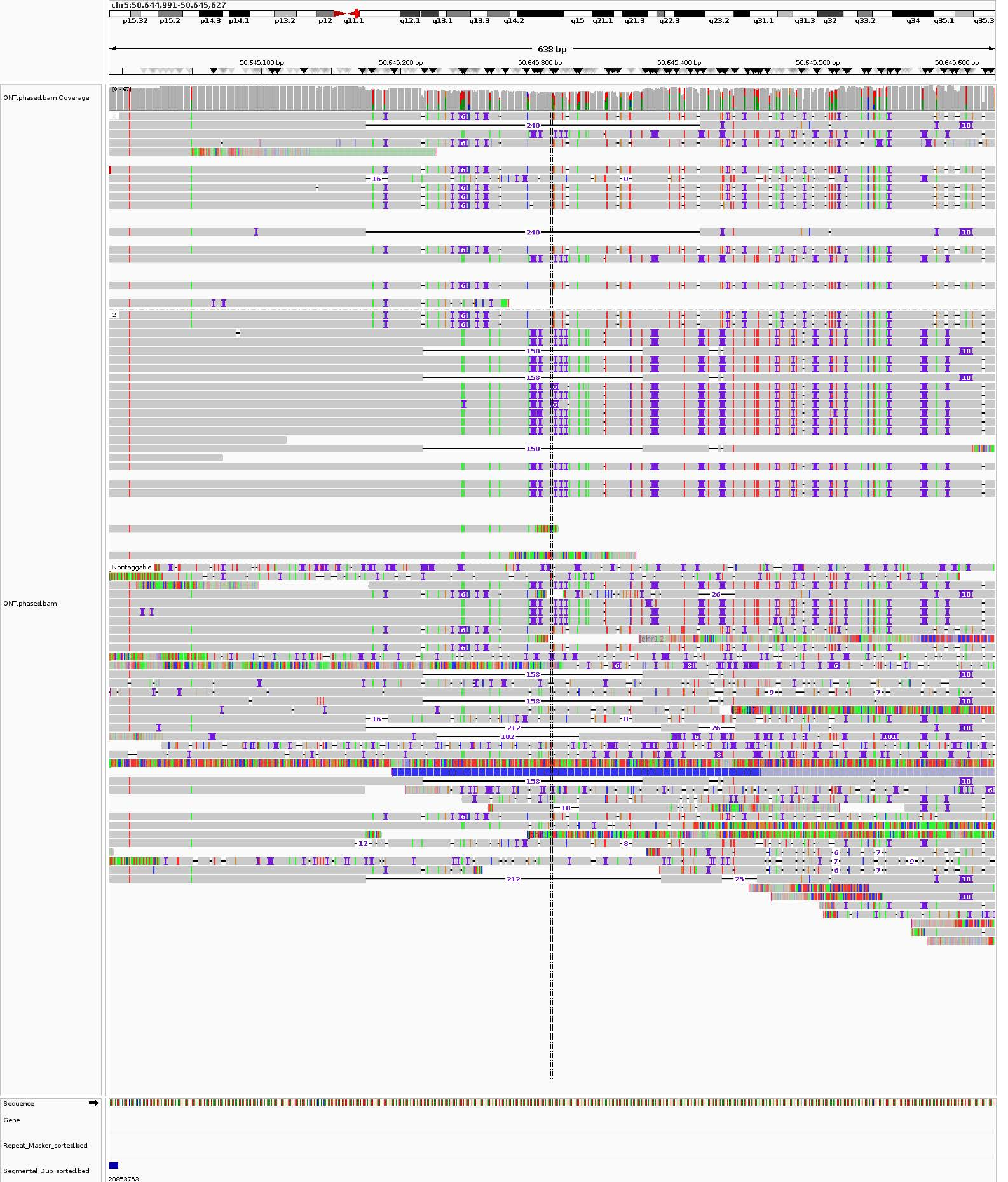
chr5:50,644,991-50,645,627
p15.32
p15.2
p14.3
p14.1
p13.2
p12
q11.1
q12.1
q13.1
q13.3
q14.2
q15
q21.1
q21.3
q22.3
q23.2
q31.1
q31.3
q32
q33.2
q34
q35.1
q35.3
638 bp
50,645,100 bp
50,645,200 bp
50,645,300 bp
50,645,400 bp
50,645,500 bp
50,645,600 bp
ONT.phased.bam Coverage
[0 - 67]
ONT.phased.bam
1
2
Nontaggable
240
158
212
102
chr12
Sequence
Gene
Repeat_Masker_sorted.bed
Segmental_Dup_sorted.bed
20853753

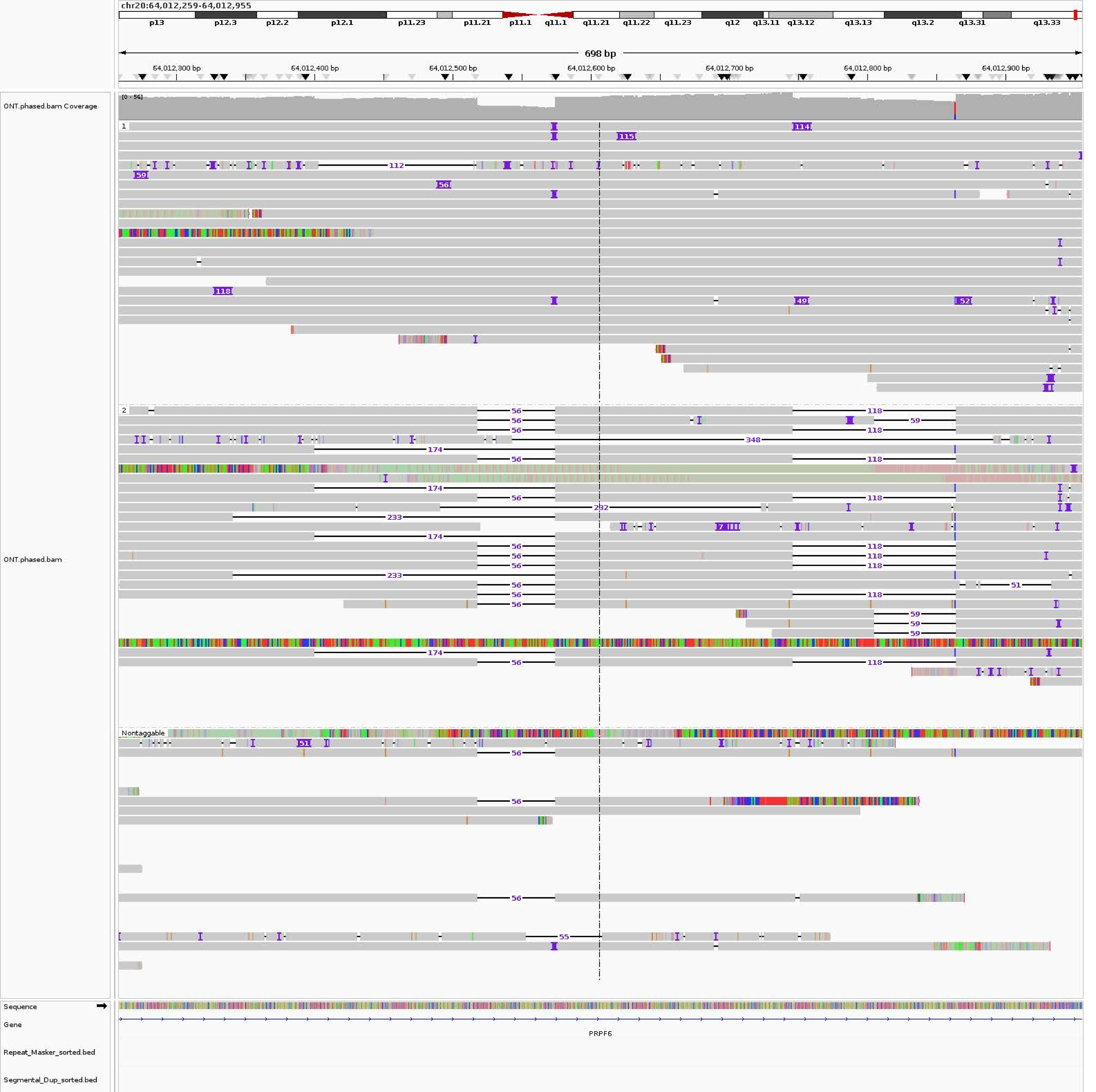
chr20:64,012,259-64,012,955
p13
p12.3
p12.2
p12.1
p11.23
p11.21
p11.1
q11.1
q11.21
q11.22
q11.23
q12
q13.11
q13.12
q13.13
q13.2
q13.31
q13.33
698 bp
64,012,300 bp
64,012,400 bp
64,012,500 bp
64,012,600 bp
64,012,700 bp
64,012,800 bp
64,012,900 bp
ONT.phased.bam Coverage
[0 - 56]
ONT.phased.bam
1
2
Nontaggable
Sequence
Gene
PRPF6
Repeat_Masker_sorted.bed
Segmental_Dup_sorted.bed

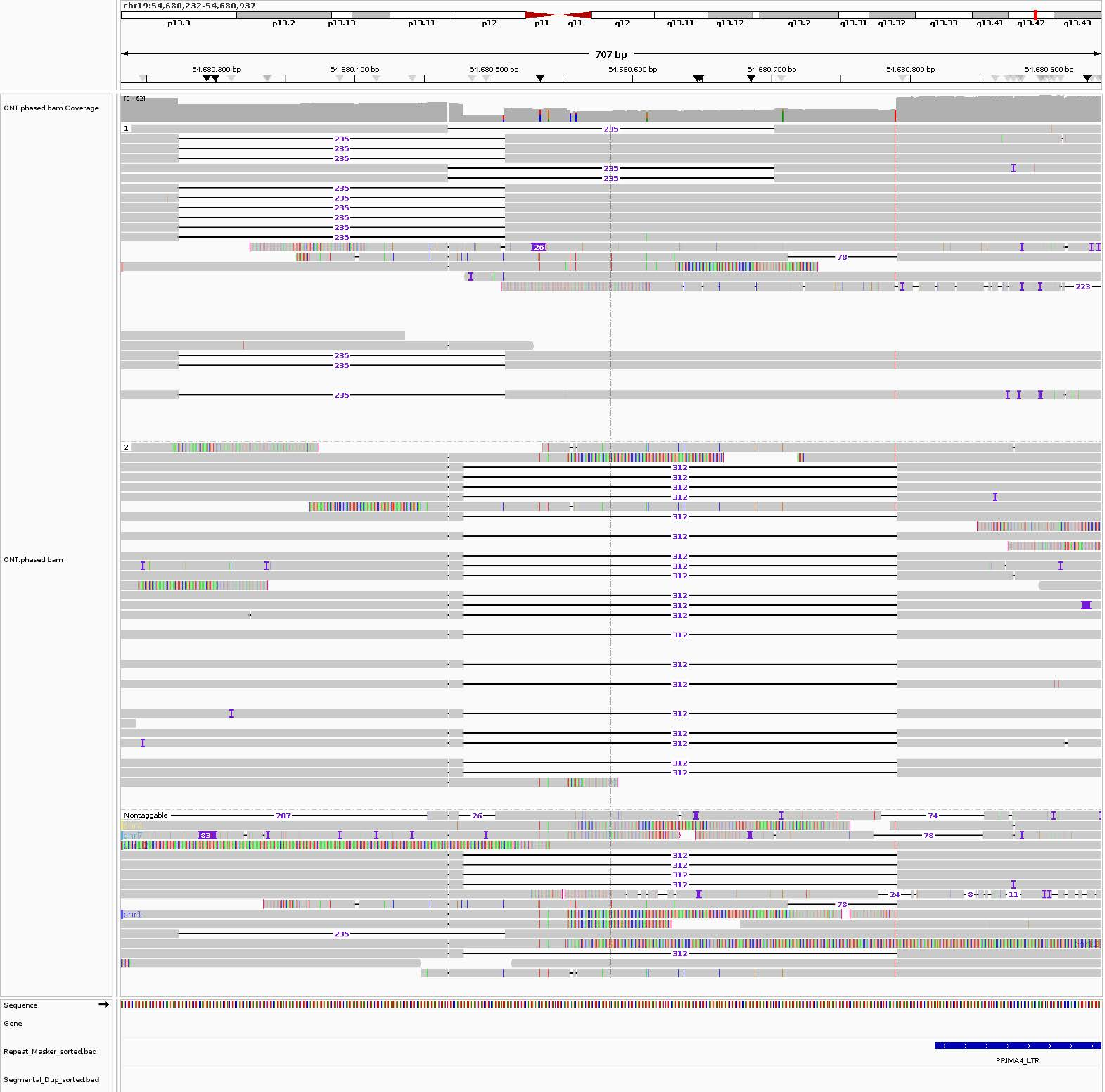

chr19:54,680,232-54,680,937
p13.3
p13.2
p13.13
p13.11
p12
p11
q11
q12
q13.11
q13.12
q13.2
q13.31
q13.32
q13.33
q13.41
q13.42
q13.43
707 bp
54,680,300 bp
54,680,400 bp
54,680,500 bp
54,680,600 bp
54,680,700 bp
54,680,800 bp
54,680,900 bp
ONT.phased.bam Coverage
[0 - 62]
ONT.phased.bam
1
2
Nontaggable
235
78
223
26
312
207
74
83
24
8
11
chr1
chr7
Sequence
Gene
Repeat_Masker_sorted.bed
PRIMA4_LTR
Segmental_Dup_sorted.bed

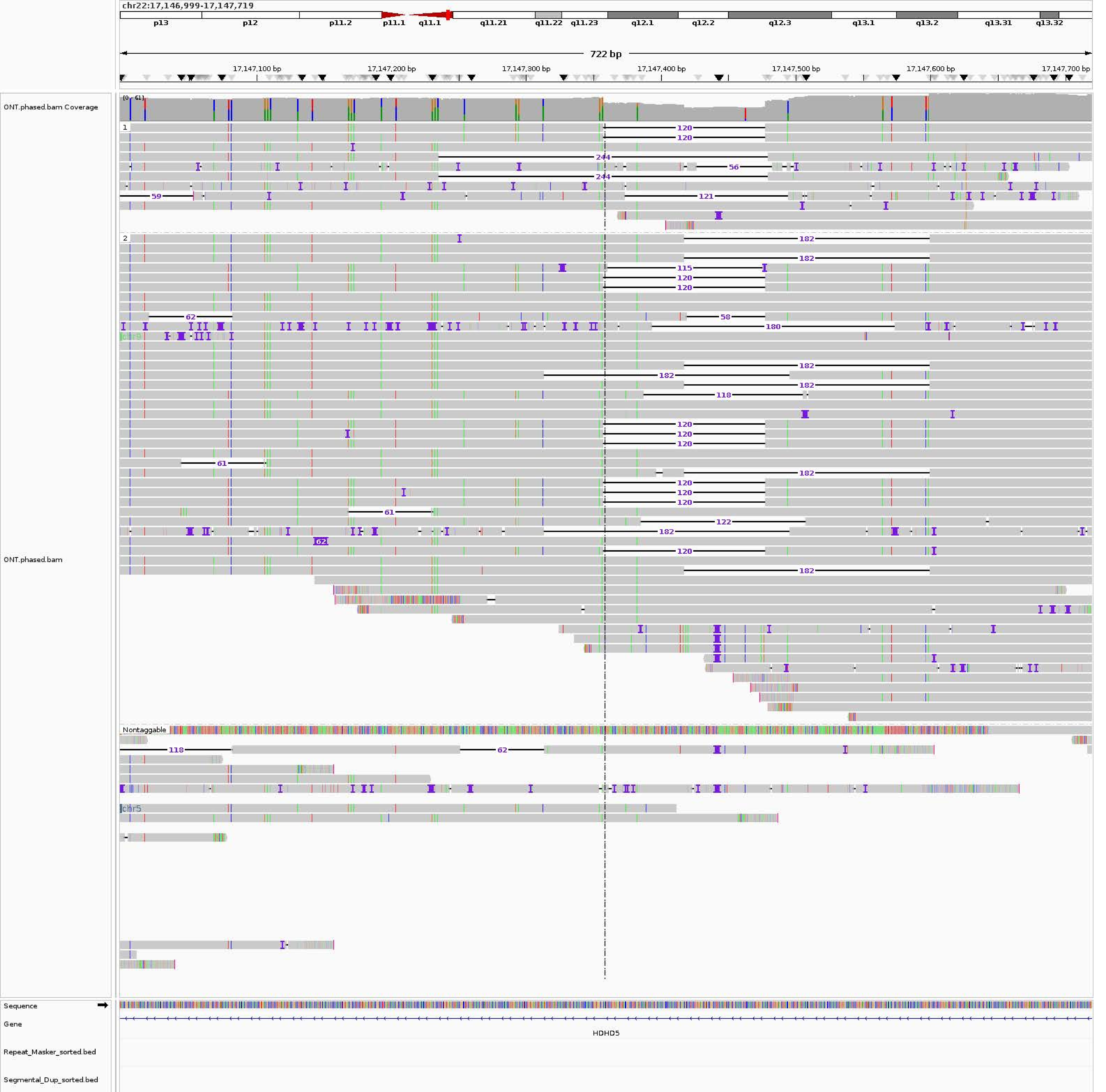
chr22:17,146,999-17,147,719
p13
p12
p11.2
p11.1
q11.1
q11.21
q11.22
q11.23
q12.1
q12.2
q12.3
q13.1
q13.2
q13.31
q13.32
722 bp
17,147,100 bp
17,147,200 bp
17,147,300 bp
17,147,400 bp
17,147,500 bp
17,147,600 bp
17,147,700 bp
ONT.phased.bam Coverage
[0 - 61]
ONT.phased.bam
Nontaggable
chr9
chr5
Sequence
Gene
HDHD5
Repeat_Masker_sorted.bed
Segmental_Dup_sorted.bed

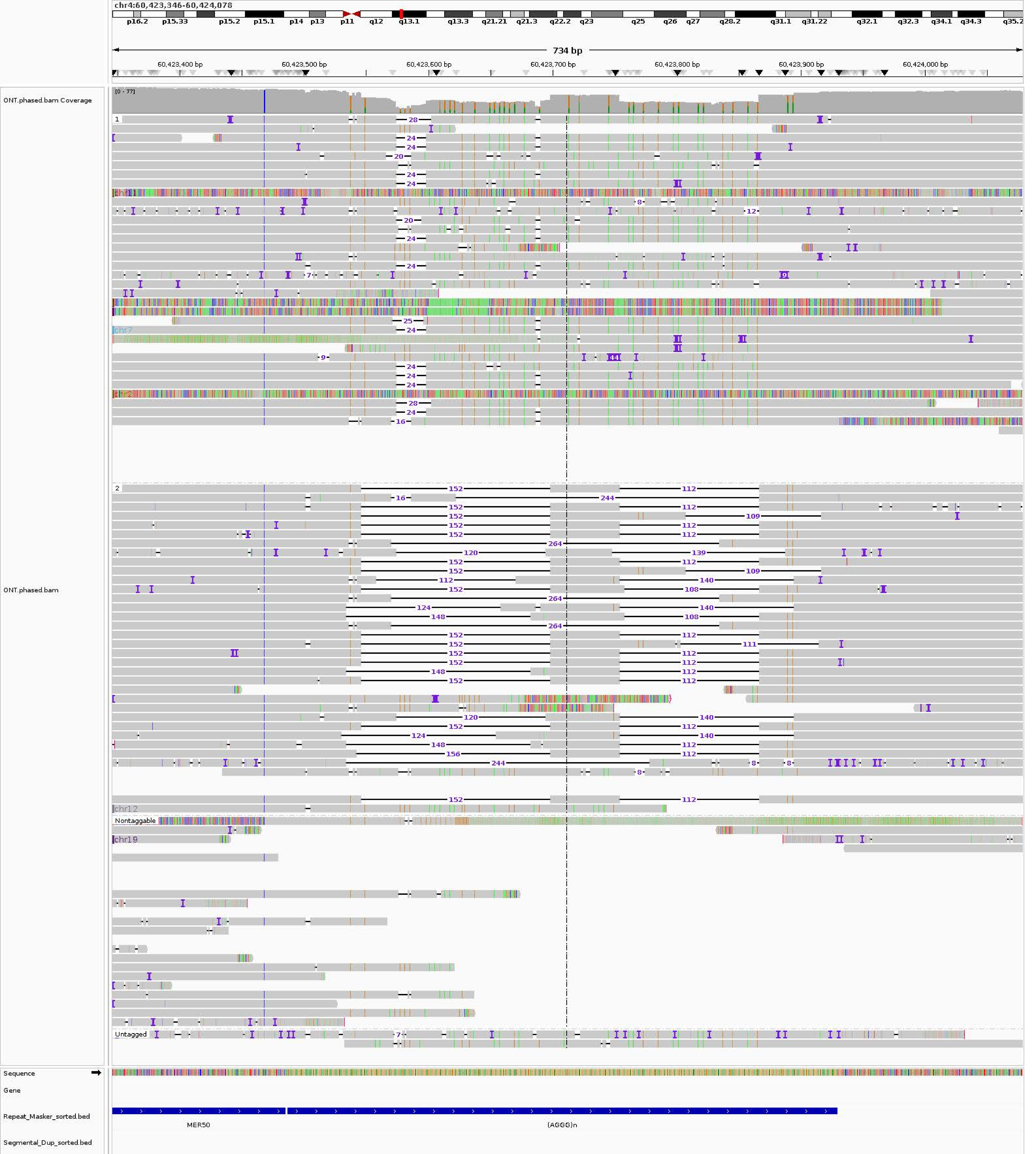
chr4:60,423,346-60,424,078
p16.2 p15.33 p15.2 p15.1 p14 p13 p11 q12 q13.1 q13.3 q21.21 q21.3 q22.2 q23 q25 q26 q27 q28.2 q31.1 q31.22 q32.1 q32.3 q34.1 q34.3 q35.2
734 bp
60,423,400 bp
60,423,500 bp
60,423,600 bp
60,423,700 bp
60,423,800 bp
60,423,900 bp
60,424,000 bp
ONT.phased.bam Coverage
[0 - 77]
ONT.phased.bam
1
2
Nontaggable
Untagged
chr11
chr7
chr2
chr12
chr19
Sequence
Gene
Repeat_Masker_sorted.bed
Segmental_Dup_sorted.bed
MER50
(AGGG)n

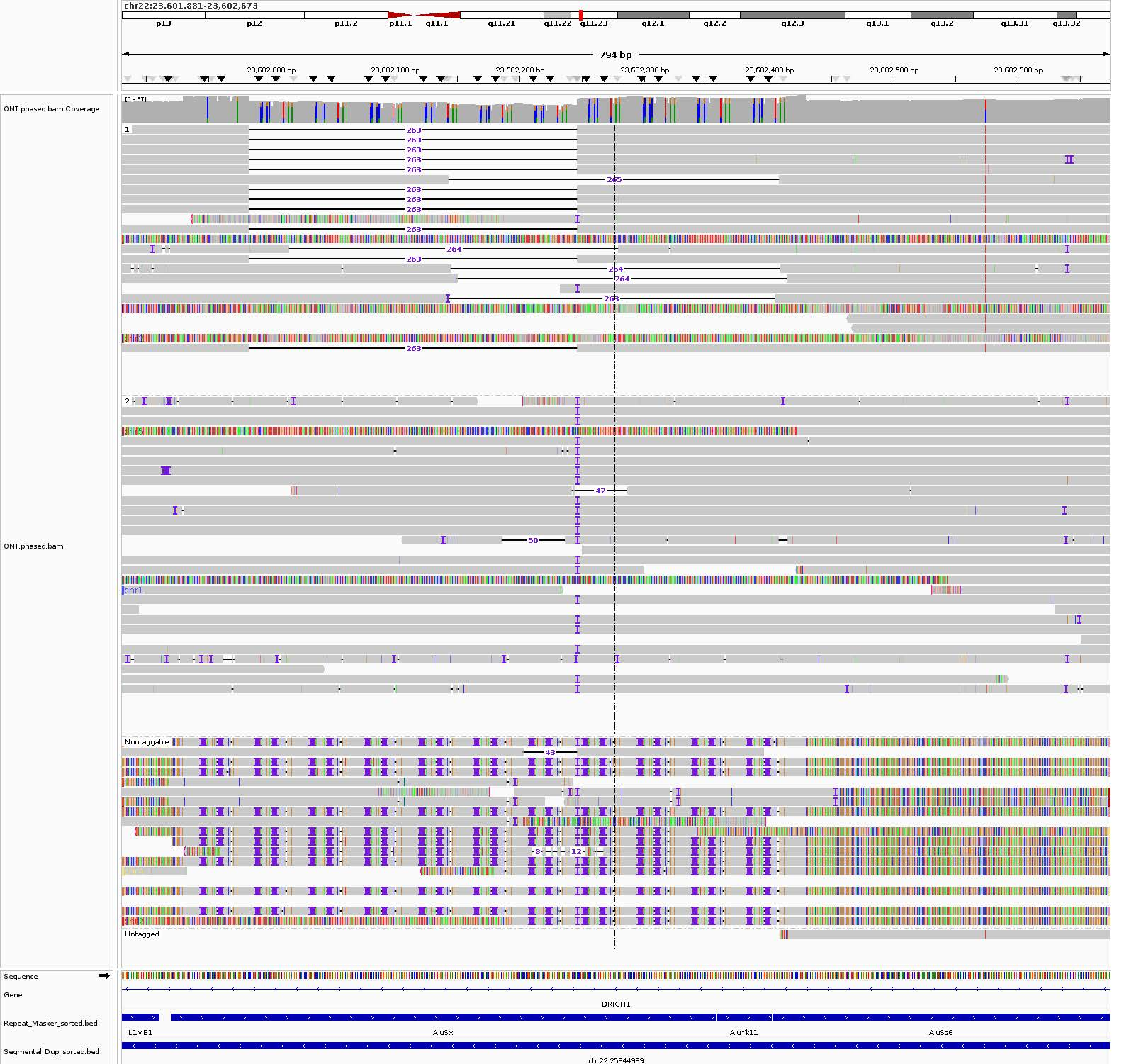
chr22:23,601,881-23,602,673
p13
p12
p11.2
p11.1
q11.1
q11.21
q11.22
q11.23
q12.1
q12.2
q12.3
q13.1
q13.2
q13.31
q13.32
794 bp
23,602,000 bp
23,602,100 bp
23,602,200 bp
23,602,300 bp
23,602,400 bp
23,602,500 bp
23,602,600 bp
ONT.phased.bam Coverage
[0 - 57]
ONT.phased.bam
1
2
263
265
264
42
50
43
8
12
chr1
Nontaggable
Untagged
Sequence
Gene
DRICH1
Repeat_Masker_sorted.bed
L1ME1
AluSx
AluYk11
AluSz6
Segmental_Dup_sorted.bed
chr22:25344989

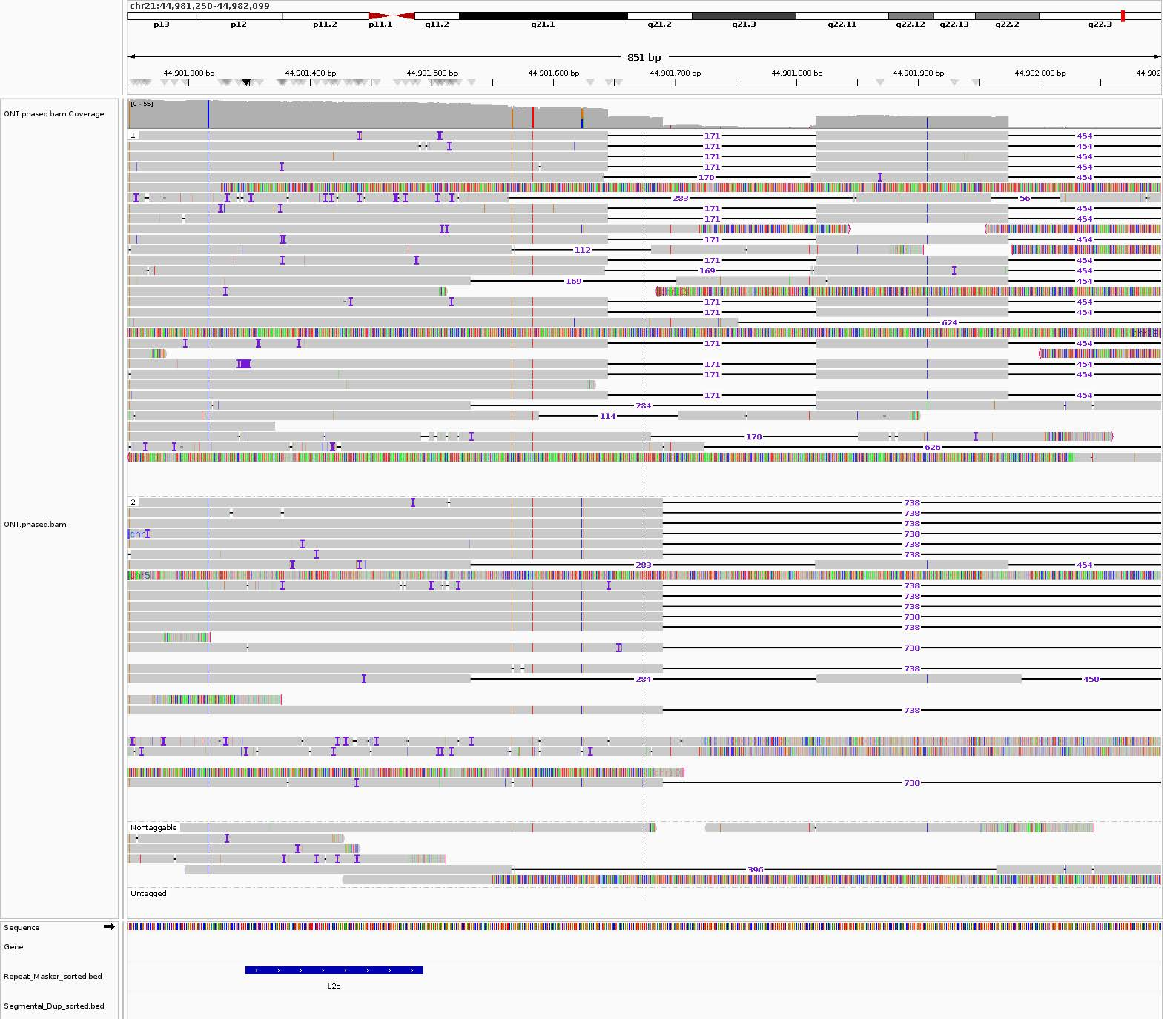

chr21:44,981,250-44,982,099
p13
p12
p11.2
p11.1
q11.2
q21.1
q21.2
q21.3
q22.11
q22.12
q22.13
q22.2
q22.3
851 bp
44,981,300 bp
44,981,400 bp
44,981,500 bp
44,981,600 bp
44,981,700 bp
44,981,800 bp
44,981,900 bp
44,982,000 bp
ONT.phased.bam Coverage
[0 - 55]
ONT.phased.bam
1
2
Nontaggable
Untagged
Sequence
Gene
Repeat_Masker_sorted.bed
L2b
Segmental_Dup_sorted.bed

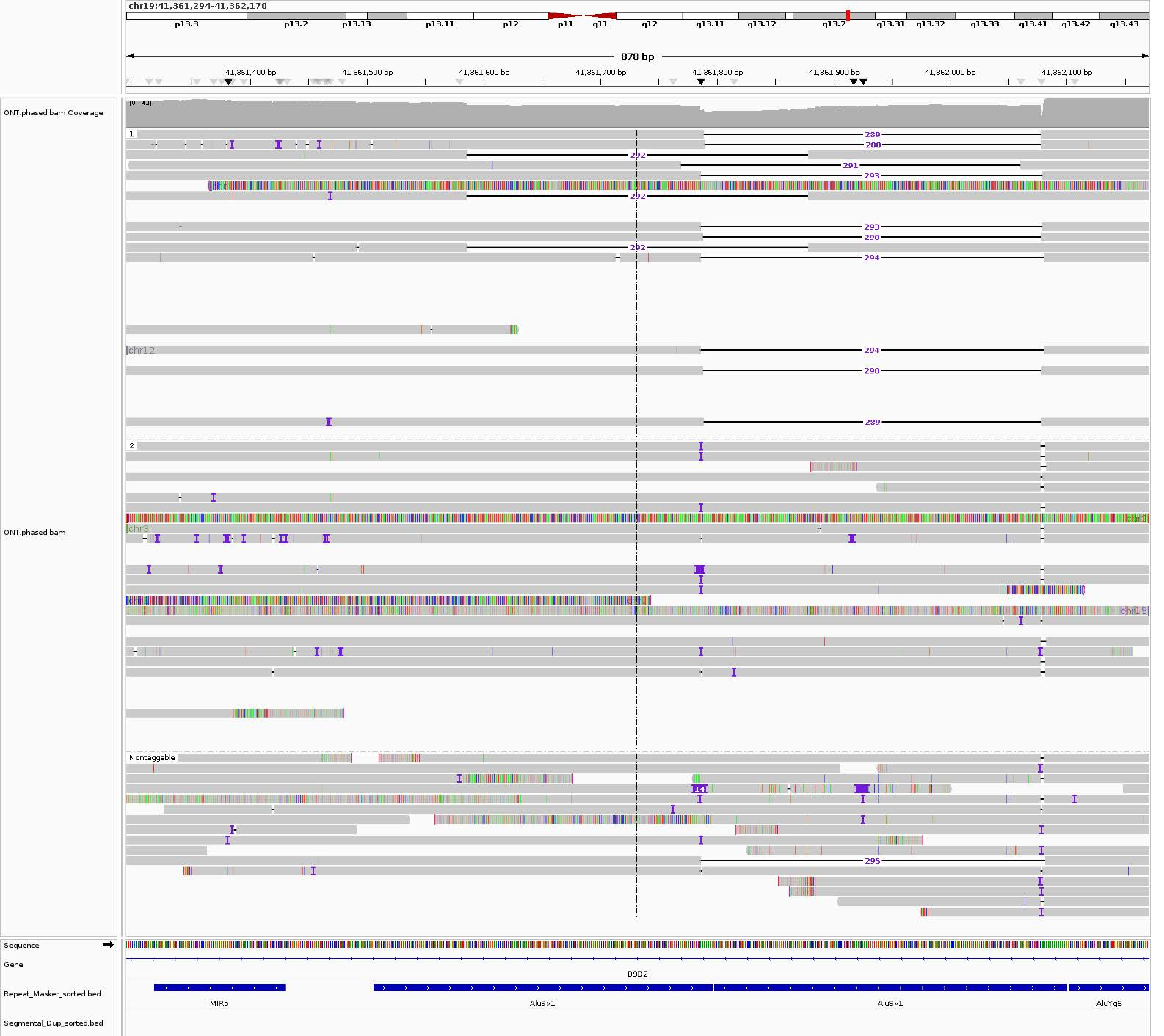

chr19:41,361,294-41,362,170
p13.3
p13.2
p13.13
p13.11
p12
p11
q11
q12
q13.11
q13.12
q13.2
q13.31
q13.32
q13.33
q13.41
q13.42
q13.43
878 bp
41,361,400 bp
41,361,500 bp
41,361,600 bp
41,361,700 bp
41,361,800 bp
41,361,900 bp
41,362,000 bp
41,362,100 bp
ONT.phased.bam Coverage
[0 - 43]
ONT.phased.bam
1
289
288
292
291
293
292
293
290
292
294
chr12
294
290
289
2
chr3
chr15
14
Nontaggable
295
Sequence
Gene
B9D2
Repeat_Masker_sorted.bed
MIRb
AluSx1
AluSx1
AluYg6
Segmental_Dup_sorted.bed

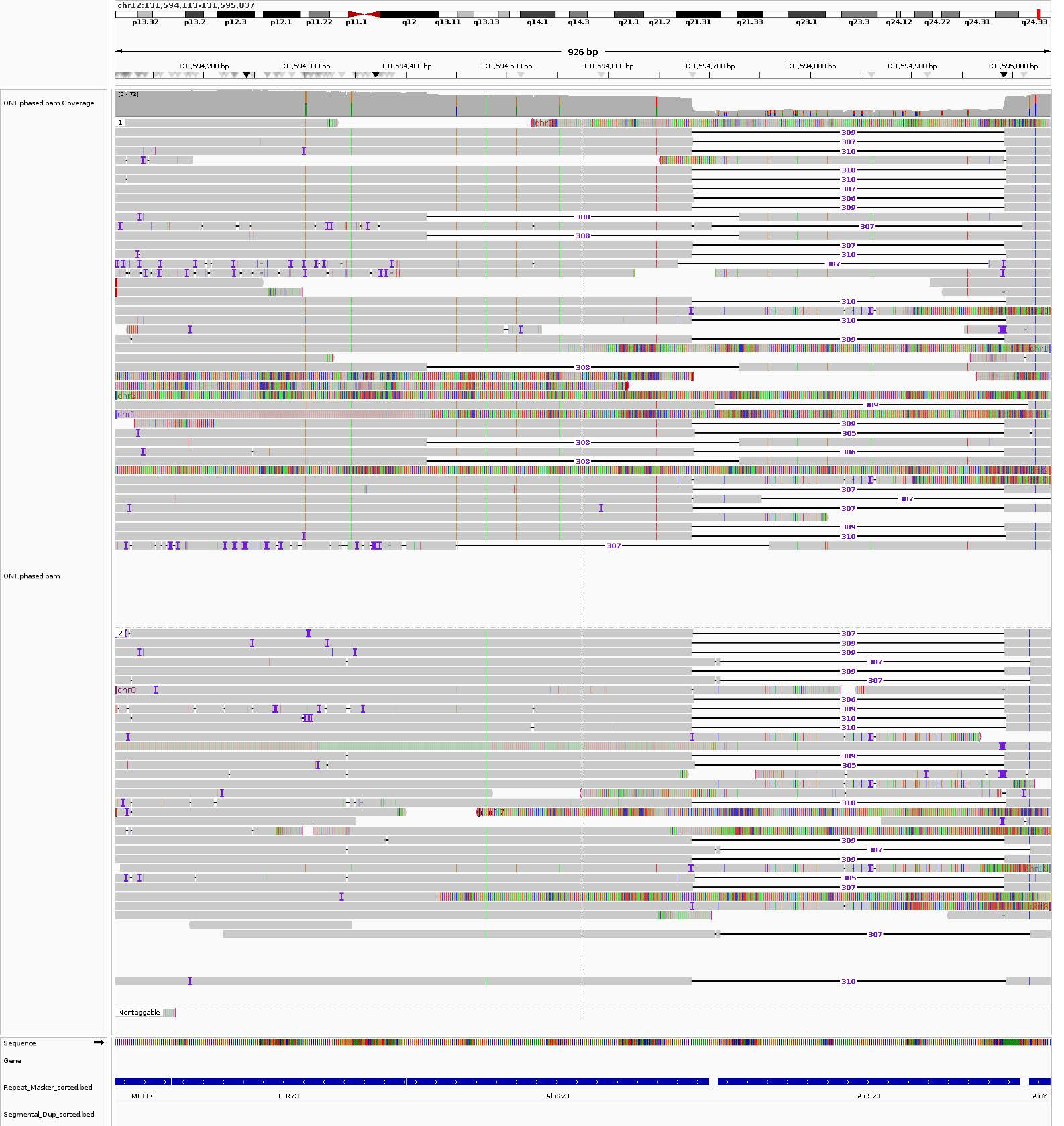
chr12:131,594,113-131,595,037
p13.32
p13.2
p12.3
p12.1
p11.22
p11.1
q12
q13.11
q13.13
q14.1
q14.3
q21.1
q21.2
q21.31
q21.33
q23.1
q23.3
q24.12
q24.22
q24.31
q24.33
926 bp
131,594,200 bp
131,594,300 bp
131,594,400 bp
131,594,500 bp
131,594,600 bp
131,594,700 bp
131,594,800 bp
131,594,900 bp
131,595,000 bp
ONT.phased.bam Coverage
[0 - 73]
ONT.phased.bam
Nontaggable
Sequence
Gene
Repeat_Masker_sorted.bed
Segmental_Dup_sorted.bed
MLT1K
LTR73
AluSx3
AluSx3
AluY

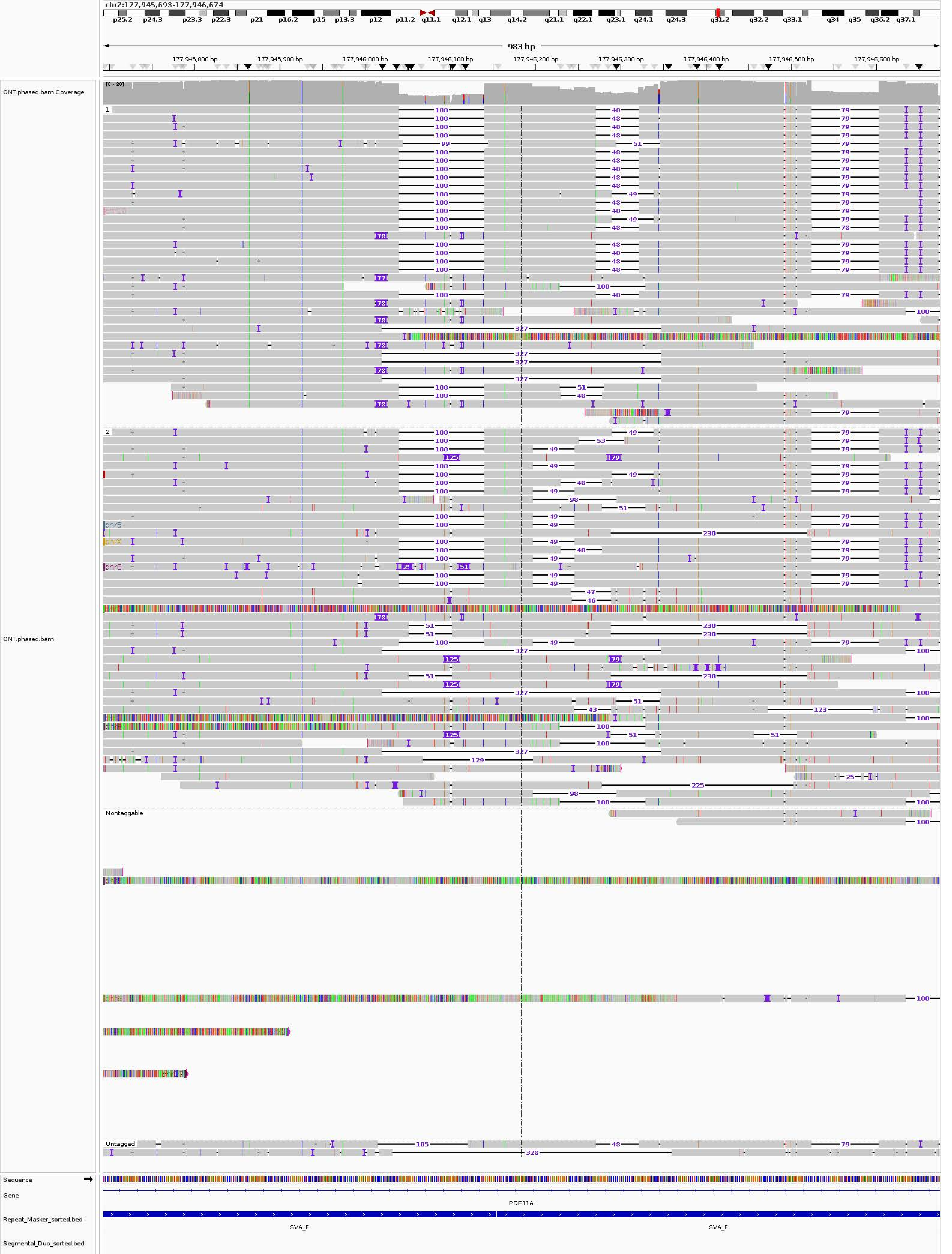
chr2:177,945,693-177,946,674
983 bp
177,945,800 bp
177,945,900 bp
177,946,000 bp
177,946,100 bp
177,946,200 bp
177,946,300 bp
177,946,400 bp
177,946,500 bp
177,946,600 bp
ONT.phased.bam Coverage
[0 - 80]
ONT.phased.bam
1
2
Nontaggable
Untagged
chr5
chrX
chr8
Sequence
Gene
PDE11A
Repeat_Masker_sorted.bed
SVA_F
SVA_F
Segmental_Dup_sorted.bed

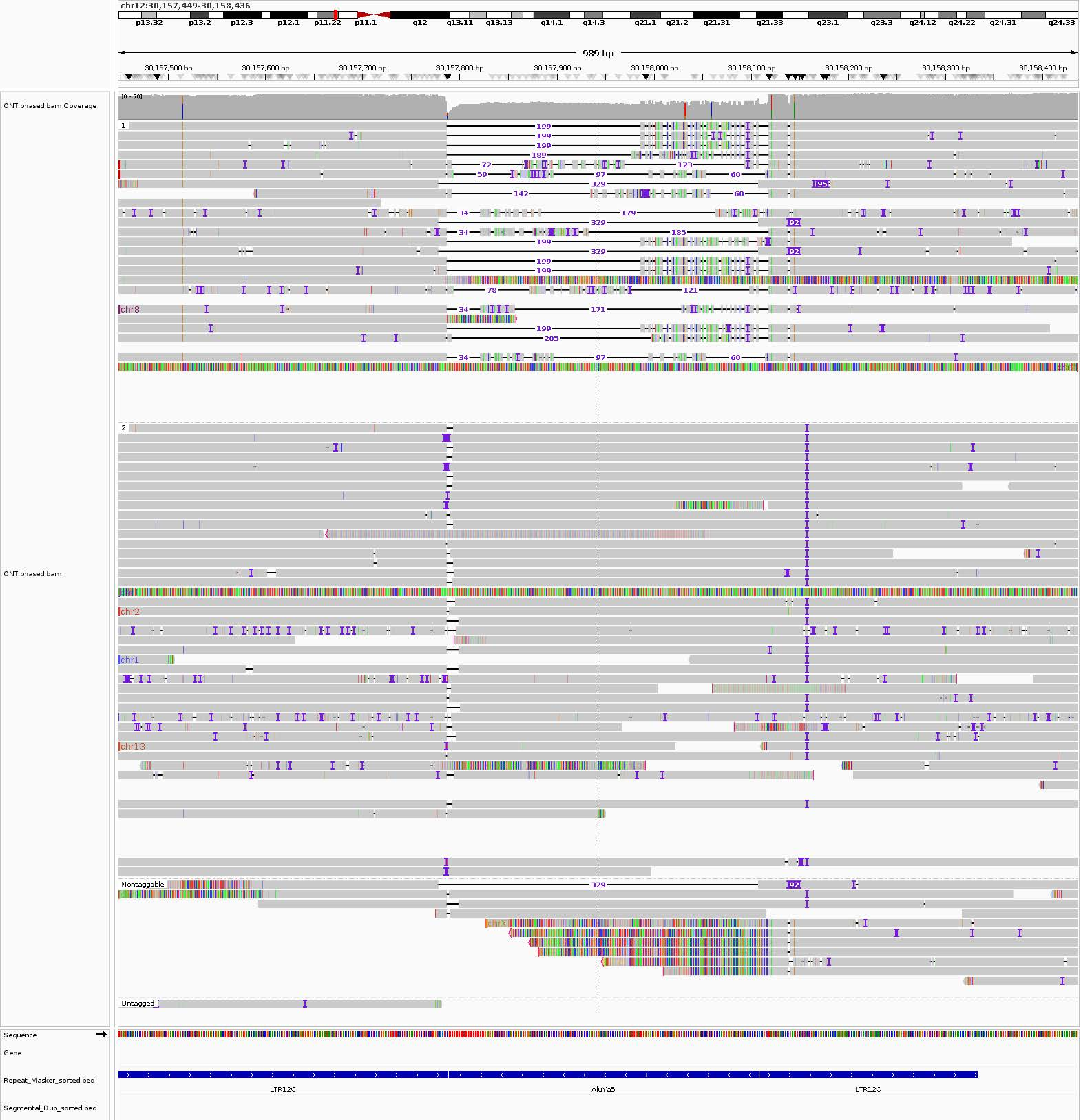

chr12:30,157,449-30,158,436
p13.32
p13.2
p12.3
p12.1
p11.22
p11.1
q12
q13.11
q13.13
q14.1
q14.3
q21.1
q21.2
q21.31
q21.33
q23.1
q23.3
q24.12
q24.22
q24.31
q24.33
989 bp
30,157,500 bp
30,157,600 bp
30,157,700 bp
30,157,800 bp
30,157,900 bp
30,158,000 bp
30,158,100 bp
30,158,200 bp
30,158,300 bp
30,158,400 bp
ONT.phased.bam Coverage
[0 - 70]
ONT.phased.bam
1
2
Nontaggable
Untagged
chr8
chr2
chr1
chr13
Sequence
Gene
Repeat_Masker_sorted.bed
Segmental_Dup_sorted.bed
LTR12C
AluYa5
LTR12C

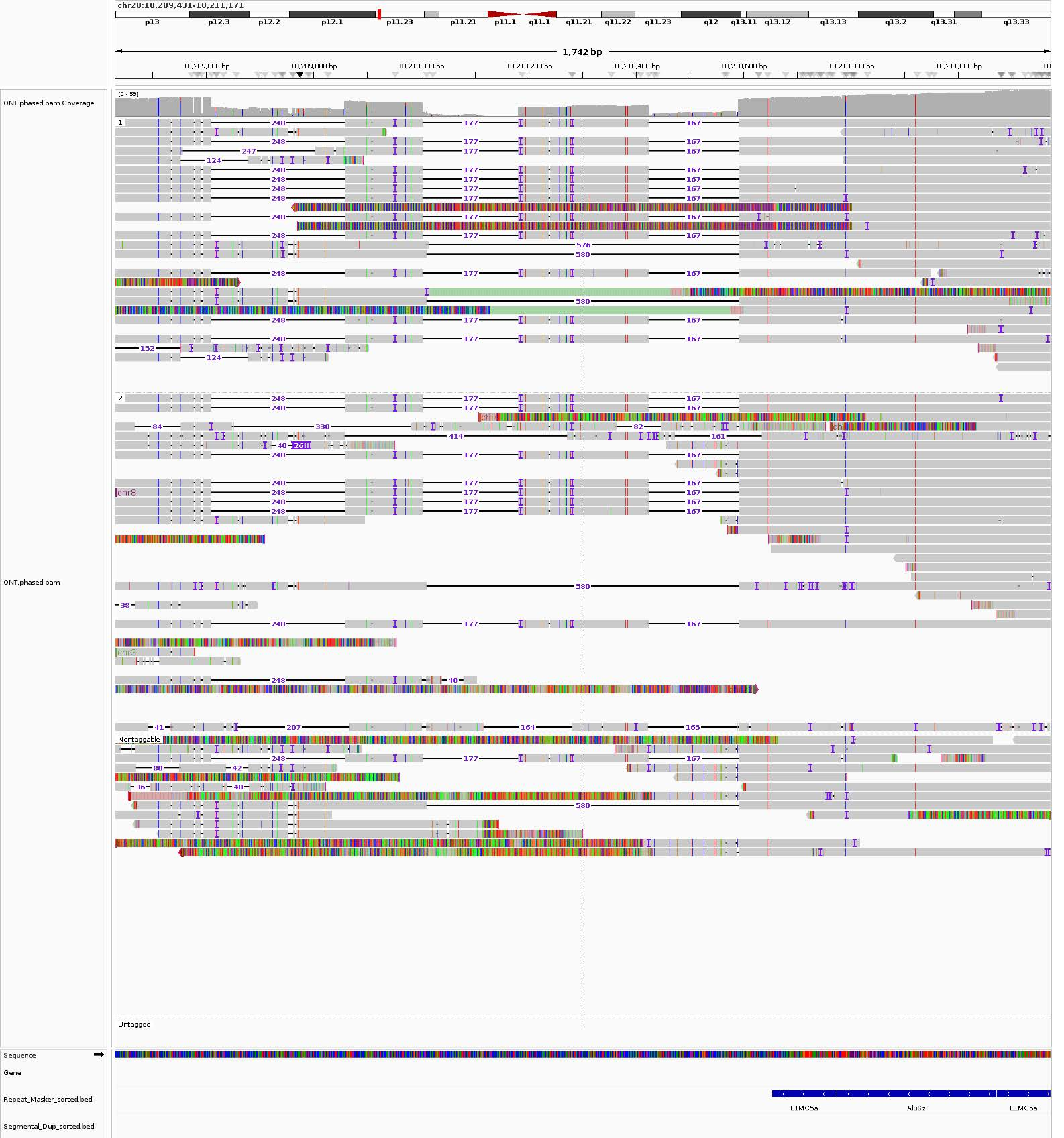
chr20:18,209,431-18,211,171
p13
p12.3
p12.2
p12.1
p11.23
p11.21
p11.1
q11.1
q11.21
q11.22
q11.23
q12
q13.11
q13.12
q13.13
q13.2
q13.31
q13.33
1,742 bp
18,209,600 bp
18,209,800 bp
18,210,000 bp
18,210,200 bp
18,210,400 bp
18,210,600 bp
18,210,800 bp
18,211,000 bp
ONT.phased.bam Coverage
[0 - 59]
ONT.phased.bam
1
2
Nontaggable
Untagged
chr8
chr3
Sequence
Gene
Repeat_Masker_sorted.bed
Segmental_Dup_sorted.bed
L1MC5a
AluSz
L1MC5a

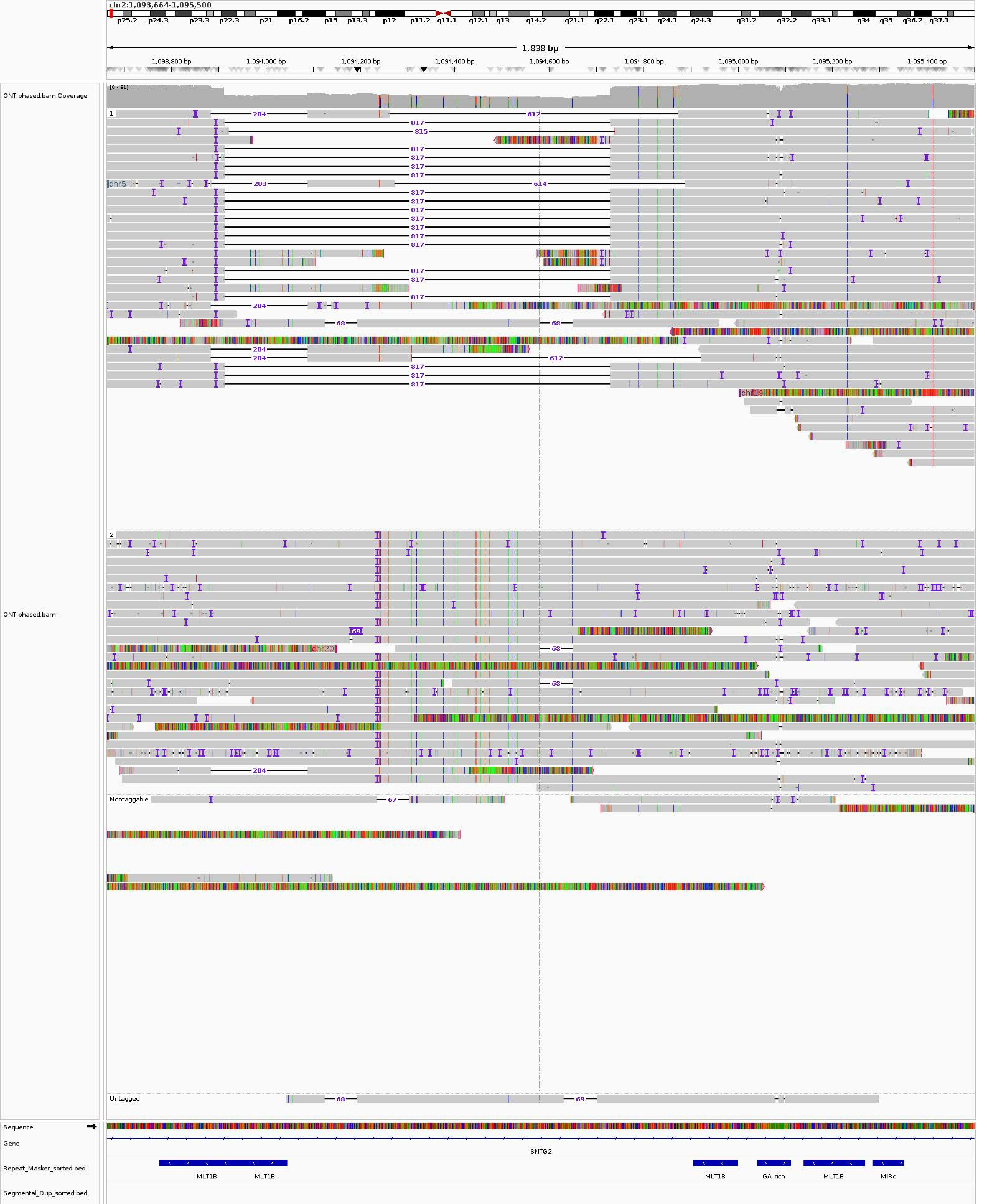

chr2:1,093,664-1,095,500
p25.2 p24.3 p23.3 p22.3 p21 p16.2 p15 p13.3 p12 p11.2 q11.1 q12.1 q13 q14.2 q21.1 q22.1 q23.1 q24.1 q24.3 q31.2 q32.2 q33.1 q34 q35 q36.2 q37.1
1,838 bp
1,093,800 bp
1,094,000 bp
1,094,200 bp
1,094,400 bp
1,094,600 bp
1,094,800 bp
1,095,000 bp
1,095,200 bp
1,095,400 bp
ONT.phased.bam Coverage
[0 - 61]
ONT.phased.bam
1
2
Nontaggable
Untagged
204
612
817
815
203
614
68
chr5
chr19
chr20
69
67
Sequence
Gene
SNTG2
Repeat_Masker_sorted.bed
MLT1B
MLT1B
MLT1B
GA-rich
MLT1B
MIRc
Segmental_Dup_sorted.bed

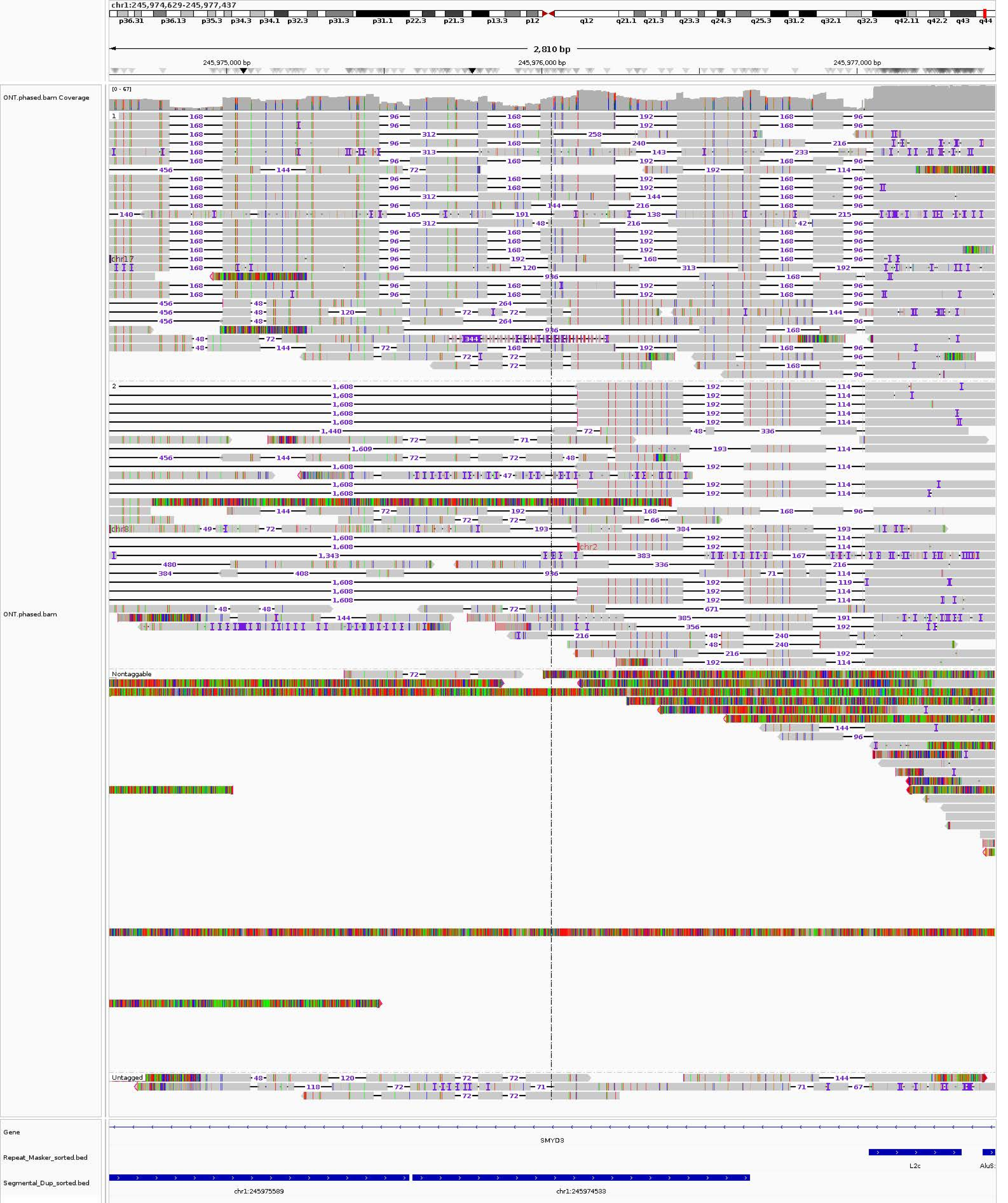

chr1:245,974,629-245,977,437
2,810 bp
245,975,000 bp
245,976,000 bp
245,977,000 bp
ONT.phased.bam Coverage
[0 - 67]
ONT.phased.bam
1
2
Nontaggable
Untagged
Gene
SMYD3
Repeat_Masker_sorted.bed
L2c
Segmental_Dup_sorted.bed
chr1:245975589
chr1:245974533

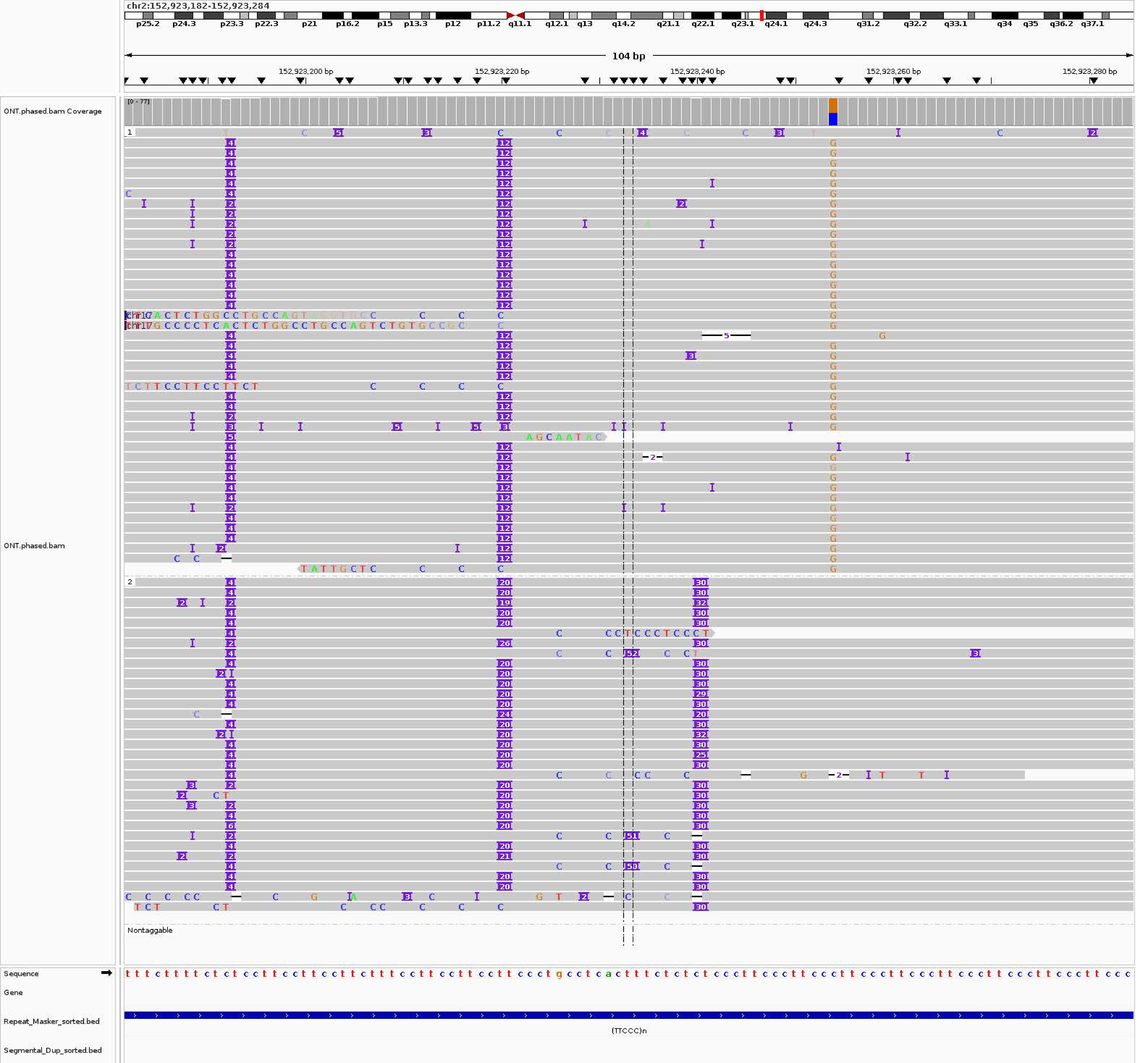
chr2:152,923,182-152,923,284
p25.2 p24.3 p23.3 p22.3 p21 p16.2 p15 p13.3 p12 p11.2 q11.1 q12.1 q13 q14.2 q21.1 q22.1 q23.1 q24.1 q24.3 q31.2 q32.2 q33.1 q34 q35 q36.2 q37.1
104 bp
152,923,200 bp
152,923,220 bp
152,923,240 bp
152,923,260 bp
152,923,280 bp
ONT.phased.bam Coverage
[0 - 77]
ONT.phased.bam
1
2
Nontaggable
Sequence
Gene
Repeat_Masker_sorted.bed
(TTCCC)n
Segmental_Dup_sorted.bed

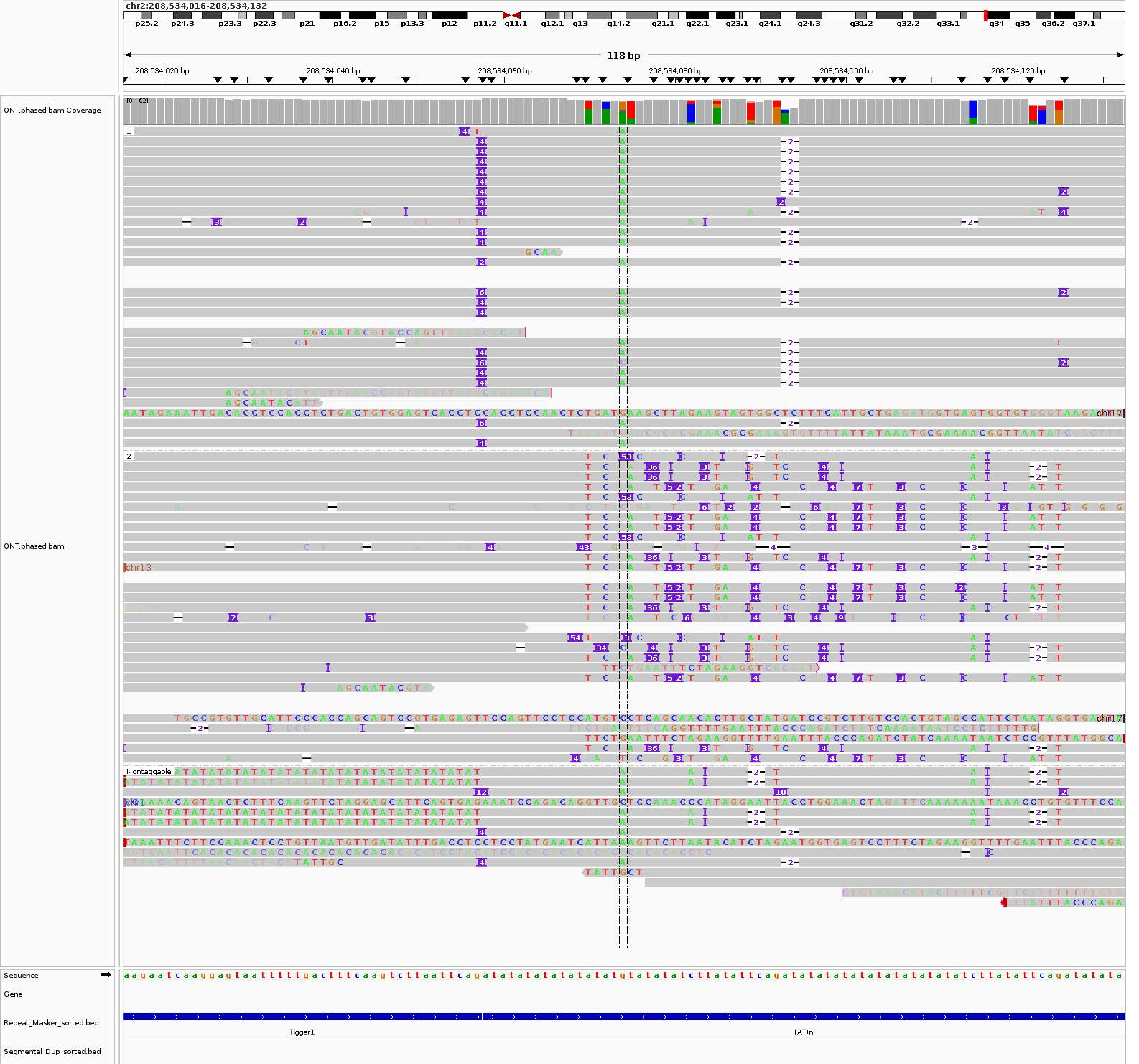
chr2:208,534,016-208,534,132
118 bp
208,534,020 bp
208,534,040 bp
208,534,060 bp
208,534,080 bp
208,534,100 bp
208,534,120 bp
ONT.phased.bam Coverage
ONT.phased.bam
Nontaggable
Sequence
Gene
Repeat_Masker_sorted.bed
Tigger1
(AT)n
Segmental_Dup_sorted.bed

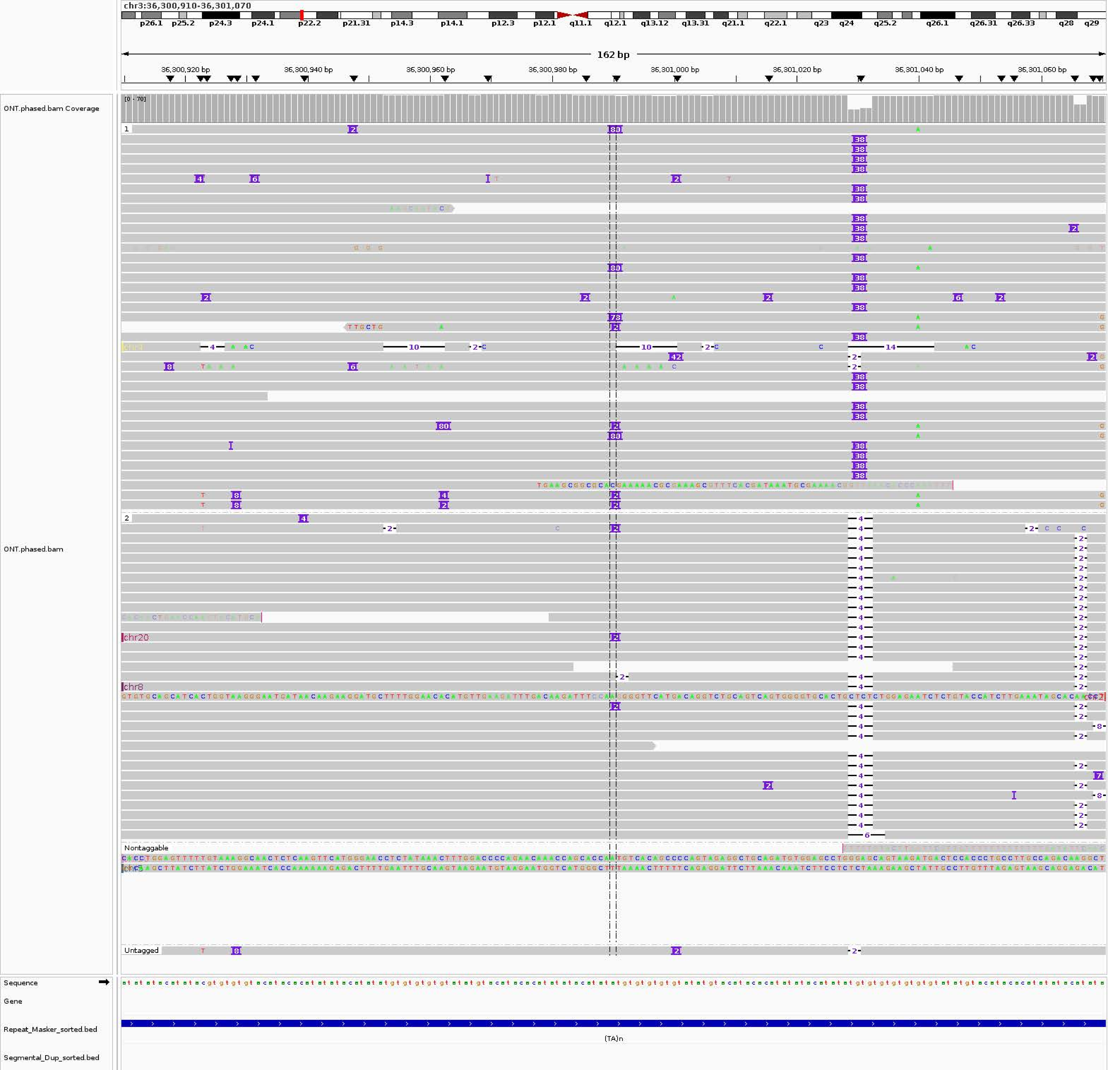

chr3:36,300,910-36,301,070
p26.1
p25.2
p24.3
p24.1
p22.2
p21.31
p14.3
p14.1
p12.3
p12.1
q11.1
q12.1
q13.12
q13.31
q21.1
q22.1
q23
q24
q25.2
q26.1
q26.31
q26.33
q28
q29
162 bp
36,300,920 bp
36,300,940 bp
36,300,960 bp
36,300,980 bp
36,301,000 bp
36,301,020 bp
36,301,040 bp
36,301,060 bp
ONT.phased.bam Coverage
[0 - 70]
ONT.phased.bam
1
2
chr20
chr8
Nontaggable
Untagged
Sequence
Gene
Repeat_Masker_sorted.bed
(TA)n
Segmental_Dup_sorted.bed

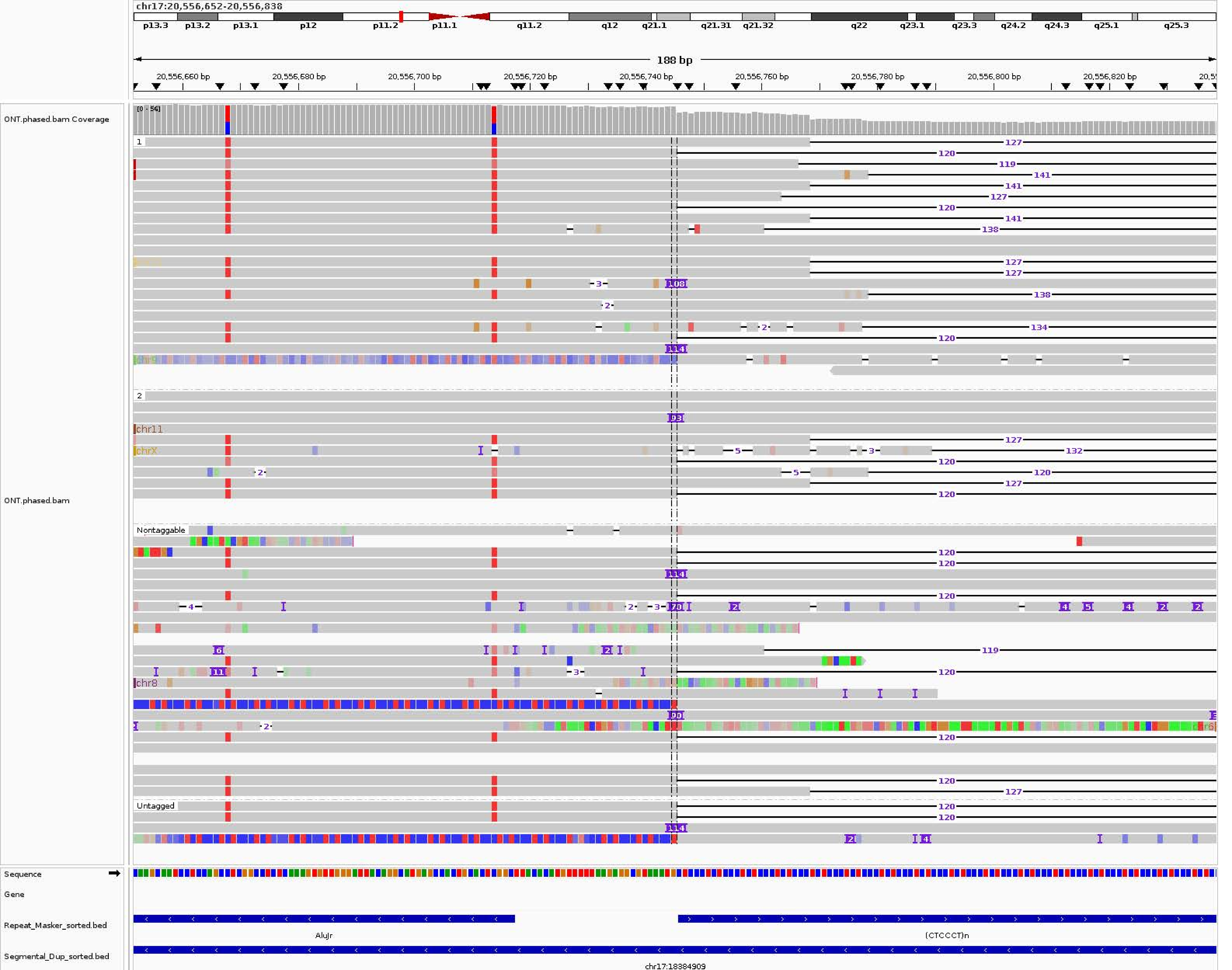

chr17:20,556,652-20,556,838
p13.3 p13.2 p13.1 p12 p11.2 p11.1 q11.2 q12 q21.1 q21.31 q21.32 q22 q23.1 q23.3 q24.2 q24.3 q25.1 q25.3
188 bp
20,556,660 bp 20,556,680 bp 20,556,700 bp 20,556,720 bp 20,556,740 bp 20,556,760 bp 20,556,780 bp 20,556,800 bp 20,556,820 bp
ONT.phased.bam Coverage
[0 - 56]
ONT.phased.bam
1
2
Nontaggable
Untagged
chr9
chr11
chrX
chr8
Sequence
Gene
Repeat_Masker_sorted.bed
AluJr
(CTCCCT)n
Segmental_Dup_sorted.bed
chr17:18384909

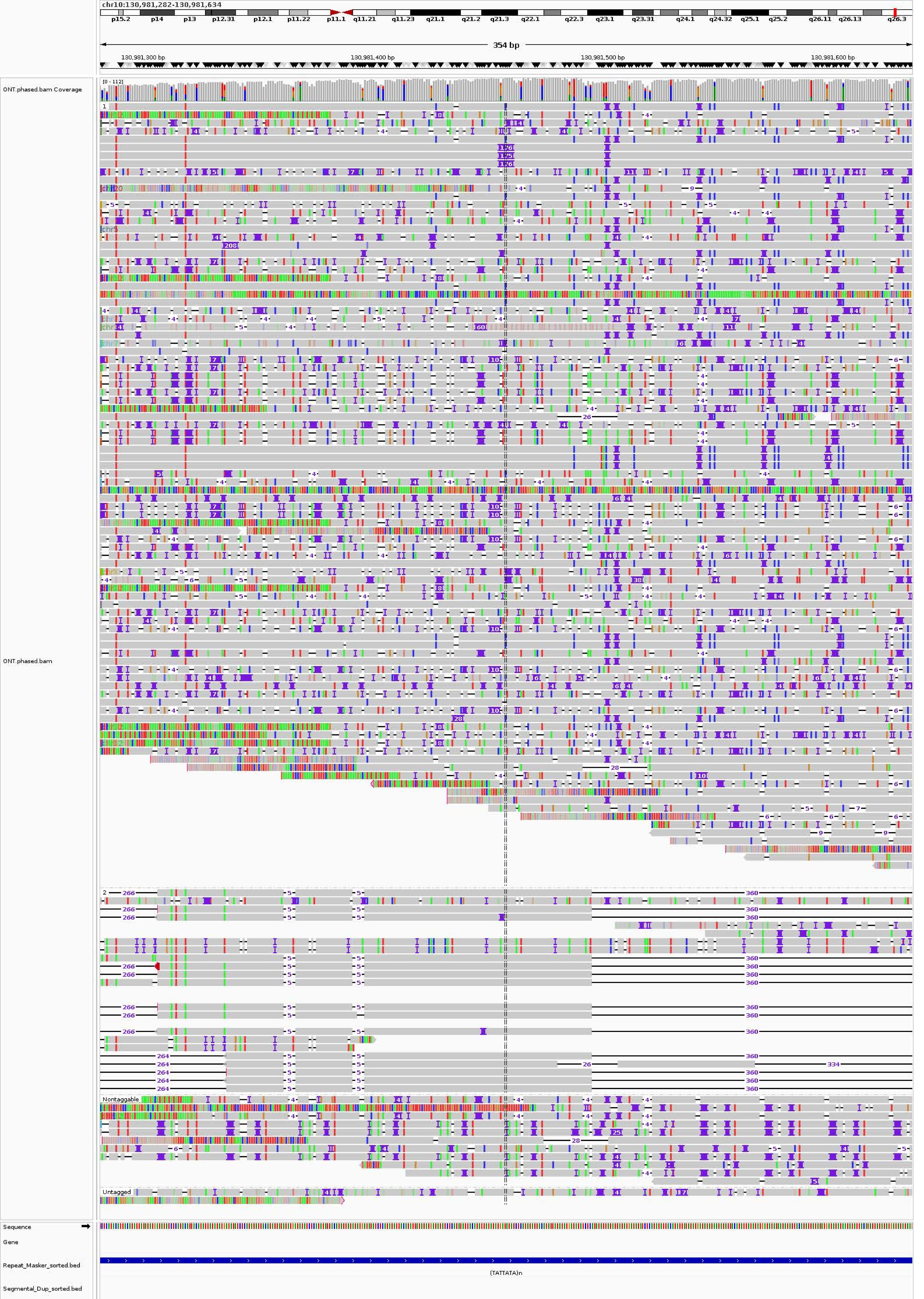

chr10:130,981,282-130,981,634
354 bp
130,981,300 bp
130,981,400 bp
130,981,500 bp
130,981,600 bp
ONT.phased.bam Coverage
[0 - 112]
ONT.phased.bam
Nontaggable
Untagged
Sequence
Gene
Repeat_Masker_sorted.bed
Segmental_Dup_sorted.bed
(TATTATA)n

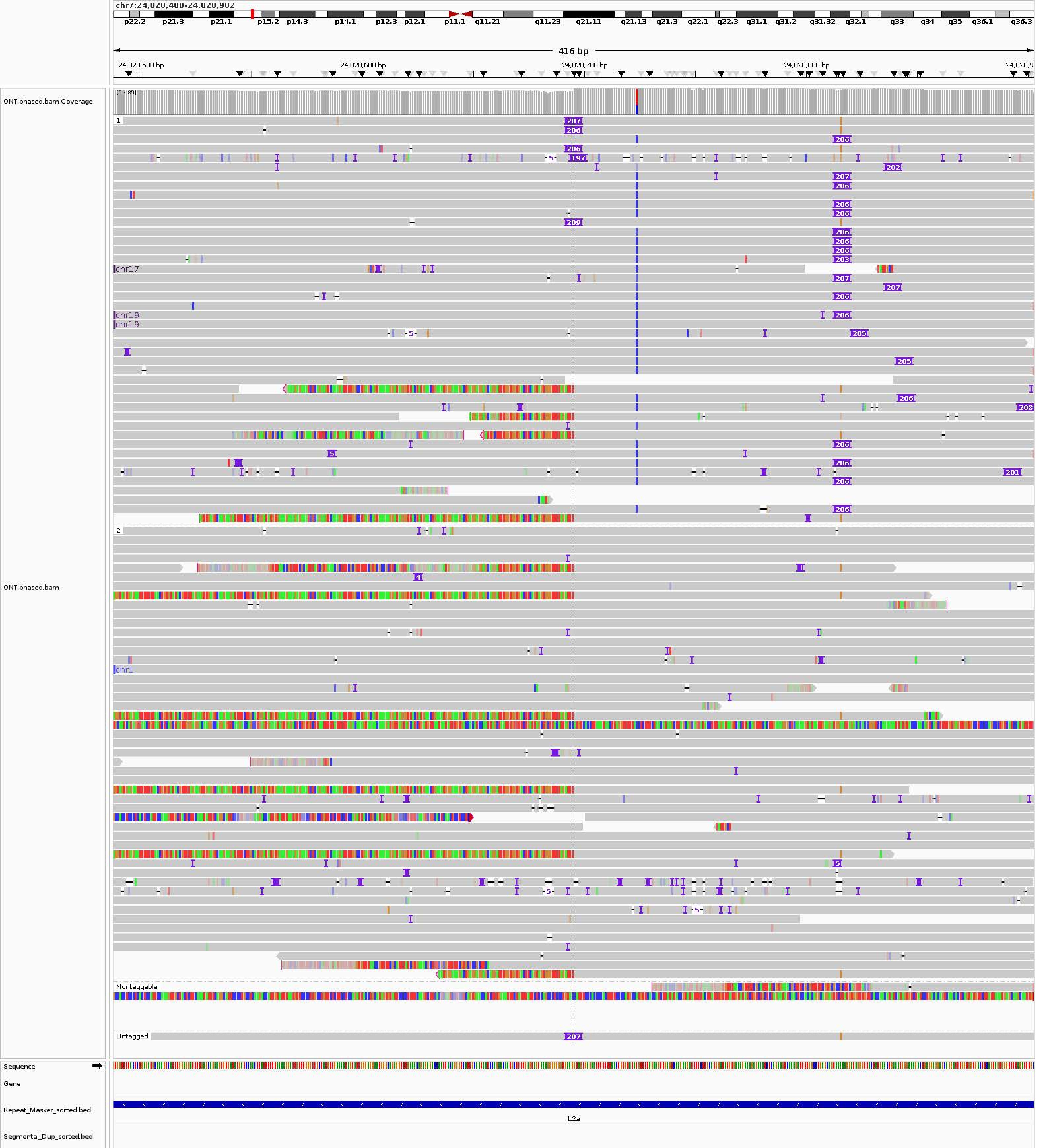
chr7:24,028,488-24,028,902
p22.2 p21.3 p21.1 p15.2 p14.3 p14.1 p12.3 p12.1 p11.1 q11.21 q11.23 q21.11 q21.13 q21.3 q22.1 q22.3 q31.1 q31.2 q31.32 q32.1 q33 q34 q35 q36.1 q36.3
416 bp
24,028,500 bp
24,028,600 bp
24,028,700 bp
24,028,800 bp
ONT.phased.bam Coverage
[0 - 89]
ONT.phased.bam
1
2
chr17
chr19
chr1
Nontaggable
Untagged
Sequence
Gene
Repeat_Masker_sorted.bed
L2a
Segmental_Dup_sorted.bed

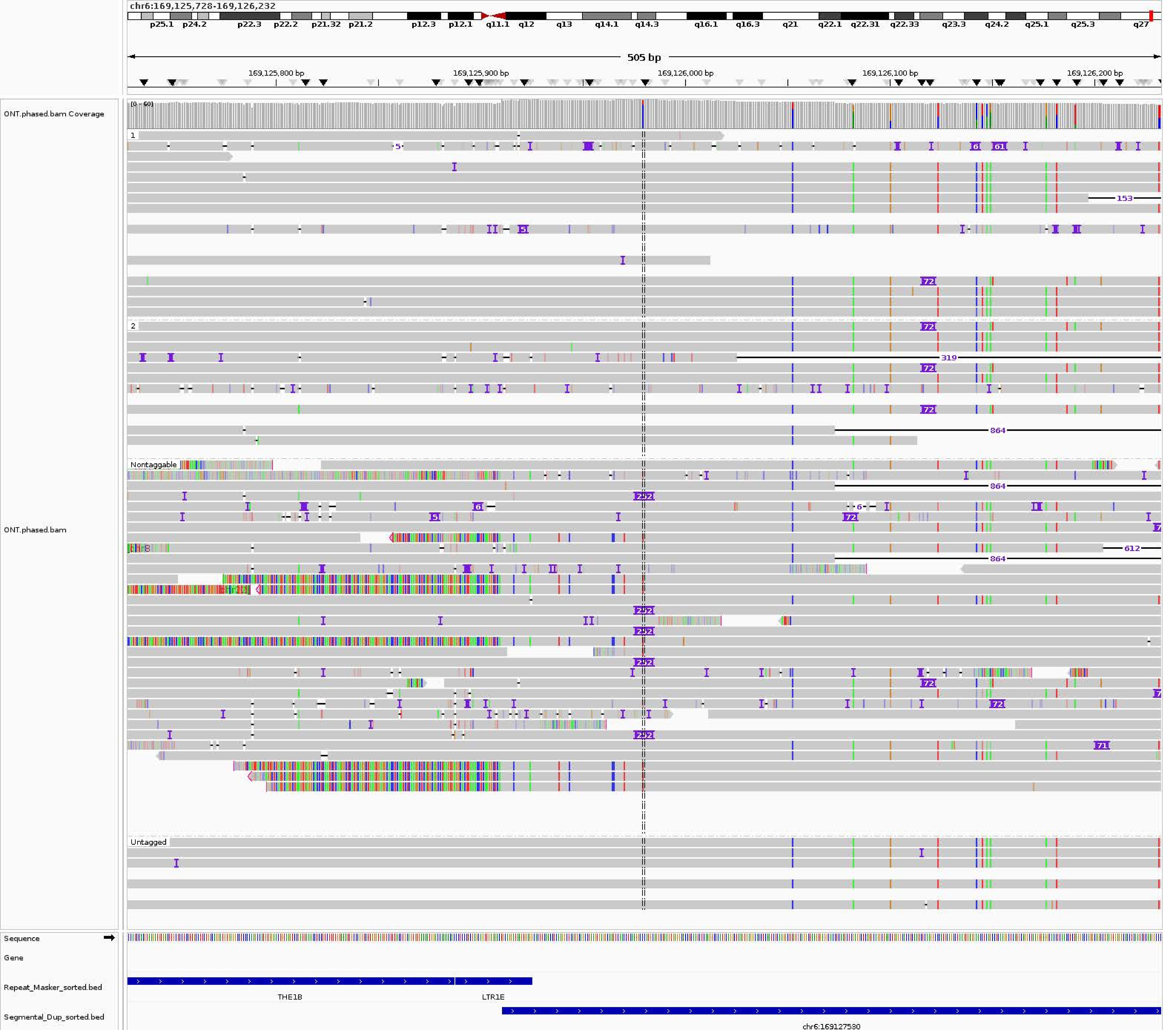
chr6:169,125,728-169,126,232
p25.1 p24.2 p22.3 p22.2 p21.32 p21.2 p12.3 p12.1 q11.1 q12 q13 q14.1 q14.3 q16.1 q16.3 q21 q22.1 q22.31 q22.33 q23.3 q24.2 q25.1 q25.3 q27
505 bp
169,125,800 bp
169,125,900 bp
169,126,000 bp
169,126,100 bp
169,126,200 bp
ONT.phased.bam Coverage
[0 - 60]
ONT.phased.bam
1
2
Nontaggable
Untagged
153
319
864
612
252
72
Sequence
Gene
Repeat_Masker_sorted.bed
THE1B
LTR1E
Segmental_Dup_sorted.bed
chr6:169127530

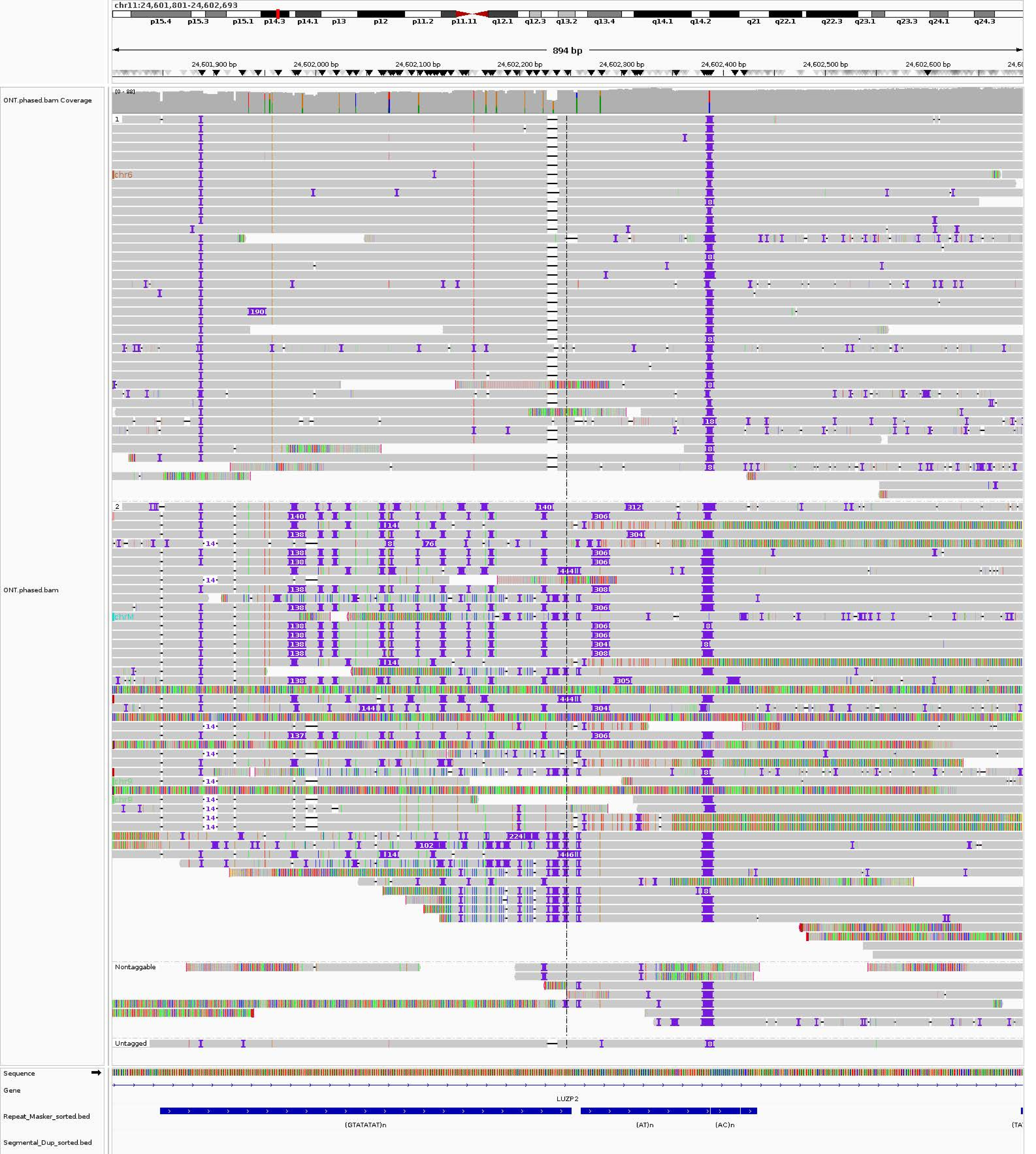
chr11:24,601,801-24,602,693
p15.4
p15.3
p15.1
p14.3
p14.1
p13
p12
p11.2
p11.11
q12.1
q12.3
q13.2
q13.4
q14.1
q14.2
q21
q22.1
q22.3
q23.1
q23.3
q24.1
q24.3
894 bp
24,601,900 bp
24,602,000 bp
24,602,100 bp
24,602,200 bp
24,602,300 bp
24,602,400 bp
24,602,500 bp
24,602,600 bp
ONT.phased.bam Coverage
[0 - 88]
ONT.phased.bam
chr6
chrM
chr9
Nontaggable
Untagged
Sequence
Gene
LUZP2
Repeat_Masker_sorted.bed
(GTATATAT)n
(AT)n
(AC)n
Segmental_Dup_sorted.bed

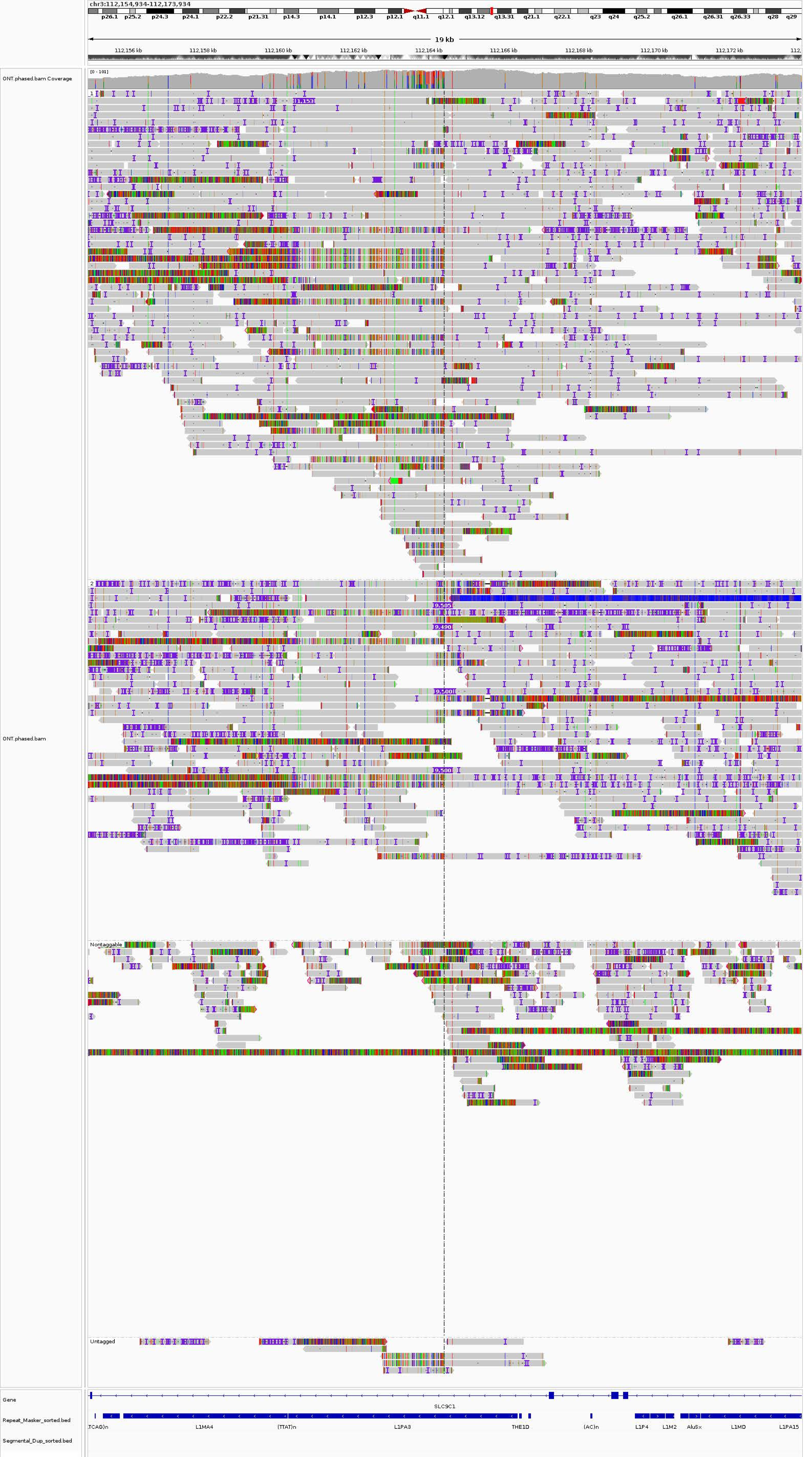
chr3:112,154,934-112,173,934
p26.1 p25.2 p24.3 p24.1 p22.2 p21.31 p14.3 p14.1 p12.3 p12.1 q11.1 q12.1 q13.12 q13.31 q21.1 q22.1 q23 q24 q25.2 q26.1 q26.31 q26.33 q28 q29
19 kb
112,156 kb 112,158 kb 112,160 kb 112,162 kb 112,164 kb 112,166 kb 112,168 kb 112,170 kb 112,172 kb
ONT.phased.bam Coverage
[0 - 101]
ONT.phased.bam
1
2
Nontaggable
Untagged
Gene
SLC9C1
Repeat_Masker_sorted.bed
Segmental_Dup_sorted.bed
TCAG)n
L1MA4
(TTAT)n
L1PA3
THE1D
(AC)n
L1P4
L1M2
AluSx
L1MD
L1PA15
