## Supplementary material for "A personalized multi-platform assessment of somatic mosaicism in the human frontal cortex": Supplementary Table 8.pdf

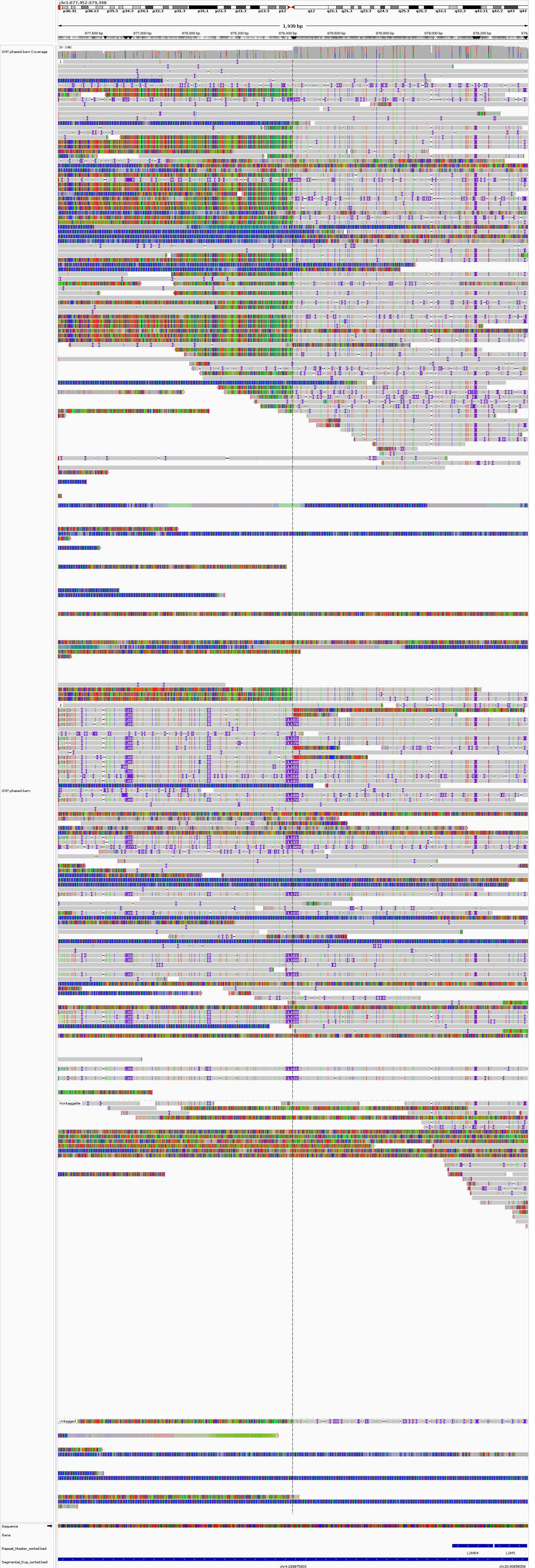
chr1:877,452-879,390
1,939 bp
877,600 bp
877,800 bp
878,000 bp
878,200 bp
878,400 bp
878,600 bp
878,800 bp
879,000 bp
879,200 bp
ONT.phased.bam Coverage
ONT.phased.bam
Nontaggable
Untagged
Sequence
Gene
Repeat_Masker_sorted.bed
Segmental_Dup_sorted.bed
L1MB4
L1M5
chr4:189975803
chr20:30856058

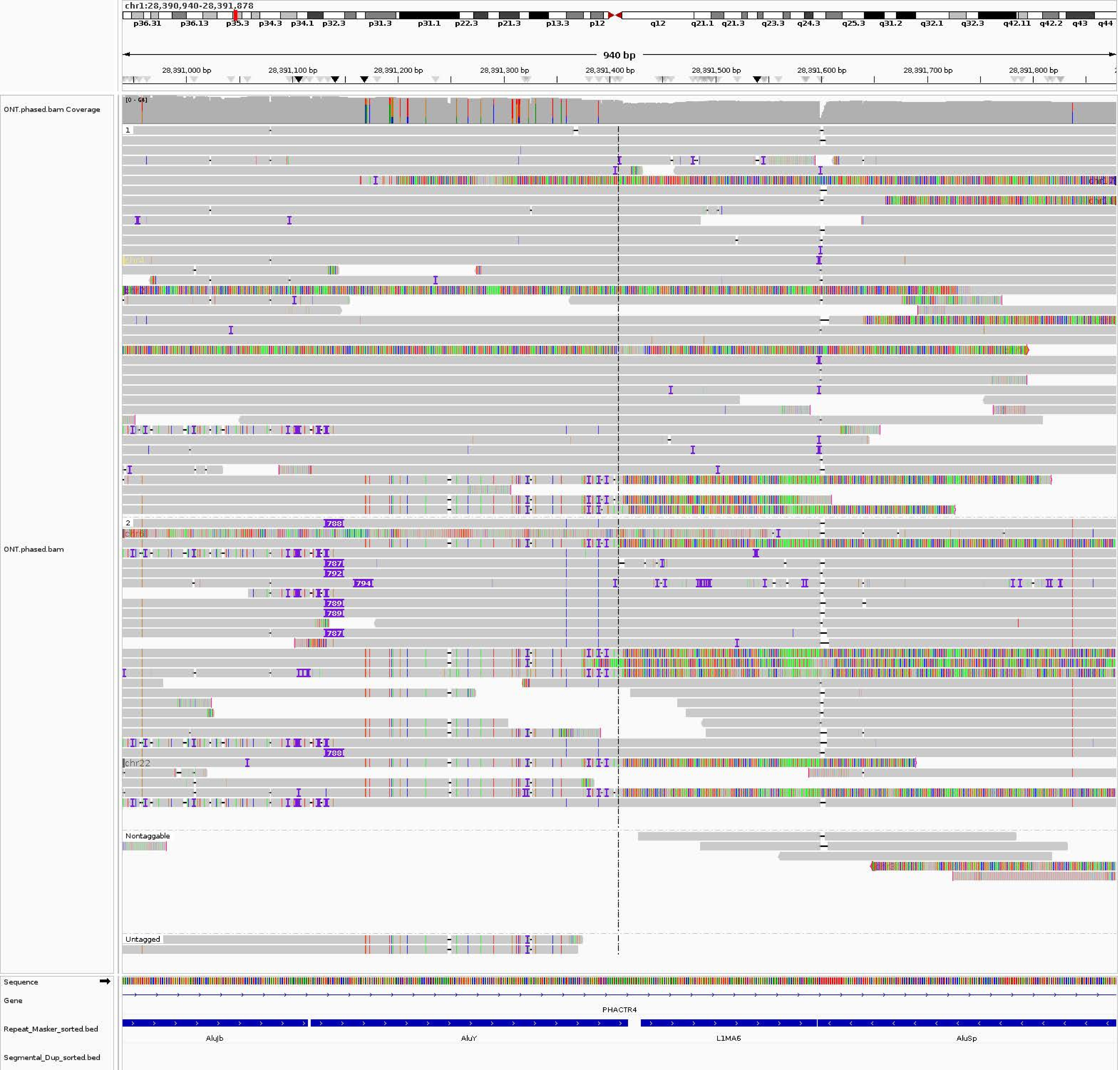
chr1:28,390,940-28,391,878
p36.31
p36.13
p35.3
p34.3
p34.1
p32.3
p31.3
p31.1
p22.3
p21.3
p13.3
p12
q12
q21.1
q21.3
q23.3
q24.3
q25.3
q31.2
q32.1
q32.3
q42.11
q42.2
q43
q44
940 bp
28,391,000 bp
28,391,100 bp
28,391,200 bp
28,391,300 bp
28,391,400 bp
28,391,500 bp
28,391,600 bp
28,391,700 bp
28,391,800 bp
ONT.phased.bam Coverage
[0 - 64]
ONT.phased.bam
1
2
788
787
792
794
789
789
787
788
chr22
Nontaggable
Untagged
Sequence
Gene
PHACTR4
Repeat_Masker_sorted.bed
AluJb
AluY
L1MA6
AluSp
Segmental_Dup_sorted.bed

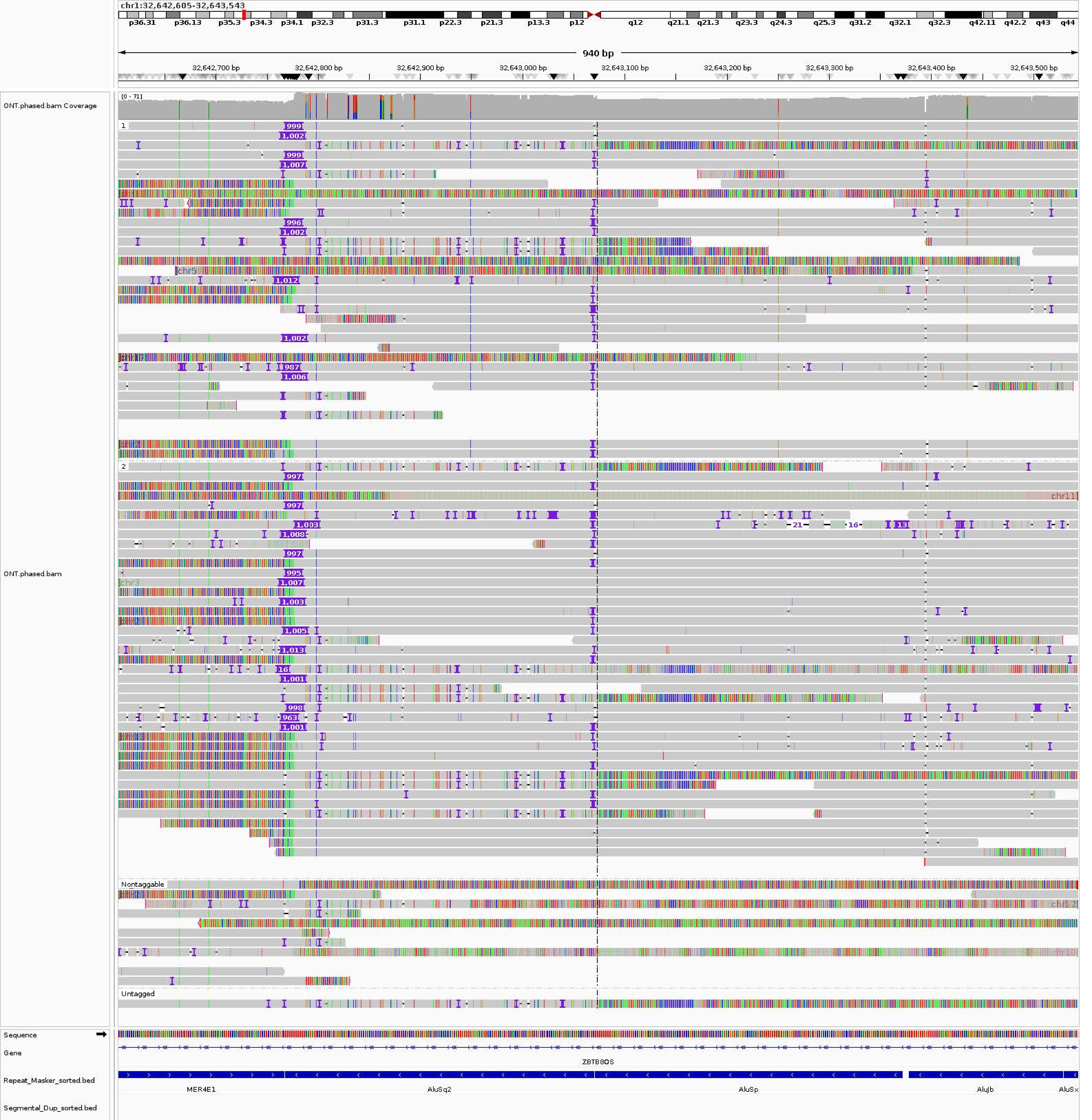

chr1:32,642,605-32,643,543
p36.31
p36.13
p35.3
p34.3
p34.1
p32.3
p31.3
p31.1
p22.3
p21.3
p13.3
p12
q12
q21.1
q21.3
q23.3
q24.3
q25.3
q31.2
q32.1
q32.3
q42.11
q42.2
q43
q44
940 bp
32,642,700 bp
32,642,800 bp
32,642,900 bp
32,643,000 bp
32,643,100 bp
32,643,200 bp
32,643,300 bp
32,643,400 bp
32,643,500 bp
ONT.phased.bam Coverage
[0 - 71]
ONT.phased.bam
1
2
Nontaggable
Untagged
chr5
chr11
chr3
chr12
Sequence
Gene
ZBTB8OS
Repeat_Masker_sorted.bed
MER4E1
AluSq2
AluSp
AluJb
AluSx
Segmental_Dup_sorted.bed

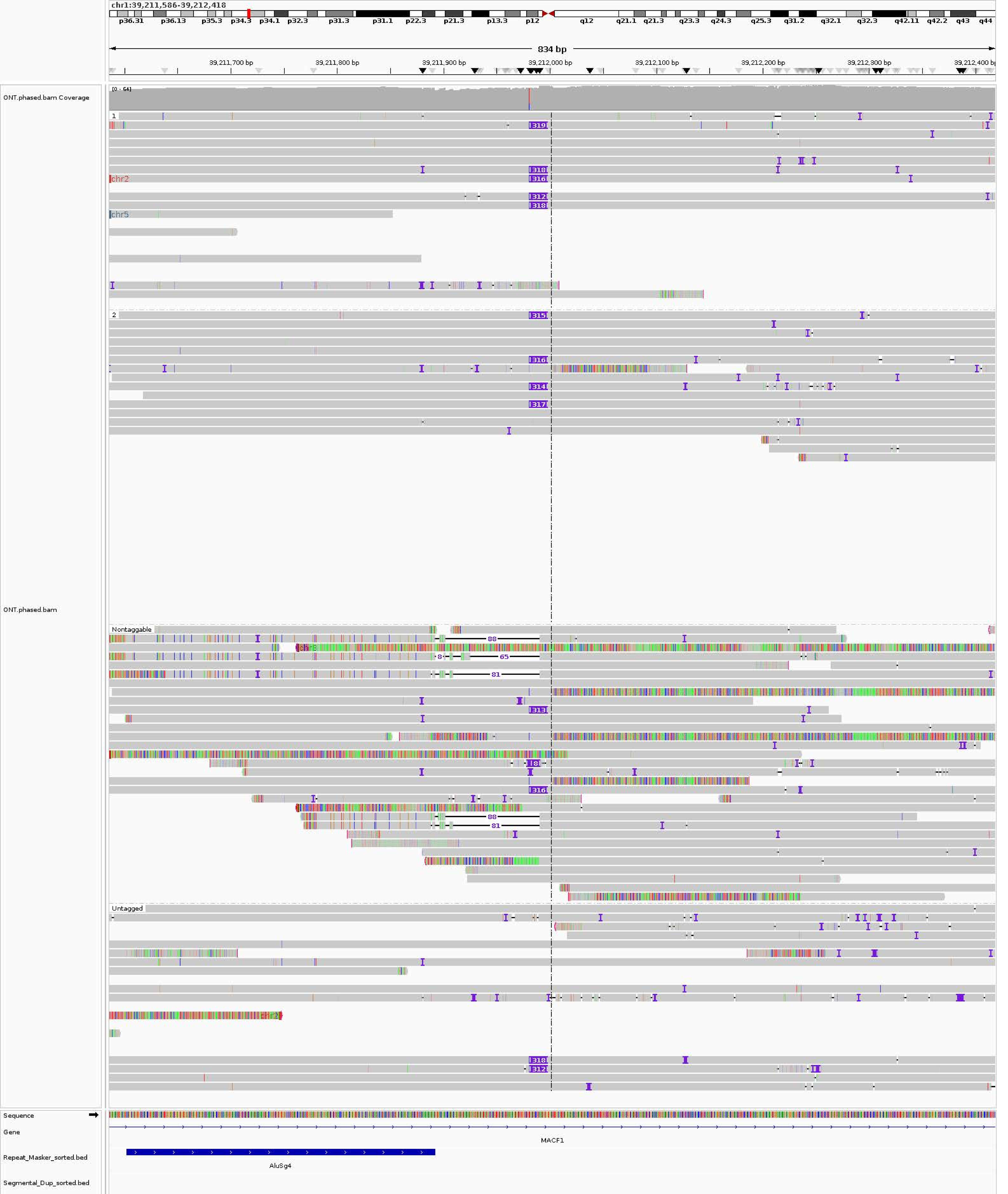
chr1:39,211,586-39,212,418
p36.31 p36.13 p35.3 p34.3 p34.1 p32.3 p31.3 p31.1 p22.3 p21.3 p13.3 p12 q12 q21.1 q21.3 q23.3 q24.3 q25.3 q31.2 q32.1 q32.3 q42.11 q42.2 q43 q44
834 bp
39,211,700 bp
39,211,800 bp
39,211,900 bp
39,212,000 bp
39,212,100 bp
39,212,200 bp
39,212,300 bp
39,212,400 bp
ONT.phased.bam Coverage
[0 - 64]
ONT.phased.bam
1
2
Nontaggable
Untagged
chr2
chr5
Sequence
Gene
MACF1
Repeat_Masker_sorted.bed
AluSg4
Segmental_Dup_sorted.bed

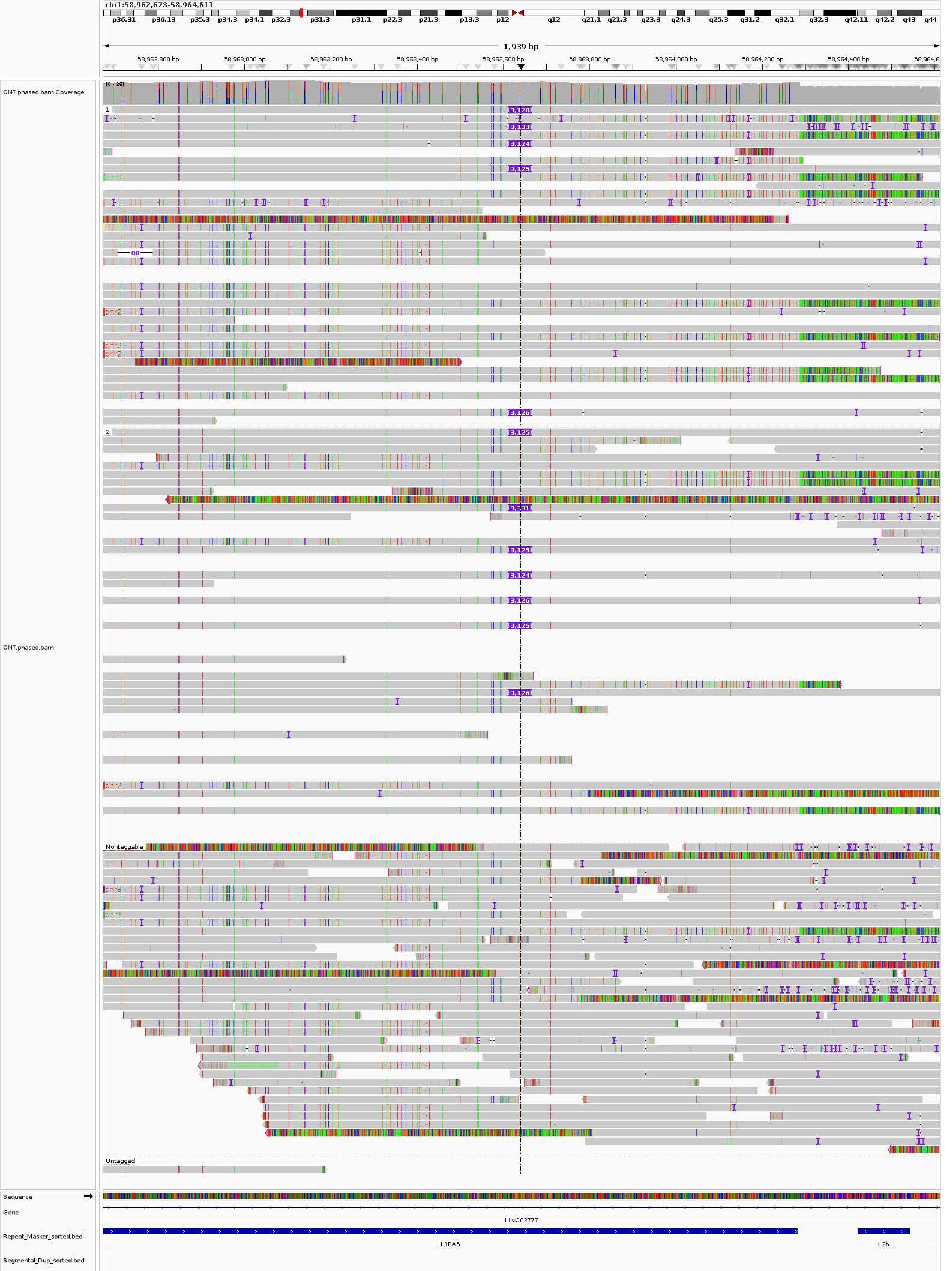
chr1:58,962,673-58,964,611
p36.31 p36.13 p35.3 p34.3 p34.1 p32.3 p31.3 p31.1 p22.3 p21.3 p13.3 p12 q12 q21.1 q21.3 q23.3 q24.3 q25.3 q31.2 q32.1 q32.3 q42.11 q42.2 q43 q44
1,939 bp
58,962,800 bp
58,963,000 bp
58,963,200 bp
58,963,400 bp
58,963,600 bp
58,963,800 bp
58,964,000 bp
58,964,200 bp
58,964,400 bp
ONT.phased.bam Coverage
[0 - 86]
ONT.phased.bam
1
2
Nontaggable
Untagged
3,128
3,133
3,124
3,125
3,126
3,331
80
chr2
chr8
chr3
Sequence
Gene
LINC02777
Repeat_Masker_sorted.bed
L1PA5
L2b
Segmental_Dup_sorted.bed

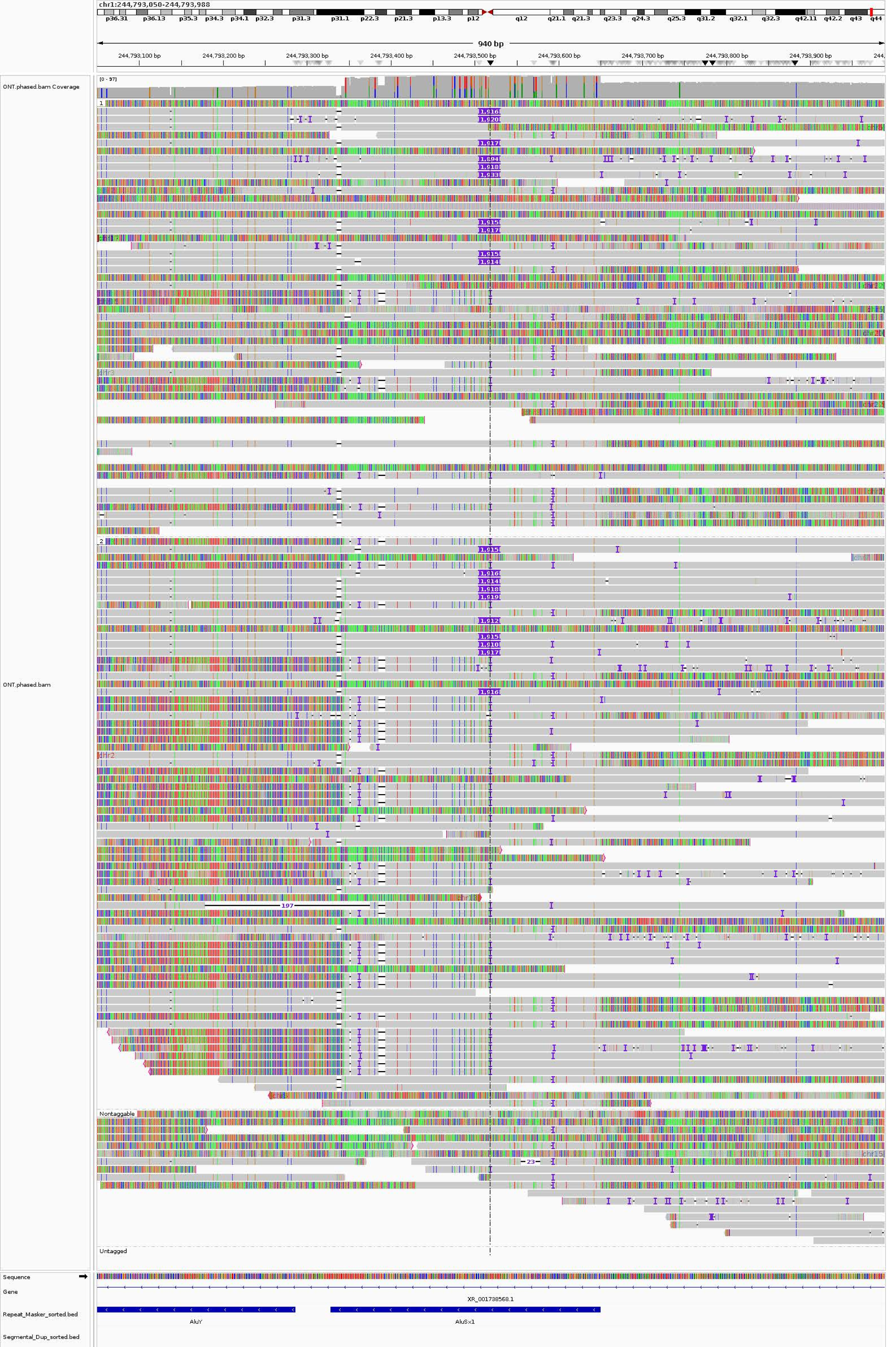
chr1:244,793,050-244,793,988
940 bp
244,793,100 bp
244,793,200 bp
244,793,300 bp
244,793,400 bp
244,793,500 bp
244,793,600 bp
244,793,700 bp
244,793,800 bp
244,793,900 bp
ONT.phased.bam Coverage
[0 - 97]
ONT.phased.bam
1
2
Nontaggable
Untagged
197
23
Sequence
Gene
XR_001738568.1
Repeat_Masker_sorted.bed
AluY
AluSx1
Segmental_Dup_sorted.bed

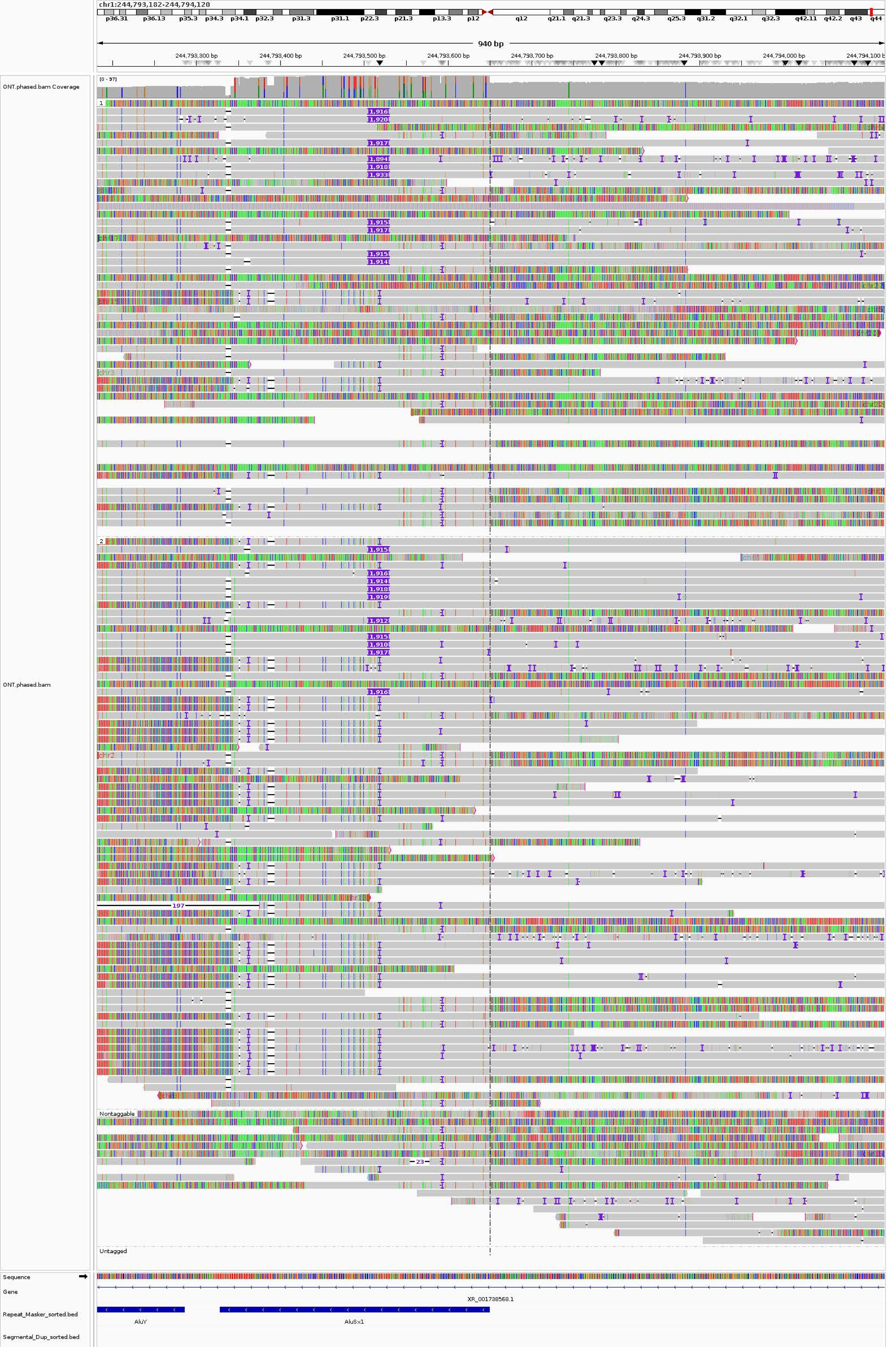

chr1:244,793,182-244,794,120
p36.31 p36.13 p35.3 p34.3 p34.1 p32.3 p31.3 p31.1 p22.3 p21.3 p13.3 p12 q12 q21.1 q21.3 q23.3 q24.3 q25.3 q31.2 q32.1 q32.3 q42.11 q42.2 q43 q44
940 bp
244,793,300 bp
244,793,400 bp
244,793,500 bp
244,793,600 bp
244,793,700 bp
244,793,800 bp
244,793,900 bp
244,794,000 bp
ONT.phased.bam Coverage
[0 - 97]
ONT.phased.bam
1
2
Nontaggable
Untagged
197
23
Sequence
Gene
XR_001738568.1
Repeat_Masker_sorted.bed
AluY
AluSx1
Segmental_Dup_sorted.bed

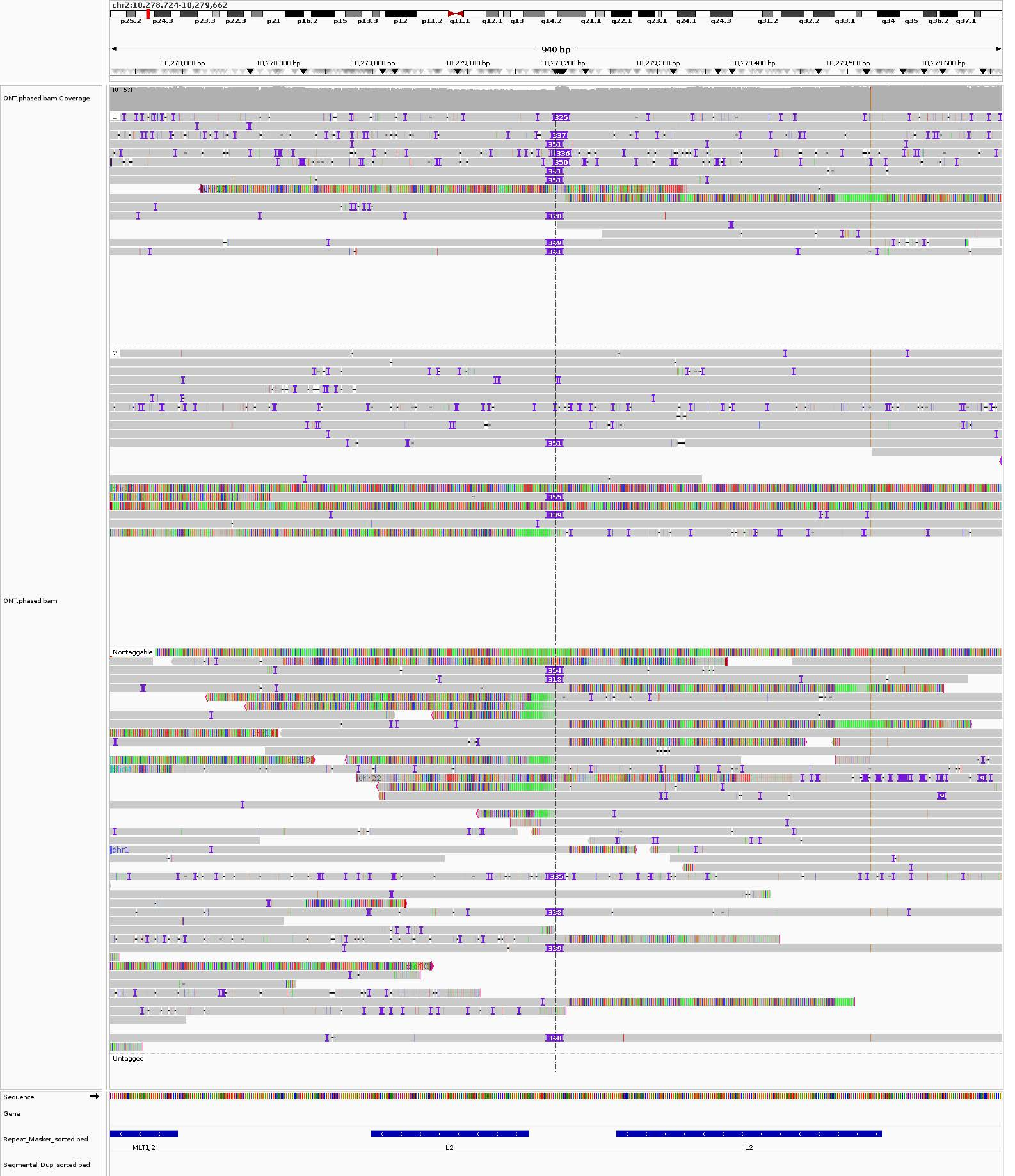
chr2:10,278,724-10,279,662
p25.2
p24.3
p23.3
p22.3
p21
p16.2
p15
p13.3
p12
p11.2
q11.1
q12.1
q13
q14.2
q21.1
q22.1
q23.1
q24.1
q24.3
q31.2
q32.2
q33.1
q34
q35
q36.2
q37.1
940 bp
10,278,800 bp
10,278,900 bp
10,279,000 bp
10,279,100 bp
10,279,200 bp
10,279,300 bp
10,279,400 bp
10,279,500 bp
10,279,600 bp
ONT.phased.bam Coverage
[0 - 57]
ONT.phased.bam
1
2
Nontaggable
Untagged
chr22
chr1
Sequence
Gene
Repeat_Masker_sorted.bed
Segmental_Dup_sorted.bed
MLT1J2
L2
L2

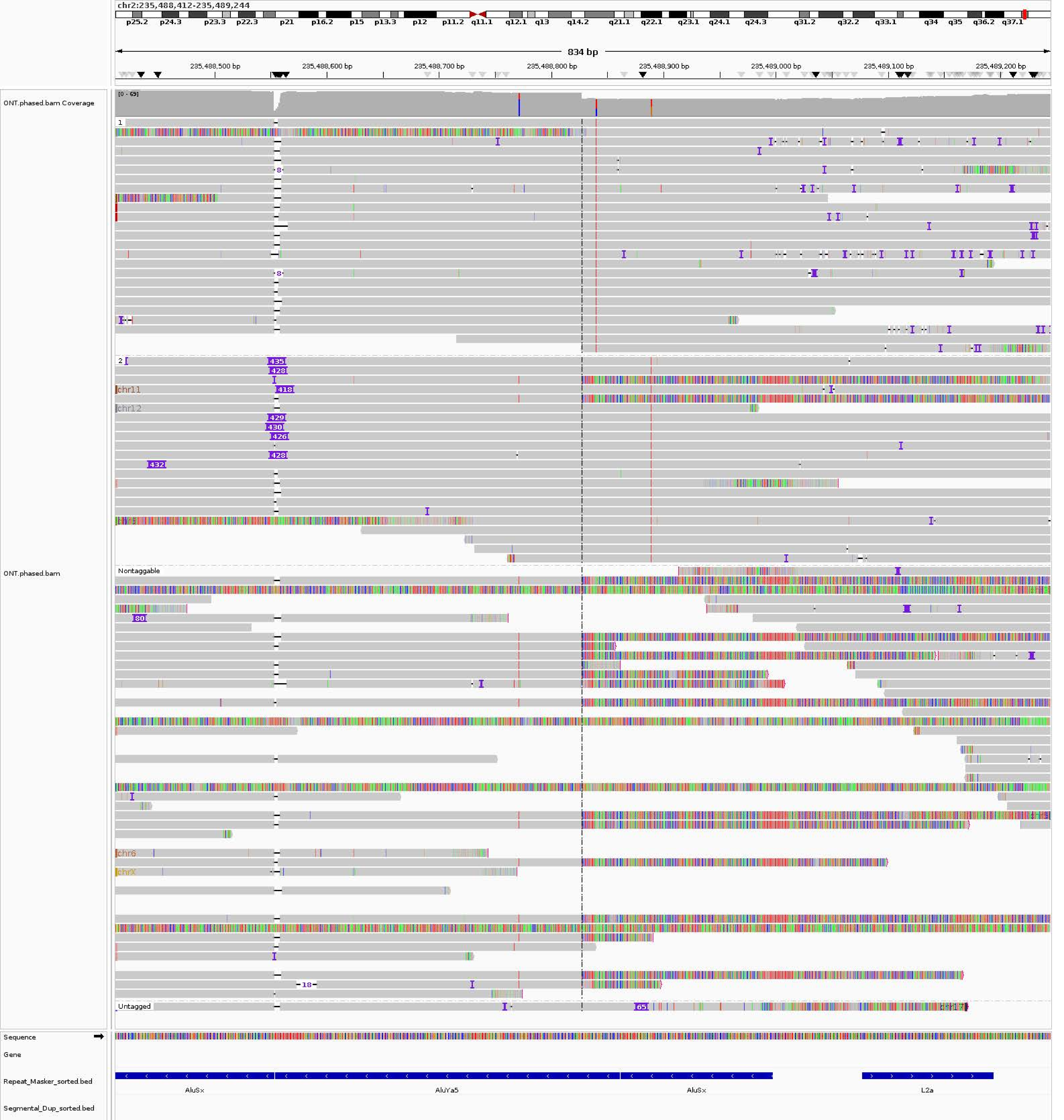

chr2:235,488,412-235,489,244
p25.2 p24.3 p23.3 p22.3 p21 p16.2 p15 p13.3 p12 p11.2 q11.1 q12.1 q13 q14.2 q21.1 q22.1 q23.1 q24.1 q24.3 q31.2 q32.2 q33.1 q34 q35 q36.2 q37.1
834 bp
235,488,500 bp
235,488,600 bp
235,488,700 bp
235,488,800 bp
235,488,900 bp
235,489,000 bp
235,489,100 bp
235,489,200 bp
ONT.phased.bam Coverage
[0 - 69]
ONT.phased.bam
1
2
435
428
418
429
430
426
428
432
chr11
chr12
Nontaggable
80
chr6
chrX
18
Untagged
65
Sequence
Gene
Repeat_Masker_sorted.bed
AluSx
AluYa5
AluSx
L2a
Segmental_Dup_sorted.bed

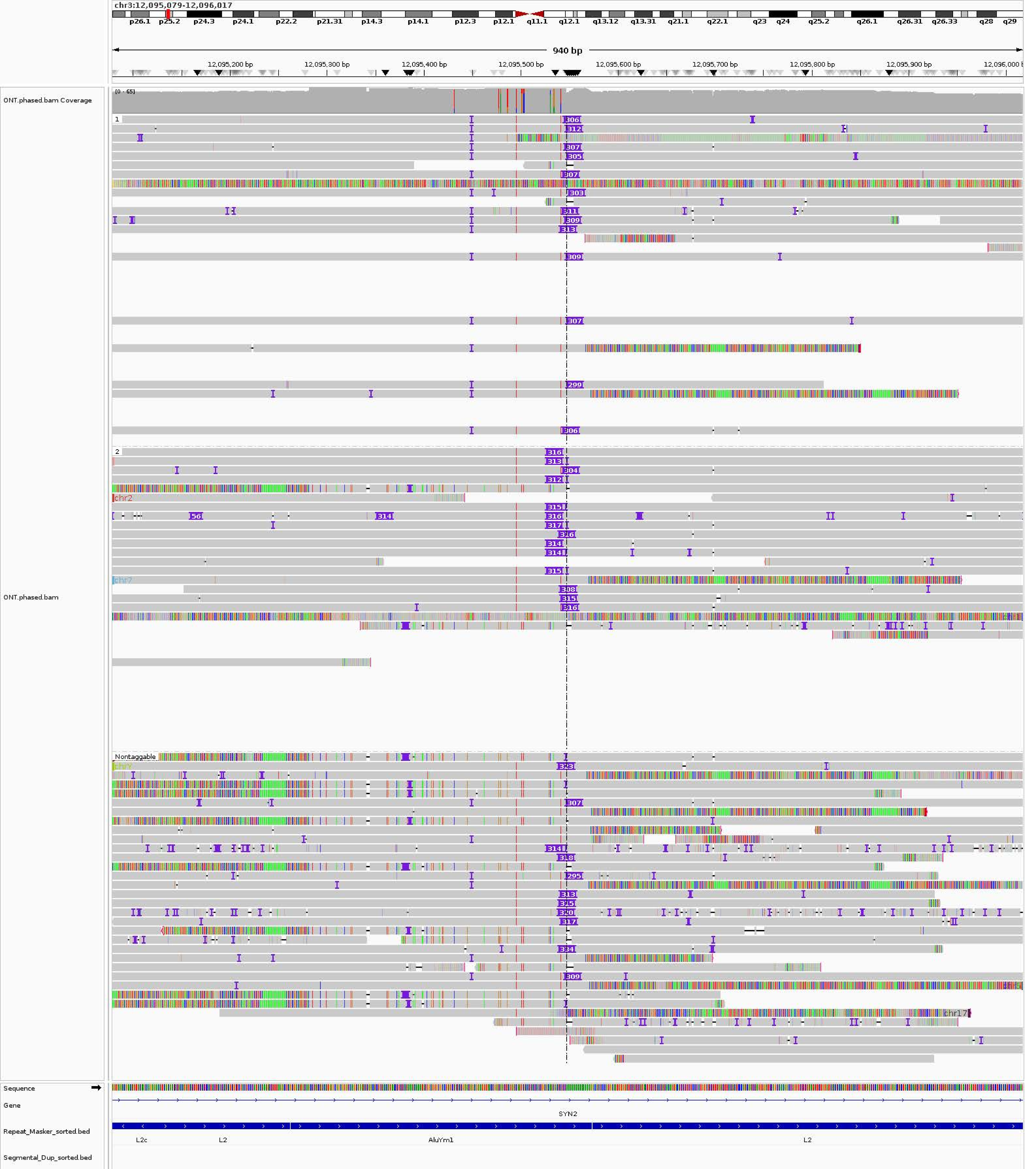
chr3:12,095,079-12,096,017
p26.1
p25.2
p24.3
p24.1
p22.2
p21.31
p14.3
p14.1
p12.3
p12.1
q11.1
q12.1
q13.12
q13.31
q21.1
q22.1
q23
q24
q25.2
q26.1
q26.31
q26.33
q28
q29
940 bp
12,095,200 bp
12,095,300 bp
12,095,400 bp
12,095,500 bp
12,095,600 bp
12,095,700 bp
12,095,800 bp
12,095,900 bp
ONT.phased.bam Coverage
[0 - 65]
ONT.phased.bam
1
2
Nontaggable
chr2
chr7
chrY
chr17
Sequence
Gene
SYN2
Repeat_Masker_sorted.bed
L2c
L2
AluYm1
L2
Segmental_Dup_sorted.bed

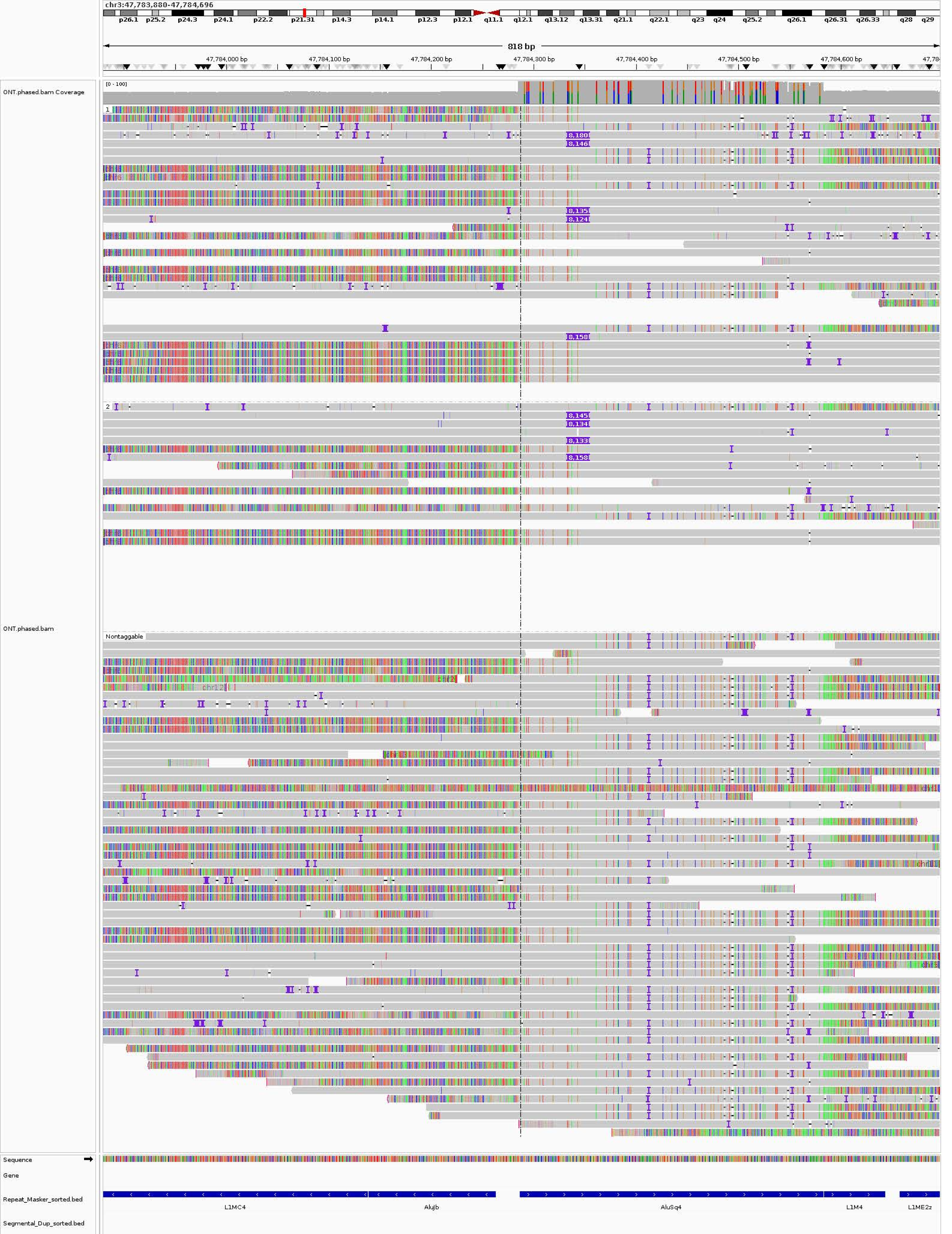
chr3:47,783,880-47,784,696
p26.1
p25.2
p24.3
p24.1
p22.2
p21.31
p14.3
p14.1
p12.3
p12.1
q11.1
q12.1
q13.12
q13.31
q21.1
q22.1
q23
q24
q25.2
q26.1
q26.31
q26.33
q28
q29
818 bp
47,784,000 bp
47,784,100 bp
47,784,200 bp
47,784,300 bp
47,784,400 bp
47,784,500 bp
47,784,600 bp
ONT.phased.bam Coverage
[0 - 100]
1
8,180
8,146
8,135
8,124
8,158
2
8,145
8,134
8,133
8,158
ONT.phased.bam
Nontaggable
chr12
Sequence
Gene
Repeat_Masker_sorted.bed
L1MC4
AluJb
AluSq4
L1M4
L1ME2z
Segmental_Dup_sorted.bed

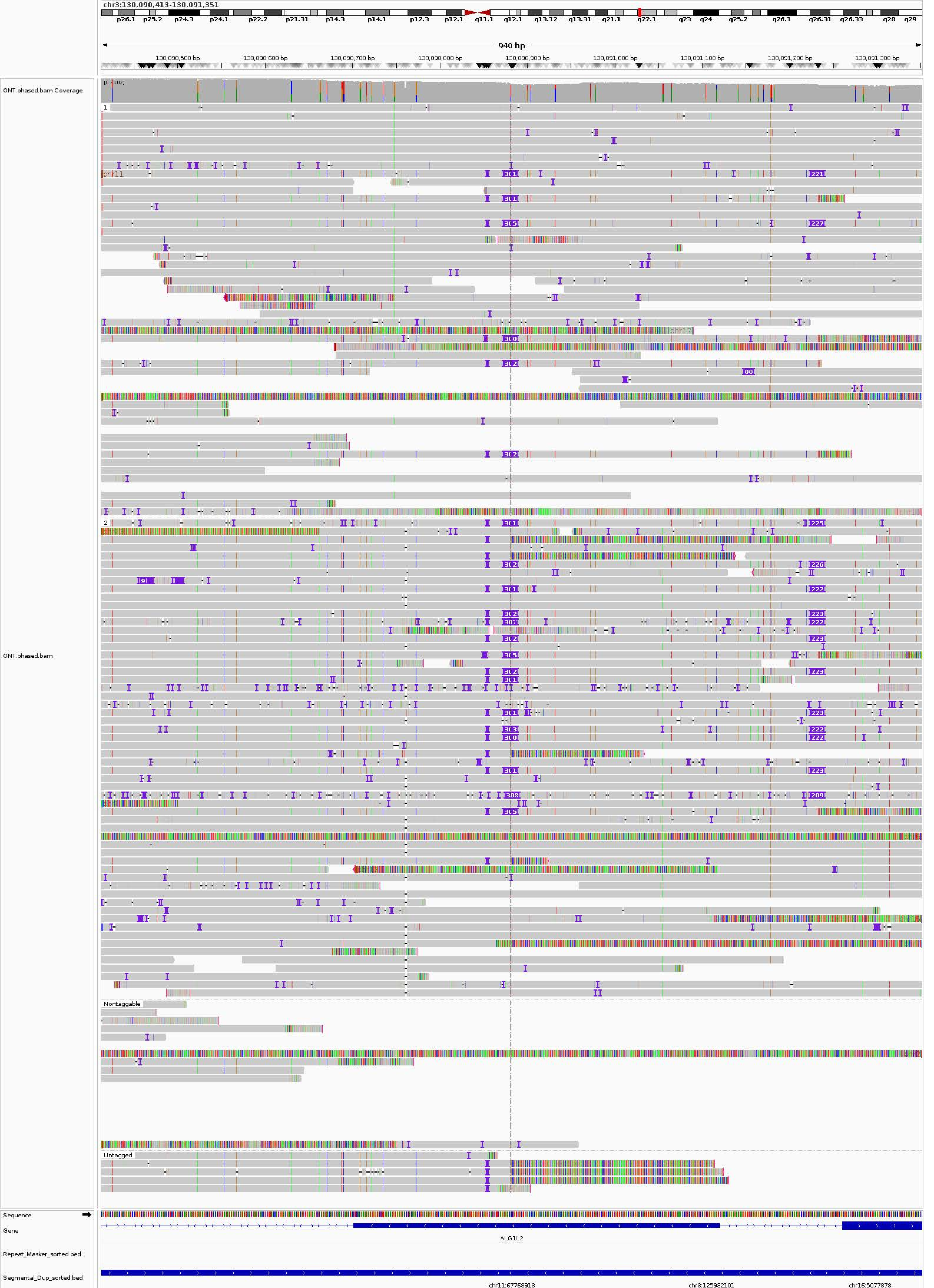
chr3:130,090,413-130,091,351
p26.1 p25.2 p24.3 p24.1 p22.2 p21.31 p14.3 p14.1 p12.3 p12.1 q11.1 q12.1 q13.12 q13.31 q21.1 q22.1 q23 q24 q25.2 q26.1 q26.31 q26.33 q28 q29
940 bp
130,090,500 bp
130,090,600 bp
130,090,700 bp
130,090,800 bp
130,090,900 bp
130,091,000 bp
130,091,100 bp
130,091,200 bp
130,091,300 bp
ONT.phased.bam Coverage
[0 - 102]
ONT.phased.bam
chr11
chr12
Nontaggable
Untagged
Sequence
Gene
ALG1L2
Repeat_Masker_sorted.bed
Segmental_Dup_sorted.bed
chr11:67768913
chr3:125932101
chr16:5077878

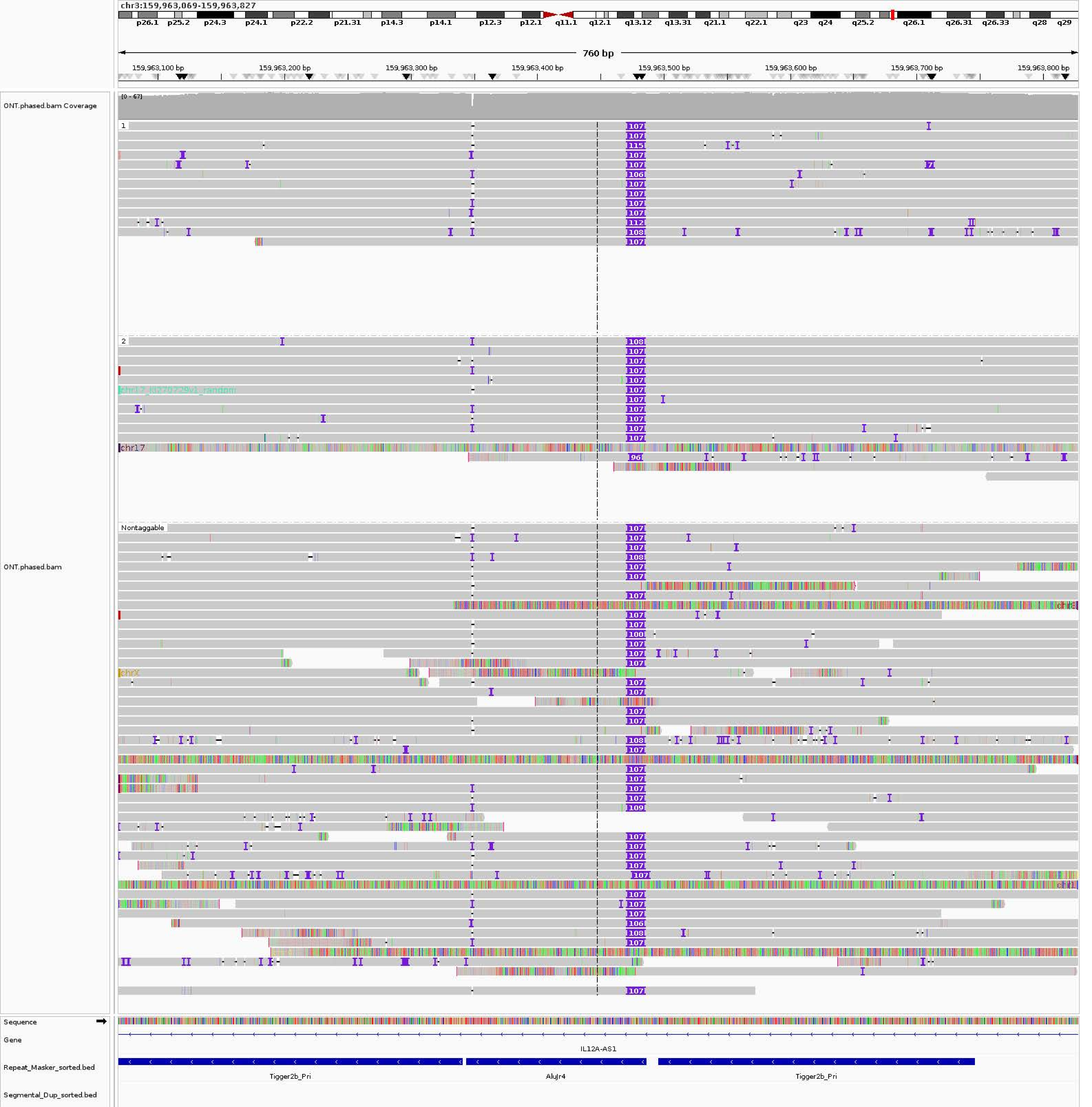
chr3:159,963,069-159,963,827
p26.1 p25.2 p24.3 p24.1 p22.2 p21.31 p14.3 p14.1 p12.3 p12.1 q11.1 q12.1 q13.12 q13.31 q21.1 q22.1 q23 q24 q25.2 q26.1 q26.31 q26.33 q28 q29
760 bp
159,963,100 bp
159,963,200 bp
159,963,300 bp
159,963,400 bp
159,963,500 bp
159,963,600 bp
159,963,700 bp
159,963,800 bp
ONT.phased.bam Coverage
[0 - 67]
ONT.phased.bam
1
2
chr12_KI270729v1_random
chr17
Nontaggable
chrX
Sequence
Gene
IL12A-AS1
Repeat_Masker_sorted.bed
Tigger2b_Pri
AluJr4
Tigger2b_Pri
Segmental_Dup_sorted.bed

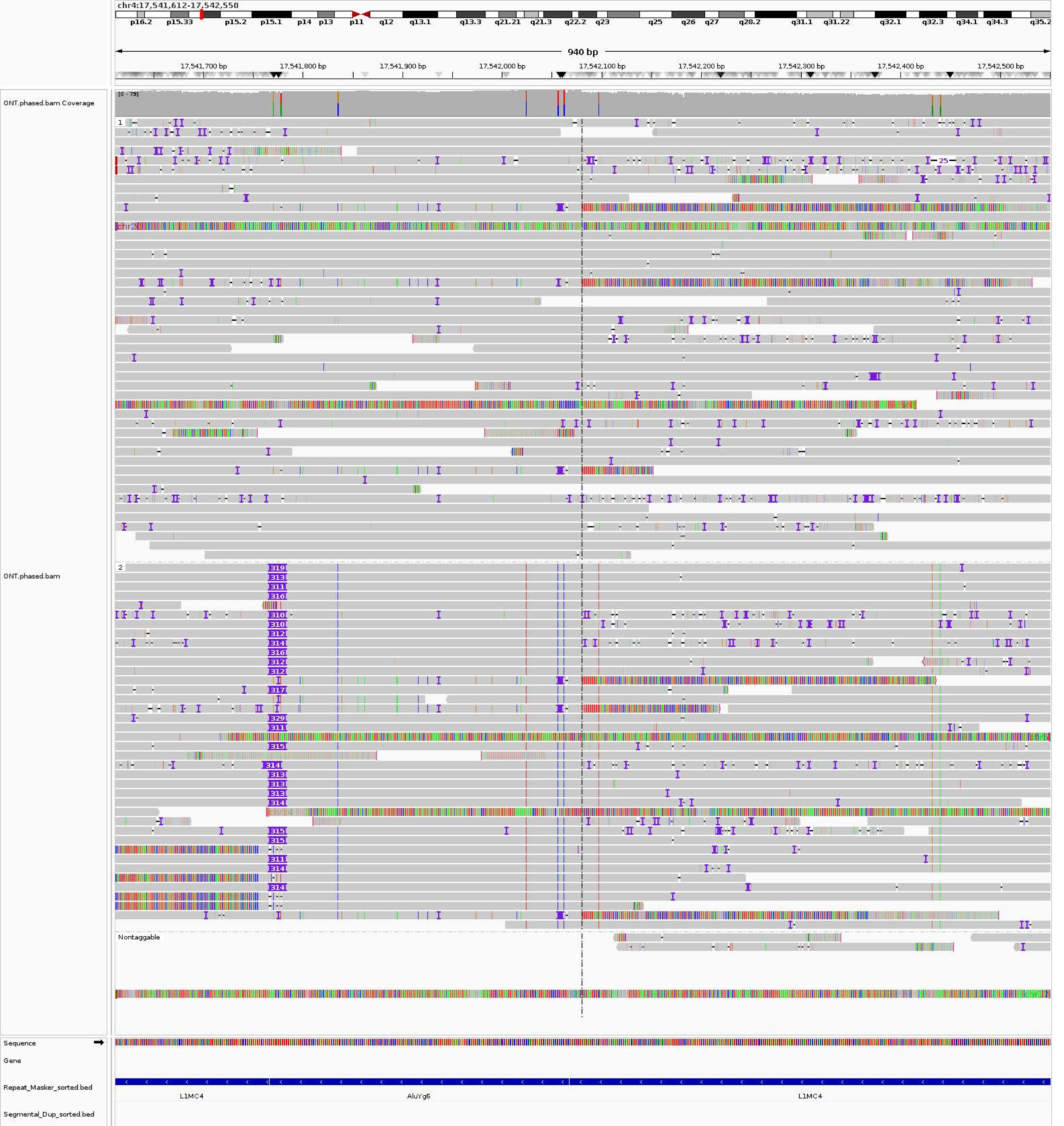

chr4:17,541,612-17,542,550
p16.2 p15.33 p15.2 p15.1 p14 p13 p11 q12 q13.1 q13.3 q21.21 q21.3 q22.2 q23 q25 q26 q27 q28.2 q31.1 q31.22 q32.1 q32.3 q34.1 q34.3 q35.2
940 bp
17,541,700 bp
17,541,800 bp
17,541,900 bp
17,542,000 bp
17,542,100 bp
17,542,200 bp
17,542,300 bp
17,542,400 bp
17,542,500 bp
ONT.phased.bam Coverage
[0 - 79]
ONT.phased.bam
1
2
Nontaggable
Sequence
Gene
Repeat_Masker_sorted.bed
L1MC4
AluYg6
L1MC4
Segmental_Dup_sorted.bed

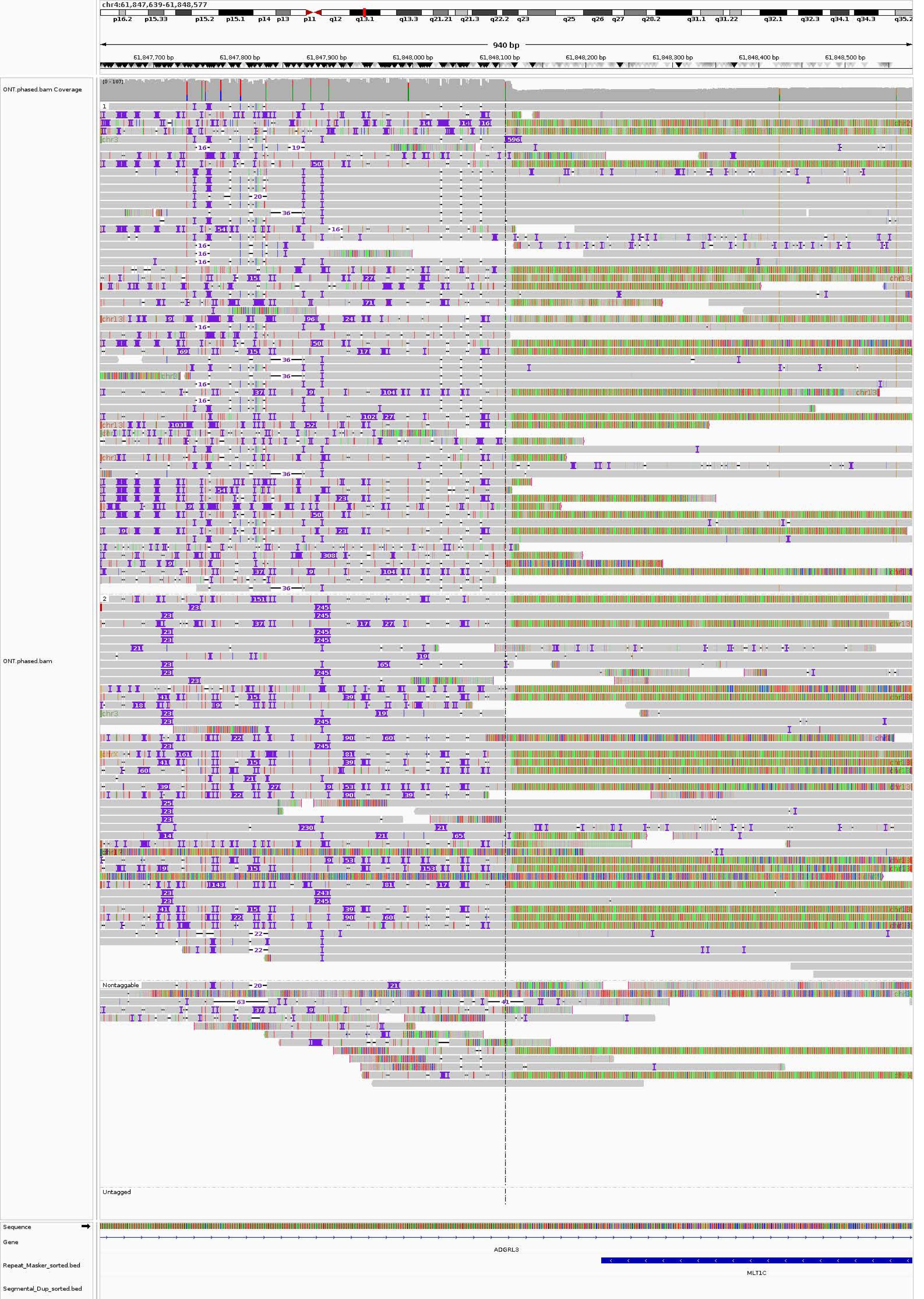
chr4:61,847,639-61,848,577
p16.2 p15.33 p15.2 p15.1 p14 p13 p11 q12 q13.1 q13.3 q21.21 q21.3 q22.2 q23 q25 q26 q27 q28.2 q31.1 q31.22 q32.1 q32.3 q34.1 q34.3 q35.2
940 bp
61,847,700 bp
61,847,800 bp
61,847,900 bp
61,848,000 bp
61,848,100 bp
61,848,200 bp
61,848,300 bp
61,848,400 bp
61,848,500 bp
ONT.phased.bam Coverage
[0 - 107]
ONT.phased.bam
1
2
Nontaggable
Untagged
Sequence
Gene
ADGRL3
Repeat_Masker_sorted.bed
MLT1C
Segmental_Dup_sorted.bed

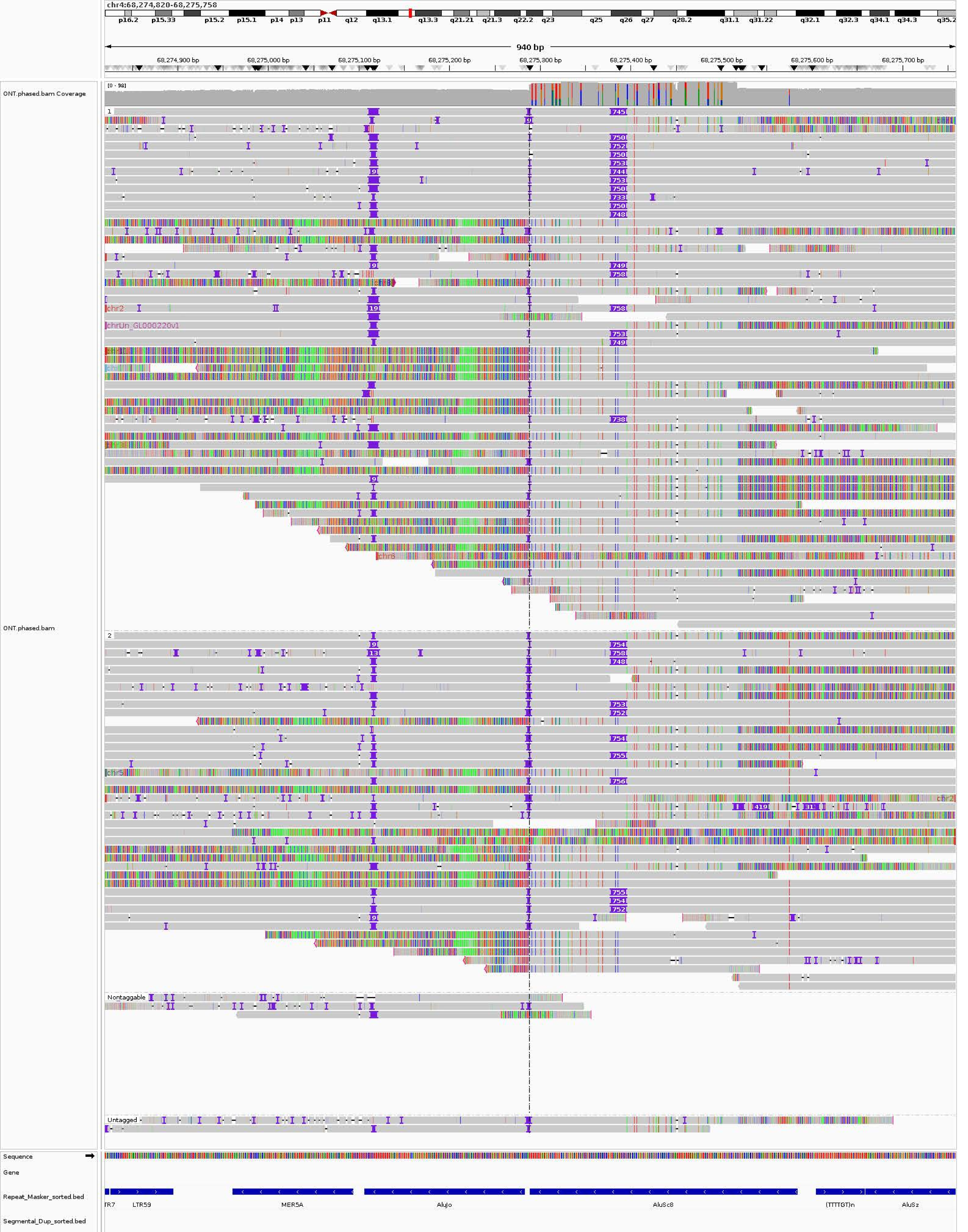
chr4:68,274,820-68,275,758
p16.2 p15.33 p15.2 p15.1 p14 p13 p11 q12 q13.1 q13.3 q21.21 q21.3 q22.2 q23 q25 q26 q27 q28.2 q31.1 q31.22 q32.1 q32.3 q34.1 q34.3 q35.2
940 bp
68,274,900 bp
68,275,000 bp
68,275,100 bp
68,275,200 bp
68,275,300 bp
68,275,400 bp
68,275,500 bp
68,275,600 bp
68,275,700 bp
ONT.phased.bam Coverage
[0 - 98]
ONT.phased.bam
1
2
Nontaggable
Untagged
chr2
chrUn_GL000220v1
chr6
chr5
Sequence
Gene
Repeat_Masker_sorted.bed
Segmental_Dup_sorted.bed
TR7
LTR59
MER5A
AluJo
AluSc8
(TTTTGT)n
AluSz

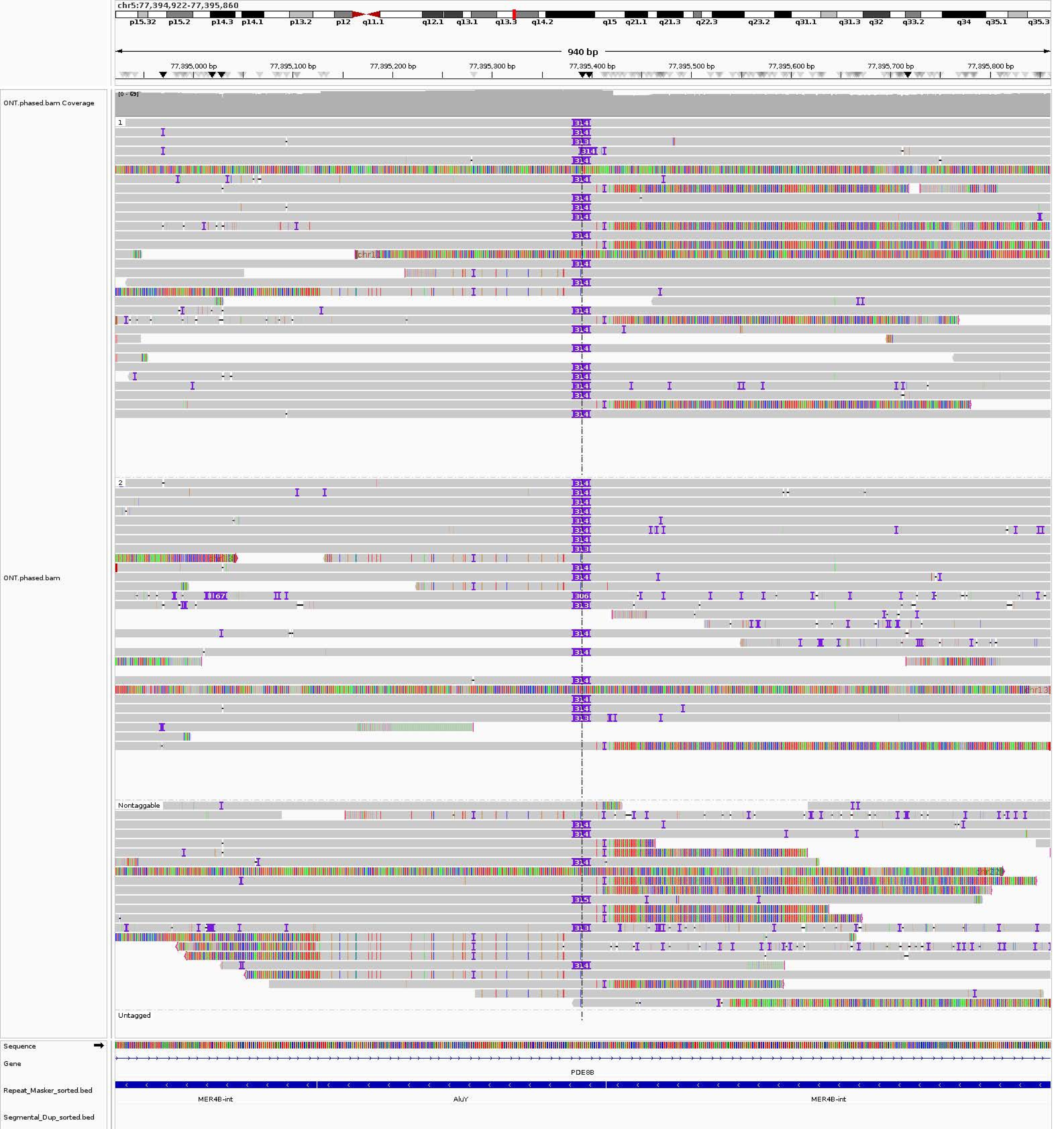
chr5:77,394,922-77,395,860
p15.32
p15.2
p14.3
p14.1
p13.2
p12
q11.1
q12.1
q13.1
q13.3
q14.2
q15
q21.1
q21.3
q22.3
q23.2
q31.1
q31.3
q32
q33.2
q34
q35.1
q35.3
940 bp
77,395,000 bp
77,395,100 bp
77,395,200 bp
77,395,300 bp
77,395,400 bp
77,395,500 bp
77,395,600 bp
77,395,700 bp
77,395,800 bp
ONT.phased.bam Coverage
[0 - 69]
ONT.phased.bam
1
2
Nontaggable
Untagged
Sequence
Gene
PDE8B
Repeat_Masker_sorted.bed
MER4B-int
AluY
MER4B-int
Segmental_Dup_sorted.bed

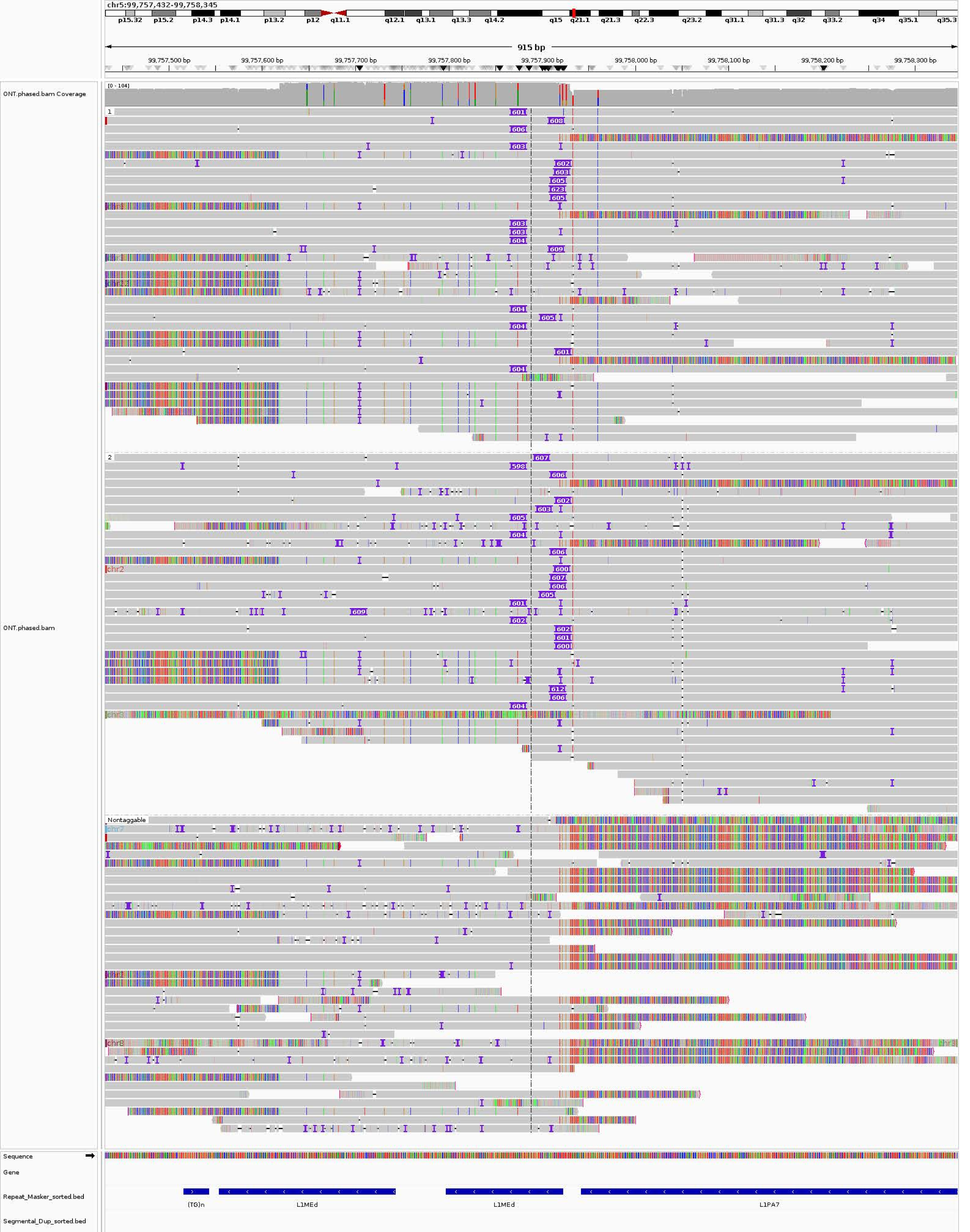

chr5:99,757,432-99,758,345
p15.32
p15.2
p14.3
p14.1
p13.2
p12
q11.1
q12.1
q13.1
q13.3
q14.2
q15
q21.1
q21.3
q22.3
q23.2
q31.1
q31.3
q32
q33.2
q34
q35.1
q35.3
915 bp
99,757,500 bp
99,757,600 bp
99,757,700 bp
99,757,800 bp
99,757,900 bp
99,758,000 bp
99,758,100 bp
99,758,200 bp
99,758,300 bp
ONT.phased.bam Coverage
[0 - 104]
ONT.phased.bam
1
2
Nontaggable
Sequence
Gene
Repeat_Masker_sorted.bed
(TG)n
L1MEd
L1MEd
L1PA7
Segmental_Dup_sorted.bed

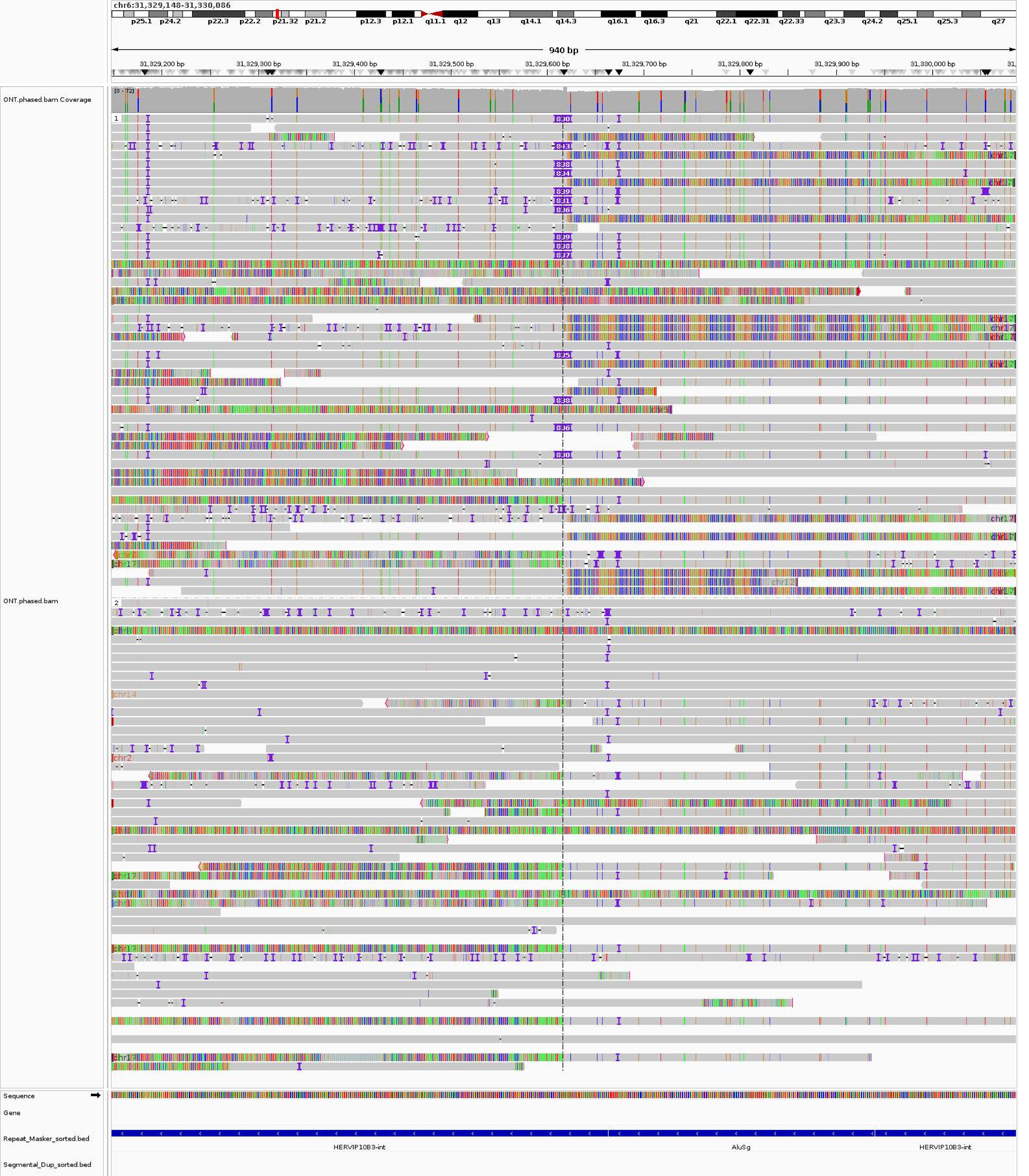
chr6:31,329,148-31,330,086
p25.1
p24.2
p22.3
p22.2
p21.32
p21.2
p12.3
p12.1
q11.1
q12
q13
q14.1
q14.3
q16.1
q16.3
q21
q22.1
q22.31
q22.33
q23.3
q24.2
q25.1
q25.3
q27
940 bp
31,329,200 bp
31,329,300 bp
31,329,400 bp
31,329,500 bp
31,329,600 bp
31,329,700 bp
31,329,800 bp
31,329,900 bp
31,330,000 bp
ONT.phased.bam Coverage
[0 - 72]
ONT.phased.bam
Sequence
Gene
Repeat_Masker_sorted.bed
HERVIP10B3-int
AluSg
HERVIP10B3-int
Segmental_Dup_sorted.bed

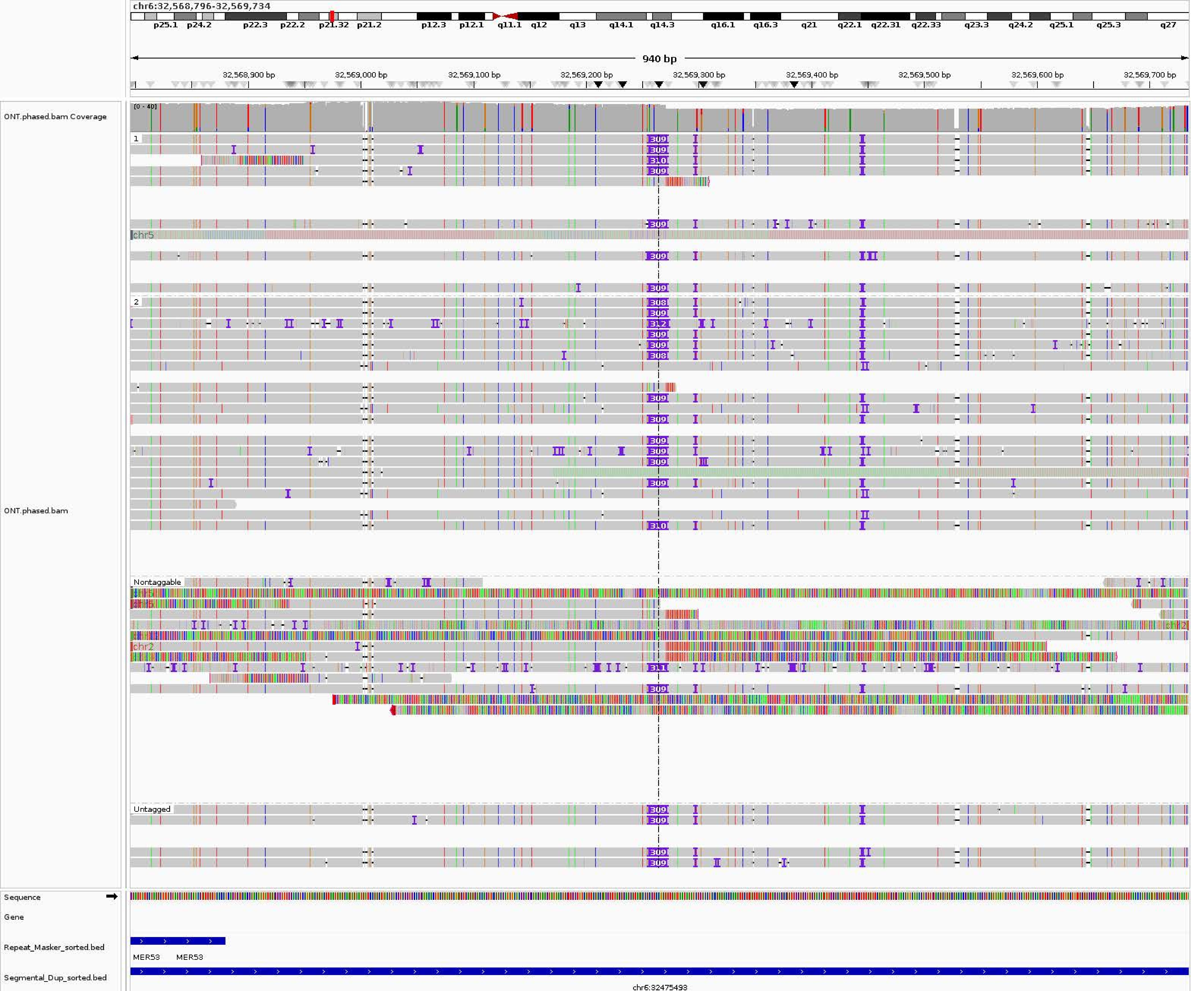

chr6:32,568,796-32,569,734
p25.1 p24.2 p22.3 p22.2 p21.32 p21.2 p12.3 p12.1 q11.1 q12 q13 q14.1 q14.3 q16.1 q16.3 q21 q22.1 q22.31 q22.33 q23.3 q24.2 q25.1 q25.3 q27
940 bp
32,568,900 bp
32,569,000 bp
32,569,100 bp
32,569,200 bp
32,569,300 bp
32,569,400 bp
32,569,500 bp
32,569,600 bp
32,569,700 bp
ONT.phased.bam Coverage
[0 - 40]
ONT.phased.bam
1
2
Nontaggable
Untagged
chr5
chr2
309
310
308
312
311
Sequence
Gene
Repeat_Masker_sorted.bed
MER53
MER53
Segmental_Dup_sorted.bed
chr6:32475493

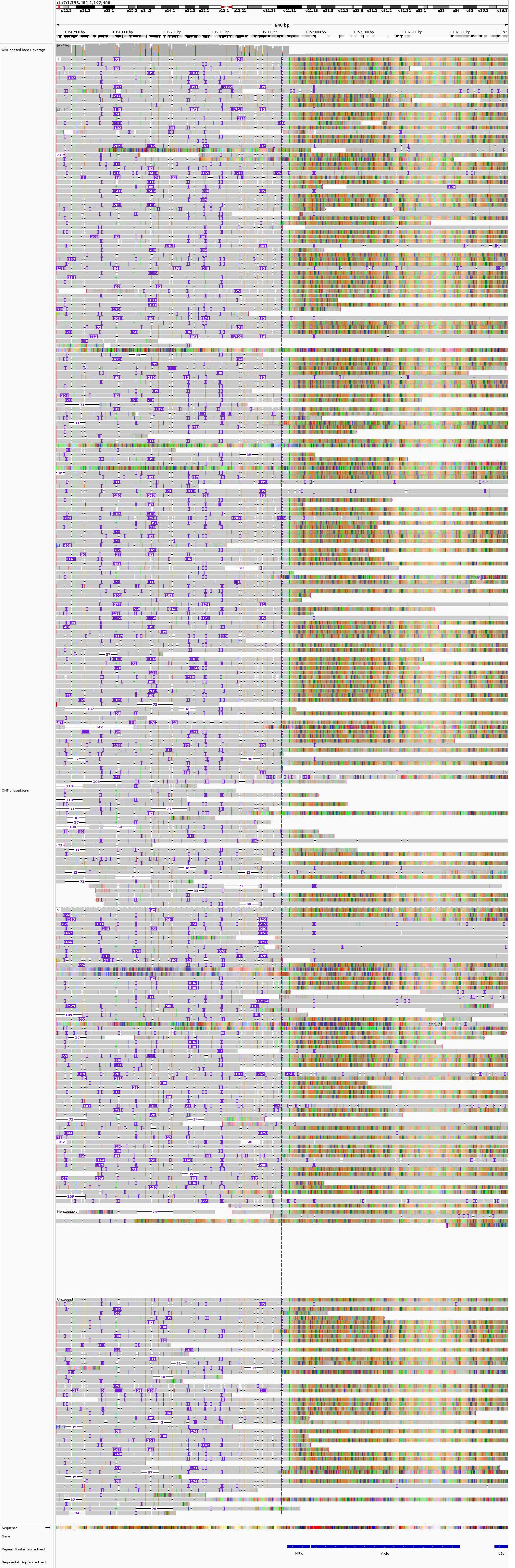

chr7:1,196,462-1,197,400
p22.2
p21.3
p21.1
p15.2
p14.3
p14.1
p12.3
p12.1
p11.1
q11.21
q11.23
q21.11
q21.13
q21.3
q22.1
q22.3
q31.1
q31.2
q31.32
q32.1
q33
q34
q35
q36.1
q36.3
940 bp
1,196,500 bp
1,196,600 bp
1,196,700 bp
1,196,800 bp
1,196,900 bp
1,197,000 bp
1,197,100 bp
1,197,200 bp
1,197,300 bp
ONT.phased.bam Coverage
ONT.phased.bam
Nontaggable
Untagged
Sequence
Gene
Repeat_Masker_sorted.bed
Segmental_Dup_sorted.bed
MIRc
AluJo
L2a

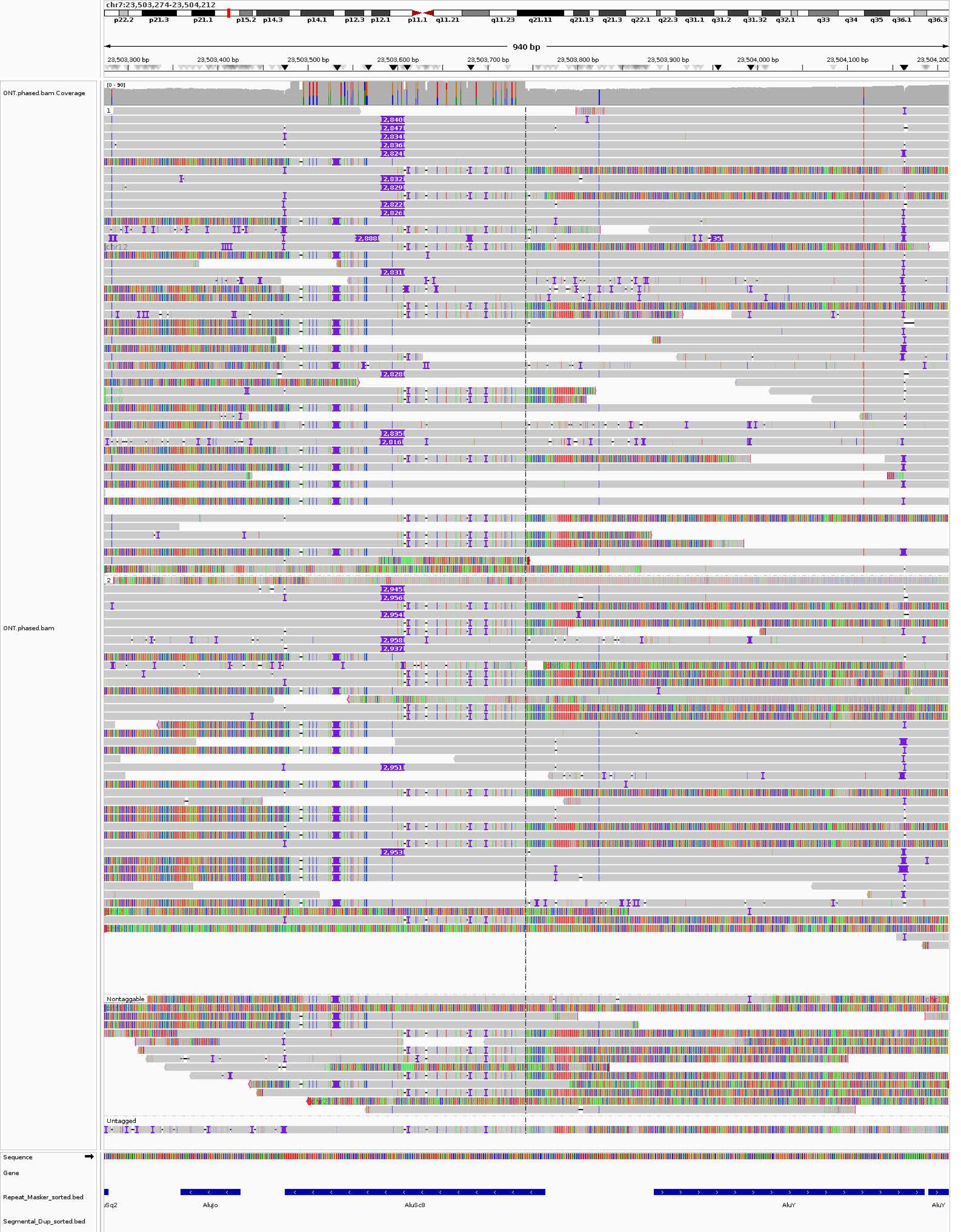
chr7:23,503,274-23,504,212
p22.2
p21.3
p21.1
p15.2
p14.3
p14.1
p12.3
p12.1
p11.1
q11.21
q11.23
q21.11
q21.13
q21.3
q22.1
q22.3
q31.1
q31.2
q31.32
q32.1
q33
q34
q35
q36.1
q36.3
940 bp
23,503,300 bp
23,503,400 bp
23,503,500 bp
23,503,600 bp
23,503,700 bp
23,503,800 bp
23,503,900 bp
23,504,000 bp
23,504,100 bp
ONT.phased.bam Coverage
[0 - 90]
ONT.phased.bam
1
2
Nontaggable
Untagged
Sequence
Gene
Repeat_Masker_sorted.bed
Segmental_Dup_sorted.bed
AluJo
AluSc8
AluY

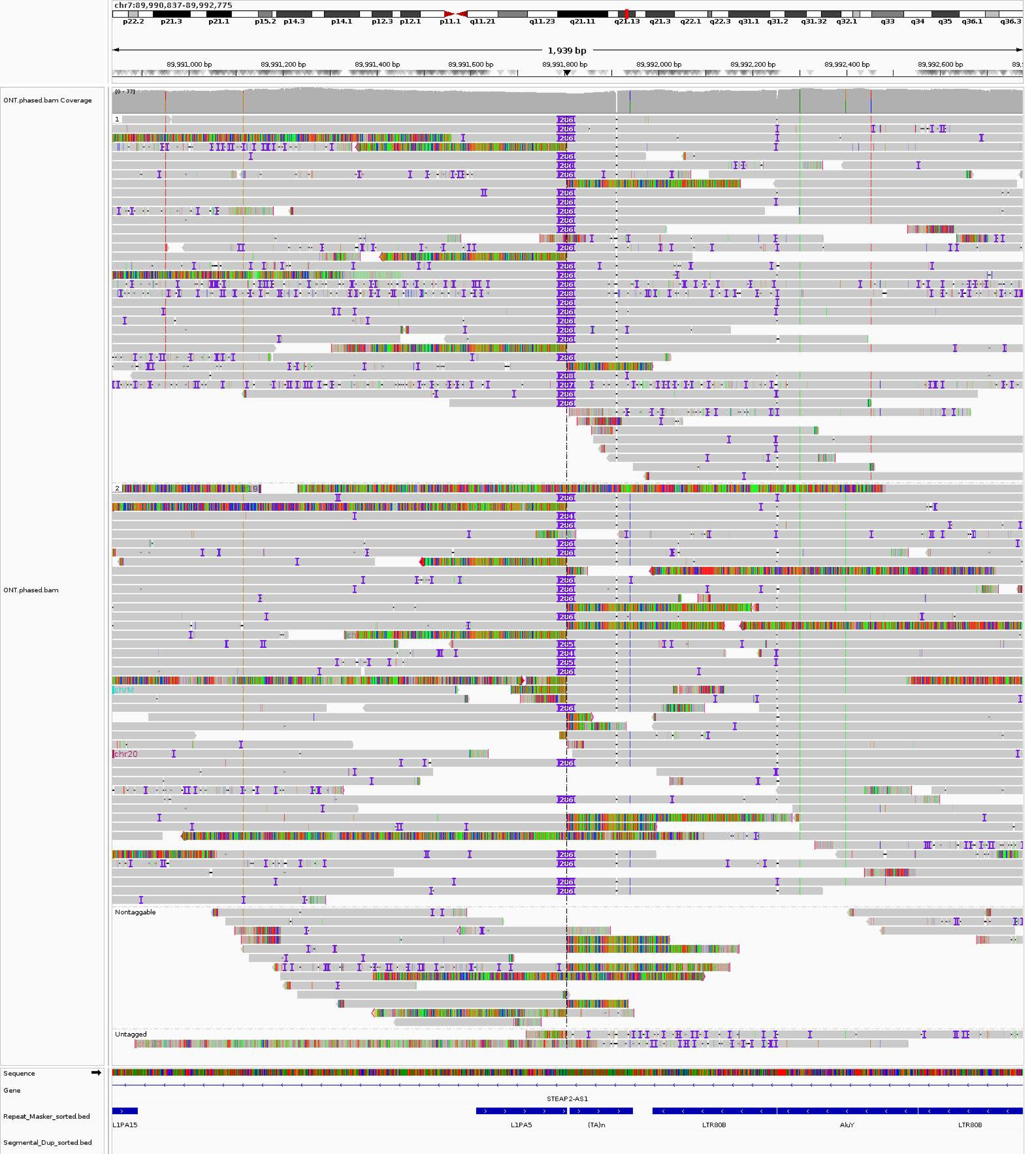

chr7:89,990,837-89,992,775
p22.2
p21.3
p21.1
p15.2
p14.3
p14.1
p12.3
p12.1
p11.1
q11.21
q11.23
q21.11
q21.13
q21.3
q22.1
q22.3
q31.1
q31.2
q31.32
q32.1
q33
q34
q35
q36.1
q36.3
1,939 bp
89,991,000 bp
89,991,200 bp
89,991,400 bp
89,991,600 bp
89,991,800 bp
89,992,000 bp
89,992,200 bp
89,992,400 bp
89,992,600 bp
ONT.phased.bam Coverage
[0 - 77]
ONT.phased.bam
1
2
chrM
chr20
Nontaggable
Untagged
Sequence
Gene
STEAP2-AS1
Repeat_Masker_sorted.bed
L1PA15
L1PA5
(TA)n
LTR80B
AluY
LTR80B
Segmental_Dup_sorted.bed

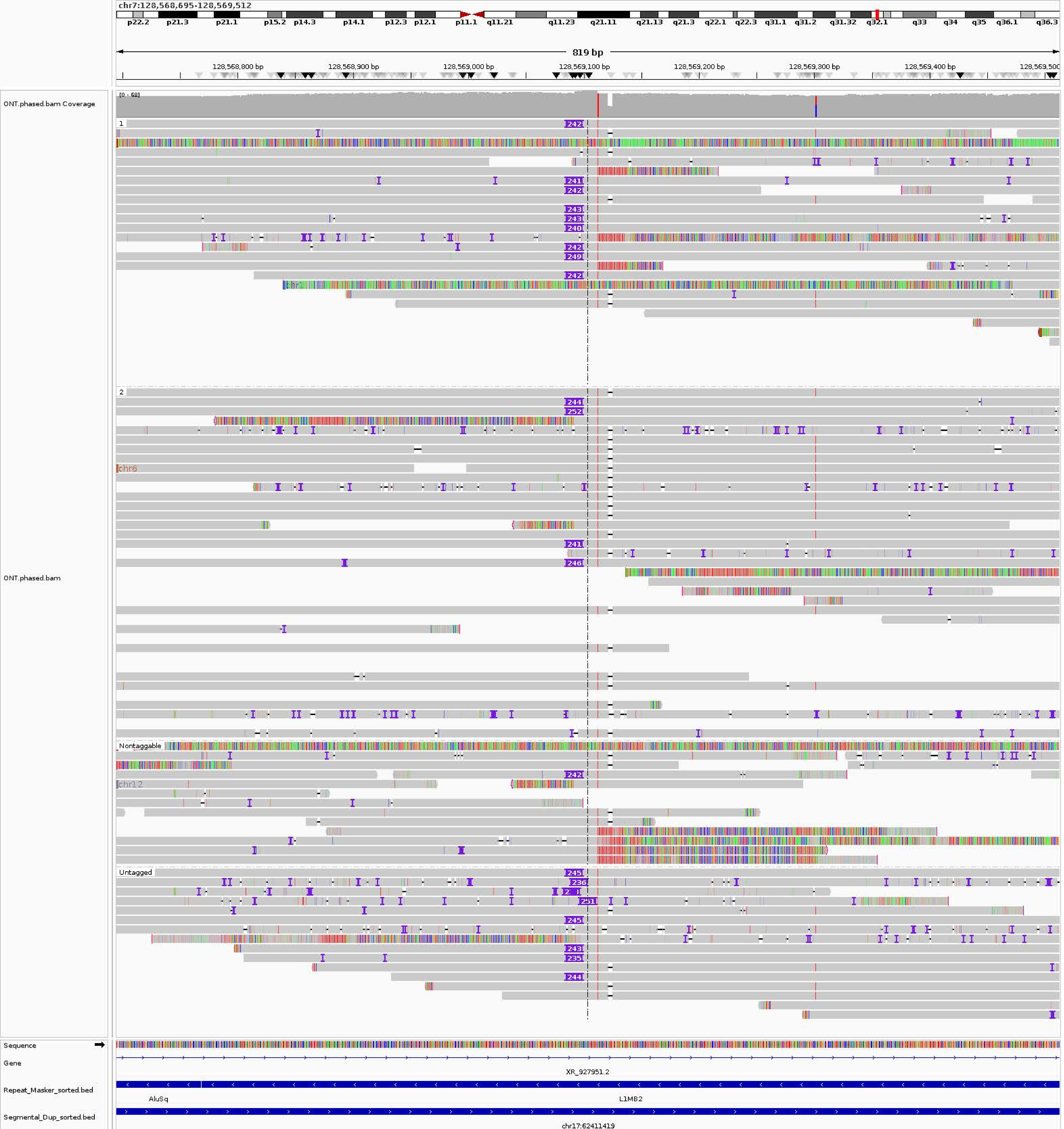
chr7:128,568,695-128,569,512
p22.2 p21.3 p21.1 p15.2 p14.3 p14.1 p12.3 p12.1 p11.1 q11.21 q11.23 q21.11 q21.13 q21.3 q22.1 q22.3 q31.1 q31.2 q31.32 q32.1 q33 q34 q35 q36.1 q36.3
819 bp
128,568,800 bp
128,568,900 bp
128,569,000 bp
128,569,100 bp
128,569,200 bp
128,569,300 bp
128,569,400 bp
128,569,500
ONT.phased.bam Coverage
[0 - 68]
ONT.phased.bam
1
2
chr6
Nontaggable
chr12
Untagged
Sequence
Gene
XR_927951.2
Repeat_Masker_sorted.bed
AluSq
L1MB2
Segmental_Dup_sorted.bed
chr17:62411419

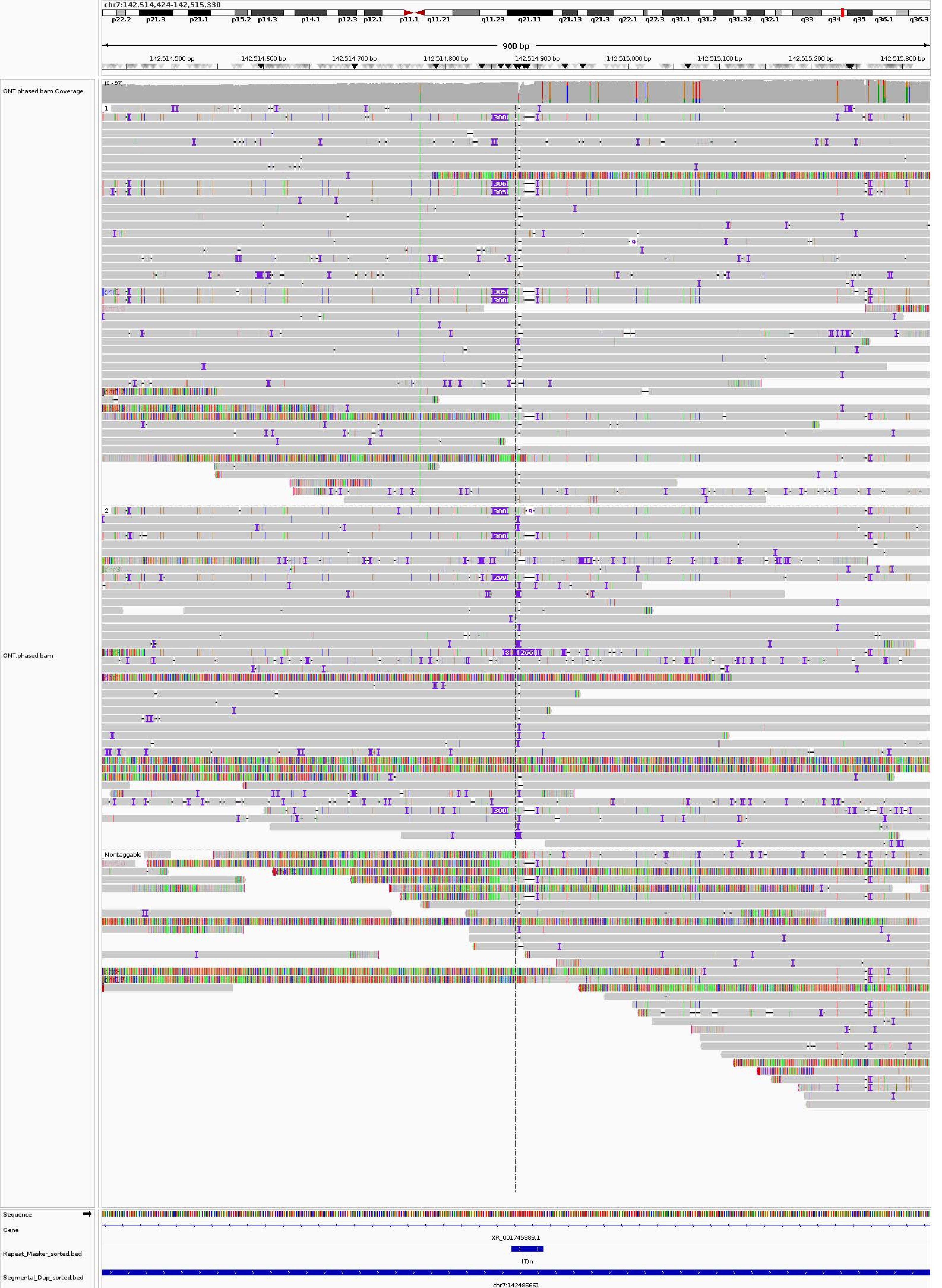
chr7:142,514,424-142,515,330
p22.2 p21.3 p21.1 p15.2 p14.3 p14.1 p12.3 p12.1 p11.1 q11.21 q11.23 q21.11 q21.13 q21.3 q22.1 q22.3 q31.1 q31.2 q31.32 q32.1 q33 q34 q35 q36.1 q36.3
908 bp
142,514,500 bp
142,514,600 bp
142,514,700 bp
142,514,800 bp
142,514,900 bp
142,515,000 bp
142,515,100 bp
142,515,200 bp
142,515,300 bp
ONT.phased.bam Coverage
[0 - 97]
ONT.phased.bam
1
2
Nontaggable
Sequence
Gene
XR_001745389.1
Repeat_Masker_sorted.bed
(T)n
Segmental_Dup_sorted.bed
chr7:142486661

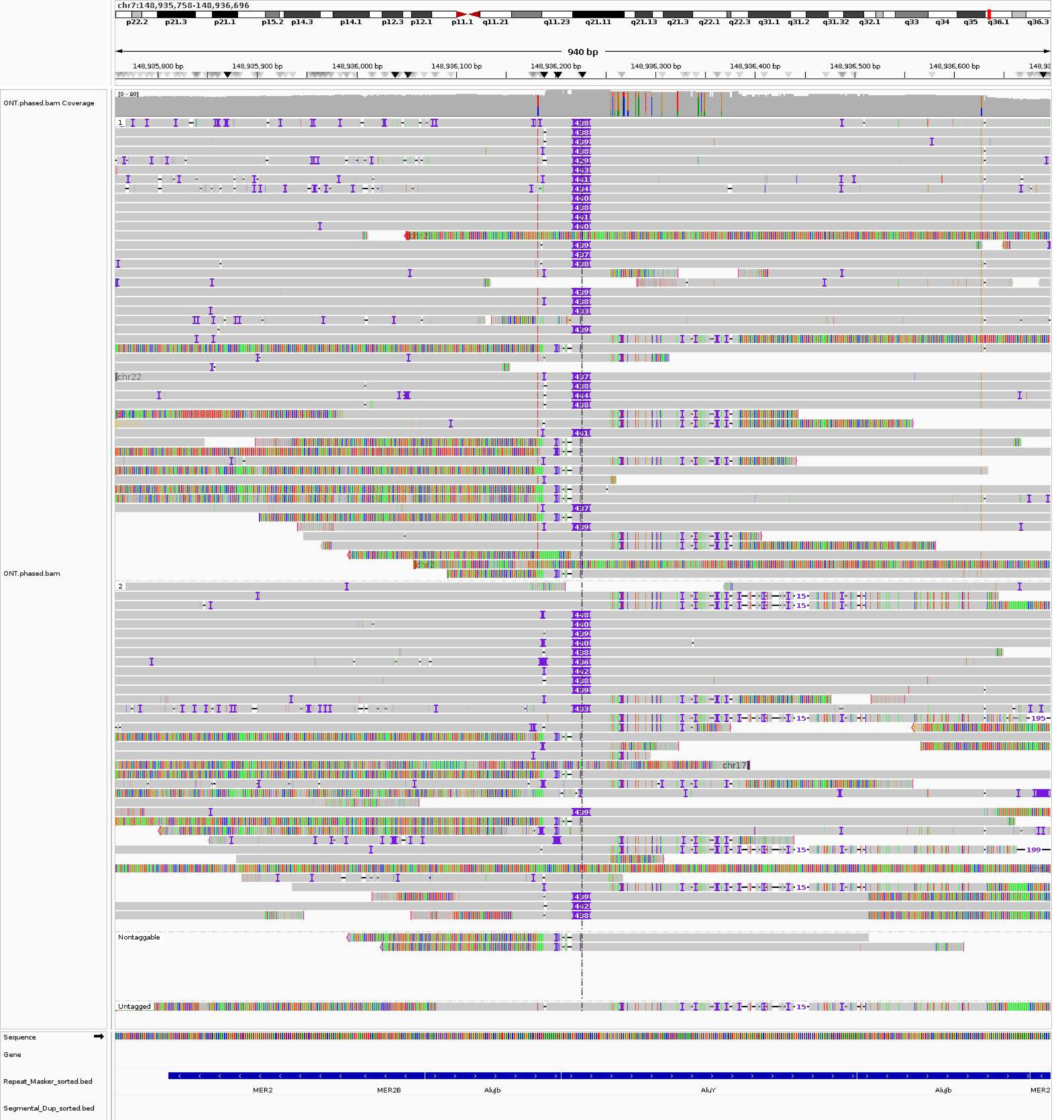
chr7:148,935,758-148,936,696
p22.2
p21.3
p21.1
p15.2
p14.3
p14.1
p12.3
p12.1
p11.1
q11.21
q11.23
q21.11
q21.13
q21.3
q22.1
q22.3
q31.1
q31.2
q31.32
q32.1
q33
q34
q35
q36.1
q36.3
940 bp
148,935,800 bp
148,935,900 bp
148,936,000 bp
148,936,100 bp
148,936,200 bp
148,936,300 bp
148,936,400 bp
148,936,500 bp
148,936,600 bp
ONT.phased.bam Coverage
[0 - 80]
ONT.phased.bam
chr22
chr17
Nontaggable
Untagged
Sequence
Gene
Repeat_Masker_sorted.bed
Segmental_Dup_sorted.bed
MER2
MER2B
AluJb
AluY
AluJb
MER2

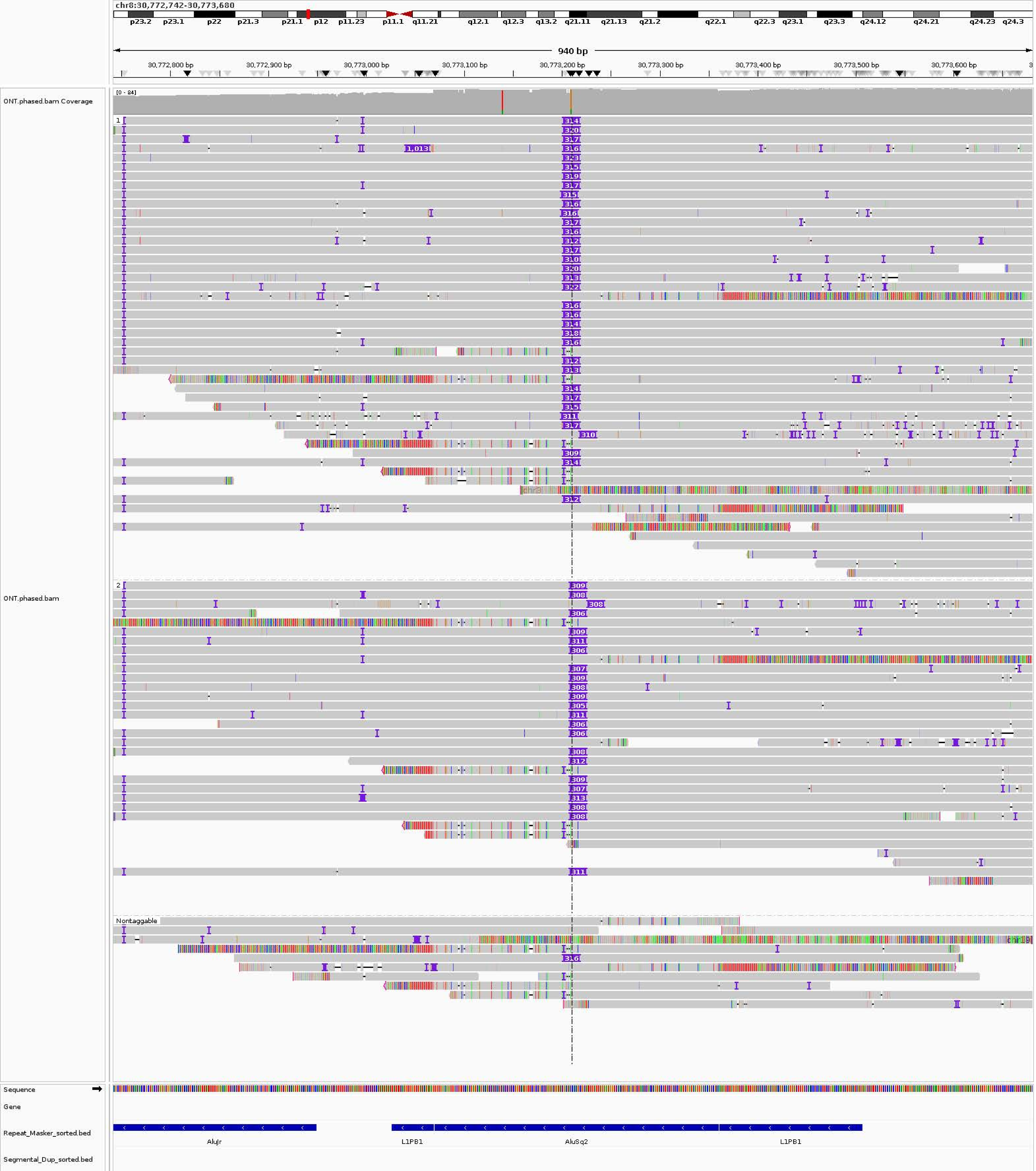
chr8:30,772,742-30,773,680
p23.2 p23.1 p22 p21.3 p21.1 p12 p11.23 p11.1 q11.21 q12.1 q12.3 q13.2 q21.11 q21.13 q21.2 q22.1 q22.3 q23.1 q23.3 q24.12 q24.21 q24.23 q24.3
940 bp
30,772,800 bp
30,772,900 bp
30,773,000 bp
30,773,100 bp
30,773,200 bp
30,773,300 bp
30,773,400 bp
30,773,500 bp
30,773,600 bp
ONT.phased.bam Coverage
[0 - 84]
ONT.phased.bam
1
2
Nontaggable
Sequence
Gene
Repeat_Masker_sorted.bed
Segmental_Dup_sorted.bed
AluJr
L1PB1
AluSq2
L1PB1

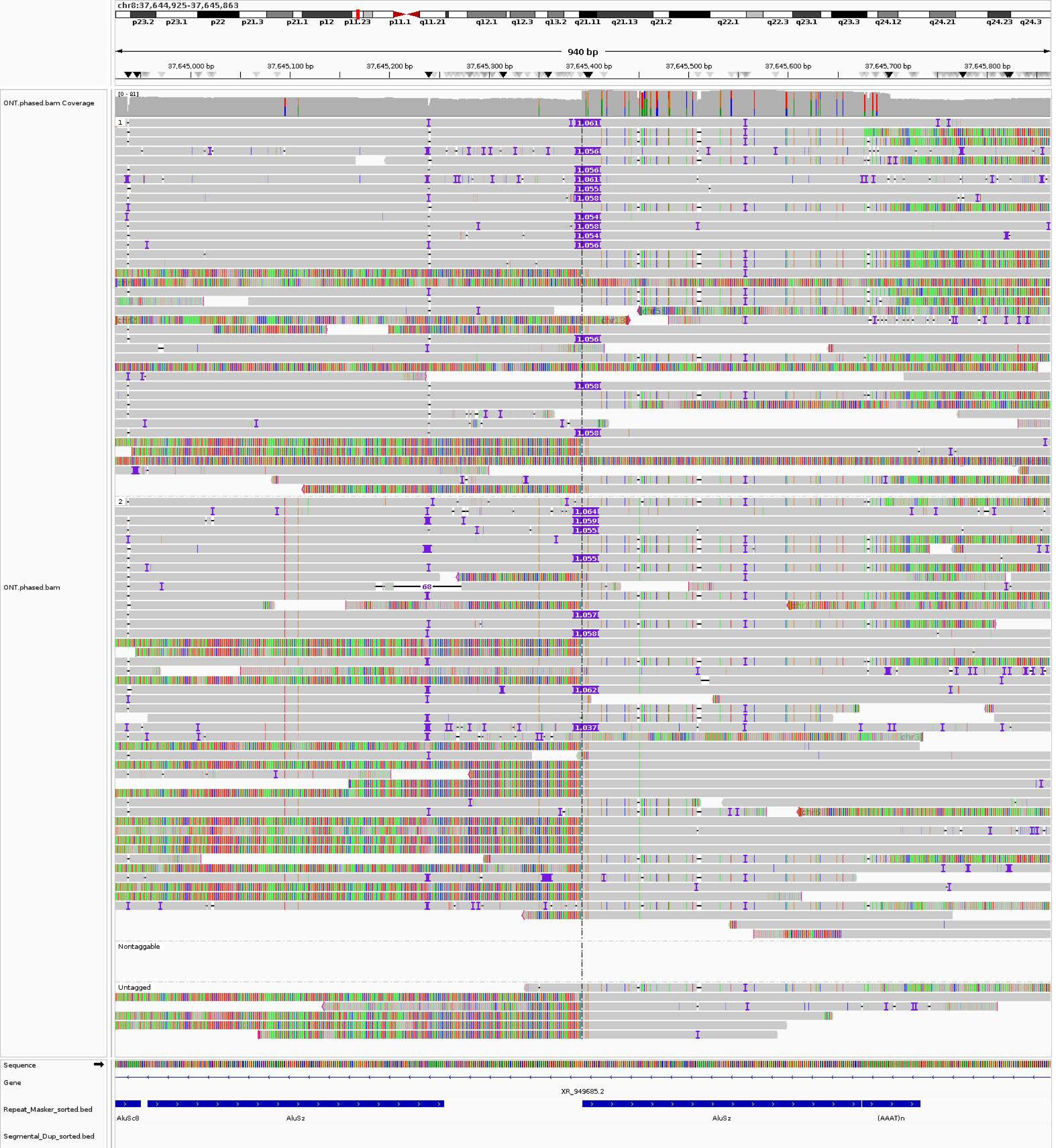
chr8:37,644,925-37,645,863
p23.2
p23.1
p22
p21.3
p21.1
p12
p11.23
p11.1
q11.21
q12.1
q12.3
q13.2
q21.11
q21.13
q21.2
q22.1
q22.3
q23.1
q23.3
q24.12
q24.21
q24.23
q24.3
940 bp
37,645,000 bp
37,645,100 bp
37,645,200 bp
37,645,300 bp
37,645,400 bp
37,645,500 bp
37,645,600 bp
37,645,700 bp
37,645,800 bp
ONT.phased.bam Coverage
[0 - 81]
ONT.phased.bam
Nontaggable
Untagged
Sequence
Gene
XR_949685.2
Repeat_Masker_sorted.bed
AluSc8
AluSz
AluSz
(AAAT)n
Segmental_Dup_sorted.bed

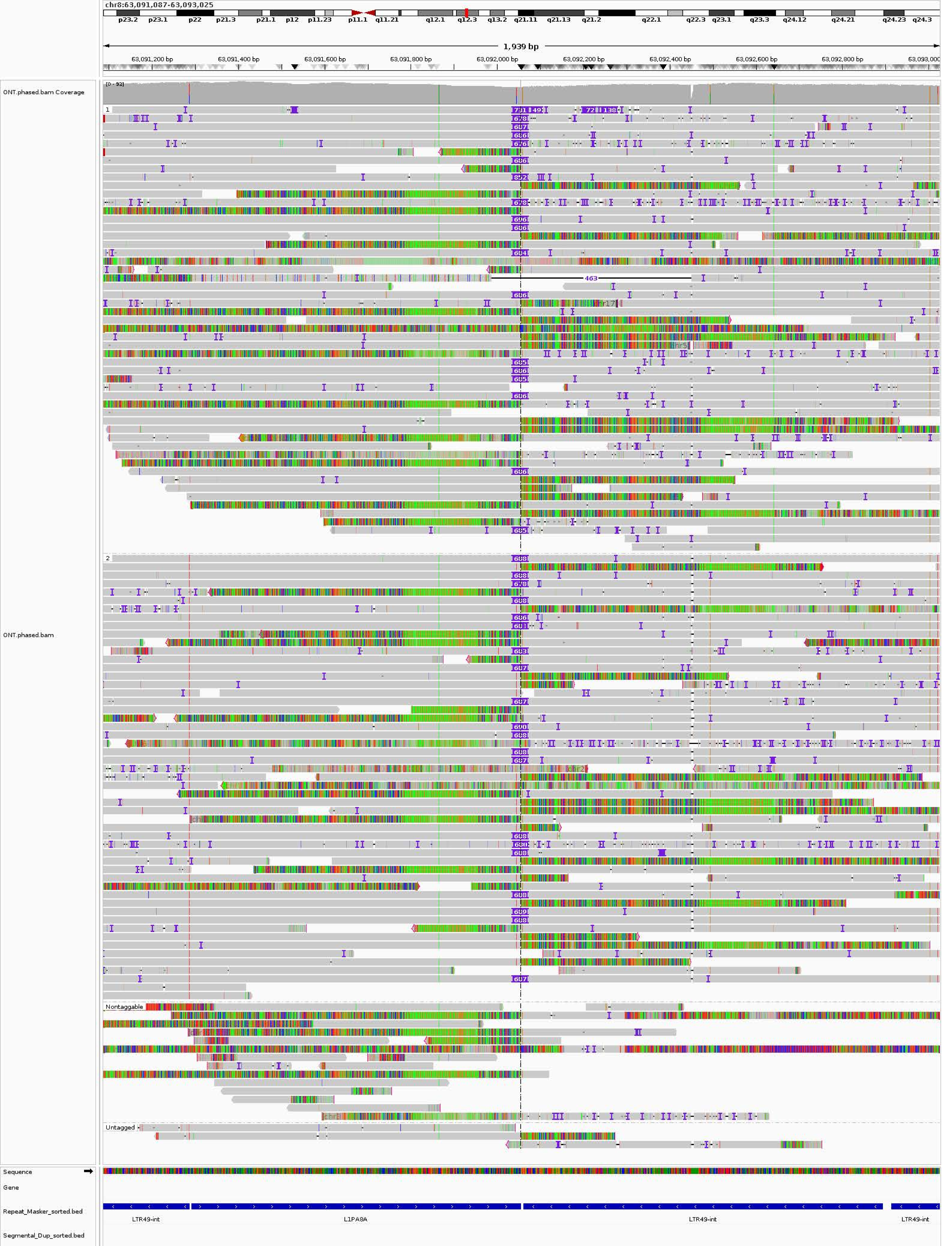
chr8:63,091,087-63,093,025
p23.2
p23.1
p22
p21.3
p21.1
p12
p11.23
p11.1
q11.21
q12.1
q12.3
q13.2
q21.11
q21.13
q21.2
q22.1
q22.3
q23.1
q23.3
q24.12
q24.21
q24.23
q24.3
1,939 bp
63,091,200 bp
63,091,400 bp
63,091,600 bp
63,091,800 bp
63,092,000 bp
63,092,200 bp
63,092,400 bp
63,092,600 bp
63,092,800 bp
63,093,000
ONT.phased.bam Coverage
[0 - 93]
ONT.phased.bam
1
2
Nontaggable
Untagged
463
Sequence
Gene
Repeat_Masker_sorted.bed
Segmental_Dup_sorted.bed
LTR49-int
L1PA8A
LTR49-int
LTR49-int

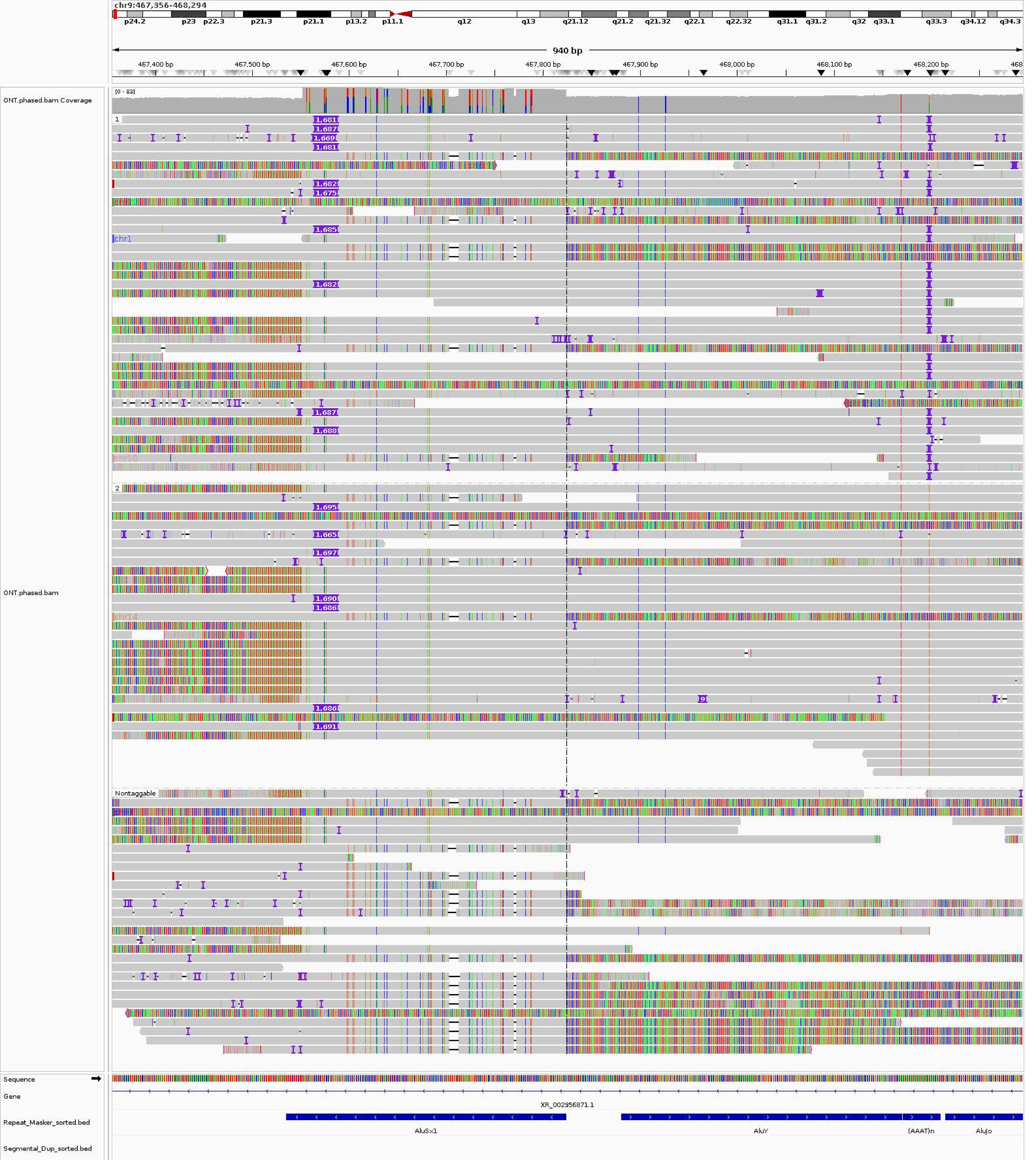
chr9:467,356-468,294
p24.2
p23
p22.3
p21.3
p21.1
p13.2
p11.1
q12
q13
q21.12
q21.2
q21.32
q22.1
q22.32
q31.1
q31.2
q32
q33.1
q33.3
q34.12
q34.3
940 bp
467,400 bp
467,500 bp
467,600 bp
467,700 bp
467,800 bp
467,900 bp
468,000 bp
468,100 bp
468,200 bp
ONT.phased.bam Coverage
[0 - 83]
1
2
Nontaggable
ONT.phased.bam
Sequence
Gene
XR_002956871.1
Repeat_Masker_sorted.bed
AluSx1
AluY
(AAAT)n
AluJo
Segmental_Dup_sorted.bed

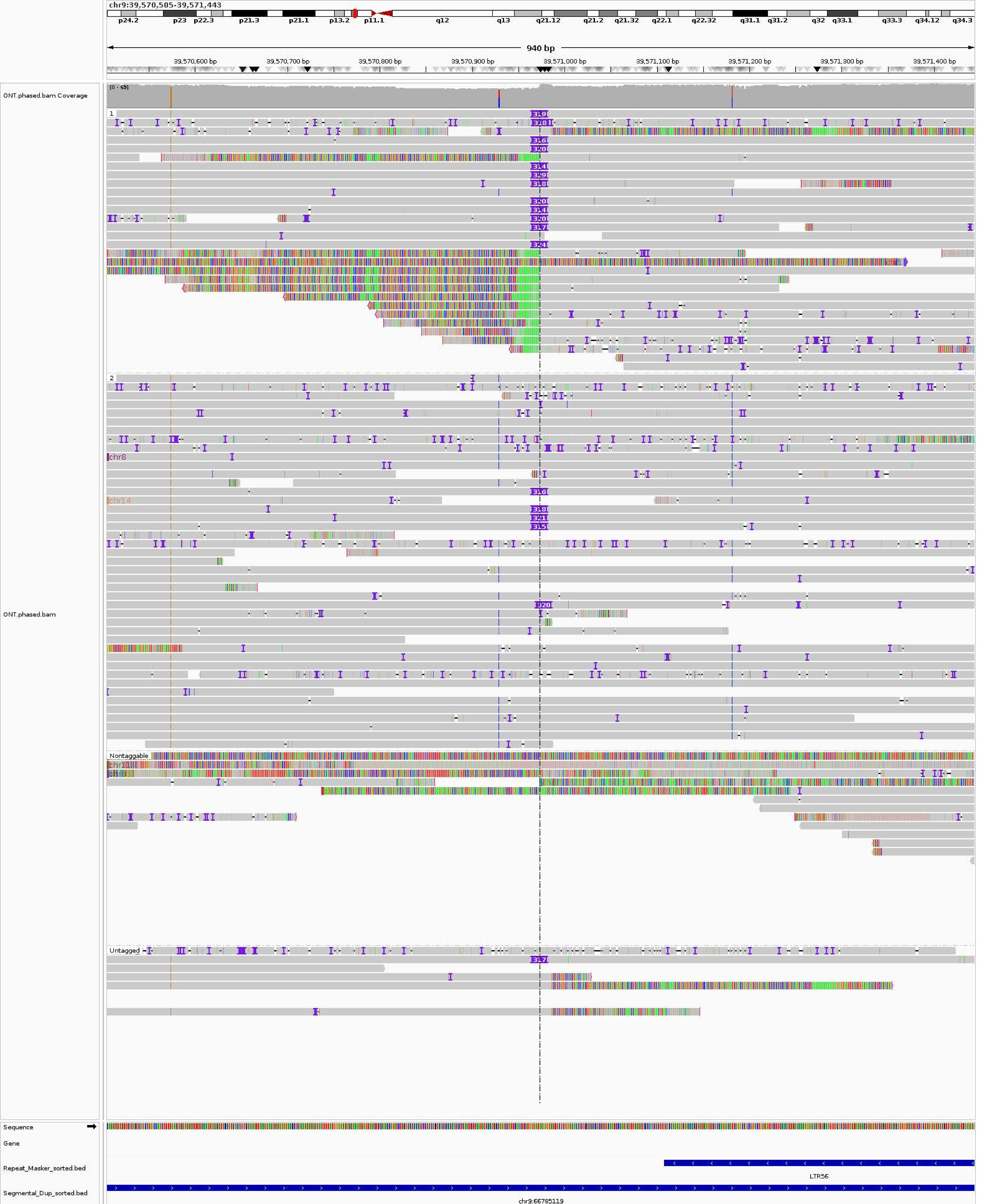
chr9:39,570,505-39,571,443
p24.2
p23
p22.3
p21.3
p21.1
p13.2
p11.1
q12
q13
q21.12
q21.2
q21.32
q22.1
q22.32
q31.1
q31.2
q32
q33.1
q33.3
q34.12
q34.3
940 bp
39,570,600 bp
39,570,700 bp
39,570,800 bp
39,570,900 bp
39,571,000 bp
39,571,100 bp
39,571,200 bp
39,571,300 bp
39,571,400 bp
ONT.phased.bam Coverage
[0 - 69]
ONT.phased.bam
1
2
chr8
chr14
Nontaggable
Untagged
Sequence
Gene
Repeat_Masker_sorted.bed
LTR56
Segmental_Dup_sorted.bed
chr9:66785119

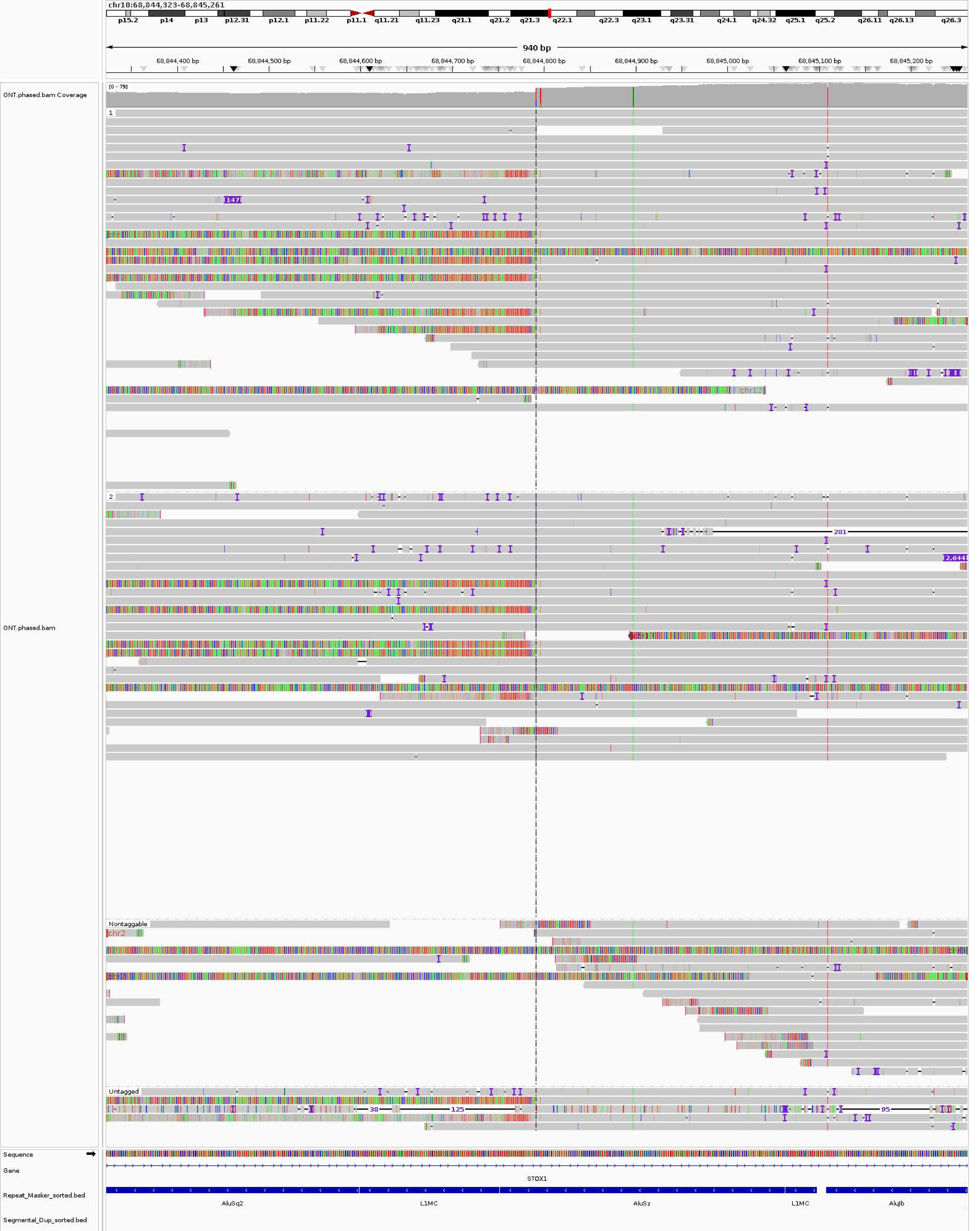
chr10:68,844,323-68,845,261
p15.2
p14
p13
p12.31
p12.1
p11.22
p11.1
q11.21
q11.23
q21.1
q21.2
q21.3
q22.1
q22.3
q23.1
q23.31
q24.1
q24.32
q25.1
q25.2
q26.11
q26.13
q26.3
940 bp
68,844,400 bp
68,844,500 bp
68,844,600 bp
68,844,700 bp
68,844,800 bp
68,844,900 bp
68,845,000 bp
68,845,100 bp
68,845,200 bp
ONT.phased.bam Coverage
[0 - 79]
1
147
chr12
2
281
2,644
ONT.phased.bam
Nontaggable
chr2
Untagged
38
125
95
Sequence
Gene
STOX1
Repeat_Masker_sorted.bed
AluSq2
L1MC
AluSz
L1MC
AluJb
Segmental_Dup_sorted.bed

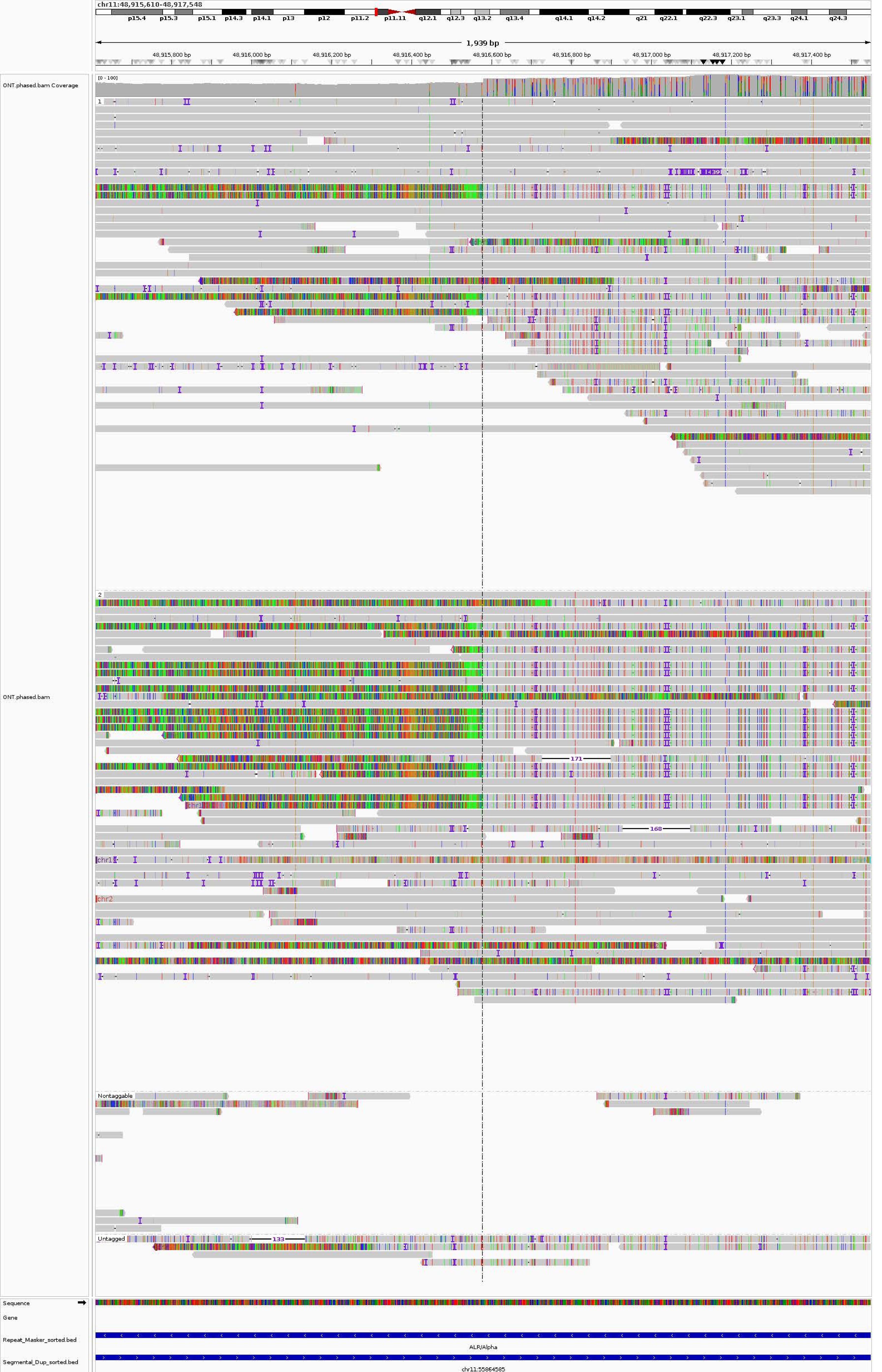

chr11:48,915,610-48,917,548
p15.4
p15.3
p15.1
p14.3
p14.1
p13
p12
p11.2
p11.11
q12.1
q12.3
q13.2
q13.4
q14.1
q14.2
q21
q22.1
q22.3
q23.1
q23.3
q24.1
q24.3
1,939 bp
48,915,800 bp
48,916,000 bp
48,916,200 bp
48,916,400 bp
48,916,600 bp
48,916,800 bp
48,917,000 bp
48,917,200 bp
48,917,400 bp
ONT.phased.bam Coverage
[0 - 100]
1
1439
2
171
168
chr1
chr2
Nontaggable
Untagged
133
ONT.phased.bam
Sequence
Gene
Repeat_Masker_sorted.bed
ALR/Alpha
Segmental_Dup_sorted.bed
chr11:55864585

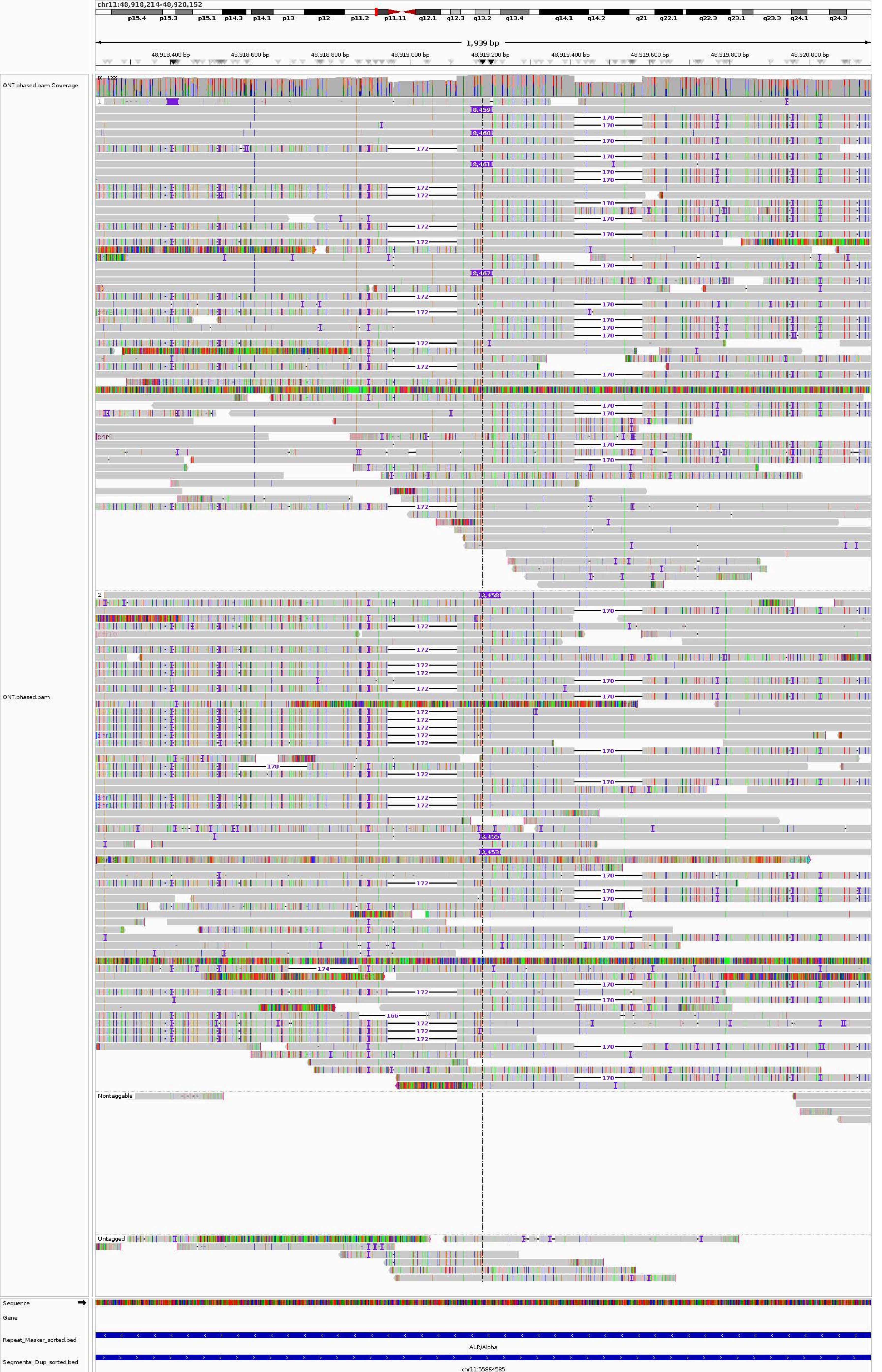
chr11:48,918,214-48,920,152
p15.4
p15.3
p15.1
p14.3
p14.1
p13
p12
p11.2
p11.11
q12.1
q12.3
q13.2
q13.4
q14.1
q14.2
q21
q22.1
q22.3
q23.1
q23.3
q24.1
q24.3
1,939 bp
48,918,400 bp
48,918,600 bp
48,918,800 bp
48,919,000 bp
48,919,200 bp
48,919,400 bp
48,919,600 bp
48,919,800 bp
48,920,000 bp
ONT.phased.bam Coverage
[0 - 122]
ONT.phased.bam
1
2
Nontaggable
Untagged
8,459
8,460
8,461
8,462
8,458
8,455
8,453
170
172
174
166
Sequence
Gene
Repeat_Masker_sorted.bed
ALR/Alpha
Segmental_Dup_sorted.bed
chr11:55864585

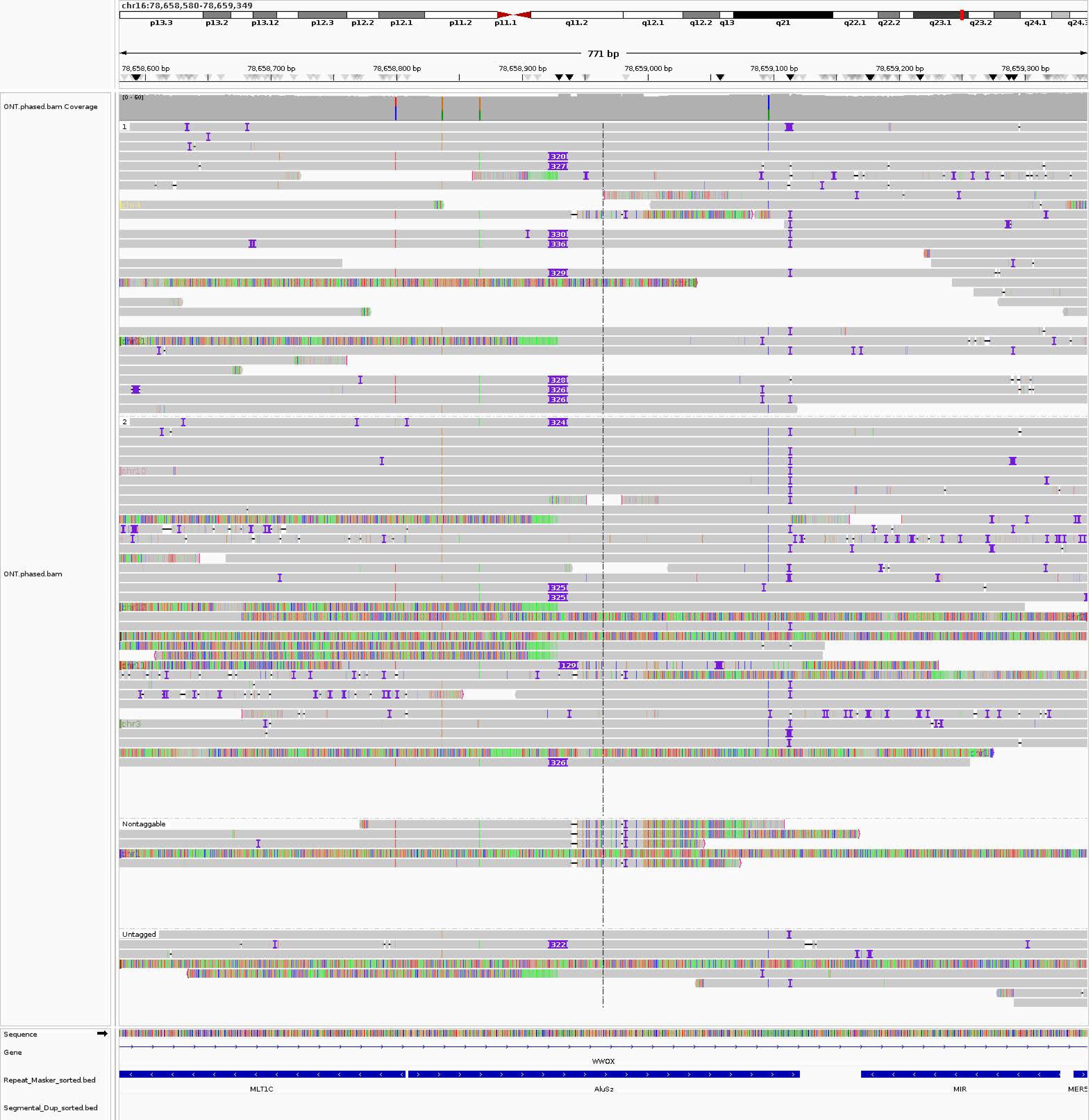
chr16:78,658,580-78,659,349
p13.3
p13.2
p13.12
p12.3
p12.2
p12.1
p11.2
p11.1
q11.2
q12.1
q12.2
q13
q21
q22.1
q22.2
q23.1
q23.2
q24.1
771 bp
78,658,600 bp
78,658,700 bp
78,658,800 bp
78,658,900 bp
78,659,000 bp
78,659,100 bp
78,659,200 bp
78,659,300 bp
ONT.phased.bam Coverage
[0 - 60]
ONT.phased.bam
1
2
Nontaggable
Untagged
Sequence
Gene
WWOX
Repeat_Masker_sorted.bed
MLT1C
AluSz
MIR
Segmental_Dup_sorted.bed

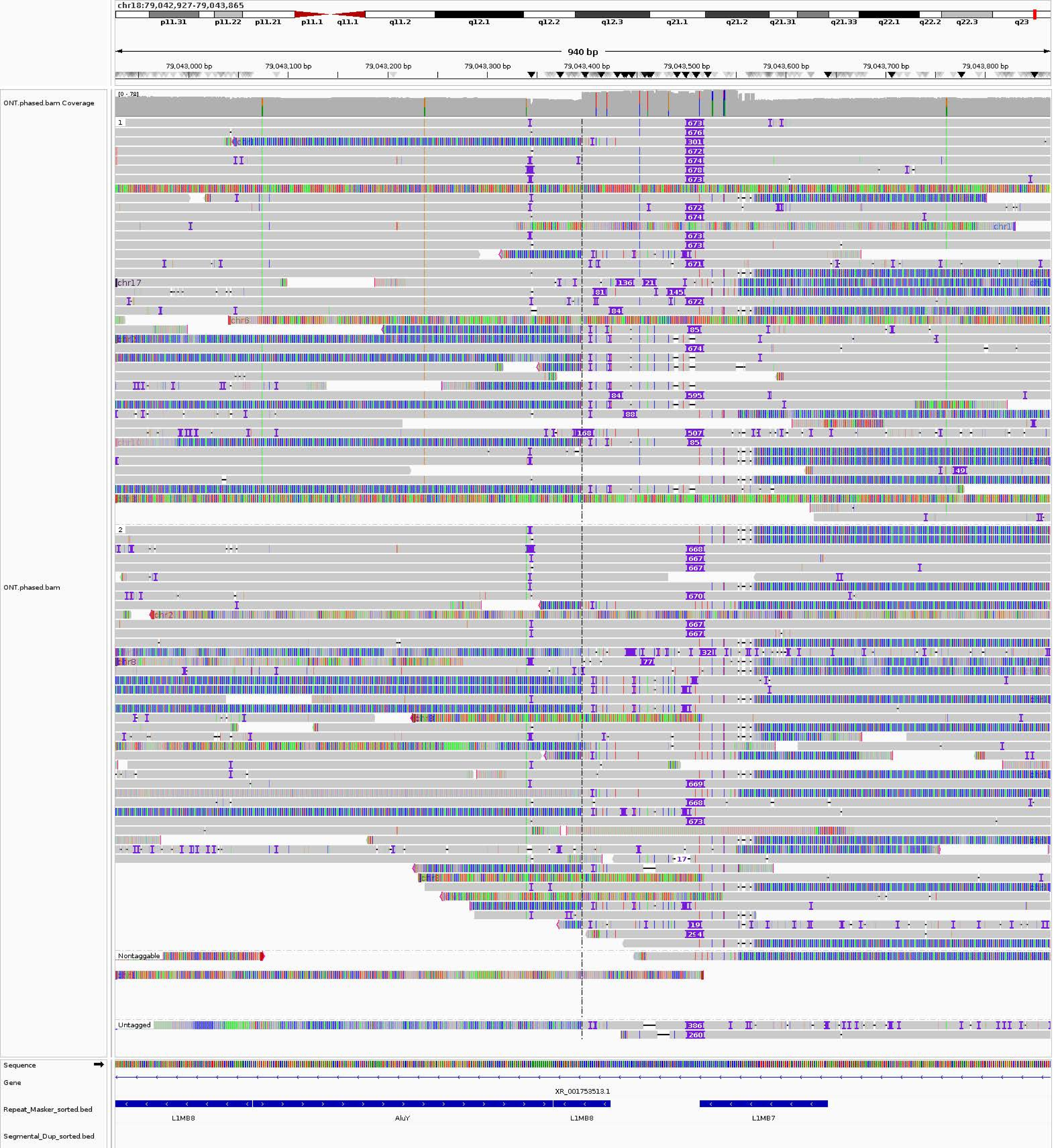
chr18:79,042,927-79,043,865
p11.31
p11.22
p11.21
p11.1
q11.1
q11.2
q12.1
q12.2
q12.3
q21.1
q21.2
q21.31
q21.33
q22.1
q22.2
q22.3
q23
940 bp
79,043,000 bp
79,043,100 bp
79,043,200 bp
79,043,300 bp
79,043,400 bp
79,043,500 bp
79,043,600 bp
79,043,700 bp
79,043,800 bp
ONT.phased.bam Coverage
[0 - 78]
ONT.phased.bam
Nontaggable
Untagged
Sequence
Gene
XR_001753513.1
Repeat_Masker_sorted.bed
L1MB8
AluY
L1MB8
L1MB7
Segmental_Dup_sorted.bed

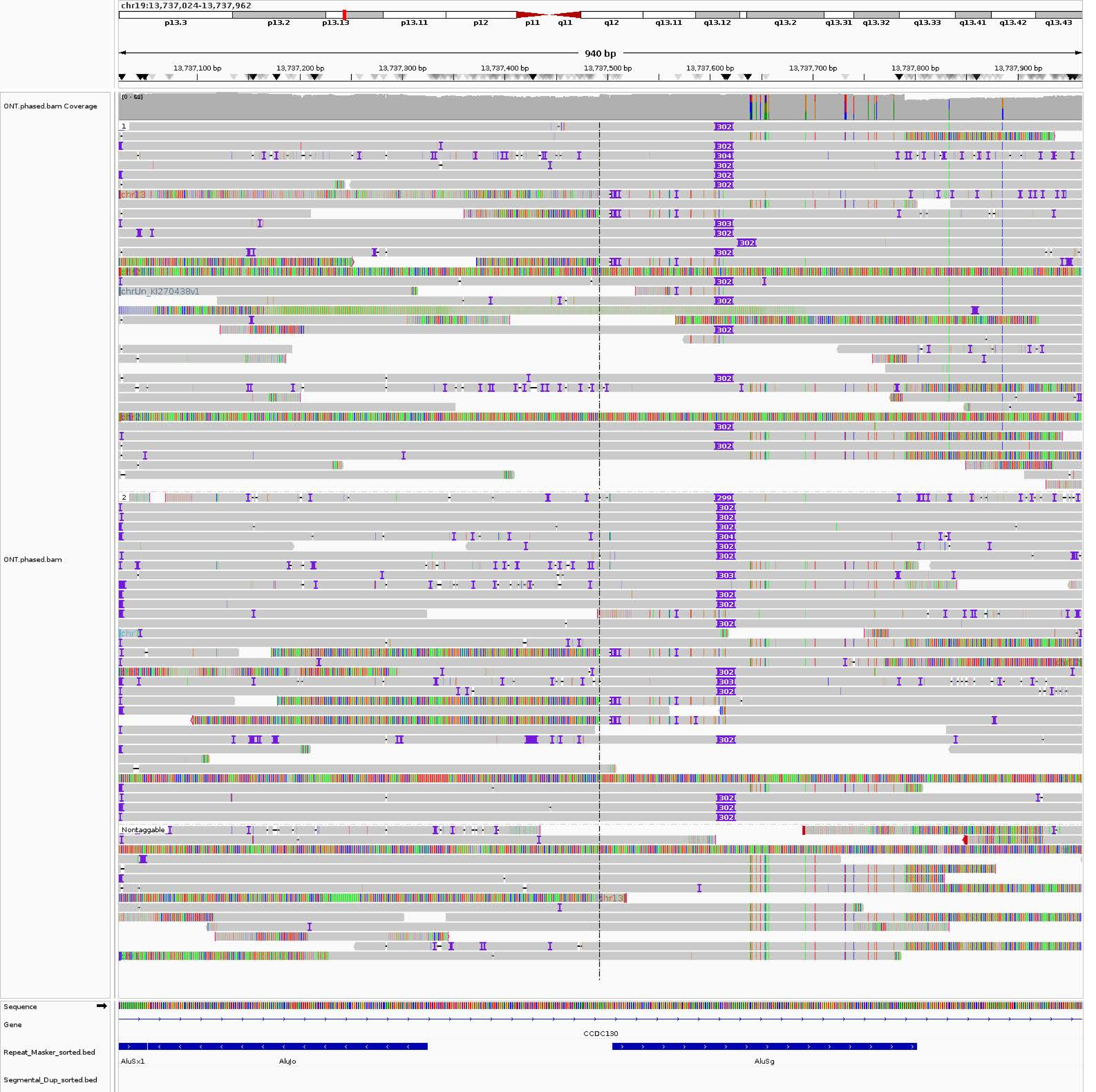
chr19:13,737,024-13,737,962
p13.3
p13.2
p13.13
p13.11
p12
p11
q11
q12
q13.11
q13.12
q13.2
q13.31
q13.32
q13.33
q13.41
q13.42
q13.43
940 bp
13,737,100 bp
13,737,200 bp
13,737,300 bp
13,737,400 bp
13,737,500 bp
13,737,600 bp
13,737,700 bp
13,737,800 bp
13,737,900 bp
ONT.phased.bam Coverage
[0 - 68]
ONT.phased.bam
chr13
chrUn_KI270438v1
Nontaggable
Sequence
Gene
CCDC130
Repeat_Masker_sorted.bed
AluSx1
AluJo
AluSg
Segmental_Dup_sorted.bed

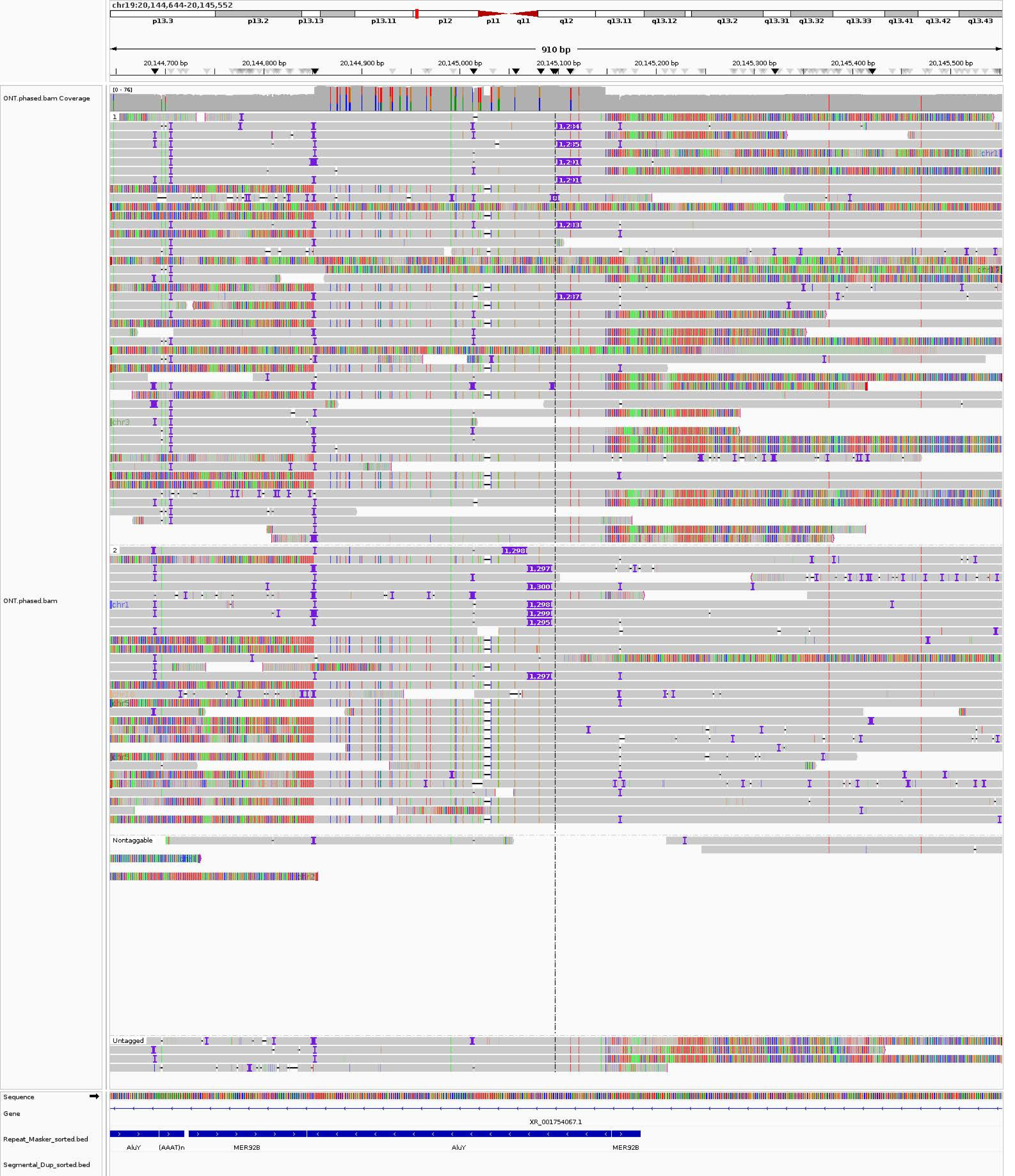

chr19:20,144,644-20,145,552
p13.3
p13.2
p13.13
p13.11
p12
p11
q11
q12
q13.11
q13.12
q13.2
q13.31
q13.32
q13.33
q13.41
q13.42
q13.43
910 bp
20,144,700 bp
20,144,800 bp
20,144,900 bp
20,145,000 bp
20,145,100 bp
20,145,200 bp
20,145,300 bp
20,145,400 bp
20,145,500 bp
ONT.phased.bam Coverage
[0 - 76]
ONT.phased.bam
1
2
Nontaggable
Untagged
Sequence
Gene
XR_001754067.1
Repeat_Masker_sorted.bed
AluY
(AAAT)n
MER92B
AluY
MER92B
Segmental_Dup_sorted.bed

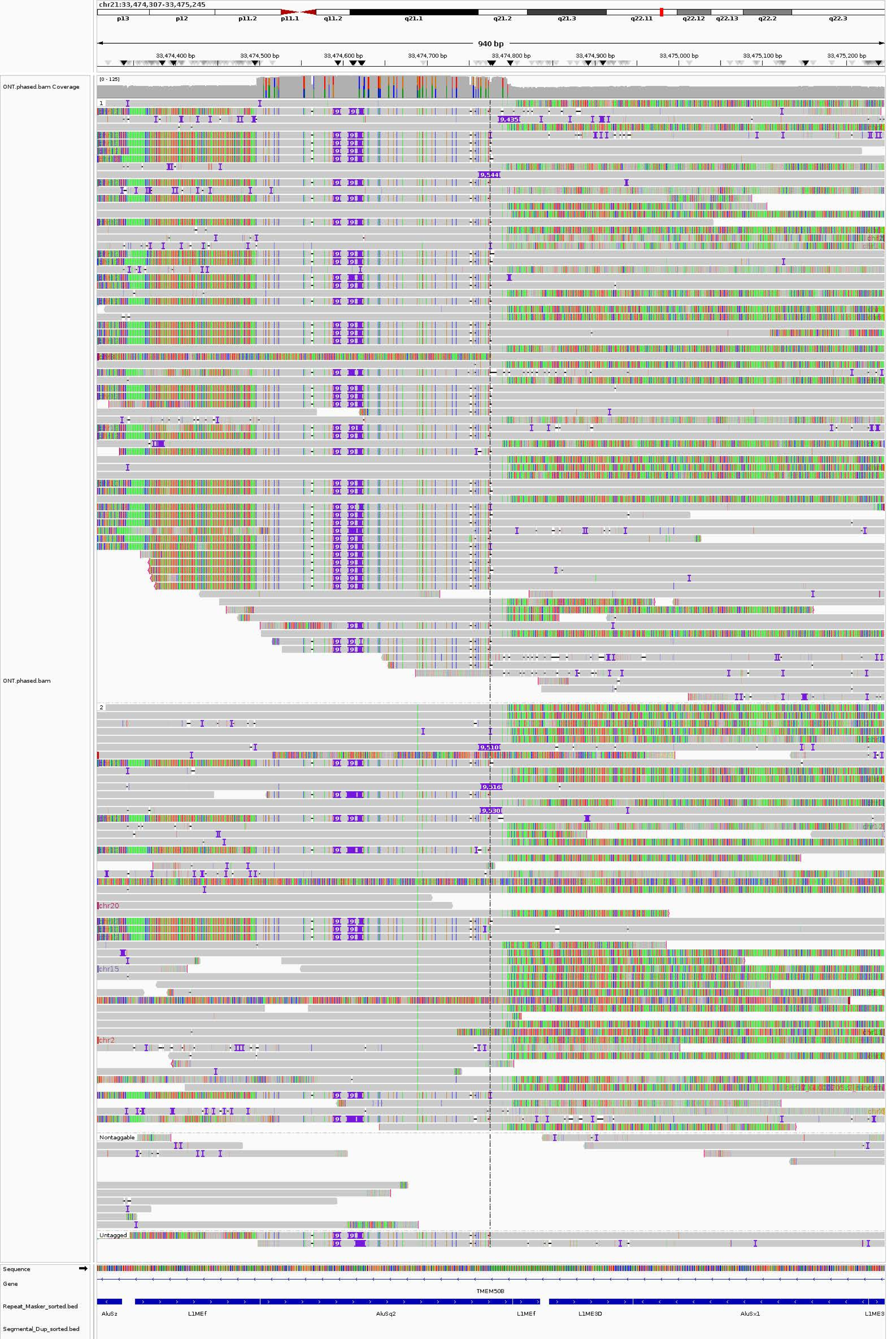
chr21:33,474,307-33,475,245
p13
p12
p11.2
p11.1
q11.2
q21.1
q21.2
q21.3
q22.11
q22.12
q22.13
q22.2
q22.3
940 bp
33,474,400 bp
33,474,500 bp
33,474,600 bp
33,474,700 bp
33,474,800 bp
33,474,900 bp
33,475,000 bp
33,475,100 bp
33,475,200 bp
ONT.phased.bam Coverage
[0 - 125]
ONT.phased.bam
1
2
chr20
chr15
chr2
Nontaggable
Untagged
Sequence
Gene
TMEM50B
Repeat_Masker_sorted.bed
AluSz
L1MEf
AluSq2
L1MEf
L1ME3D
AluSx1
L1ME3
Segmental_Dup_sorted.bed

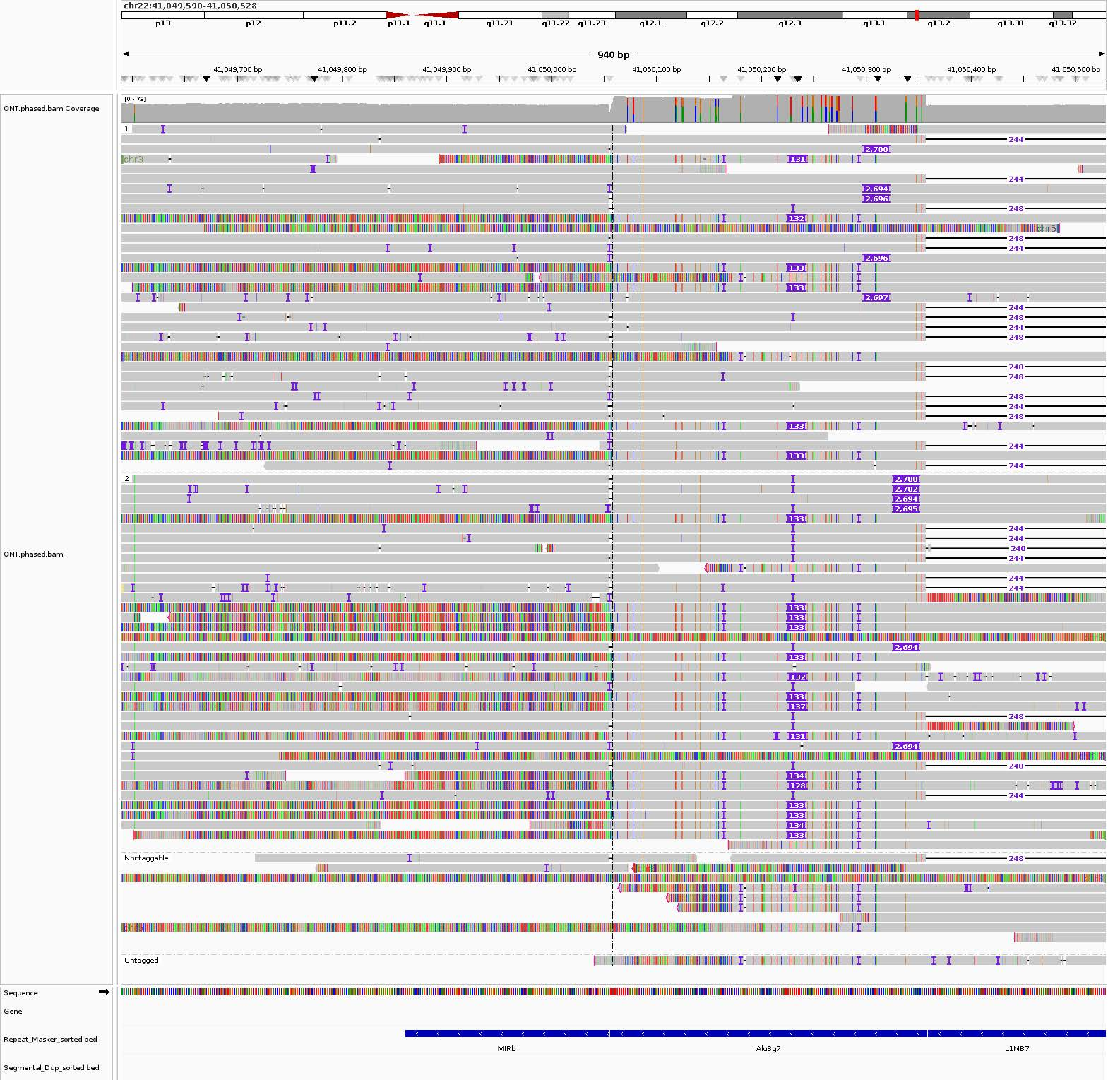
chr22:41,049,590-41,050,528
p13
p12
p11.2
p11.1
q11.1
q11.21
q11.22
q11.23
q12.1
q12.2
q12.3
q13.1
q13.2
q13.31
q13.32
940 bp
41,049,700 bp
41,049,800 bp
41,049,900 bp
41,050,000 bp
41,050,100 bp
41,050,200 bp
41,050,300 bp
41,050,400 bp
41,050,500 bp
ONT.phased.bam Coverage
[0 - 73]
ONT.phased.bam
1
2
Nontaggable
Untagged
chr3
chr5
Sequence
Gene
Repeat_Masker_sorted.bed
Segmental_Dup_sorted.bed
MIRb
AluSg7
L1MB7
