## Supplementary figures and images for "A personalized multi-platform assessment of somatic mosaicism in the human frontal cortex"

### Supplementary Figure 1.pdf

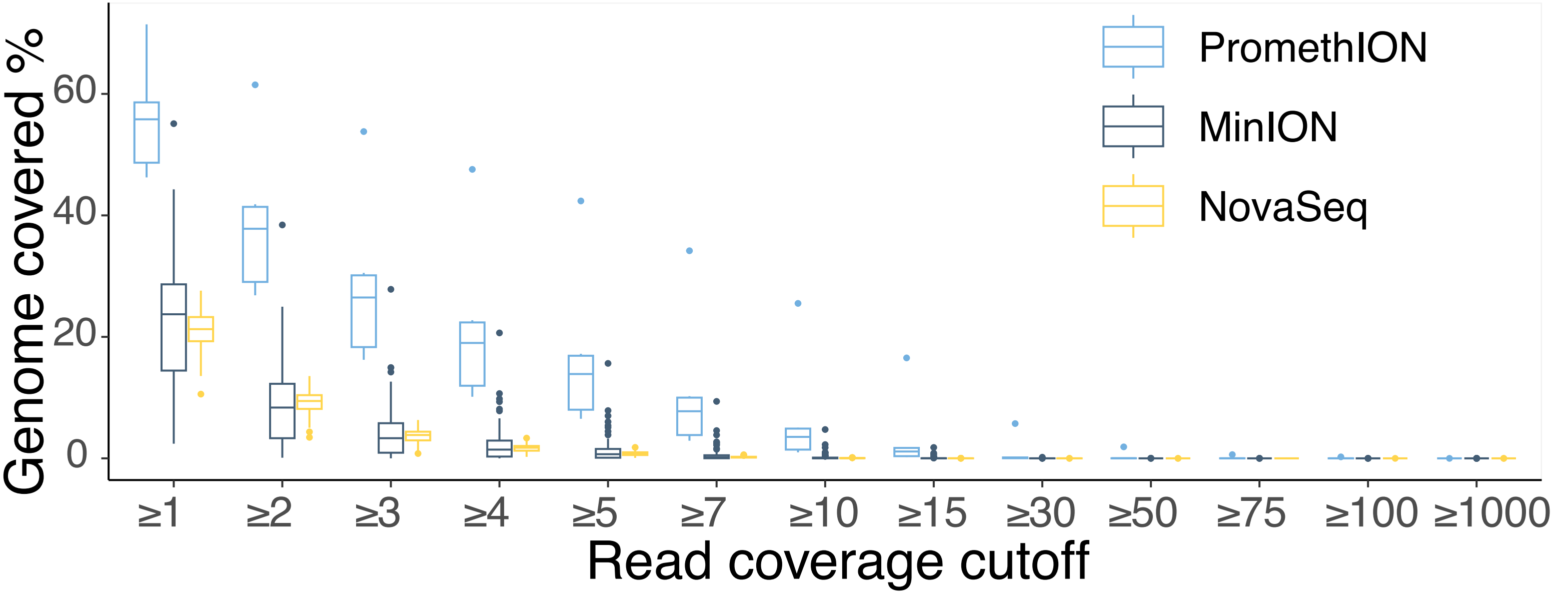

### Supplementary Figure 2.pdf

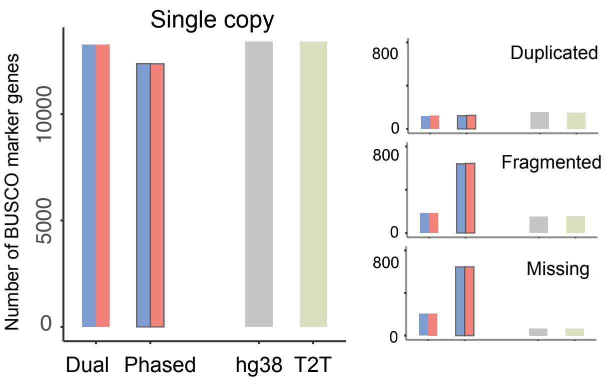

### Supplementary Figure 3.pdf

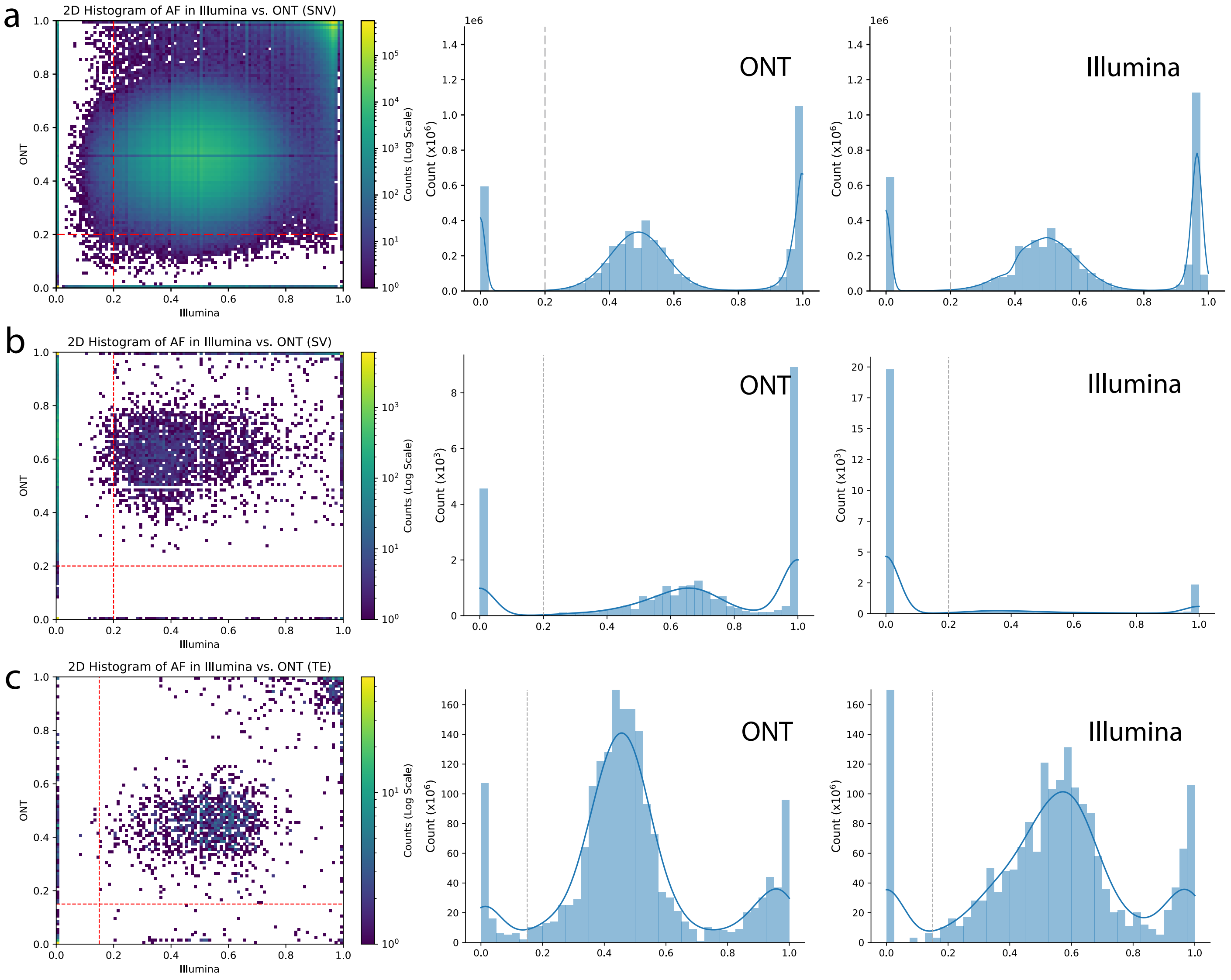

### Supplementary Figure 4.pdf

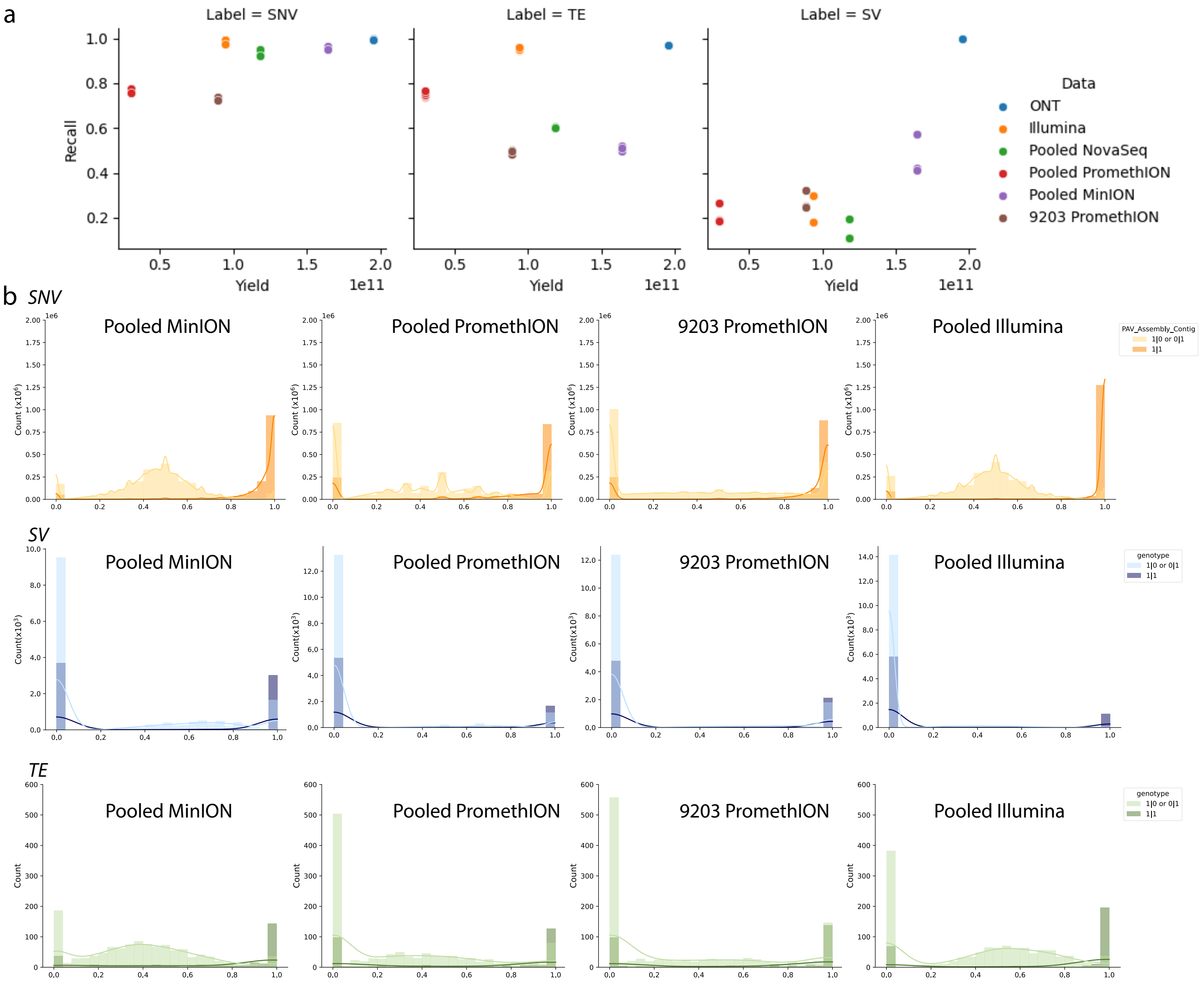

### Supplementary Figure 5.pdf

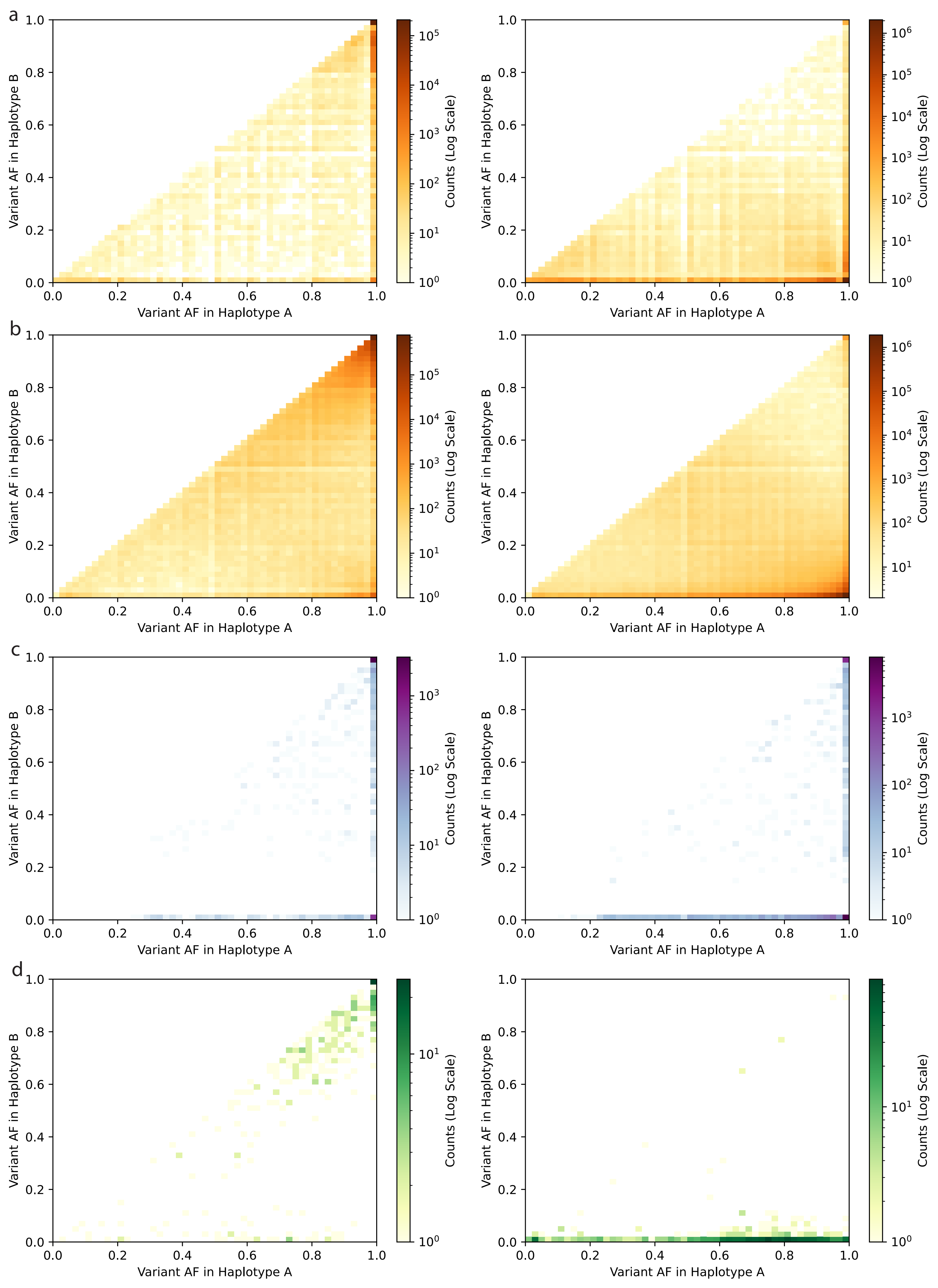

### Supplementary Figure 6.pdf

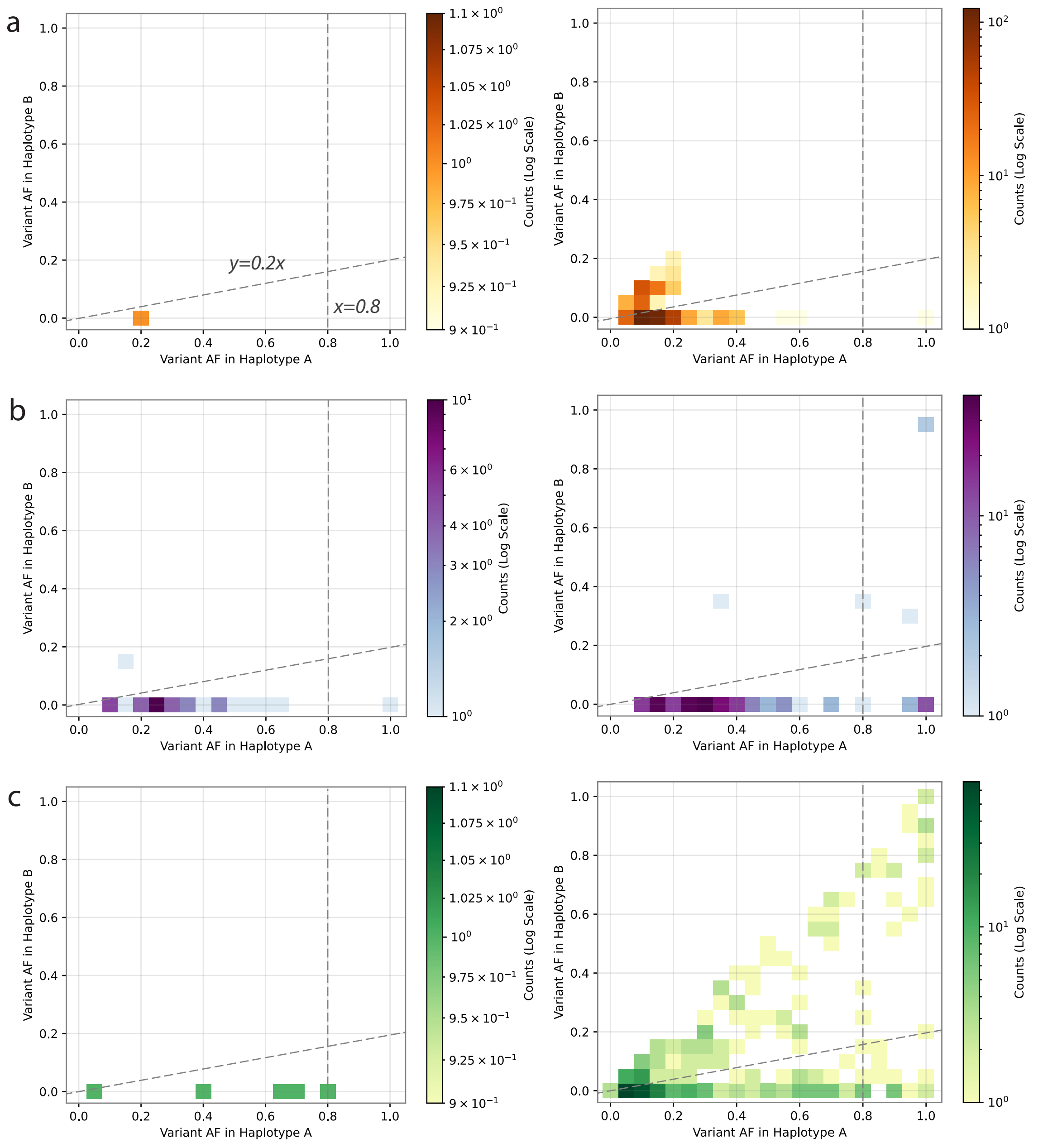

### Supplementary Figure 7.pdf

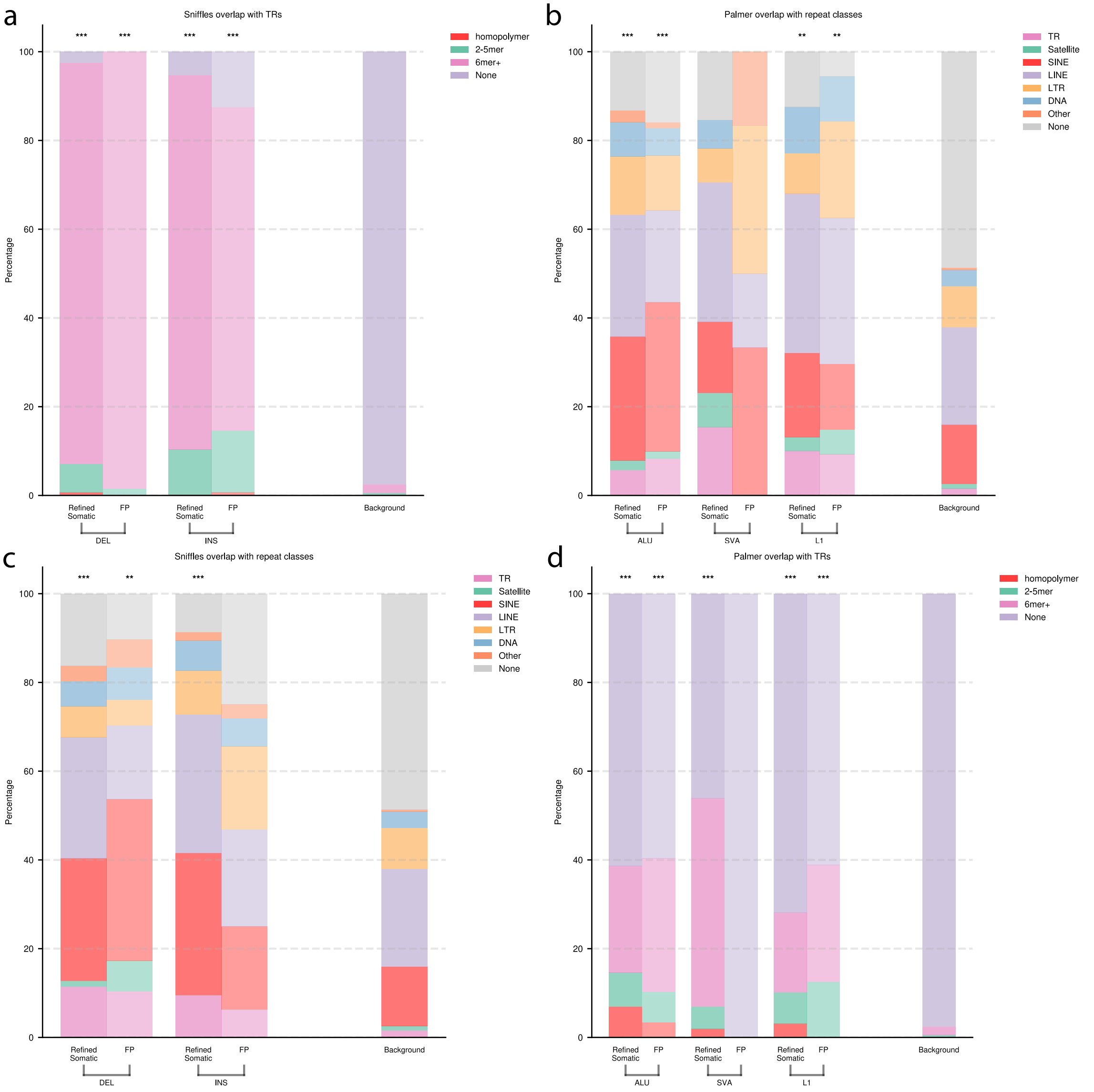

### Supplementary Figure 8.pdf

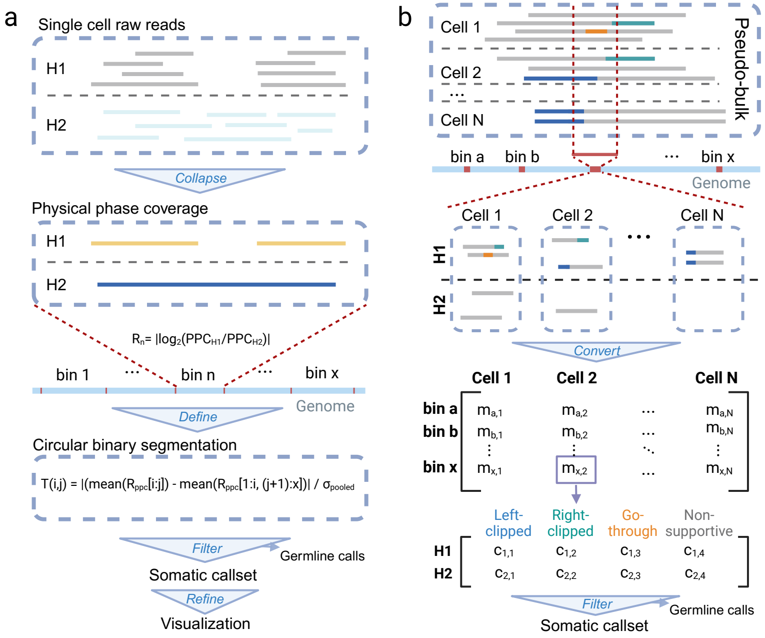
